# What is the correct genomic structure of the complex chromosomal rearrangement at the *Fm* locus in Silkie chicken?

**DOI:** 10.1101/2024.02.05.578760

**Authors:** Ashutosh Sharma, Nagarjun Vijay

**Affiliations:** Computational Evolutionary Genomics Lab, Department of Biological Sciences, IISER Bhopal, Bhauri, Madhya Pradesh, India

## Abstract

ARISING FROM Zhu, F., Yin, ZT., Zhao, QS. et al. Communications Biology https://doi.org/10.1038/s42003-023-05619-y (2023)

High-quality chromosome-level genome assemblies for numerous avian species promise to address longstanding questions in bird evolution and biology. In a recent issue of *Communications Biology*, Zhu, F., Yin, ZT., Zhao, QS. et al. (ZYZSJ)^1^ presented a chromosome-level assembly for the Silkie chicken using a multi-platform high-coverage dataset to obtain accurate and complete sequences spanning the entire genome. A key finding from their genomic analysis is the reconstruction of the structure of the complex rearrangement at the *Fm* locus, the primary genetic change underlying the rare and conspicuous dermal hyperpigmentation phenotype generally called Fibromelanosis. However, in contrast to their identification of the *\*Fm_1* scenario, several previously published studies^2–6^ claim that *\*Fm_2* is the valid scenario. Our re-analysis of ZYZSJ’s new assembly (CAU_Silkie) using long-read data from multiple black-bone chickens demonstrates that *\*Fm_2* is the correct scenario. The *\*Fm_1* scenario favoured by ZYZSJ results from an assembly error caused by mosaic haplotypes generated during the de novo assembly step. We recommend post-assembly validation and correction in genome projects to prevent misinterpretation due to assembly artefacts. Enhancing the assembly of haplotypes in such complex regions is essential for unravelling the genetic foundations of traits governed by genes within these areas.

## Introduction

The fibromelanosis phenotype, found in Silkie and other black-bone chicken breeds such as Yeonsan Ogye and Kadaknath, is caused by a complex chromosomal rearrangement (*Fm* locus) on chromosome 20 with a common origin in all black-bone chicken breeds^2–9^ (**Supplementary Text** and **Supplementary Figure S1-4**). However, the correct genomic structure of this rearrangement has been challenging to establish based on short-read sequencing, PCR and genetic analysis of crossing experiments alone^10^. Three possible hypothetical scenarios (*\*Fm_1, *Fm_2*, and *\*Fm_3*, see **Fig. 1**) have been proposed^3^ based on the identified rearrangement junctions. Genetic analysis of crosses between individuals with *Fm* and wild-type (*\*N*) has favoured the second scenario (*\*Fm_2*)^3,10^. However, subsequent studies utilising limited long-read sequencing for de novo assembly have failed to determine the correct scenario^11–13^ (**Supplementary Text**).

**Figure 1:**
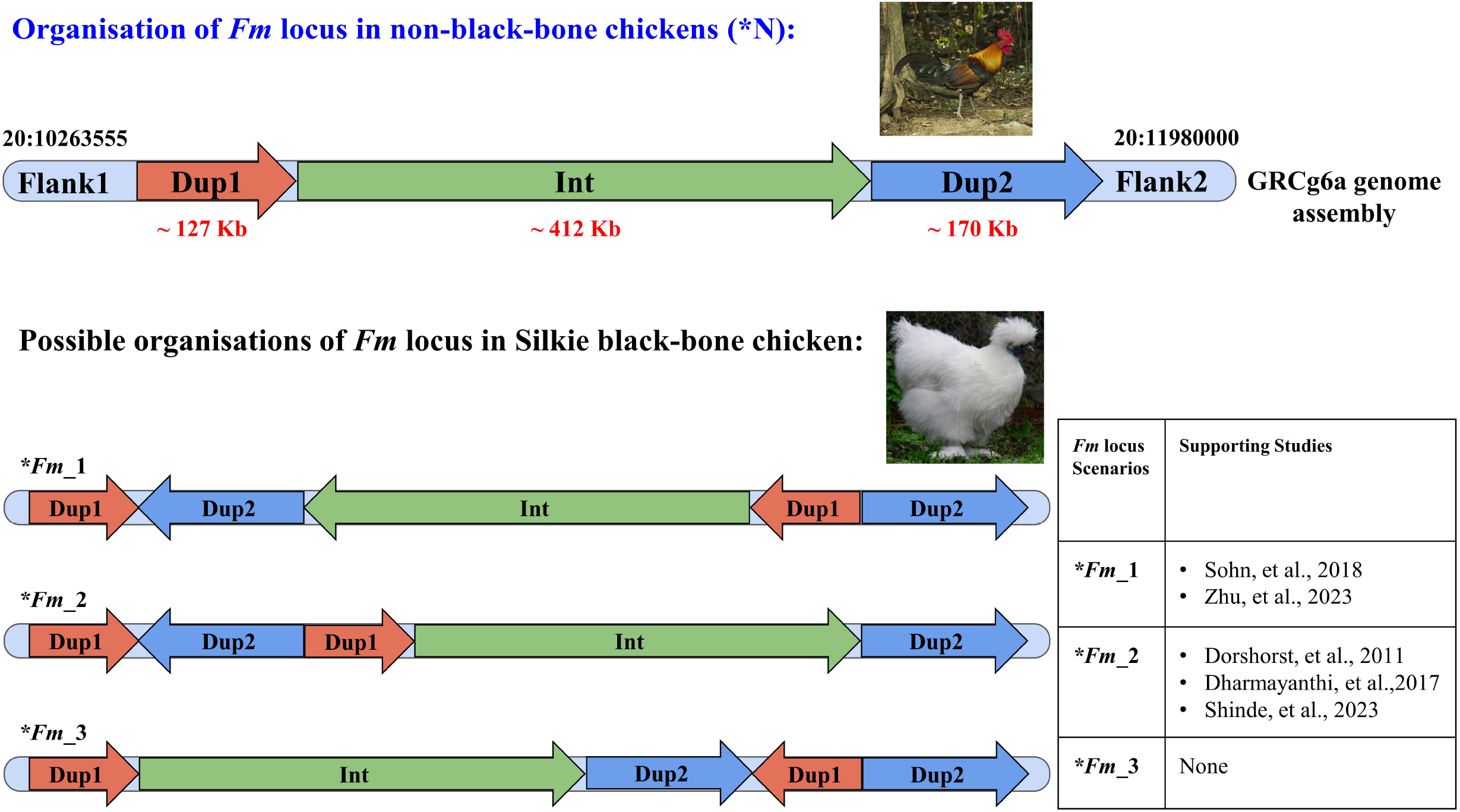
Organisation of *Fm* locus in Red junglefowl and Silkie black-bone chicken. The *Fm* locus located on chromosome 20 consists of two distinct non-paralogous regions designated as Duplication 1 (Dup1 ∼127 Kb) and Duplication 2 (Dup2 ∼170 Kb), separated by an intervening Intermediate (Int ∼412 Kb) region. The organisation of the *Fm* locus in non-BBC (*\*N*) neither involves duplication nor rearrangements. In black-bone chicken breeds, the *Fm* locus has undergone a complex rearrangement event involving duplications and inversions. Although the junctions between these regions have been identified based on short-read sequencing and PCR, the overall organisation of the *Fm* locus has been challenging to decipher. Three possible scenarios (*\*Fm_1, *Fm_2*, and *\*Fm_3*) representing the hypothetical organisation of the *Fm* locus have been proposed by Dorshorst et al., 2011 based on the junction sequences identified. Dorshorst and most subsequent studies have primarily supported the *\*Fm_2* scenario. However, some studies have favoured the *\*Fm_1* scenario.

## Results

ZYZSJ generated a de novo high-quality multi-platform chromosome-level genome (CAU_Silkie) from a single *Fm* homozygote (*Fm/Fm*) silkie individual. In ZYZSJ’s Figs. 1b and Supplementary Fig. 11-15, they identified *\*Fm_1* scenario (which the authors call FM2) as the arrangement at the *Fm* locus by relying upon this genome assembly. In a parallel study^5^, we opted for a haplotype phasing approach to resolve the structure of the *Fm* locus. In this approach, the silkie long-reads mapped to the GRCg6a genome were assigned to two haplotypes (**Fig. 2A**) corresponding to the two copies of the duplicated regions. The reads mapping at the boundaries of the duplicated region consists of two types: those that spanned the duplicated-unduplicated junctions and those that aligned in the duplicated region but were soft-clipped at the junctions (**Supplementary Figure S4a**). The genomic sequences contiguous with these two types of reads were distinguished based on the allelic states at haplotype-defining positions (HDPs). HDPs are sites that separate haplotype-consistent long-read tiling paths extending into the flanks of the duplicated regions. The read assignment step identified two haplotypes (**Fig. 2B**) spanning Dup1 and Dup2 regions and extending into the flanking regions. Step-wise extension of these haplotype-specific tiling paths identified *\*Fm_2* scenario as the correct arrangement of the *Fm* locus (**Fig. 2C**).

**Figure 2:**
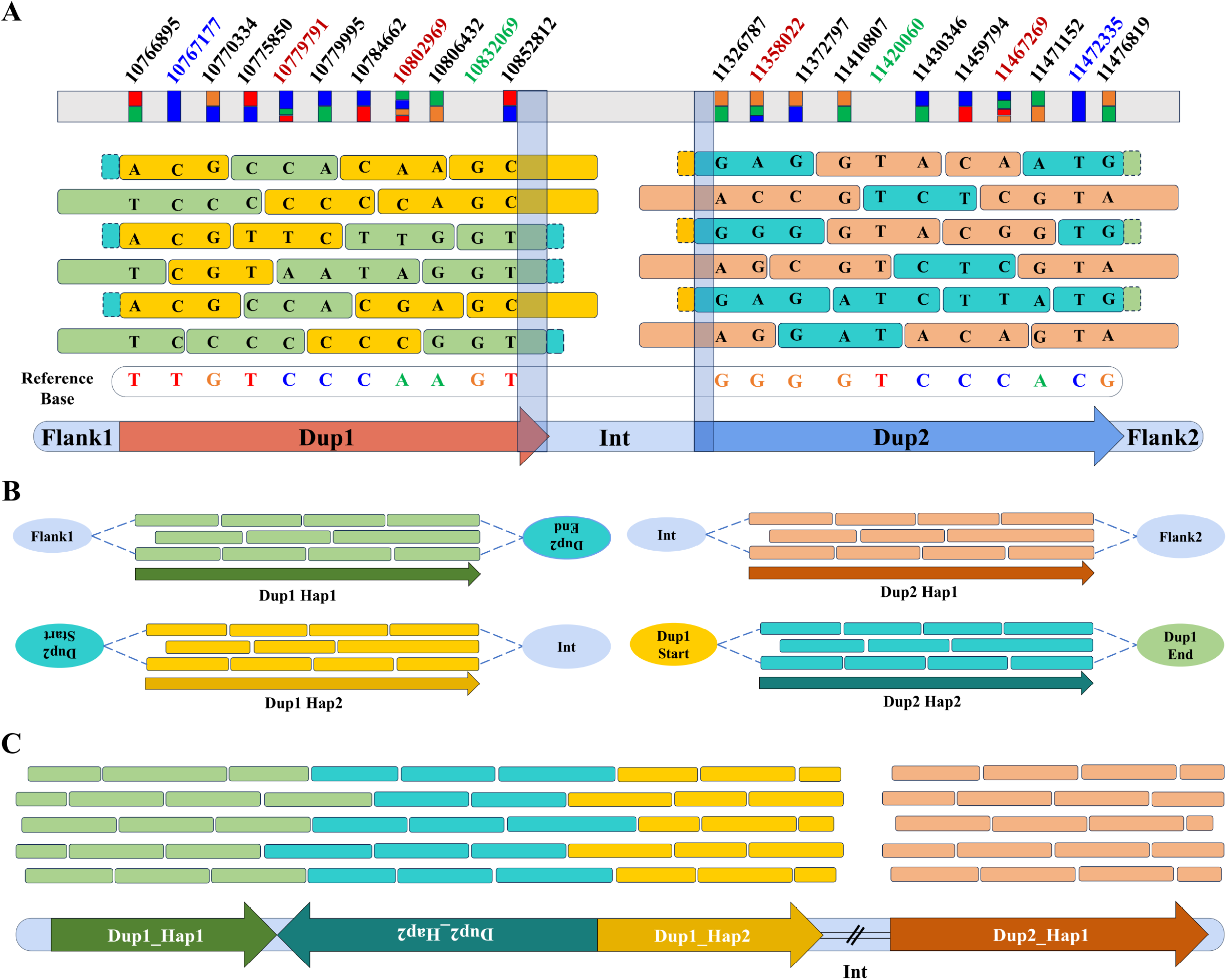
Graphical representation of haplotype-based phasing at the *Fm* locus in CAU Silkie using GRCg6a genome assembly. (**A**) Alignment of CAU_Silkie long-reads to the GRCg6a genome assembly at Dup1 and Dup2 regions: Reads spanning the junction between Flank1 and Dup1, and clipping at the end of Dup1 are displayed in light green. Reads beginning within Dup1 and spanning the Dup1-Int junction are shown in yellow. For the Dup2 region, reads spanning the junction between Flank2 and Dup2 and extending into Int are represented in light orange, while reads that are soft-clipped at the start and end of Dup2 are depicted in teal. Soft-clipped reads at the start of Dup1 (illustrated with dotted lines) are connected to the start of Dup2, and those soft-clipped at the end of Dup1 are connected to the end of Dup2. Nucleotide positions marked in black denote Haplotype Defining Positions (HDP) within the Dup1 and Dup2 regions, which distinguish the haplotypes. Light pink nucleotide positions indicate tri- and tetra-allelic sites. Blue marks homozygous positions in the CAU Silkie that differ from the GRCg6a genome assembly, while green marks homozygous positions in both CAU Silkie and GRCg6a. The vertical outlines represent the HDP-poor regions. (**B**) Resolution of Haplotypes: Based on the phasing data, the start and end points of each haplotype in Dup1 and Dup2 are identified. (**C**) Identification of *\*Fm_2* Scenario based on boundaries of each haplotype: The rearrangement of the *Fm* locus is illustrated based on the defined haplotype boundaries.

Despite utilising ZYZSJ’s long-read datasets (ONT and PacBio), our analysis consistently supported the *\*Fm_2* scenario (**Supplementary Text, Supplementary Figures S5-121** and **Supplementary Tables S1-19**). Our haplotype phasing approach excels in resolving the rearrangement at the *Fm* locus for several reasons: (1) we identify specific haplotype-defining positions (HDPs) spanning the entire length of the *Fm* locus, (2) we quantify long-read support for each pair of HDPs, and (3) we independently validate the results using ONT and PacBio long-reads. Our method’s intuitive nature and data visualisation at various steps led us to the correct (*\*Fm_2*) conclusion, which can be verified by examining the raw data (**Fig. 2**). We observed that the end of the Dup1 region and the start of the Dup2 region have insufficient HDPs (**Supplementary Figure S9a and S10a**). Therefore, we relied upon haplotype-specific reads that span these HDP-poor regions.

Evaluating long-read support for the CAU_Silkie assembly revealed multiple haplotype switching events (**Supplementary Figure S122**). The absence of raw read support for the published CAU_Silkie genome assembly suggests that the de novo assembly approach has failed to resolve the haplotypes. We observed that the allelic states either did not differ between haplotypes or were inconsistent with the long-read dataset (**Supplementary Text, Supplementary Figure S123-163** and **Supplementary Tables S20-27**). Hence, the CAU_Silkie assembly of the *Fm* locus consists of a mosaic haplotype. Due to a mosaic haplotype in the CAU_Silkie assembly, the previously identified 49 HDPs were insufficient for haplotype phasing. So, we identified 211 new HDPs while re-analysing the long-read data using CAU_Silkie as the reference. All 260 HDPs are concordant with the haplotypes we reconstructed earlier^5^, reinforcing our result that the *\*Fm_2* is the correct structure of the *Fm* locus (**Supplementary Table S20-27**).

## Discussion

Based on the evidence presented in our re-analysis and the findings from our previously published study^5^, we propose that consistent with early studies^3,10^, *\*Fm_2* represents the accurate arrangement at the *Fm* locus, with no support for scenarios *\*Fm_1* and *\*Fm_3*. Our evidence that *\*Fm_2* is the correct structure of the *Fm* locus will be vital to resolving future questions about gene regulation in this region. In a broader context, a black-box approach to genome assembly, lacking downstream validation, can introduce erroneous bases into genome assemblies, potentially impacting evolutionary genomic analysis^14^. Given the increasing capability of long-read sequencing methods to resolve complex structural variations, we advocate for caution when employing genome assembly tools. To enhance the reliability of results, we recommend thorough post-assembly evaluation and correction of the genomic sequence, utilising all raw data used in the assembly process and supplementing it with alternative sources of information such as population genetics and methylation-based phasing.

## Methods

A schematic description of HDP identification and haplotype-specific read partitioning using the GRCg6a genome is provided in **Supplementary Figure S120 & S121**. In our re-analysis, we scrutinised the CAU_Silkie assembly to assess whether the sequencing raw data substantiated the integrity of the genome assembly. Initially, we examined the allelic states at the 49 HDPs identified in our recent study^5^ within the CAU_Silkie assembly after the liftover of positions from GRCg6a. We could not find haplotype-consistent tiling paths based on these HDPs. Considering the possibility that the genome assembler employed a distinct set of HDPs for reconstructing the correct structure of the *Fm* locus, we identified 211 additional sites that differed between the haplotypes of the CAU_Silkie assembly. We adapted our read-backed phasing script to assess long-read support for each pair of these sites. The complete set of HDPs (previous 49 + newly identified 211 positions) consistently recovered the same haplotypes in mapped long reads irrespective of whether the GRCg6a or CAU_Silkie assembly was used.

## Acknowledgements

We thank the University Grants Commission for supporting AS with a Ph.D. scholarship. The Department of Biotechnology, Ministry of Science and Technology, India (Grant no. BT/11/IYBA/2018/03) and Science and Engineering Research Board (Grant no. ECR/2017/001430) provided funds for computational resources (i.e., Har Gobind Khorana Computational Biology cluster) used.

## Author contributions

Ashutosh Sharma: Conceptualisation, Formal analysis, Investigation, Visualisation, Validation, Writing - original draft, Writing - review & editing. Nagarjun Vijay: Conceptualization, Resources, Writing - original draft, Writing - review & editing, Funding acquisition, Project administration, Supervision.

## Data Availability

Scripts and data are available at https://github.com/Ashu2195/Fm_locus.

