## Supplementary Text for "What is the correct genomic structure of the complex chromosomal rearrangement at the *Fm* locus in Silkie chicken?"

ARISING FROM Zhu, F., Yin, ZT., Zhao, QS. et al. Communications Biology
https://doi.org/10.1038/s42003-023-05619-y (2023)

### 1. Common origin of *Fm* locus

The genome assembly of one Silkie individual is used to resolve the *Fm* locus in the ZYZSJ 2023 paper. However, several breeds of black-bone chicken are described in different parts of the world. Since the focus of the ZYZSJ 2023 paper has been the Silkie chicken, they have not considered other black-bone chicken breeds. Moreover, the possibility of different individuals within the Silkie breed having different arrangements at the *Fm* locus has not been explored by ZYZSJ 2023. Generating high-quality chromosome-level assemblies for hundreds of individuals from different breeds remains prohibitively expensive. Moreover, the assembly of the *Fm* locus region in each of these individuals will be challenging due to the complex nature of this region.

In this context, we relied upon complementary sources of information to explore whether the *Fm* locus arrangement is shared across individuals of Silkie and those of other black-bone breeds. Apart from understanding the history of the black-bone chicken breeds, evaluating the origins of the *Fm* locus will allow the integration and comparison of assemblies of different black-bone breeds generated by independent groups worldwide. This approach allows us to evaluate the organization of the *Fm* locus across large numbers of individuals without generating high-quality genome assemblies for such a large sample size. The study of diverse black-bone chicken breeds has already suggested a "Single origin of the *Fm* phenotype"[1]. Based on sequence divergence in the Dup1 and Dup2 regions, Dharmayanthi 2017 suggests "a rough estimate of divergence time (0.6–0.8 myr) implies that these EDN3 alleles in fact originated in the ancestral RJF population". We identified ~260 HDPs (Haplotype-Defining Positions) in the ~300 Kb region, which is probably underestimated. Our goal was not to identify all possible HDPs but to find enough HDPs to perform haplotype phasing. Nonetheless, we find approximately 1 HDP per Kb. These numbers are comparable to those Dharmayanthi (2017) reported and support an ancient origin of the *Fm* locus. Such convincing evidence of a common origin for the *Fm* locus allows the integration of data from other black-bone chicken breeds to resolve the structure of this complex genomic region. We cannot help but wonder about the implications of such a common and ancient origin of the *Fm* locus in understanding chicken domestication. Another possible explanation for the divergence between the two copies of the duplicated regions is differential gene flow due to the limited recombination of one copy with wild-type (*N**) in contrast to a higher recombination rate at the other.

Approaches to using local ancestry information to investigate the "common origin for gamecocks from around the world" have been highly successful[2]. We rely upon local ancestry inference through principal component analysis (PCA) and phylogenetic trees to reconstruct the evolutionary relationship among black-bone and non-black-bone breeds at the *Fm* locus. To obtain a comprehensive picture, we included 139 chickens, including 61 black-bone chickens (13 black-bone breeds) (see **Supplementary Table S1**). Our local PCA of the Dup1 and Dup2 regions separates all black-bone from non-black-bone chicken (see **Supplementary Figures S1 and S2**). The local phylogenies of these regions also clearly distinguish black-bone from non-black-bone chickens, irrespective of the breed (see **Supplementary Figures S3 and S4**). These results provide an independent confirmation of a common origin of black-bone specific *Fm* locus.

In our recent study of Kadaknath[3] and other black-bone chicken breeds, we used local PCA, phylogeny, admixture, and pairwise population genetic parameters like $F_{ST}$ to establish a common origin for the *Fm* locus across black-bone breeds. Additionally, the junction sequences are the same (https://www.frontiersin.org/files/Articles/1180658/fgene-14-1180658-HTML/image_m/fgene-14-1180658-g007.jpg) in all the black-bone breeds, and the remarkable two-fold increase in short-read coverage at these precise locations (at the base-pair level) further supports the shared junctions across the breeds.

### 2. Earlier published studies of black-bone chicken

The genomic organization of the *Fm* locus was explored in the Silkie chicken more than a decade ago[4–6]. Subsequent studies of Silkie and other black-bone chicken breeds have focused on understanding genomic history, copy number variation and diversity patterns at the *Fm* locus[1,7–16]. The hyperpigmentation in Silkie chicken affects other traits, such as altered immune development[17,18]. A recent study of Xichuan black-bone chicken[19] reported finding "an inversion at 10.97–11.52 Mb and a duplication at 10.74–12.55 Mb" overlapping the *Fm* locus. The detailed study of the *Fm* locus has made it a prominent example that has been used as a textbook example of a striking phenotype associated with copy number variation[20,21]. A recent slew of genomic studies has generated a wealth of genomic data for diverse black-bone chicken breeds.

1. The earlier genomes of Silkie2 (GCA_024679325.1) and Silkie3 (GCA_024653025.1) reporting the De Novo Assembly of 20 Chicken Genomes in the journal ***Molecular Biology and Evolution***, 2022[22] did not manage to assemble the *Fm* locus completely and resolve its correct structure. The Silkie3 individual was sequenced at relatively high coverage (i.e., SMRT (~91x)/NGS (~65x)/Hi-C (~113x)) using multiple-sequencing strategies. Silkie2 was sequenced using only ~134x NGS data.
2. The genome assembly of Yeonsan Ogye (GCA_002798355.1/ Ogye1.0) generated by Sohn et al. (2018) and published in ***Gigascience***[23] does not complete this region's assembly with low-depth (~9.7x coverage) PacBio data. The authors state, "In this study, the draft genome of YO was assembled using a hybrid de novo assembly method that takes advantage of high-depth Illumina short reads (376.6X) and low-depth Pacific Biosciences (PacBio) long reads (9.7X)." Despite concerted efforts, the correct scenario could not be identified among the three possible scenarios. However, this study supported the **Fm_1* scenario as this requires the least number of rearrangement events from wild-type.
3. The genome assembly of Silkie (GCA_033088195.1) by CAU (i.e., CAU_Silkie_1.0) generated in the ZYZSJ study is the first de novo assembly-based study to claim resolution of the *Fm* locus based on de novo assembly and supports the **Fm_1* scenario. The study was published in ***Communications Biology*** [24] and concluded, "In our case, the high-quality Silkie chicken assembly solves FM traits which had not been investigated by large-scale

chicken pan-genome assemblies, especially based on the second-generation whole-genome sequencing data, and insufficient third-generation data.".

The advances in sequencing technology and the associated genome assembly tools have made near-perfect assemblies a reality for many organisms, including those with gigabase-sized, repeat-rich genomes. It is worth noting that the attributes that are aspired from high-quality reference genomes, such as those being generated by the vertebrate genome project, are:

1. Be complete. i.e., without any gaps
2. Accurate. i.e., base calls and structure should be correct and error-free
3. Haplotype resolved in all regions, including repeats and other complex regions
4. Representative of the species or breed and not just a single individual

Yet, the complex, hard-to-sequence, and hard-to-assemble regions remain challenging for automated pipelines and need special attention. Identifying and resolving miss-assemblies in de novo assembled genomes has received considerable attention since the early days of genomics, as demonstrated by tools like AMOSValidate[25], which used the ACE file format (output by various assembly programs) to validate genome assemblies. Recent tools like REAPR[26] have used post-assembly read mapping to validate genome assembly. The Flagger[27] read-based assembly evaluation pipeline highlights the relevance of haplotype-resolved assembly validation in telomere-to-telomere genome assembly.

### 3. <u>Recent genome assemblies available on NCBI</u>

Much more recently, as part of BioProject# PRJNA1000217, the genomes of Silkie, Hailanhe and Lueyang breed chickens have been sequenced using PacBio Sequel and assembled using hifiasm v. 0.19.4-r575. These assemblies were posted to NCBI on Dec 21, 2023.

1. The GCA_034509865.1 (Silkie genome from Northwest A&F University) assembly has partially assembled the *Fm* locus with scaffold JAVDCC010000086.1 assembling the *N** arrangement (Flank1-Dup1-Int-Dup2-Flank2) and the scaffold JAVDCC010000529.1 assembling the Dup1-Dup2-Dup1 regions. The partial assembly along the scaffold JAVDCC010000529.1 is inconsistent with the **Fm_1* scenario favoured by Zhu et al. and instead supports the **Fm_2* scenario. The independent Silkie PacBio dataset generated and assembled by Northwest A&F University, China, further demonstrates the challenge posed

by the *Fm* locus to automated de novo assembly of this region. The data reported for this assembly has a limited (~22x) genome coverage.

2. The de novo assembly of the Lueyang breed chicken (GCA_034509885.1), another black-bone breed from China, has a complete assembly of the *Fm* locus. Interestingly, this assembly has reconstructed the **Fm_2* scenario using a de novo assembly approach. (See alignment of Lueyang assembly with GRCg6a: **Supplementary Figure S5**). The data reported for this assembly has an intermediate (~41x) genome coverage.

### 4. Haplotype-phased denovo assembly

First, we obtained all ONT reads corresponding to the *Fm* locus region to perform a local re-assembly. For a detailed comparison of the long-read assembly programs for overcoming uncollapsed haplotypes, we refer the reader to the recent study by [28]. An important result from this study is that "The assemblers we tested (Canu, Flye, NextDenovo, Ra, Raven, Shasta and wtdbg2) exhibited strikingly different behaviors when dealing with highly heterozygous regions, resulting in variable amounts of uncollapsed haplotypes".

**Shasta assemblies**:

We tried assembling the ONT reads from the *Fm* locus in the default mode of Shasta using all the reads from the *Fm* locus, only those from Dup1, only those from Dup2, only those from Dup1 and Dup2, and only those from Dup1, Dup2 and Int. The assembly graphs of these assemblies demonstrate the challenge in resolving the *Fm* locus with the default mode of Shasta, as both haplotypes of Dup1 and Dup2 occur in a single collapsed edge with twice the coverage of the flanking regions (see **Supplementary Figure S6-8**).

Separating the haplotypes of Dup1 and Dup2 is essential to overcome this limitation. Therefore, in addition to the default assembly mode, we tried assembly mode 2 with the config file Nanopore-UL-Phased-Nov2022. The assemblies generated using assembly mode 2 provide a "New implementation of phased assembly (Mode 2) improves the quality of phased assemblies with fewer artifacts".

The resulting assemblies were inspected in Bandage[29] to evaluate the assembly quality and the ability of the assemblers to separate the haplotypes of Dup1 and Dup2. We found that assembly mode 2 could separate the haplotypes over a majority of the length of the sequence (**Supplementary Figures S9 and S10**). However, similar to the challenges the mapping-based phasing method faces, the results demonstrate the difficulty of separating the haplotypes in the HDP-poor regions (**Supplementary Figure S9a and S10a**).

To compare de novo assembly results with those of read-backed phasing, we mapped the sequences of the edges assembled by Shasta in mode2 to the GRCg6a genome assembly. We found that sequences of the de novo assembled edges (from mode 2 of Shasta) differed from each other at the HDPs and were consistent with the haplotype-specific allelic states we identified based on read-backed phasing. In the GRCg6a genome, the ~127Kb long Dup1 region starts at 10766772 and ends at 10894151.

The two edges assembled in the Dup1 region are ~111Kb long and do not span the ends of Dup1. The node PR.0.1.0.1 is 1,11,782 base pair long and starts aligning to the genome at 10766895 and stops aligning at 10878704. Similarly, the node PR.0.1.0.0 is 1,11,772 base pair long and starts aligning to the genome at 10766896 and stops aligning at 10878703. The alignment of these node sequences starts precisely at the first HDP (10766895) in Dup1 and extends ~25Kb beyond the last HDP (10852812) identified in our read-backed phasing approach (**Supplementary Figure S11-13**). Upon carefully comparing the de novo assembled sequences, we found one previously unidentified HDP at 10852841 supported by the long-read data (**Supplementary Figure S14**). Another major difference between the sequences of the two edges was seen towards the end of Dup1 at position 10874946 (**Supplementary Figure S15**). However, the raw reads do not support this difference between the de novo assembled edge sequences. On the other hand, at several HDPs, the edges have the same base in both sequences, while the raw read supports the two haplotypes (**Supplementary Figure S16 and S17**). These analyses of Shasta assembled Dup1 sequences further demonstrate the challenge of assembling the complex region spanning the *Fm* locus.

The two edges assembled in the Dup2 region are ~135Kb long and do not span the entirety of Dup2, which is ~170Kb in length (**Supplementary Figures S18 and S19**). The node PR.0.1.0.1 is 1,35,088 base pair long and starts aligning to the genome at 11332861 and stops aligning at

11467885. Similarly, the node PR.0.1.0.0 is 1,35,030 base pair long and starts aligning to the genome at 11332860 and stops aligning at 11467886 (**Supplementary Figure S20 and S21**). The alignment of these node sequences starts at 11332861, a previously unidentified putative HDP and occurs after the second HDP identified in the Dup2 region. The third HDP in the Dup2 region also differs between the two assembled edges (**Supplementary Figure S21a**). The Dup2 sequence edges align up to 11467885 and fall short of the last HDP at 11476819 by ~9Kb. Similar to the Dup1 region at several HDPs, the Dup2 sequence edges have the same base in both sequences, while the raw read supports the two haplotypes (**Supplementary Figure S22-24**). Overall, neither Dup1 nor Dup2 haplotypes could be resolved entirely using the de novo approach using Shasta mode 2. The availability of even higher coverage ultra-long reads and the prevalence of more HDPs in this region could allow automated haplotype-resolved de novo assembly of these regions.

**Raven Assembler:**

Our attempts at de novo assembly of the *Fm* locus region with the Raven assembler resulted in entangled graphs that could not separate the two haplotypes. The assembler differences have already been established in previous studies[28]. The assembly graphs generated by Raven for different regions of the *Fm* locus are provided in (**Supplementary Figure S25-29**).

### 5. Motivation for using a mapping-based approach to phasing

Given the contradictory assemblies (CAU_Silkie, Lueyang hifiasm assembly, our Shasta-based haplotype-resolved assemblies of Silkie) of the *Fm* region and entangled graphs generated by de novo assembly methods, the de novo approach poses a perplexing conundrum. Therefore, a de novo assembly-based approach will never be able to provide a conclusive answer if the assembly quality cannot be validated. Robust studies of segmental duplications in humans[30] have addressed potential haplotype-phasing errors using parent-child trios (i.e., locally phased using trio-binning) and cell lines with a single paternal haplotype. Our mapping-based approach to phasing can be validated by inspecting the long-read support at each HDP.

Irrespective of which method (de novo or reference-based) is used, the underlying data (PacBio & ONT) remains the same. The structural variation resolution method must rely upon sequence-level differences between the two copies of Dup1 and Dup2. The two copies of Dup1 are ~99.878% similar, and those of Dup2 are ~99.939% similar (**Supplementary Figure S30**). In addition to the high similarity between the copies within Silkie, the Dup1 and Dup2 regions of Silkie are highly similar (>99% similar) to the GRCg6a reference (**Supplementary Figures S31 and S32**). Given such high sequence similarity levels, the reference bias issue is expected to be minimal. Reference bias or genome mapping bias occurs when reads that contain non-reference alleles fail to align with their true point of origin, leading to inaccurate results.

### 6. Investigation of Reference bias at the *Fm* locus

Reference bias can be identified as regions with lower mapping quality scores or lower coverage in the reads containing the non-reference bases. We evaluated if any of these reference bias-indicating characters could be seen in the *Fm* locus.

**Read coverage-based evaluation of reference bias**:

GRCg6a reference:

Read coverage of long-read (both ONT and PacBio) mapped to the *Fm* locus of GRCg6a is consistent and has a sharp increase in coverage at the duplicated (Dup1 and Dup2) regions (**Supplementary Figure S33 and 34**). Similarly, the read coverage of one Silkie individual short-read (SRA Run ID # SRR17968701) has a similar pattern of sequencing coverage. The two-fold increase in coverage at the Dup1 and Dup2 regions is consistent with both haplotypes mapping to the GRCg6a reference genome without any reference bias. This coverage pattern is consistent regardless of the read mapper (bwa or minimap2) and method used to calculate coverage (bedtools coverage or MosDepth) (**Supplementary Figures S35 and S36**).

Read coverage for reads assigned to each haplotype and those not assigned to any haplotype were calculated for ONT and PacBio reads mapped using bwa and minimap2. While the total coverage is comparable across the duplicated regions, the coverage of the reads assigned to specific haplotypes drops at the Dup1-end and Dup2-start. Concordant with this difference in coverage between haplotype-assigned reads and total coverage, the coverage of unassigned reads increases at the end of Dup1 and Dup2 start. The drop in coverage in these regions among haplotype-assigned reads results from a lack of HDPs in this region. The haplotype-specific coverage demonstrates that read mapping to the GRCg6a reference is not affected by reference bias, as both haplotypes map equally well irrespective of the read mapper (bwa or minimap2) and method used to calculate coverage (bedtools coverage or MosDepth).

To quantify the mapping location of pairs of the reads that map at HDPs and contain haplotype-specific bases, we analyzed the HI-C data from the CAU Silkie (**Supplementary Figure S36a**). The pairs of reads of Dup1 Hap1 show an increase in coverage at the Flank1 region, while the pairs of reads of Dup1 Hap2 show an increase in coverage at the Dup2 start region. The coverage patterns of read pairs suggest that Dup1 Hap1 is adjacent to Flank1, and Dup1 Hap2 is positioned beside the Dup2 start. Similarly, the read pairs of Dup2 Hap1 exhibit increased coverage at the Flank2 and Int regions, whereas the read pairs of Dup2 Hap2 show increased coverage at the Dup1 region. This coverage pattern indicates that Dup2 Hap1 is between Flank2 and Int, while Dup2 Hap2 is between Dup1 Hap1 and Dup1 Hap2. These findings further support the **Fm_2* scenario.

<u>CAU_Silkie reference</u>:

Read coverage of long-read (both ONT and PacBio) and short-read mapped to the *Fm* locus of CAU_silkie is consistent and has no abrupt increase in coverage at the duplicated (Dup1 and Dup2) regions (**Supplementary Figure S37** and **S38**) as both copies are assembled separately. The total coverage of long and short-read data along the Dup1 and Dup2 regions does not increase compared to the Int or Flank regions in CAU_Silkie. It suggests that reads from only one of the haplotypes map to the genome at any given position. This uniform coverage across the *Fm* locus is expected for haplotype-specific assembly. However, upon separating the reads based on the haplotypes, we found that the start of Dup1 (see **panels A and B**) has a higher coverage of hap1 reads than hap2 reads. In contrast, the HDP-poor region near the end of Dup1 has a higher coverage of hap2 reads than hap1 reads. Notably, the Dup1r copy has a higher coverage of hap2 reads at the Dup1r-Dup2 junction than the hap1 reads. The hap1 read coverage is higher than the hap2 read coverage at the HDP-poor region near the Int-Dup1r junction.

Even in the Dup2 (see **panels C and D**) region, we see a trend similar to Dup1 with a change in haplotype-specific coverage in the HDP-poor region. The end of the Dup2 region has higher coverage for hap1 reads than hap2 reads. This trend reverses to a higher coverage of hap2 reads than hap1 reads in the HDP-poor Dup2 start region. Similarly, the coverage of hap1 reads is higher than hap2 reads near the Dup2r-Int junction. This trend reverses towards the Dup2r-Dup1 junction with higher coverage for hap2 reads than hap1 reads. This switch in haplotype-specific coverage along the length of Dup1 and Dup2 already hints at the possibility of mosaic assembly due to haplotype-switching in the HDP-poor regions.

In contrast to the GRCg6a reference, the CAU_silkie genome has clear signatures of reference mapping bias.

Similar to the GRCg6a analysis, we examined the CAU Silkie HI-C data with the CAU Silkie genome assembly (**Supplementary Figure S38a**). The read pairs of Dup1 Hap1 show an increase in coverage at the Flank1 region, while the read pairs of Dup1 Hap2 show an increase in coverage at the start regions of Dup2r and Dup2. Interestingly, despite Dup1r being located near Dup2 in the CAU Silkie genome, there is increased coverage in Dup2r, which is unexpected. Additionally, the read pairs of Dup2 Hap1 show increased coverage at the Flank2 and Int regions compared to Dup1r. Since Dup1r is much closer to Dup2 than the Int region, this finding is again inconsistent with the genome assembly. Moreover, the read pairs of Dup2 Hap2 show an increase in coverage at both the Dup1 and Dup1r regions at the same level, even though Dup1r is much farther away from Dup2r. This inconsistent read pair coverage in the CAU Silkie genome assembly further rejects the **Fm_1* scenario and supports the **Fm_2* scenario.

**Mapping quality-based evaluation of reference bias:**

Read mapping quality indicates the accuracy with which reads are mapped to a specific genomic location. Our evaluation of the mean mapping quality (in 100 base pair windows) along the GRCg6a reference found mapping qualities close to 60 throughout the length of the *Fm* locus (see **Supplementary Figure S39**). Our previous analysis of haplotype-specific coverage has demonstrated that reads from both haplotypes are mapped to the GRCg6a reference in comparable numbers. The evaluation of mean mapping quality rejects the possibility of reference bias as reads from both haplotypes are mapped with comparable mapping quality.

The CAU_silkie genome has two copies of Dup1 and Dup2 assembled along the *Fm* locus, making accurate read mapping challenging. Our evaluation of the mean mapping quality (in 100 base pair windows) along the CAU_silkie reference found mapping qualities close to 0 in the Dup1 and Dup2 regions (see **Supplementary Figure S40**). In contrast, the mapping qualities in the rest of the *Fm* locus, including the junction regions, are close to 60. The mapping quality analysis also suggests that the GRCg6a reference has better mapping than CAU_silkie and is unaffected by reference bias.

In our case, the two sets of reads mapped to the same location in the GRCg6a reference correspond to the two haplotypes, not some form of unknown mapping bias. We can be confident of this as the long reads from PacBio and ONT form haplotype-consistent tiling paths through the entire lengths of Dup1 and Dup2 regions. Such haplotype-consistent paths are not expected in the case of reference bias.

### 7. Individual read-based support for the haplotypes

To further demonstrate the validity of the haplotypes we have reconstructed, the actual long-reads from each haplotype are visualized in **Supplementary Figure S41-44**. At each HDP, the colour represents whether the base in a particular read matches the haplotype-specific base. The tiling path of reads covers the entire Dup1 and Dup2 regions in haplotype-consistent paths. In contrast to looking at pairs of HDPs, this representation demonstrated the consistency of the haplotypes across the entire length of the long-reads containing several HDPs.

Dup1 example reads:

For example, long-read#SRR17968711.1911295 spans 102 HDPs in the Dup1 region and is assigned to haplotype-1. Of the 102 HDPs, we found that the base of the read at 99 HDPs matches the assigned haplotype. Only at one HDP does the base in the read not match any haplotype, and at two HDPs, the base in the read matches haplotype-2. **Supplementary Table S2** provides the full list of HDPs and the base in the read at each HDP.

Similarly, the long-read#SRR17968711.1022612 spans 108 HDPs in the Dup1 region and is assigned to haplotype-2. Of the 108 HDPs, we found that the base of the read at 104 HDPs matches the assigned haplotype. Only at one HDP does the base in the read not match any haplotype, and at three HDPs, the base in the read matches haplotype-1. **Supplementary Table S3** provides the full list of HDPs and the base in the read at each HDP.

Dup2 example reads:

The long-read#SRR17968711.852855 spans 78 HDPs in the Dup2 region and is assigned to haplotype-1. Of the 78 HDPs, we found that the base of the read at 73 HDPs matches the assigned haplotype. Only at two HDPs does the base in the read not match any haplotype, and at three

HDPs, the base in the read matches haplotype-2. **Supplementary Table S4** provides the full list of HDPs and the base in the read at each HDP.

The long-read#SRR17968711.2221154 spans 64 HDPs in the Dup2 region and is assigned to haplotype-2. Of the 64 HDPs, we found that the base of the read at 60 HDPs matches the assigned haplotype. Only at two HDPs does the base in the read not match any haplotype, and at two HDPs, the base in the read matches haplotype-1. **Supplementary Table S5** provides the full list of HDPs and the base in the read at each HDP.

These individual reads with high concordance with the assigned haplotype at such a large number of HDPs cannot be explained by sequencing errors and support phasing of the haplotypes over several Kb. In contrast to short-reads, such long-read-based phasing has the unique advantage of disentangling haplotypes over such long distances.

### 8. Junction coverage can identify if the individual is heterozygote or homozygote

An important point to consider in the context of coverage at the junctions is that the coverage at the four junctions can determine if the individual under consideration is a heterozygote or homozygote for the *Fm* allele. The original Dorshorst paper estimates whether the individuals are heterozygote or homozygote using qPCR (see **Figure 3** in the Dorshorst paper[4])

The CAU_Silkie individual assembled is a homozygote (*Fm/Fm*). As expected in the case of a homozygote, the coverage at all four internal junctions is very similar in both the PacBio and ONT data. Even the flanking junctions (Flank1-Dup1 and Dup2-Flank2) have coverage similar to that of the four internal junctions.

| | **Flanking Junctions** | | **Internal Junctions** | | | |
|---|---|---|---|---|---|---|
| | Found in both *N** and *Fm* rearrangement scenarios | | | | Found only in the *Fm* rearrangement scenarios | |
| Sequencing Technology | Flank1_Dup1 | Dup2_Flank2 | Dup1_Int (Junction-3) | Int_Dup2 (Junction-2) | Dup1_Dup2_R (Junction-1) | Dup1_R_ Dup2 (Junction-4) |
| PacBio Reads | 411 | 377 | 563 | 558 | 424 | 460 |
| ONT Reads | 56 | 42 | 52 | 47 | 42 | 37 |

Suppose the individual used was a heterozygote; junctions 2 (Int_Dup2) and 3 (Dup1_Int) coverage should be twice that at junctions 1 (Dup1_Dup2_R) and 4 (Dup1_R_Dup2) because junctions 2 and 3 occur in both *N** and *Fm* while junctions 1 and 4 occur only in the *Fm* arrangement. The flanking junctions (Flank1-Dup1 and Dup2-Flank2) coverage should also be twice that at junctions 1 and 4 because the flanking junctions occur in both *N** and *Fm* while junctions 1 and 4 occur only in the *Fm* arrangement. However, this is not true in the PacBio or ONT junction spanning reads. Therefore, the Silkie individual used for assembly is a homozygote (*Fm/Fm*) and cannot be a heterozygote. Haplotype phasing over the entire length of Dup1 and Dup2 would not have been possible if the individual was a (*N*/Fm*) heterozygote.

### 9. Reliability of GRCg6a chicken genome at the *Fm* locus

The GRCg6a/GGA6a/galGal6 genome is constructed using 'RJF #256' from an inbred line (UCD 001). Various established genomic regions within GRCg6a are known to contain assembly errors and gaps[31]. However, the *Fm* locus region in wild-type birds (**N*) does not exhibit such well-known assembly errors (https://www.ncbi.nlm.nih.gov/grc/tpf/chicken/chr20). To further evaluate the genome assembly quality at the *Fm* locus, we obtained the ONT long-reads of the bioproject# PRJNA693184 generated from chicken of the Huxu breed. These ONT long-reads from wild-type (**N*) birds were mapped to the GRCg6a chicken reference genome using the bwa read mapper. The resulting read alignments were converted from bam to bed format and visualised using the UCSC genome browser. Our independent verification of the *Fm* locus region, employing the ONT long-read dataset from wild-type birds, reveals robust tiling paths, providing substantial support. There is no indication of collapsed assemblies or missing/gap regions in this specific genomic area in GRCg6a (see **Supplementary Figure S45**).

Few zero-depth 100-bp windows are located in the region 20:11,298,343-11,299,157. Viewing this region in IGV clearly shows that this region contains a ~600bp deletion specific to the Huxu breed (**Supplementary Figure S46**). In contrast to the Huxu breed (SRR15421342), this deletion is not found in other chicken long-read data (for example, Arbor Acres breeder roosters (SRR13494713)) (**Supplementary Figure S47**). Since the deletion is Huxu breed-specific, it is unsurprising that it is not visible when the Huxu reads are mapped to the Huxu genome. Similarly, a few 100-bp, 10-bp, and 5-bp windows with zero depth and Huxu breed-specific deletions are located across the *Fm* locus (**Supplementary Figure S48-52**). As the *Fm* locus has a good quality assembly, using the GRCg6a chicken reference as an anchor for the read-backed haplotype-phasing of the *Fm* locus region is unaffected by genome assembly errors.

**10. Sequence context & homopolymer errors at HDPs in read-backed haplotype-phasing**

Identifying HDPs and quantifying long-read support at pairs of HDPs relies upon counting the number of reads consistent with the haplotypes. Some variants of ONT have an issue of systematic errors in long homopolymer runs, as reported earlier[32]. Therefore, systematic base calling errors at the same bases in ONT reads can potentially masquerade as HDPs.

To evaluate the effect of homopolymer runs, we searched for homopolymer runs of 2 or more consecutive bases within ten base pair flanks on both sides of all the HDPs. Of the 122 HDPs in the Dup1 region, 37 homopolymer runs, and among the 138 HDPs in the Dup2 region, 45 homopolymer runs were identified. The five most pronounced homopolymer runs from Dup1 (**Supplementary Figure S53-57**) and Dup2 (**Supplementary Figure S58-62**) were visually inspected in IGV to ensure that the reads that support a given base do not result from systematic base calling errors.

**11. Fraction & count of reads supporting HDP pairs**

To evaluate the long-read support for individual haplotypes, we quantified the number of reads supporting each of the 16 possible combinations of bases for every pair of HDPs. The expectation is that most of the reads should support the two combinations corresponding to the two haplotypes. However, some reads may support other combinations due to sequencing errors at either or both positions. Moreover, within-haplotype polymorphisms can increase support for the third and fourth combinations. We excluded tri-allelic and tetra-allelic sites when selecting sites as HDPs. Yet, some of the within-haplotype polymorphisms could have escaped this filtering step and can support the third and fourth combinations. The fraction of reads supporting every pair of bases at consecutive HDPs was calculated based on the total number of reads spanning every HDP pair. The most well-supported combination of bases tends to have 30 to 70% of the reads, and the second most well-supported combination tends to have 20 to 50%. While the third combination is supported by < 25% of reads, the fourth and fifth combinations have < 10 % of reads (**Supplementary Figure S63-66**). Few pairs with considerable support for the third combination could be within-haplotype polymorphisms.

We quantified the fraction of reads supporting different combinations over the length of Dup1 and Dup2 regions. Over the entire length of the duplicated regions, the first and second most well-supported combinations had a much higher fraction of reads supporting them than the third, fourth and fifth combinations (**Supplementary Figure S67-74**). The few examples in which the second and third most well-supported combinations had similar support happened to be within-haplotype polymorphisms (**Supplementary Figure S75-83**).

The fraction of reads supporting a particular combination does not provide a measure of magnitude as the numbers are expressed as a fraction. Therefore, we also compared the number of reads supporting each combination (**Supplementary Figure S84-95**). Especially in the PacBio dataset, hundreds of reads supported the third and, in some rare cases, the fourth combination. These sites correspond to within-haplotype polymorphisms.

We further looked for long-read support for three consecutive HDPs at a time. In contrast to pairs of HDPs, looking at HDP triplets provides greater confidence in the reconstructed haplotypes (**Supplementary Figure S96-119**). The strong support for HDP pairs and HDP triplets nullified any possibility of sequencing errors masquerading as haplotypes.

In our read-backed haplotype-phasing approach, we employ the GRCg6a genome solely as an anchor to identify haplotype-consistent tiling paths using haplotype-defining positions within long-read datasets. The use of GRCg6a as the reference is comparable to the PanGenome graph approaches pioneered in analysing structural variants in human and chicken genomes[27,33]. The PanGenome assembly approaches rely on T2T genomes and use the highest-quality references to build PanGenome graphs. Read mapping-based phasing of genome assemblies is a fairly common practice for human genomes[34,35]. Several popular software tools (WhatsHap[36], WhatsHap polyphase[37], HapCut[38], HapCut2[39], Haptree[40], ProbHap[41], HapCompass[42], H-BOP[43], MixSIH[44], LongPhase[45], nPhase[46], HAT[47], Ranbow[48], flopp[49], Hap10[50], EHTLD[51], etc.) are available to perform such phasing. Simulation-based studies have compared the performance of haplotype-phasing approaches in resolving polyploid genomes[52]. Our recent paper[3], which performs haplotype phasing of the *Fm* locus, demonstrates a special case of using a read-backed haplotype phasing strategy to resolve a complex chromosomal rearrangement. Haplotype phasing has been used to reconstruct large-scale complex chromosomal rearrangements to resolve the karyotypic changes associated with cancers[53].

### 12. High sequence similarity in the Dup1 & Dup2 regions of GRCg6a and CAU_Silkie genome

We performed multiple sequence alignment of the two copies of the duplicated regions from the CAU_Silkie genome along with the corresponding region from the GRCg6a genome assembly using CLUSTALW[54]. The analysis revealed a remarkable level of sequence similarity, exceeding 99%. Subsequently, we utilised progressiveMauve[55] to perform the same alignments independently and visualise the local changes in sequence similarity along the length of the sequences. Our examination of the GRCg6a genome assembly compared to the CAU_Silkie genome reveals a striking similarity in the regions, devoid of substantial insertion/deletion events. The variations are confined to nucleotide sequence differences between the two duplicated regions' copies. The genomes exhibit a high degree of comparability with only minor divergence, amounting to a few hundred nucleotide differences, except for the identified chromosomal rearrangement.

### 13. Read-backed phasing of Dup1 and Dup2 regions based on GRCg6a reference

Our earlier paper[3] includes the mapping locations (in .bed format https://github.com/ceglabsagarshinde/Kadaknath_Project/tree/main) of each ONT read (with read IDs) by haplotype, enabling easy loading and visualisation in the UCSC genome browser. The decision to use the GRCg6a genome was influenced by the unavailability of a high-quality genome for the Silkie breed at the time of our study, coupled with the convenient availability of various visualisation options for the GRCg6a genome. Using the GRCg6a genome also makes it easy to compare the corresponding nucleotide states between the haplotypes directly. HDPs were identified by mapping the ONT and PacBio long reads from CAU Silkie to the GRCg6a genome assembly. A detailed flowchart outlining the methodology explains the steps for obtaining the final set of HDPs (**Supplementary Figure S120**). A graphical example of long-read-based haplotype phasing provides an overview of read-backed haplotype phasing (**Supplementary Figure S121**).

The annotated code used for this haplotype-phasing process is provided below:

```
#################################################################
##Mapping
#The CAU Silkie long-read data was mapped to GRCg6a using BWA.
```

```
bwa index Gallus_gallus.GRCg6a.dna_sm.toplevel.fa.gz
for i in *.fastq.gz
do
bwa bwasw -t 24 -a2 -b3 -q2 -r2 -z1 Gallus_gallus.GRCg6a.dna_sm.toplevel.fa.gz $i > "$i".sam
samtools view -bhS -@ 24 "$i".sam > "$i".bam
samtools sort "$i".bam -@ 24 -o "$i".sorted.bam
samtools index  "$i".sorted.bam
done
```

**##BAM read-count**

**#We used the Bam read-count utility to identify positions with more than ten reads for more than one allele.**

**#Excluded the tri and tetra allelic positions to get the potential set of HDps.**

```
bam-readcount -f Gallus_gallus.GRCg6a.dna_sm.toplevel.fa -w 0 bamfile -l dup1_dup2.bed > bam_readcount.txt
for j in *txt;do cut -f 1-4,6-9 $j|sed 's/:/\t/g'|cut -f 1-4,5,6,19,20,33,34,47,48 > $j.count;done
for i in `cat bam_readcount.txt.count|sed 's/\t/_/g'`
do
val1=`echo $i|cut -f 6,8,10,12 -d '_'|tr "_" "\n"|sort -k1n,1|tail -n 1`
val2=`echo $i|cut -f 6,8,10,12 -d '_'|tr "_" "\n"|sort -k1n,1|tail -n 2|head -n 1`
echo $val1 $val2|awk -v rat=$i '$1>10 && $2>10{print rat}'|sed 's/_/\t/g' >> atleast_10.txt
done
```

**#For each Potential HDP identified in GRCg6a, we created an ATGC.tsv file with all four bases (A, T, G, C), which can be present in haplotypes.**

```
for i in `cat GRC_pos`
do
  kl="SRR17968711_12"
  echo  "20@$i@A@HapA\n" | sed 's/@/\t/g' > ATGC.tsv
  echo  "20@$i@T@HapT\n" | sed 's/@/\t/g' >> ATGC.tsv
  echo  "20@$i@G@HapG\n" | sed 's/@/\t/g' >> ATGC.tsv
```

```
  echo  "20@$i@C@HapC\n" | sed 's/@/\t/g' >> ATGC.tsv
```

**#Use the biostar214299.jar tool to process the BAM file with the positions in ATGC.tsv to define the haplotypes based on the given base information at each HDP and output a BAM file.**

```
  java -jar biostar214299.jar --samoutputformat BAM -p ATGC.tsv
SRR17968711_12_CAU_Silkie_Fm_subseted.bam > "$i"_"$kl".bam

  samtools sort -@16 "$i"_"$kl".bam -o "$i"_"$kl".sorted.bam

  samtools index "$i"_"$kl".sorted.bam
```

**#Extract reads for each haplotype from the sorted BAM file and convert to FASTA format**

```
  for hapcount in A T G C

  do

    samtools view -@16 -b -r Hap"$hapcount" "$i"_"$kl".sorted.bam > "$i"_hap"$hapcount".bam

    samtools fasta "$i"_hap"$hapcount".bam > "$i"_hap"$hapcount".fa
```

**#Extract the SRR ID for each haplotype from the FASTA file.**

```
  grep ">" "$i"_hap"$hapcount".fa | sed 's/>//g' | sort -u > "$i"_hap"$hapcount".list

  done

  mv *.list readlists

  rm "$i"_"$kl".bam "$i"_"$kl".sorted.bam "$i"_"$kl".sorted.bam.bai "$i"_hap*.bam
"$i"_hap*.fa

done
```

**#For two consecutive HDPs, we count all 16 base pair combinations and supporting read count for Dup1 and Dup2**

**# Dup1 Phasing**

```
for i in `cat dup1_pair`

do

  pos1=`echo $i | sed 's/_/\t/g' | cut -f1`

  pos2=`echo $i | sed 's/_/\t/g' | cut -f2`

  count_AA=`cat "$pos1"_hapA.list "$pos2"_hapA.list | sort | uniq -c | awk '$1>1{print $0}' | wc
-l`

  count_AT=`cat "$pos1"_hapA.list "$pos2"_hapT.list | sort | uniq -c | awk '$1>1{print $0}' | wc -
l`
```

```
  count_AG=`cat "$pos1"_hapA.list "$pos2"_hapG.list | sort | uniq -c | awk '$1>1{print $0}' | wc
-l`
  count_AC=`cat "$pos1"_hapA.list "$pos2"_hapC.list | sort | uniq -c | awk '$1>1{print $0}' | wc
-l`
  count_TA=`cat "$pos1"_hapT.list "$pos2"_hapA.list | sort | uniq -c | awk '$1>1{print $0}' | wc -
l`
  count_TT=`cat "$pos1"_hapT.list "$pos2"_hapT.list | sort | uniq -c | awk '$1>1{print $0}' | wc -
l`
  count_TG=`cat "$pos1"_hapT.list "$pos2"_hapG.list | sort | uniq -c | awk '$1>1{print $0}' | wc -
l`
  count_TC=`cat "$pos1"_hapT.list "$pos2"_hapC.list | sort | uniq -c | awk '$1>1{print $0}' | wc -
l`


  count_GA=`cat "$pos1"_hapG.list "$pos2"_hapA.list | sort | uniq -c | awk '$1>1{print $0}' | wc
-l`
  count_GT=`cat "$pos1"_hapG.list "$pos2"_hapT.list | sort | uniq -c | awk '$1>1{print $0}' | wc -
l`
  count_GG=`cat "$pos1"_hapG.list "$pos2"_hapG.list | sort | uniq -c | awk '$1>1{print $0}' | wc
-l`
  count_GC=`cat "$pos1"_hapG.list "$pos2"_hapC.list | sort | uniq -c | awk '$1>1{print $0}' | wc
-l`
  count_CA=`cat "$pos1"_hapC.list "$pos2"_hapA.list | sort | uniq -c | awk '$1>1{print $0}' | wc
-l`
  count_CT=`cat "$pos1"_hapC.list "$pos2"_hapT.list | sort | uniq -c | awk '$1>1{print $0}' | wc -
l`
  count_CG=`cat "$pos1"_hapC.list "$pos2"_hapG.list | sort | uniq -c | awk '$1>1{print $0}' | wc
-l`
  count_CC=`cat "$pos1"_hapC.list "$pos2"_hapC.list | sort | uniq -c | awk '$1>1{print $0}' | wc -
l`
  echo $count_AA AA > temp.counts
  echo $count_AT AT >> temp.counts
  echo $count_AG AG >> temp.counts
  echo $count_AC AC >> temp.counts
```

```
  echo $count_TA TA >> temp.counts
  echo $count_TT TT >> temp.counts
  echo $count_TG TG >> temp.counts
  echo $count_TC TC >> temp.counts
  echo $count_GA GA >> temp.counts
  echo $count_GT GT >> temp.counts
  echo $count_GG GG >> temp.counts
  echo $count_GC GC >> temp.counts
  echo $count_CA CA >> temp.counts
  echo $count_CT CT >> temp.counts
  echo $count_CG CG >> temp.counts
  echo $count_CC CC >> temp.counts
  # Sort and filter haplotype combinations with counts greater than 10
  haps=`sort -k1n,1 temp.counts | awk '$1>10{print $0}' | sed 's/ /_/g' | tr "\n" ","`
  # Output the results for pos1 and pos2 along with counts and haplotypes
  echo $pos1 $pos2 $count_AA $count_AT $count_AG $count_AC $count_TA $count_TT $count_TG $count_TC $count_GA $count_GT $count_GG $count_GC $count_CA $count_CT $count_CG $count_CC $haps
done
# Dup2 Phasing
for i in `cat dup2_pair`
do
  pos1=`echo $i | sed 's/_/\t/g' | cut -f1`
  pos2=`echo $i | sed 's/_/\t/g' | cut -f2`
  count_AA=`cat "$pos1"_hapA.list "$pos2"_hapA.list | sort | uniq -c | awk '$1>1{print $0}' | wc -l`
  count_AT=`cat "$pos1"_hapA.list "$pos2"_hapT.list | sort | uniq -c | awk '$1>1{print $0}' | wc -l`
  count_AG=`cat "$pos1"_hapA.list "$pos2"_hapG.list | sort | uniq -c | awk '$1>1{print $0}' | wc -l`
```

```
count_AC=`cat "$pos1"_hapA.list "$pos2"_hapC.list | sort | uniq -c | awk '$1>1{print $0}' | wc -l`
count_TA=`cat "$pos1"_hapT.list "$pos2"_hapA.list | sort | uniq -c | awk '$1>1{print $0}' | wc -l`
count_TT=`cat "$pos1"_hapT.list "$pos2"_hapT.list | sort | uniq -c | awk '$1>1{print $0}' | wc -l`
count_TG=`cat "$pos1"_hapT.list "$pos2"_hapG.list | sort | uniq -c | awk '$1>1{print $0}' | wc -l`
count_TC=`cat "$pos1"_hapT.list "$pos2"_hapC.list | sort | uniq -c | awk '$1>1{print $0}' | wc -l`
count_GA=`cat "$pos1"_hapG.list "$pos2"_hapA.list | sort | uniq -c | awk '$1>1{print $0}' | wc -l`
count_GT=`cat "$pos1"_hapG.list "$pos2"_hapT.list | sort | uniq -c | awk '$1>1{print $0}' | wc -l`
count_GG=`cat "$pos1"_hapG.list "$pos2"_hapG.list | sort | uniq -c | awk '$1>1{print $0}' | wc -l`
count_GC=`cat "$pos1"_hapG.list "$pos2"_hapC.list | sort | uniq -c | awk '$1>1{print $0}' | wc -l`
count_CA=`cat "$pos1"_hapC.list "$pos2"_hapA.list | sort | uniq -c | awk '$1>1{print $0}' | wc -l`
count_CT=`cat "$pos1"_hapC.list "$pos2"_hapT.list | sort | uniq -c | awk '$1>1{print $0}' | wc -l`
count_CG=`cat "$pos1"_hapC.list "$pos2"_hapG.list | sort | uniq -c | awk '$1>1{print $0}' | wc -l`
count_CC=`cat "$pos1"_hapC.list "$pos2"_hapC.list | sort | uniq -c | awk '$1>1{print $0}' | wc -l`
echo $count_AA AA > temp.counts
echo $count_AT AT >> temp.counts
echo $count_AG AG >> temp.counts
echo $count_AC AC >> temp.counts
echo $count_TA TA >> temp.counts
echo $count_TT TT >> temp.counts
echo $count_TG TG >> temp.counts
```

```
  echo $count_TC TC >> temp.counts
  echo $count_GA GA >> temp.counts
  echo $count_GT GT >> temp.counts
  echo $count_GG GG >> temp.counts
  echo $count_GC GC >> temp.counts
  echo $count_CA CA >> temp.counts
  echo $count_CT CT >> temp.counts
  echo $count_CG CG >> temp.counts
  echo $count_CC CC >> temp.counts
  haps=`sort -k1n,1 temp.counts | awk '$1>10{print $0}' | sed 's/ /_/g' | tr "\n" ","`
echo $pos1 $pos2 $count_AA $count_AT $count_AG $count_AC $count_TA $count_TT
$count_TG $count_TC $count_GA $count_GT $count_GG $count_GC $count_CA $count_CT
$count_CG $count_CC $haps
done
################################################################
```

In the current analysis, we have extended our read-backed phasing of Dup1 and Dup2 regions based on all 260 HDPs identified. Using GRCg6a as a reference, we phased the ONT and PacBio long-read data from CAU_Silkie (see **Supplementary Tables S6-S9**), PacBio data from another Silkie individual (**Supplementary Tables S10** and **S11**), and Lueyang individual (**Supplementary Tables S12** and **S13**) generated by Northwest A&F University. For Northwest A&F University Silkie, 258 (122 for Dup1 and 136 for Dup2 (3 new + 133 previous) HDPs have been used. For the Northwest A&F University Lueyang individual, 259 HDPs (122 for Dup1 and 137 for Dup2 (22 new + 115 previous) have been used to phase the Lueyang PacBio data. The read-backed phasing of all three individuals was consistent and supported the **Fm_2* scenario.

#### 14. <u>Liftover of HDPs identified in GRCg6a to CAU_Silkie, Lueyang & Huxu genomes</u>

The corresponding positions between GRCg6a and the other three reference genomes were identified based on pairwise chain alignments generated by lastz. Since the CAU_Silkie and Lueyang reference has two copies of Dup1 and Dup2 regions, we masked one copy of Dup1 and Dup2 to generate the first set of alignments and masked the other copy to get the other set of alignments. These two sets of alignments were used to liftover the HDPs from GRCg6a to the

CAU_Silkie and Lueyang reference genome using the liftOver utility. As Huxu has only one copy of Dup1 and Dup2, masking was not required.

### 15. Read-backed Phasing of Dup1 and Dup2 Regions Using Huxu & Lueyang Genomes

CAU_Silkie ONT long reads were aligned to both the Huxu and Lueyang reference genome assemblies for read-backed phasing, using HDPs identified through liftover. The HDPs identified in Huxu successfully phased the CAU_Silkie ONT data and supported the GRCg6a-based phasing (**Supplementary Tables S14 & S15**). Similarly, the HDPs identified in Lueyang were also able to phase the Dup1 and Dup2 regions (**Supplementary Tables S16-19**). The read-backed phasing performed with both genome assemblies was consistent, reinforcing the **Fm_2* scenario.

### 16. Read-backed phasing of Dup1 and Dup2 regions based on CAU_Silkie genome

The recent release[24] of the high-quality chromosome-level CAU_Silkie genome allows us to reassess our analysis with this updated genomic resource. Hence, our current re-analysis used the CAU_Silkie genome to assess the haplotype phasing spanning HDPs. The CAU_Silkie long-reads are assigned haplotypes based on the read-backed phasing of reads mapped to the GRCg6a genome. The same reads mapped to the CAU_Silkie genome lack support for the sequences assembled as Dup1, Dup2r, Dup1r and Dup2 (**Supplementary Figure S122**). At the beginning of Dup1, the HDPs (11016732, 11021560 and 11029748) are correctly assembled in the CAU_Silkie and are supported by the raw read with reads from a single haplotype mapped to the genome (**Supplementary Figure S123-125**). However, towards the end of Dup1, near the HDP-poor region, the genomic sequence of CAU_Silkie has alleles from the other haplotype at several HDPs (11051834, 11066868 and 11102690). Upon inspection of read support for the genomic sequence at these HDPs, reads from both haplotypes were found to be mapping. More importantly, the allelic state of the genomic sequence at consecutive HDPs does not match the allelic state of the long-reads at the same HDPs (**Supplementary Figure S126-130**). A similar trend of mosaic haplotypes is evident near HDP-poor regions in Dup2r (11191958, 11195286, 111209872, 11275674, and 11294797), Dup1r (11768362, 11814782 and 11831962) and Dup2 (11874547, 11894857, 11920571, and 11973510) (**Supplementary Figure S131-155**). However, the regions distant from the HDP-poor regions (Dup2r: 11144700 and 11161186; Dup1r: 11841295, 11849483 and 11854311; Dup2: 12005042, 12014943 and 12024652) are properly assembled in the CAU_Silkie genome and are well-supported by long-reads (**Supplementary Figure S156-163**).

This re-analysis shows that **(A)** Continuous raw read support for the CAU_Silkie is missing at the identified haplotype-switch locations. **(B)** Raw read support from ONT and PacBio data independently supports the haplotypes inferred by our approach. Using CAU_Silkie genome assembly, we identified additional HDPs supporting the **Fm_2* scenario based on the read-backed phasing approach. **(C)** The reads from both haplotypes are mapped at the locations with mosaic haplotypes and can be evaluated in read mapping to the CAU_Silkie genome (see **Supplementary Table S20-27**). The haplotype-consistent paths we identify are well supported irrespective of the genome assembly used for the inference and demonstrate that **Fm_2* is the correct scenario.

#### 17. Limitations and Future Directions

Our ability to perform haplotype-resolved genome assembly of complex rearrangements depends on the haplotype-specific sequence level differences. In the case of the *Fm* locus, such haplotype-specific sequences are identified as HDPs or Haplotype Defining Positions. As noted earlier, the main challenge to successfully assembling the entire rearrangement is the paucity of such HDPs at the end of the Dup1 region and the start of the Dup2 region. The availability of ONT long-reads in CAU_Silkie allowed us to resolve the rearrangement by relying upon the ability of these reads to span across several Kb in the HDP-poor region. Notably, the hifiasm-based assembly of the Lueyang black-bone chicken using PacBio reads alone is possible due to additional HDPs at the HDP-poor region in the individual used for genome sequencing. Another solution to spanning such HDP-poor regions is using methylation-determined HDPs or MD-HDPs. While the nucleotide states at MD-HDPs are identical in both haplotypes, the methylation state would be haplotype-specific. Tools such as MethPhaser[56] provide a ready-to-use setup to perform such MD-HDP-based haplotype phasing. Unfortunately, the methylation data associated with the CAU_Silkie data is currently not publicly available and could not be used to resolve the *Fm* locus.

**Longer read lengths**:

Further increments in the length of long-reads and reduction in their error rates are expected with technological advances. For instance, the average length of ONT reads in the Huxu breed dataset is higher than that in the CAU_Silkie data. Such improvements in read length will allow the phasing of haplotypes with fewer HDPs due to the availability of longer-range connectivity information.

**Optical Mapping**:

The use of optical mapping technology to improve the quality of genomes and discover structural variation has seen increasing success in the past few years[57]. Although the two copies of the duplicated regions are highly similar at the *Fm* locus, using optical mapping to span these regions will identify the unique flanking regions for each copy. Hence, optical mapping provides another approach to validate the genomic organisation of the *Fm* locus.

**Trio-binning**:

The use of parent-child trios to perform haplotype phasing has been widely used to reconstruct genomes with better contiguity and accurate resolution of haplotypes[58]. Resolution of complex rearrangements, such as the *Fm* locus, is well-suited for trio-binning. Several crosses (*Fm/Fm* vs *N*/N**, *Fm/Fm* vs *Fm/N** and *Fm/Fm* vs *Fm/Fm*) are possible for trio-binning at the *Fm* locus. We envisage that the assembly of the *Fm* locus can serve as a benchmark for evaluating different genome assembly approaches and tools in the future.

Several genome assembly tools provide output as primary and alternative assembly fasta sequences to represent the assembly of haplotypes. Some recent assemblers also provide a graph-based output file in the GFA format. While the GFA format version 1.0 has several useful features that allow exploration of the sequences assembled, the details of the sequencing reads used for assembling different parts of the genome are not readily available. The ACE format used by some of the early genome assemblers provided information regarding the reads used for assembly. The availability of such information is handy for assembly validation and troubleshooting.
