## Supplementary Figures for "What is the correct genomic structure of the complex chromosomal rearrangement at the *Fm* locus in Silkie chicken?"

ARISING FROM Zhu, F., Yin, ZT., Zhao, QS. et al. Communications Biology https://doi.org/10.1038/s42003-023-05619-y (2023)

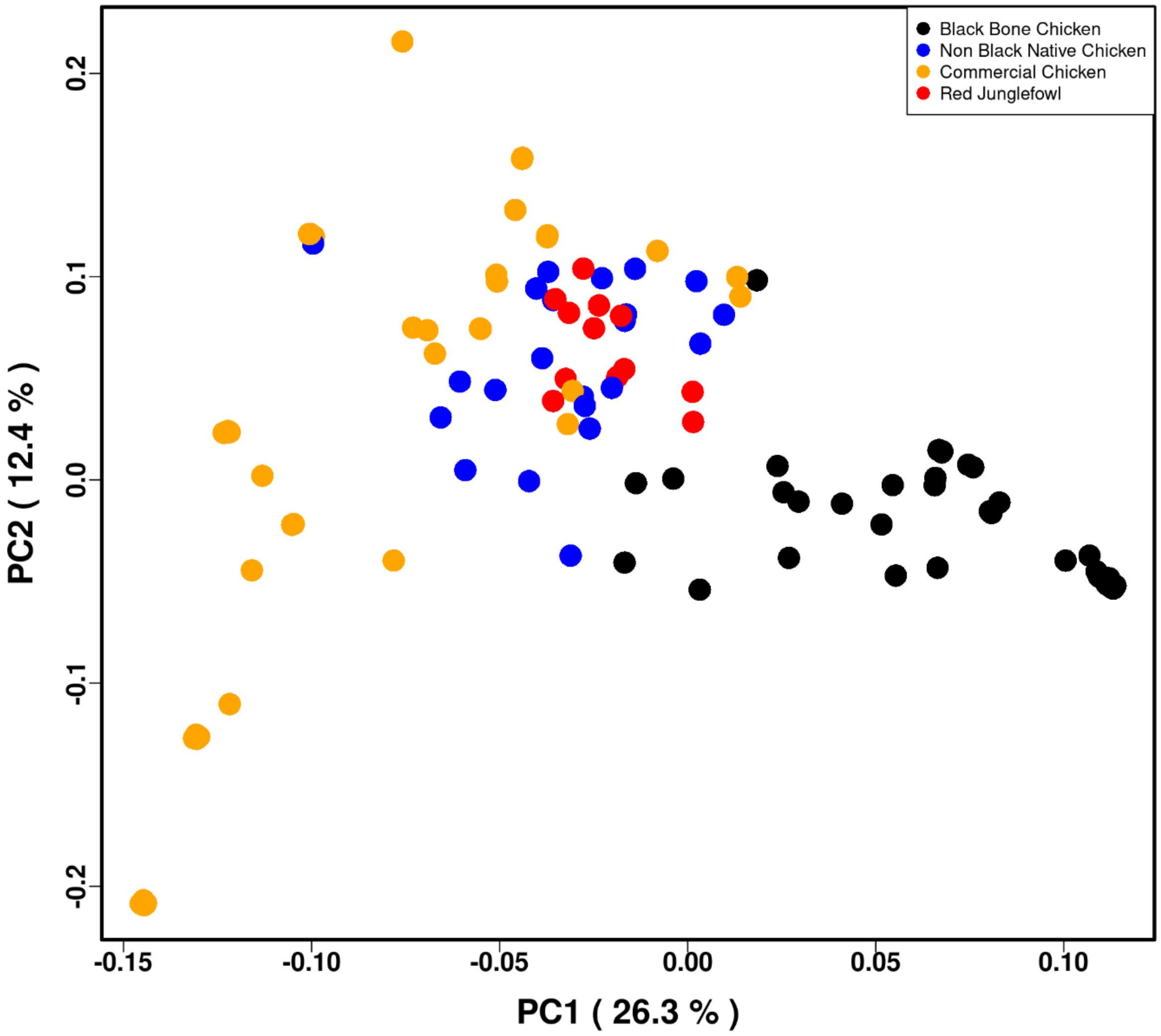


**Supplementary Figure. S1: Principal Component Analysis (PCA) of Genetic Relationships Among 139 Chicken Individuals Based on the Dup1 Region:** This PCA plot illustrates the genetic relationships among 139 chicken individuals, including 61 Black-bone and 78 non-Black bone chickens, based on the Dup1 region. The PC1 accounts for 26.3% of the variance, while the PC2 accounts for 12.4%. Different chicken groups are color-coded for clarity: Black-bone chicken breeds are shown in black, non-Black-bone native chicken breeds in blue, commercial chicken breeds in orange, and wild red junglefowl in red.

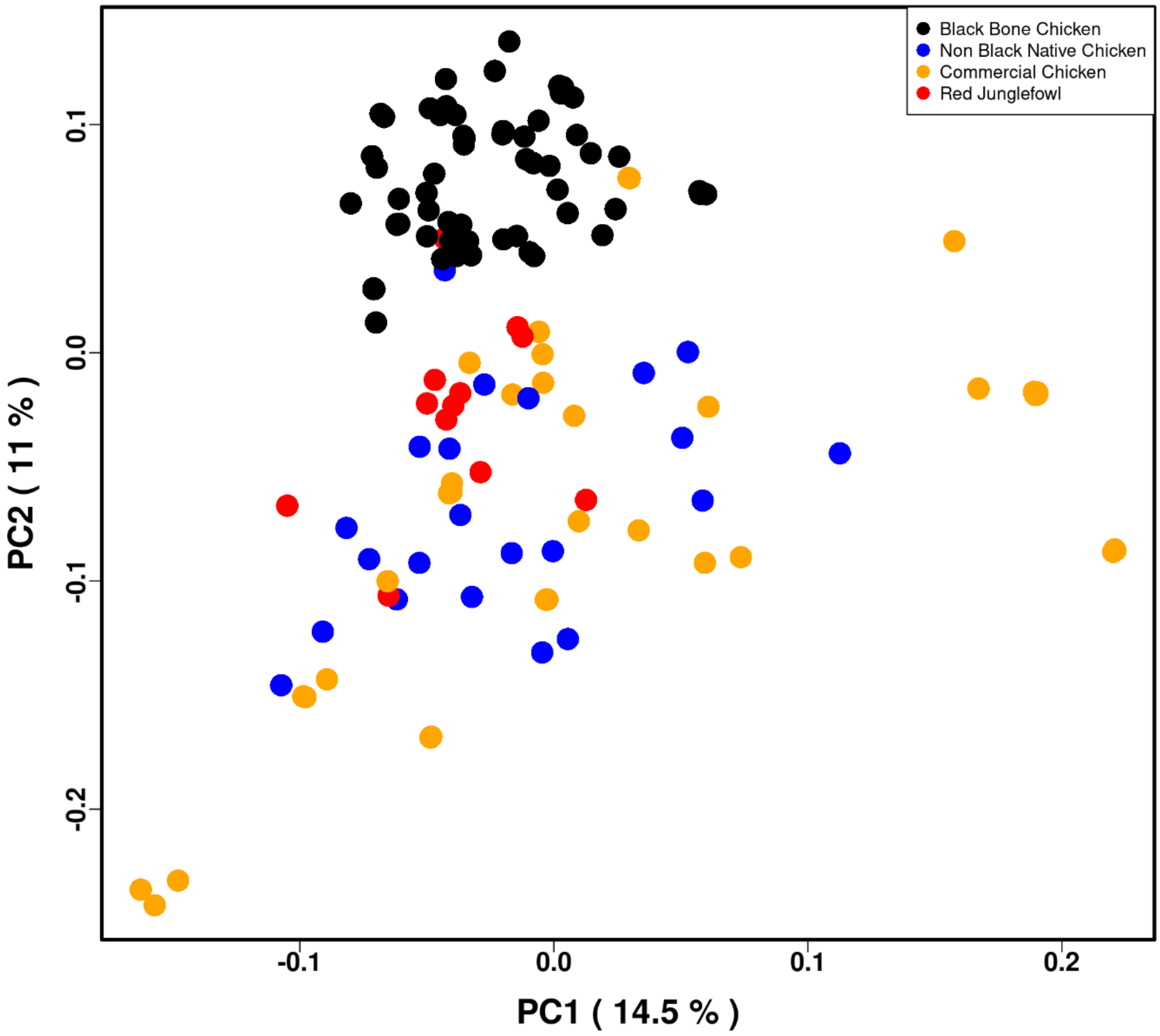


**Supplementary Figure. S2: Principal Component Analysis (PCA) of Genetic Relationships Among 139 Chicken Individuals Based on the Dup2 Region:** This PCA plot illustrates the genetic relationships among 139 chicken individuals, including 61 Black-bone and 78 non-Black bone chickens, based on the Dup2 region. The PC1 accounts for 14.5% of the variance, while the PC2 accounts for 11%. Different chicken groups are color-coded for clarity: Black-bone chicken breeds are shown in black, non-Black-bone native chicken breeds in blue, commercial chicken breeds in orange, and wild red junglefowl in red.

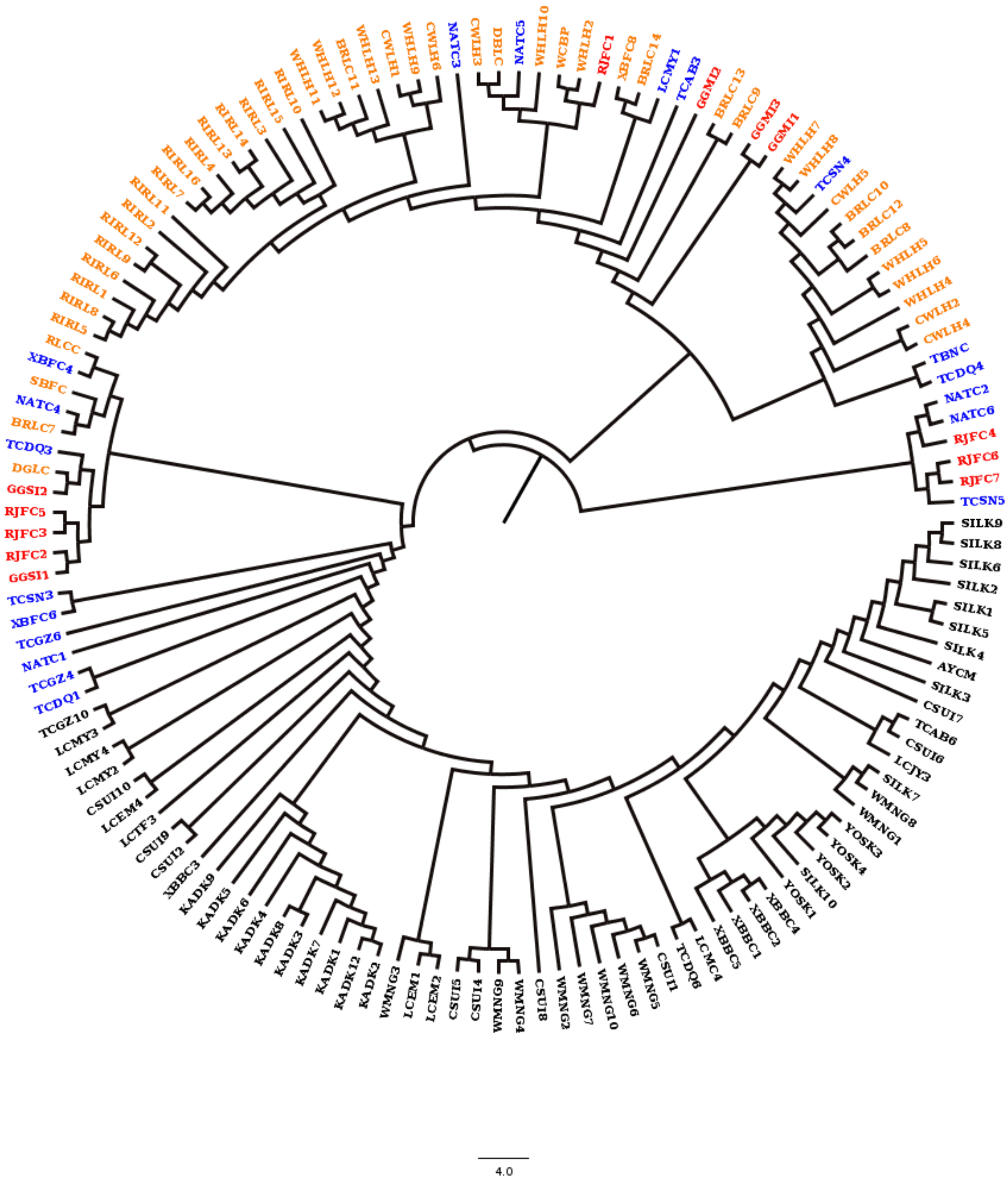


**Supplementary Figure. S3: Local Phylogeny of 139 Chicken Individuals Based on the Dup1 Region:** Phylogenetic tree of SNPs within the Dup1 region constructed for 139 chicken individuals, comprising 61 Black-bone and 78 non-Black-bone chickens. The tree was generated using iqtree2 and visualized with FigTree. Black-bone chicken breeds are depicted in black, non-black-bone native breeds in blue, commercial breeds in orange, and wild red junglefowl in red.

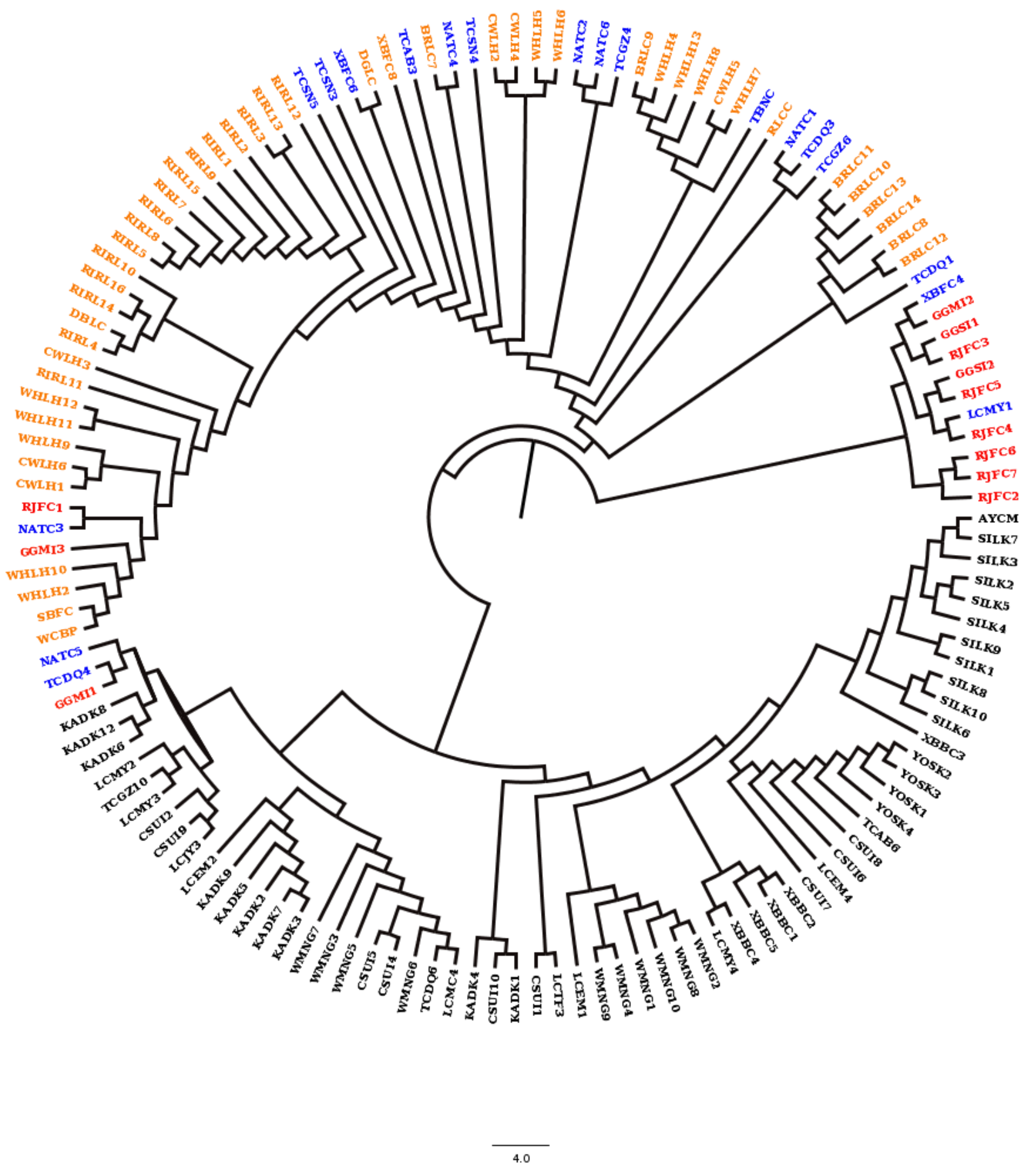

**Supplementary Figure. S4: Local Phylogeny of 139 Chicken Individuals Based on the Dup2 Region:** Phylogenetic tree of SNPs within the Dup2 region constructed for 139 chicken individuals, comprising 61 Black-bone and 78 non-Black-bone chickens. The tree was generated using iqtree2 and visualized with FigTree. Black-bone chicken breeds are depicted in black, non-black-bone native breeds in blue, commercial breeds in orange, and wild red junglefowl in red.

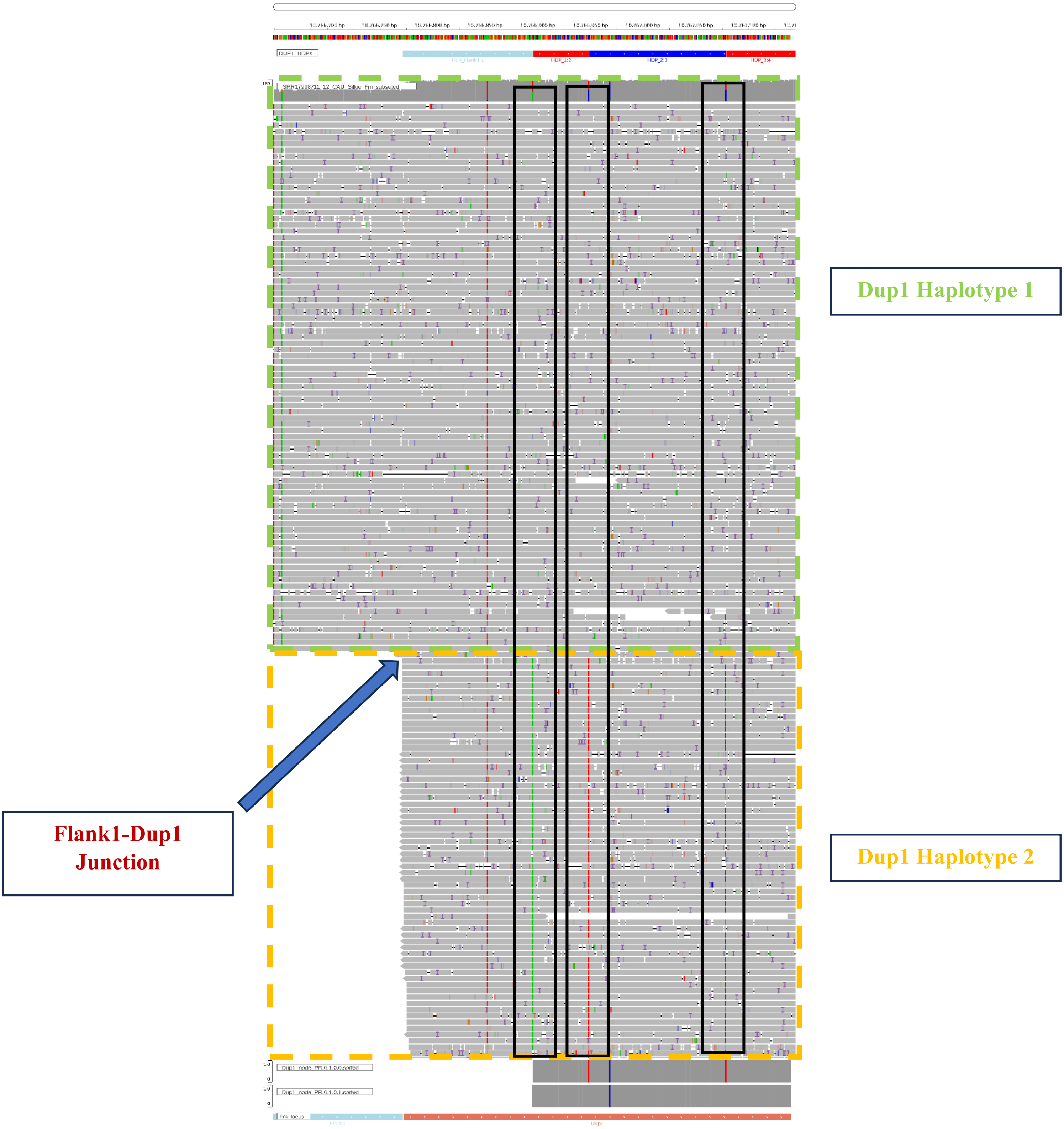


**Supplementary Fig. S4a:** This figure displays two sets of haplotype reads at the Flank1-Dup1 junction in CAU Silkie. Reads from Dup1 haplotype 2 show soft clipping at the junction. Three HDPs are highlighted within black vertical boxes.

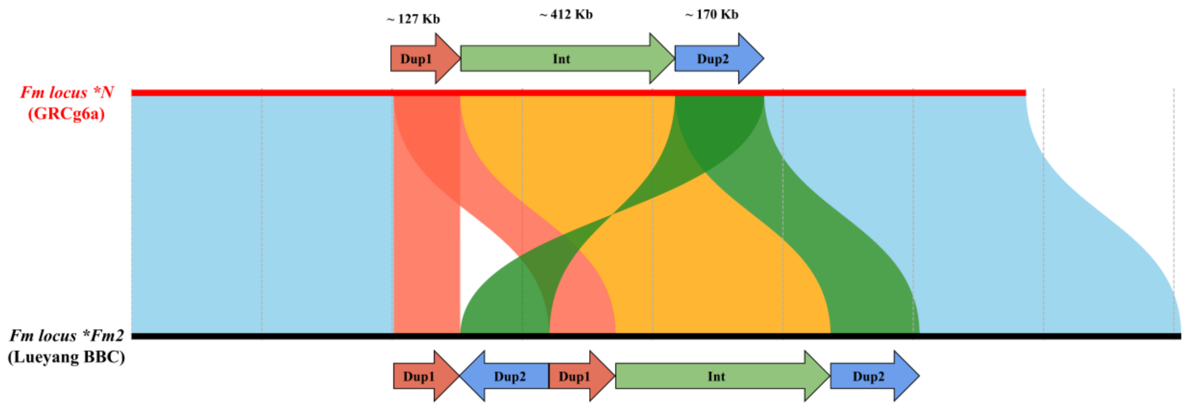


**Supplementary Figure. S5: Organization of the *Fm* Locus in GRCg6a and Lueyang Genome Assemblies:** The genomic organization of the *Fm* locus is elucidated through a visual depiction of the pairwise genome alignment between the GRCg6a reference and the Lueyang black-bone chicken genome at the *Fm* locus. This alignment was performed using minimap2, syri, and plotsr. The figure highlights the Dup1, Int, and Dup2 regions to emphasize the **Fm_2* arrangement. Identifying these regions provides insight into the structural organization and variation present in the *Fm* locus between the two genome assemblies.

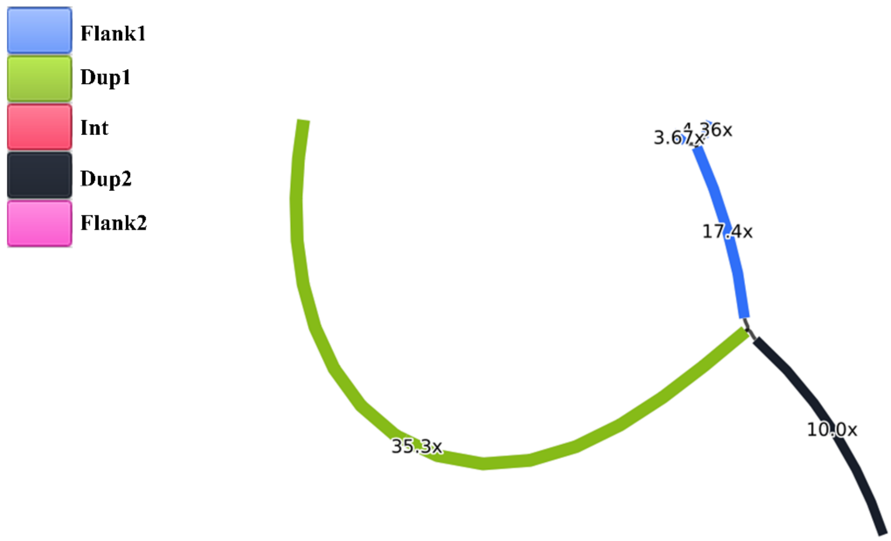


**Supplementary Figure. S6:** The de novo assembly of the Dup1 region, using Shasta, does not resolve the two haplotypes. Edges are color-coded based on BLAST-based annotation. The graph has been visualized using Bandage.

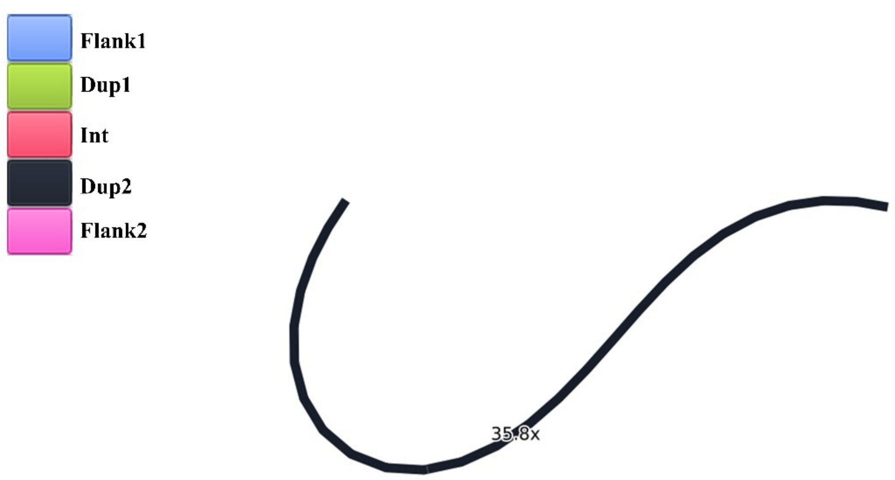


**Supplementary Figure. S7:** The de novo assembly of the Dup2 region, using Shasta, does not resolve the two haplotypes. Edges are color-coded based on BLAST-based annotation. The graph has been visualized using Bandage.

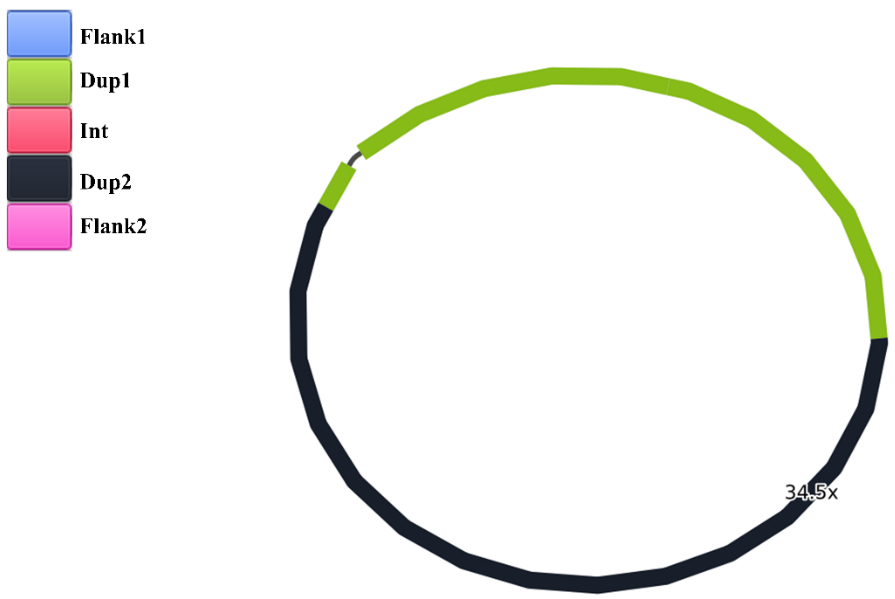


**Supplementary Figure. S8:** The de novo assembly of the Dup1 and Dup2 region together, using Shasta, does not resolve the haplotypes of Dup1 and Dup2. Edges are color-coded based on BLAST-based annotation. The graph has been visualized using Bandage.

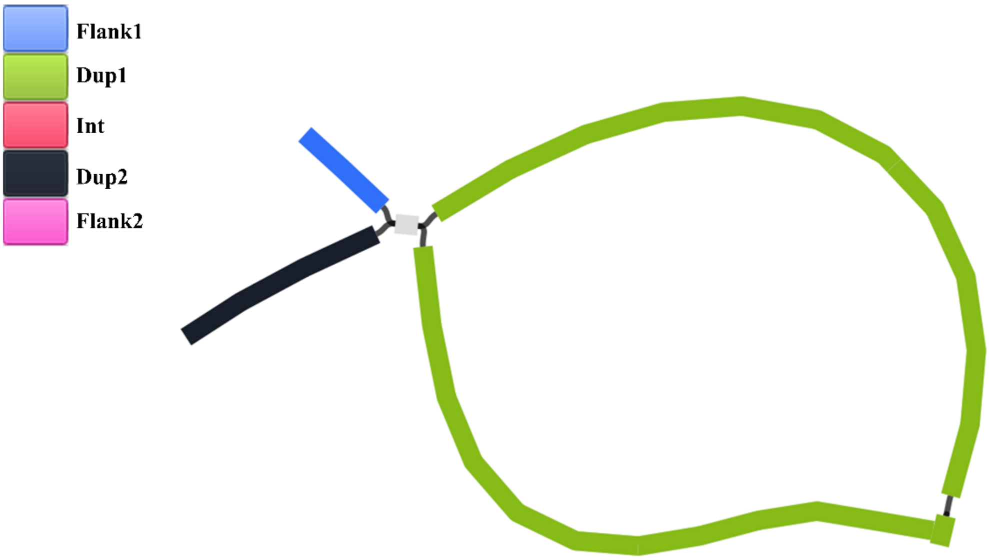


**Supplementary Figure. S9:** The de novo assembly of the Dup1 region using Shasta resolves the two haplotypes within the Dup1 region in mode 2. Edges are color-coded based on BLAST-based annotation. However, the ends of the Dup1 region remain unresolved, particularly in the HDP-poor region. The graph has been visualized using Bandage.

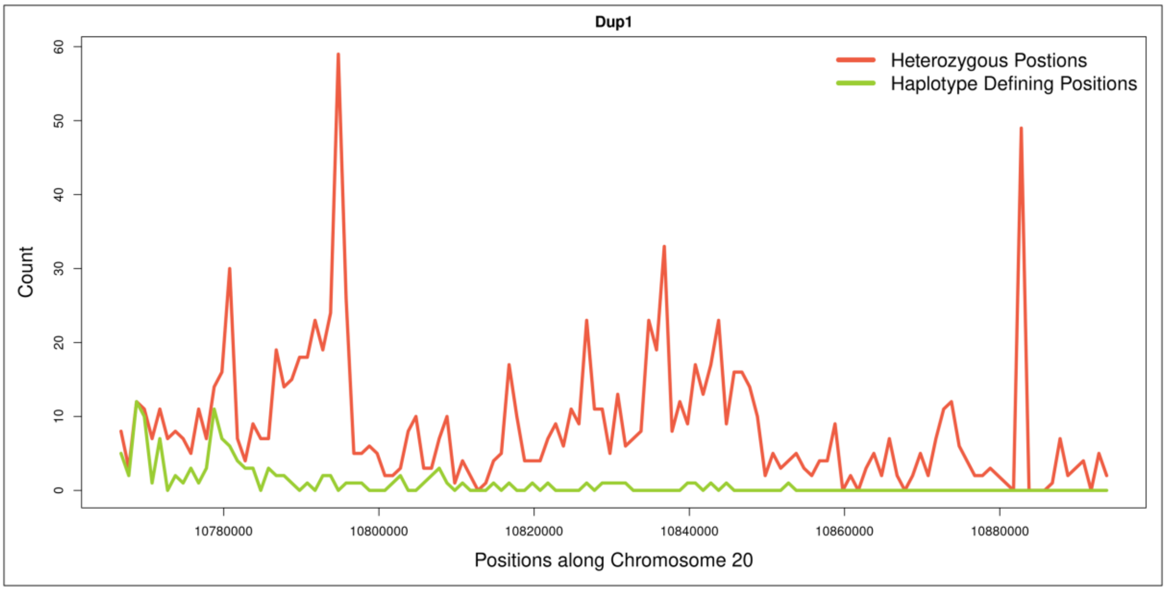


**Supplementary Figure. S9a:** Distribution of heterozygous and Haplotype Defining Positions (HDPs) across Dup1. Compared to Dup1 start, Dup1 end is a HDP poor region.

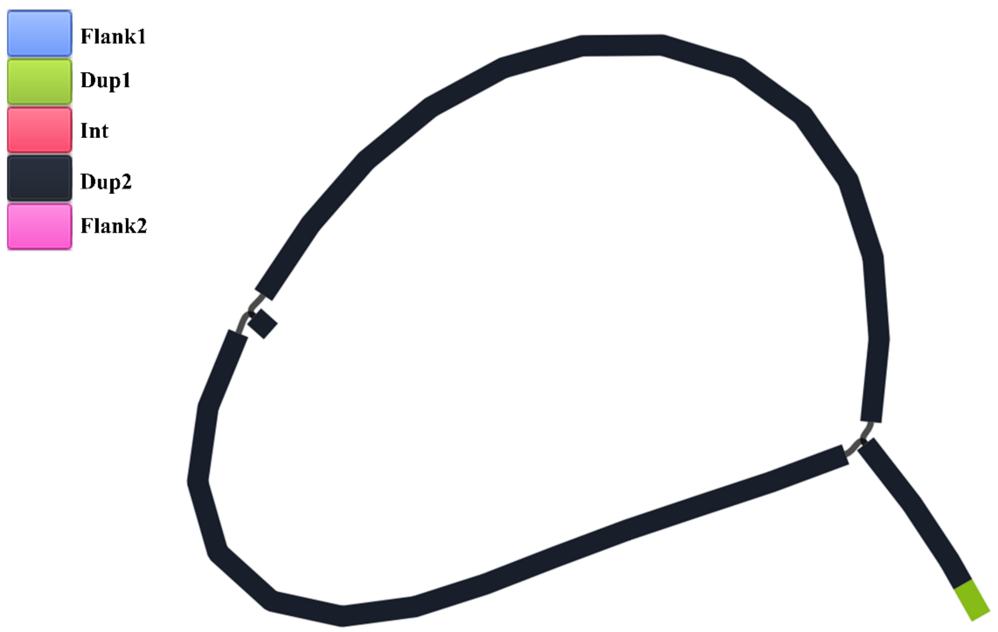


**Supplementary Figure. S10:** The de novo assembly of the Dup2 region using Shasta resolves the two haplotypes within the Dup2 region in mode 2. Edges are color-coded based on BLAST-based annotation. However, the Start of the Dup2 region remains unresolved, particularly in the HDP-poor region. The graph has been visualized using Bandage.

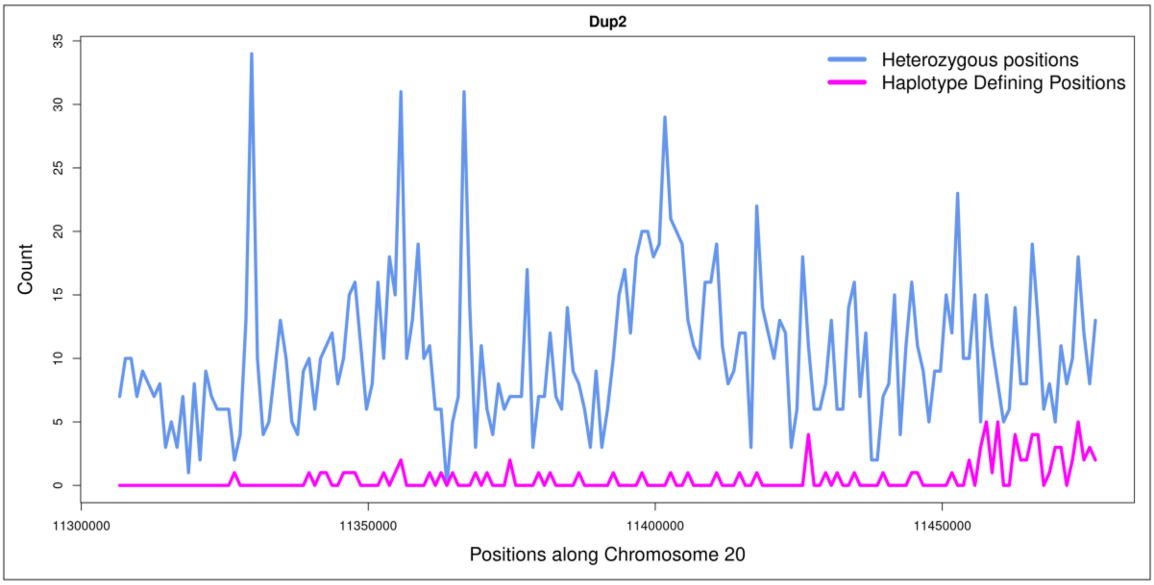


**Supplementary Figure. S10a:** Distribution of heterozygous and Haplotype Defining Positions (HDPs) across Dup1. Compared to the Dup2 end, the Dup2 start is a HDP poor region.

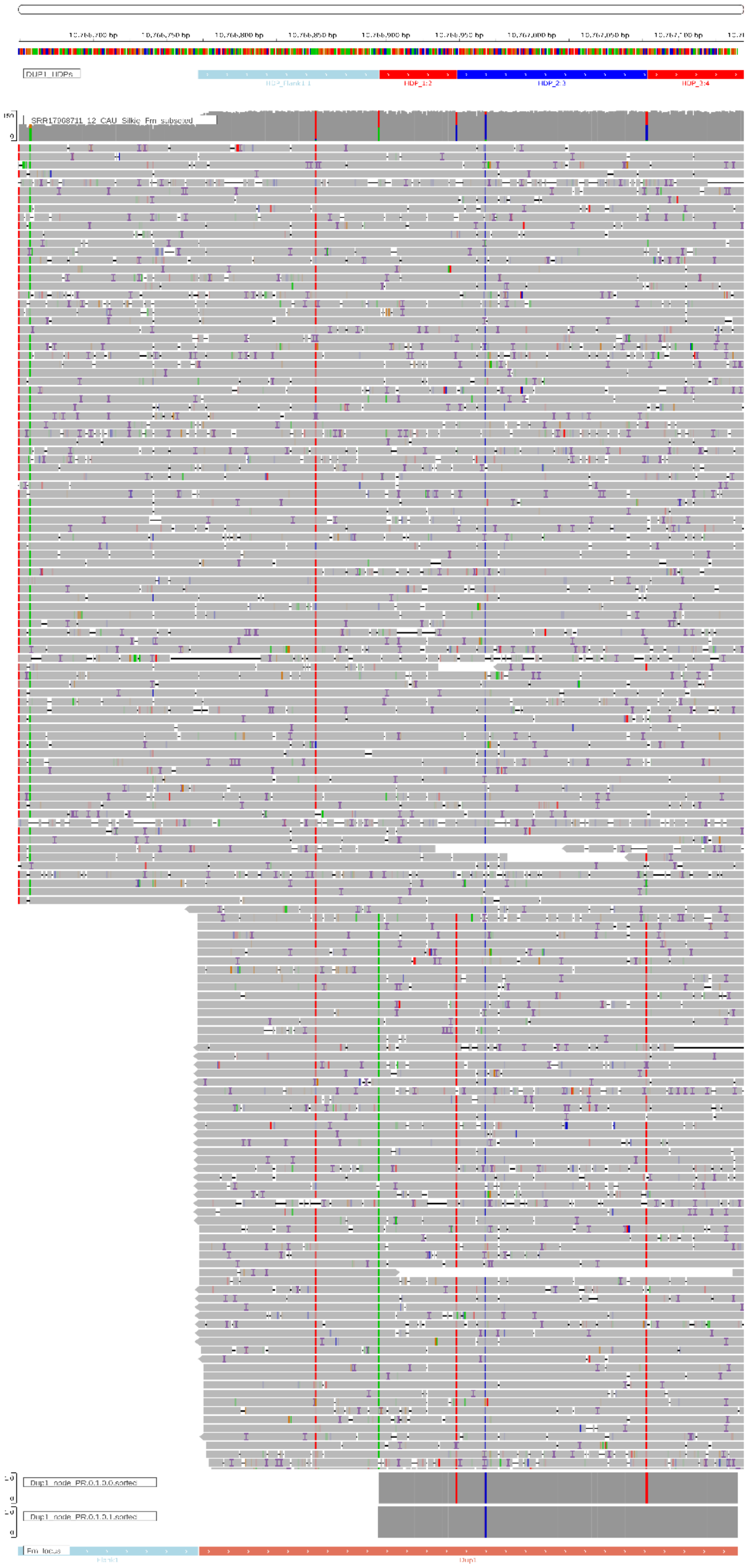

**Supplementary Figure. S11:** The Dup1 edge PR.0.1.0.1 assembled in Shasta starts at position 10766895, while Dup2 edge PR.0.1.0.0 starts at position 10766896, as shown in the lower panel.

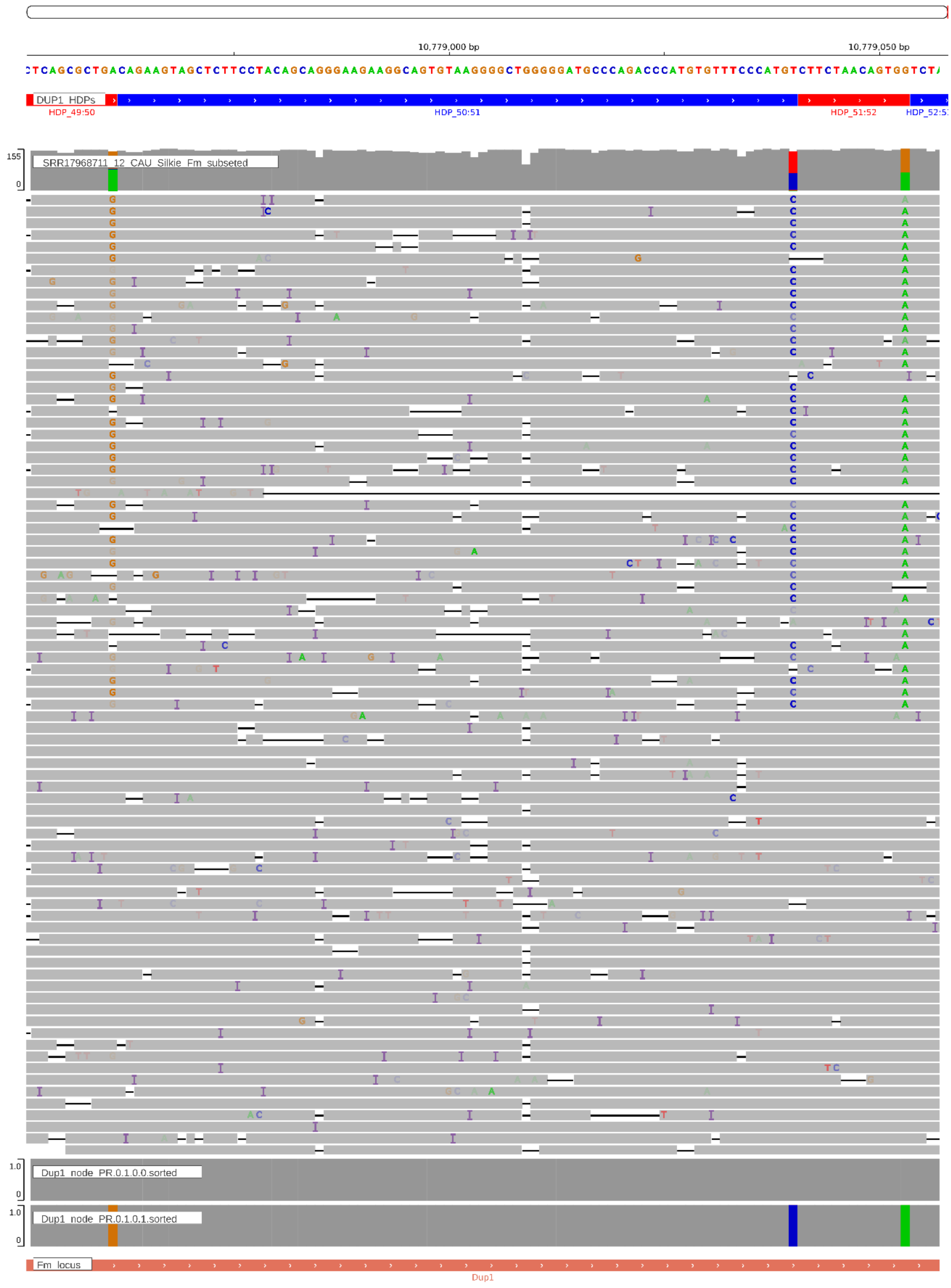


**Supplementary Figure. S12:** IGV screenshot of the 2 HDPs present in the mid-region of Dup1. The edges assembled in Shasta are shown in the lower panel.

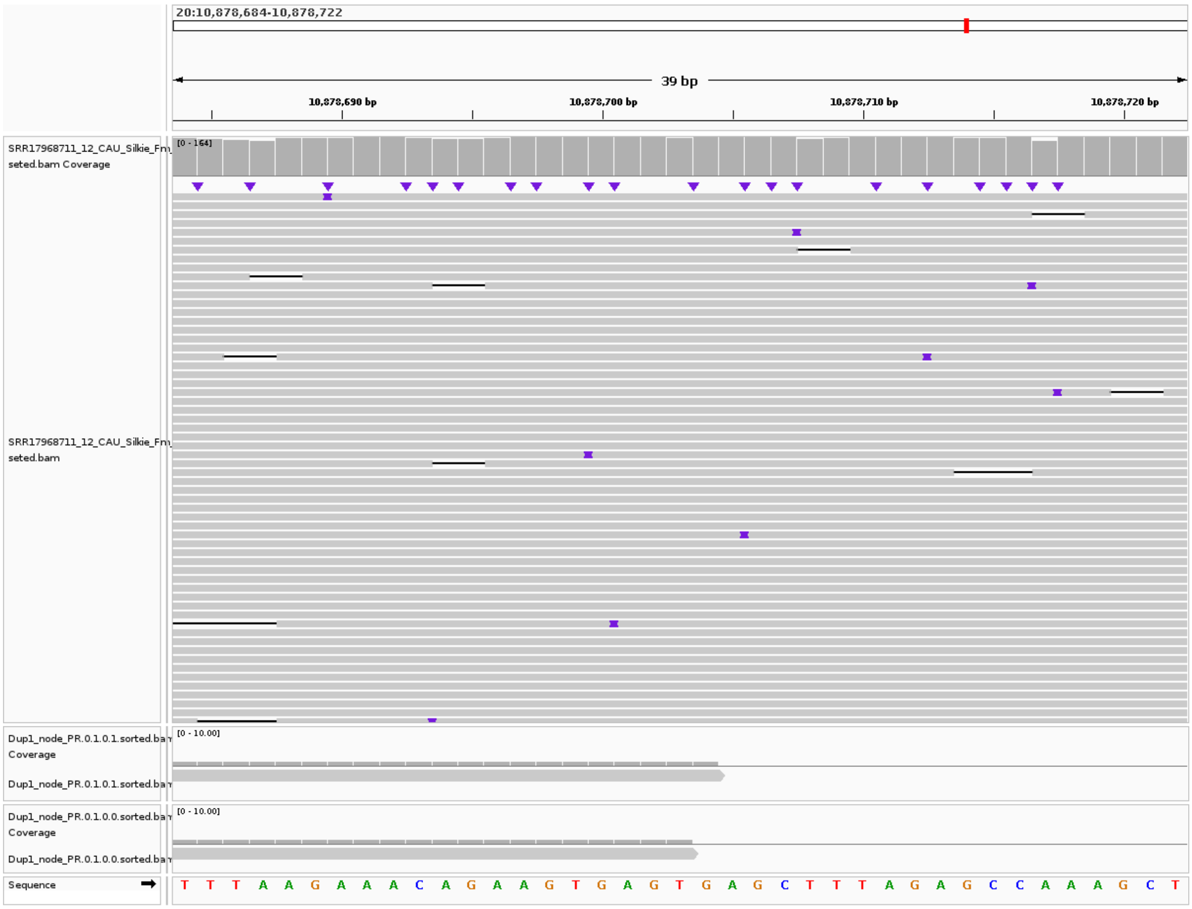


**Supplementary Figure. S13:** The Dup1 edge PR.0.1.0.1 assembled in Shasta ends at position 10878704, while Dup2 edge PR.0.1.0.0 starts at position 10878703, as shown in the lower panel.

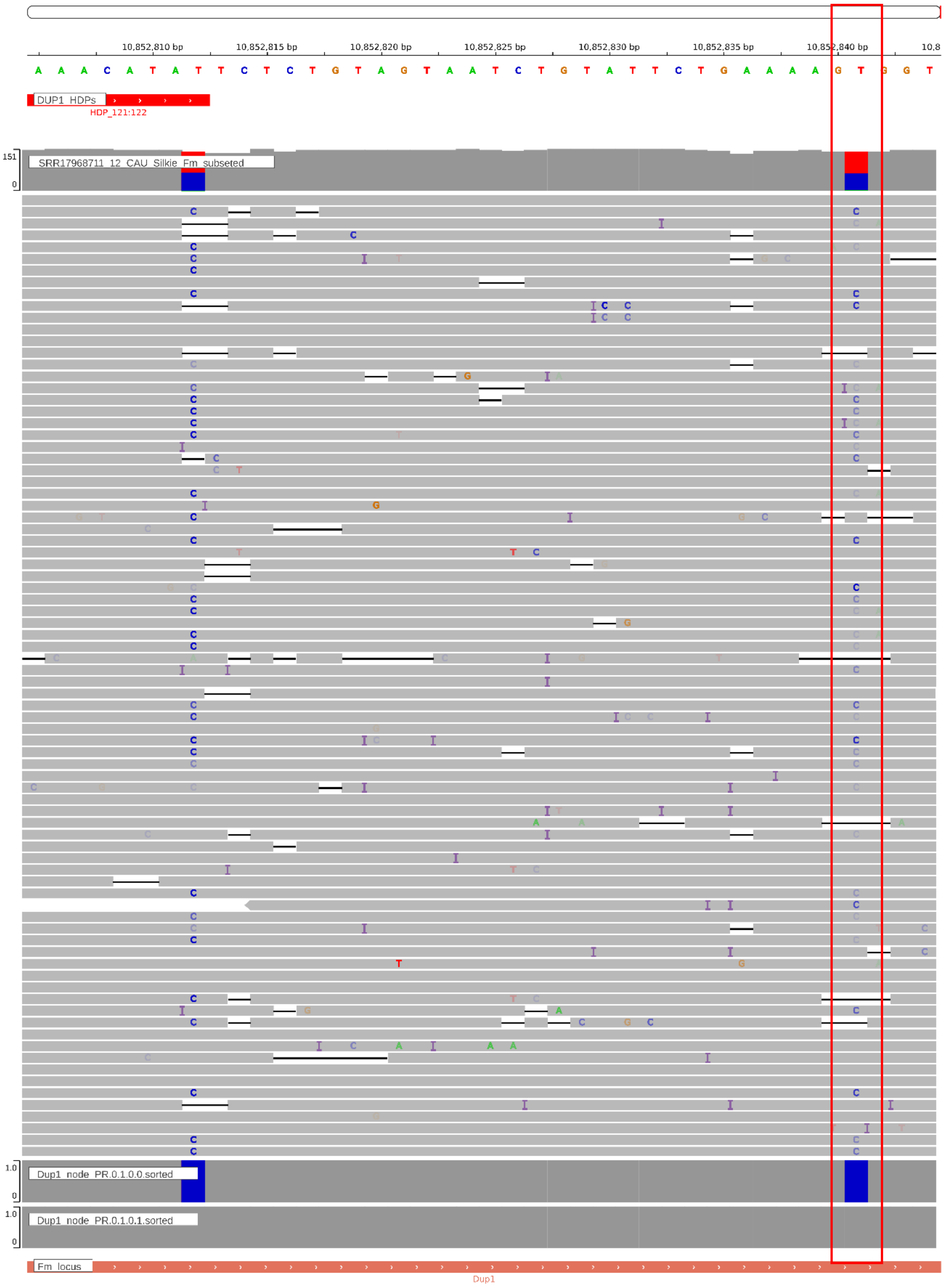


**Supplementary Figure. S14:** IGV screenshot of the last HDP present at the Dup1. The edges assembled in Shasta span beyond the last HDP are shown in the lower panel. The previously unidentified HDP at 10852841 is shown in a red vertical box.

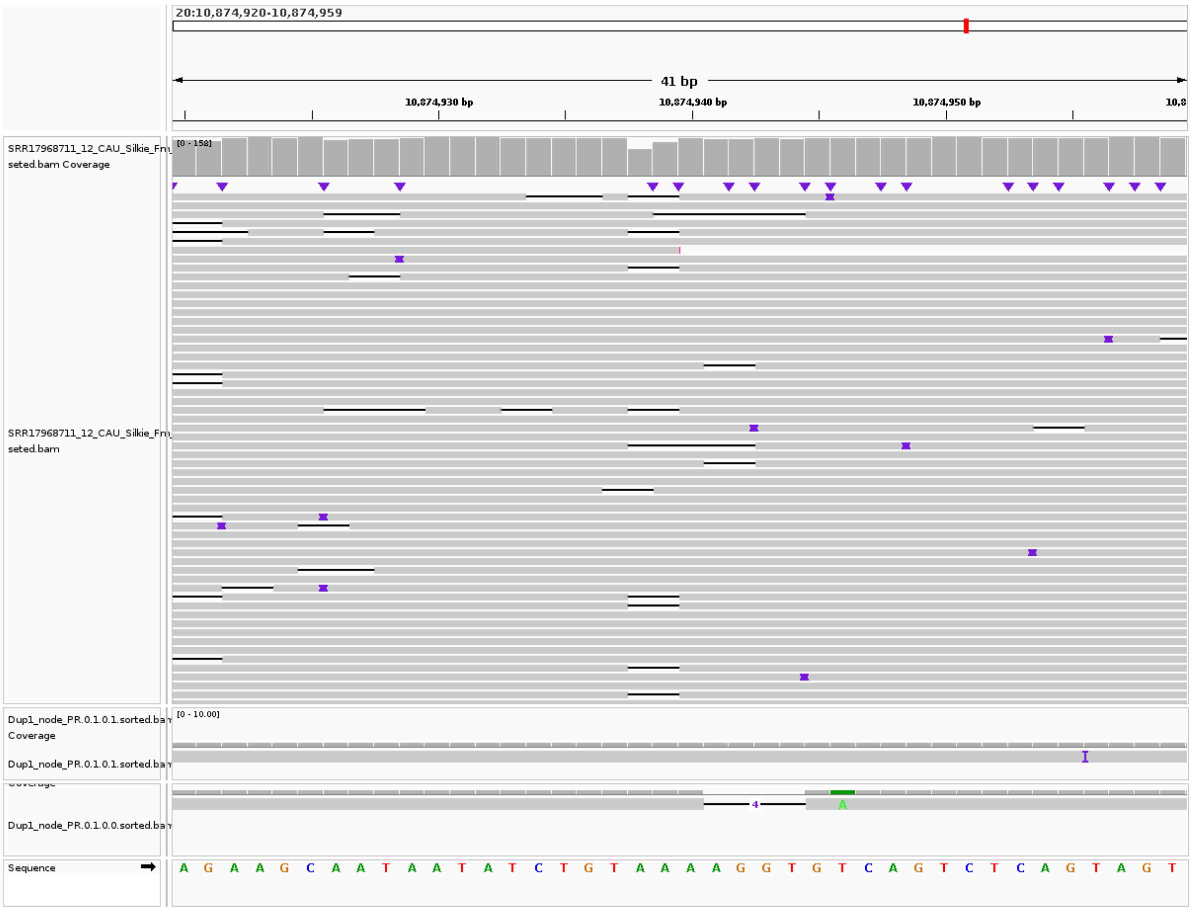


**Supplementary Figure. S15:** IGV screenshot of the Dup1 edges assembled in Shasta, showing a collapse. All reads at position 10874946 are homozygous, while Shasta assembled two haplotypes.

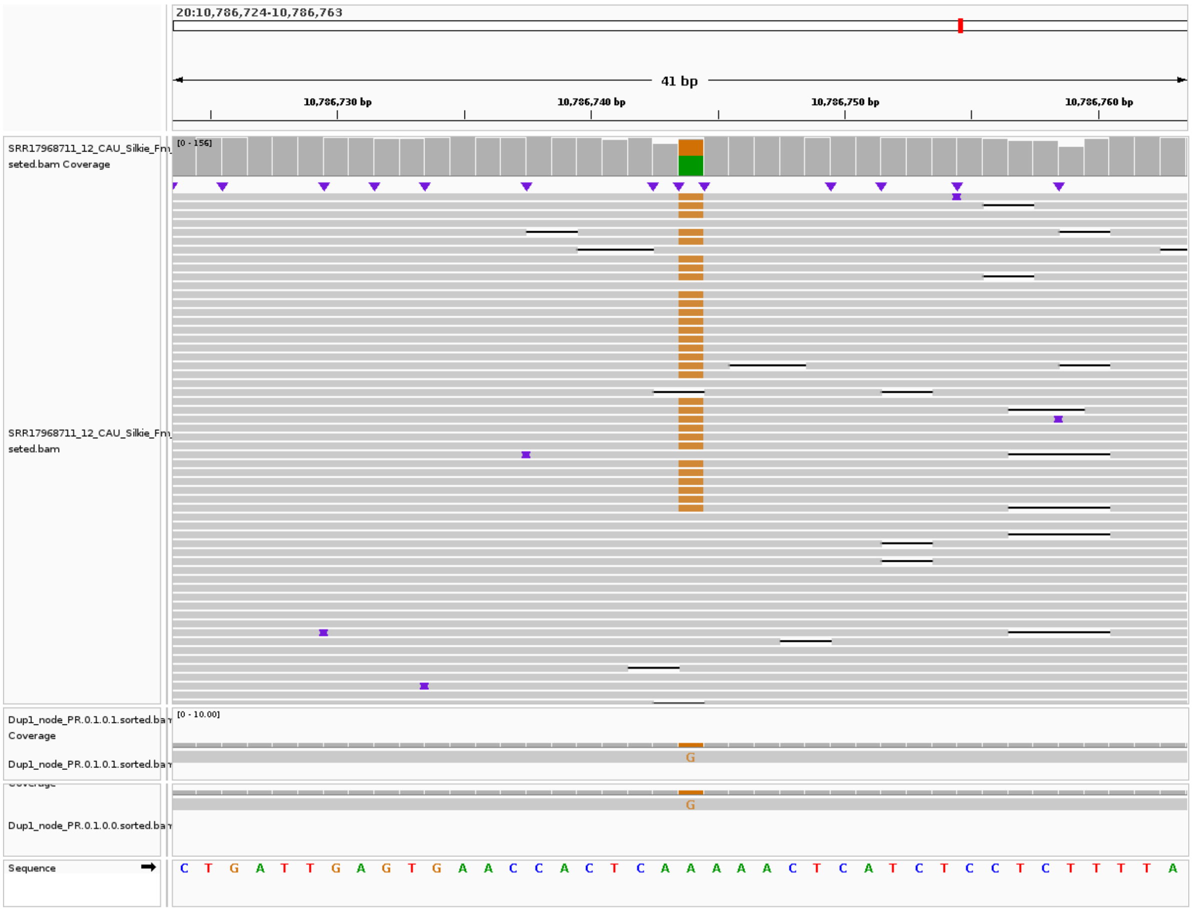


**Supplementary Figure. S16:** IGV screenshot of the Dup1 edges assembled in Shasta, showing a collapse. Despite the comparable proportion of the two haplotypes in the raw reads at position 10786744, Shasta assembled only one haplotype.

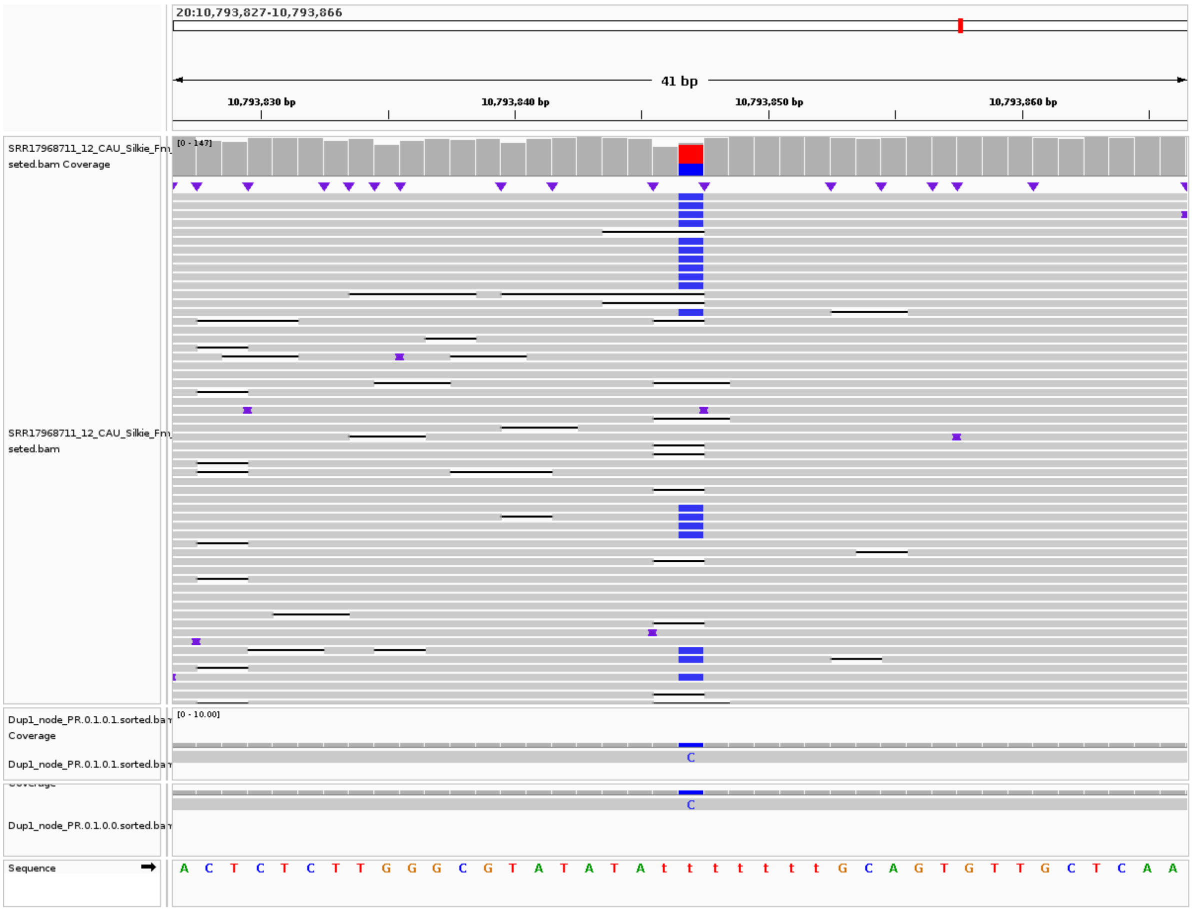


**Supplementary Figure. S17:** IGV screenshot of the Dup1 edges assembled in Shasta, showing a collapse. Despite the comparable proportion of the two haplotypes in the raw reads at position 10793847, Shasta assembled only one haplotype.

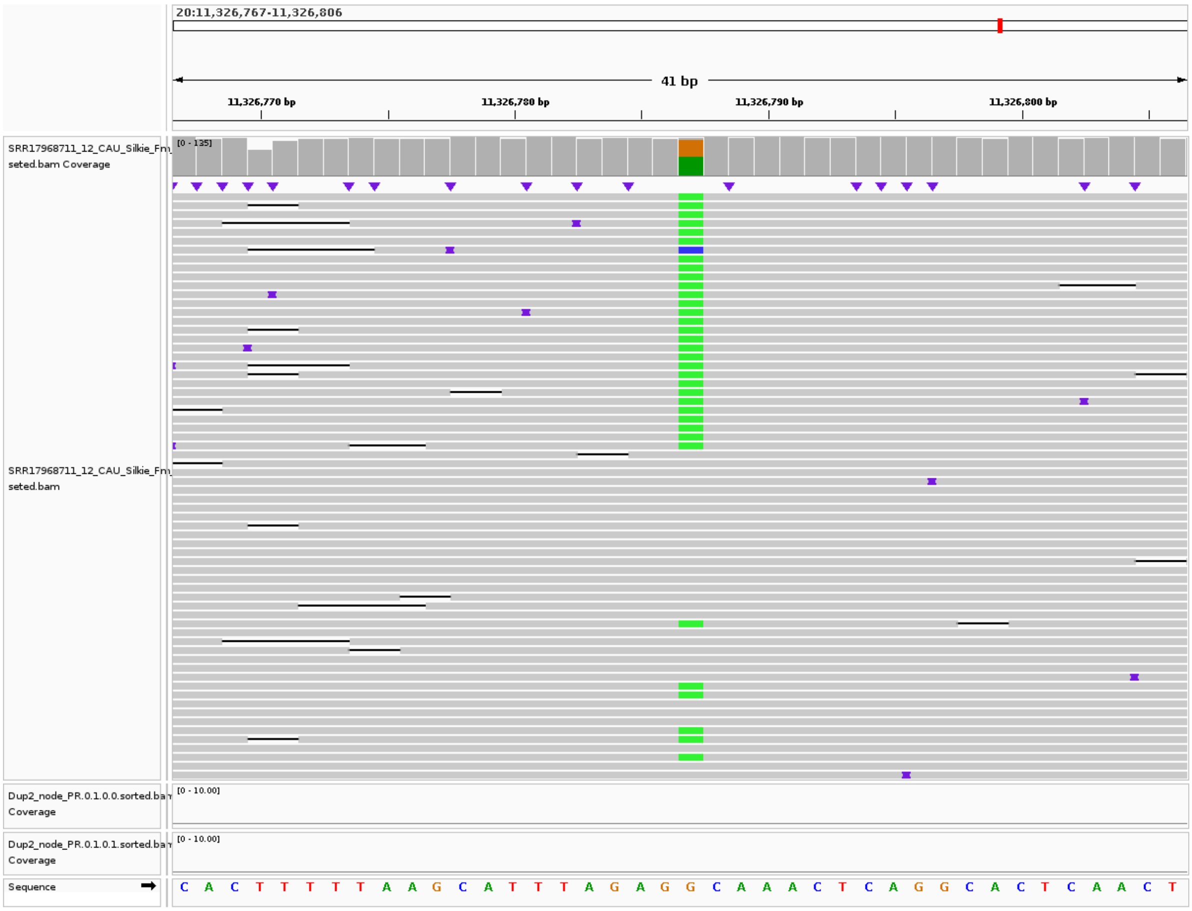


**Supplementary Figure. S18:** IGV screenshot of the first HDP present at the Dup2 start. The edges assembled in Shasta do not span the first HDP.

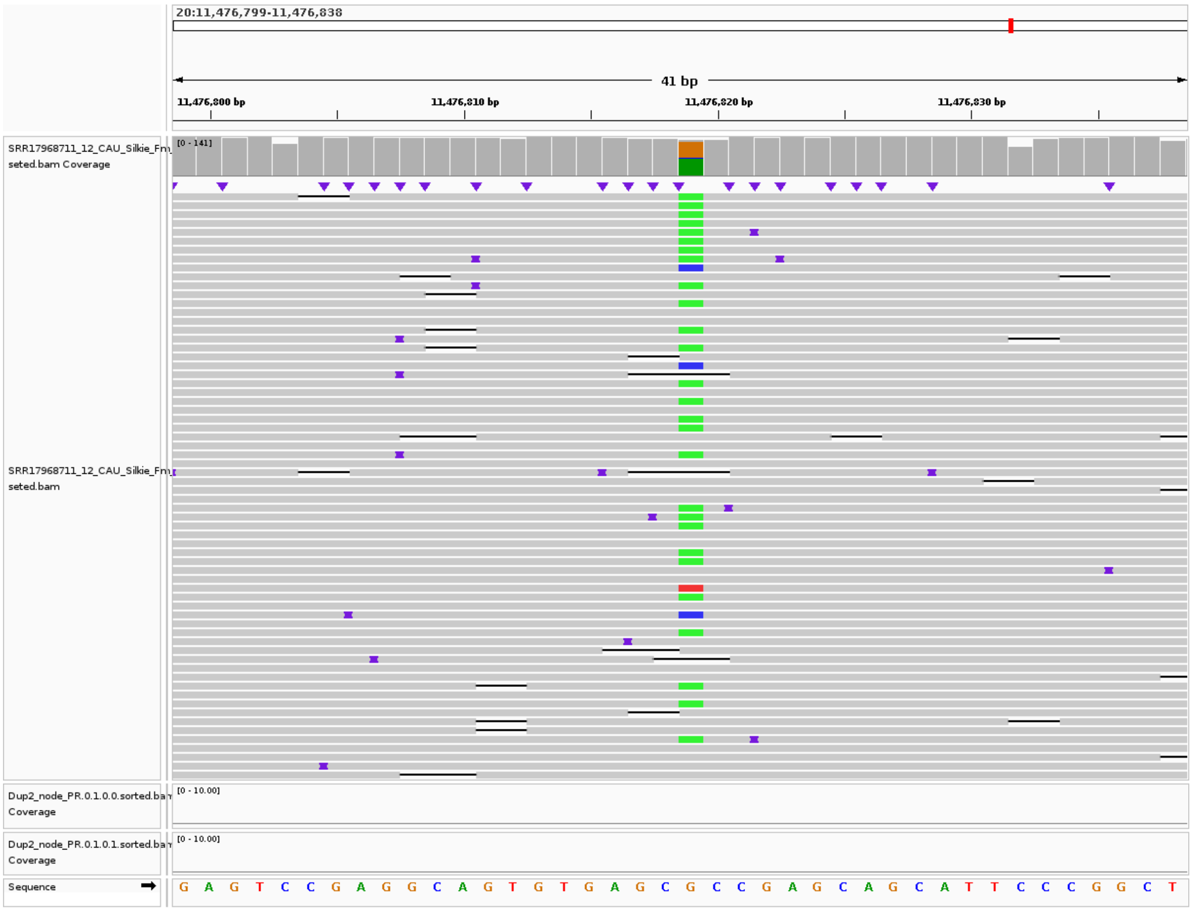


**Supplementary Figure. S19:** IGV screenshot of the last HDP present at the Dup2 end. The edges assembled in Shasta do not span the last HDP.

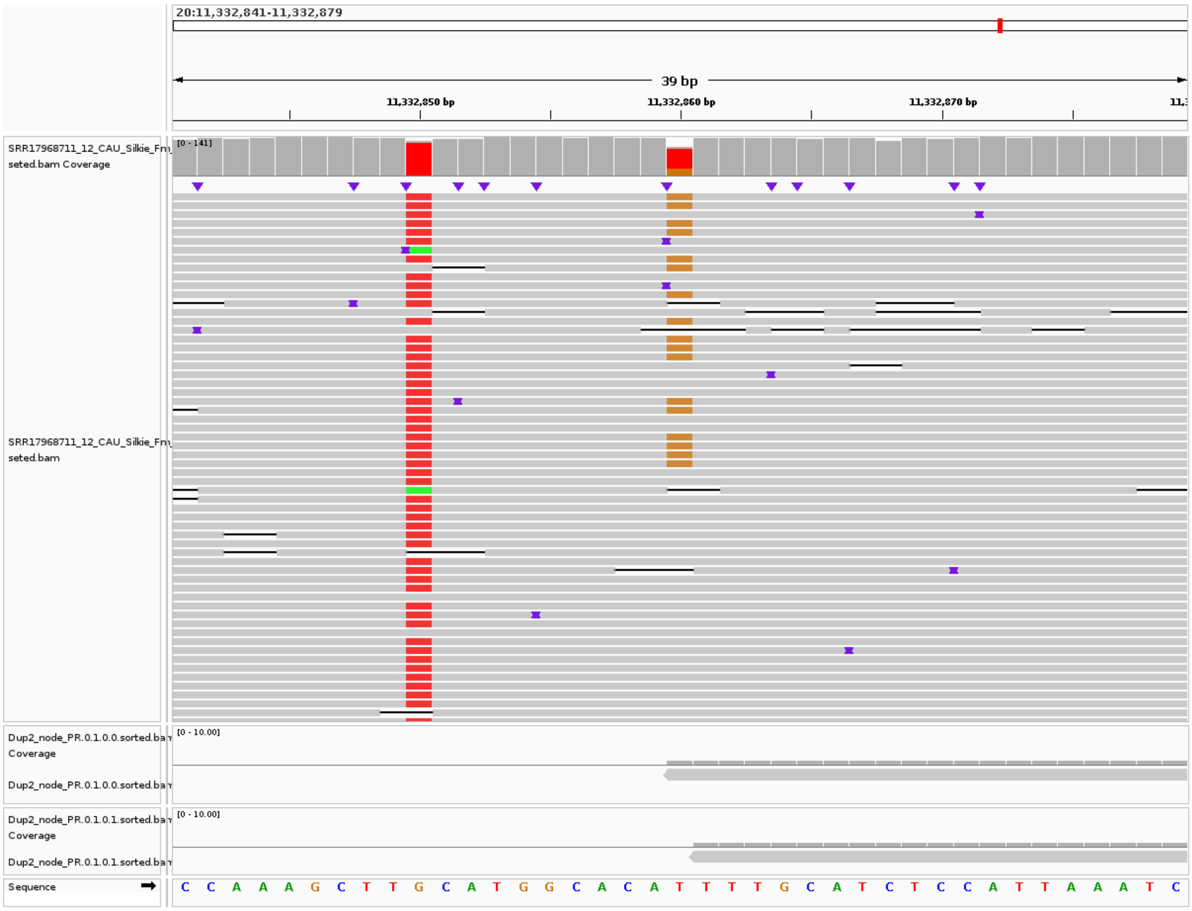


**Supplementary Figure. S20:** The Dup2 edge PR.0.1.0.0 assembled in Shasta starts at position 11,332,860, while Dup2 edge PR.0.1.0.1 starts at position 11,332,861, as shown in the lower panel.

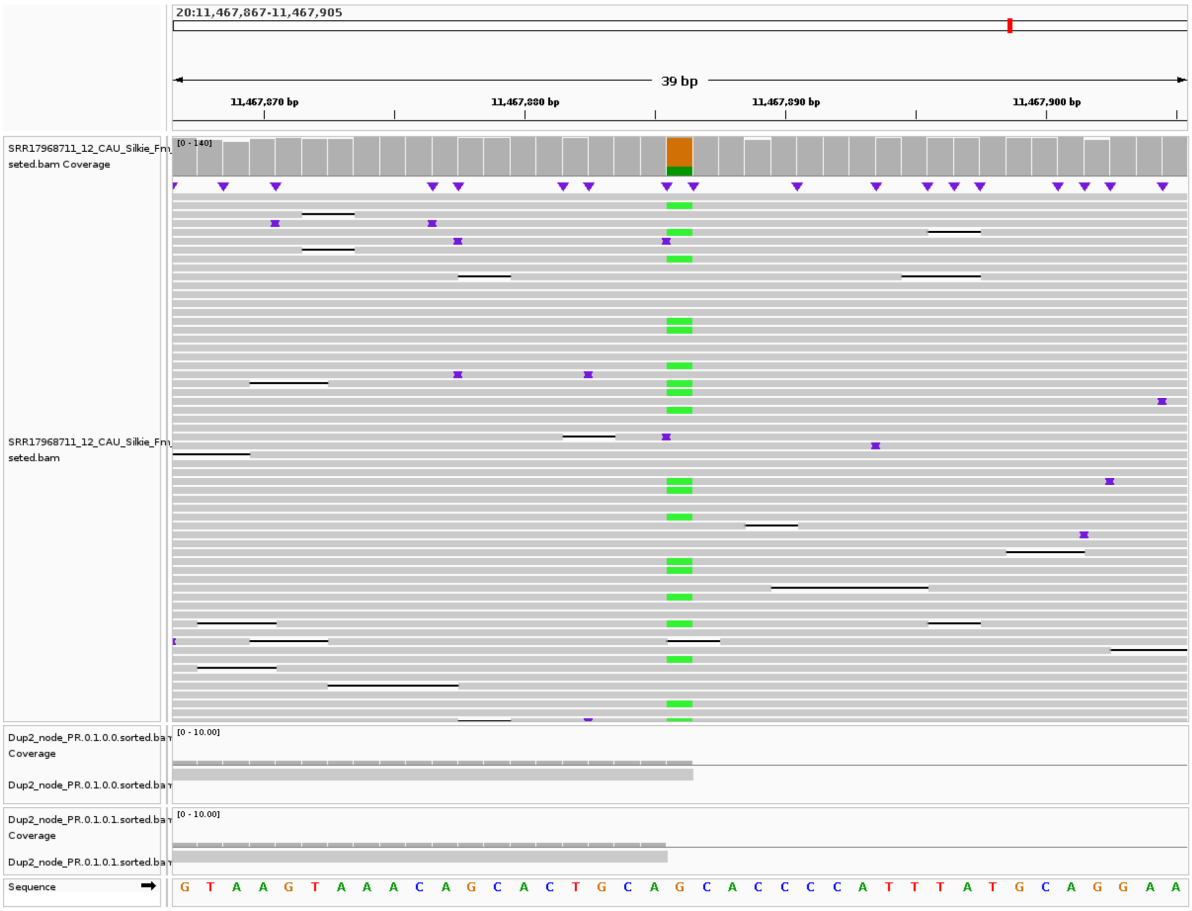


**Supplementary Figure. S21:** The Dup2 edge PR.0.1.0.0 assembled in Shasta ends at position 11467886, while Dup2 edge PR.0.1.0.1 starts at position 11467885, as shown in the lower panel.

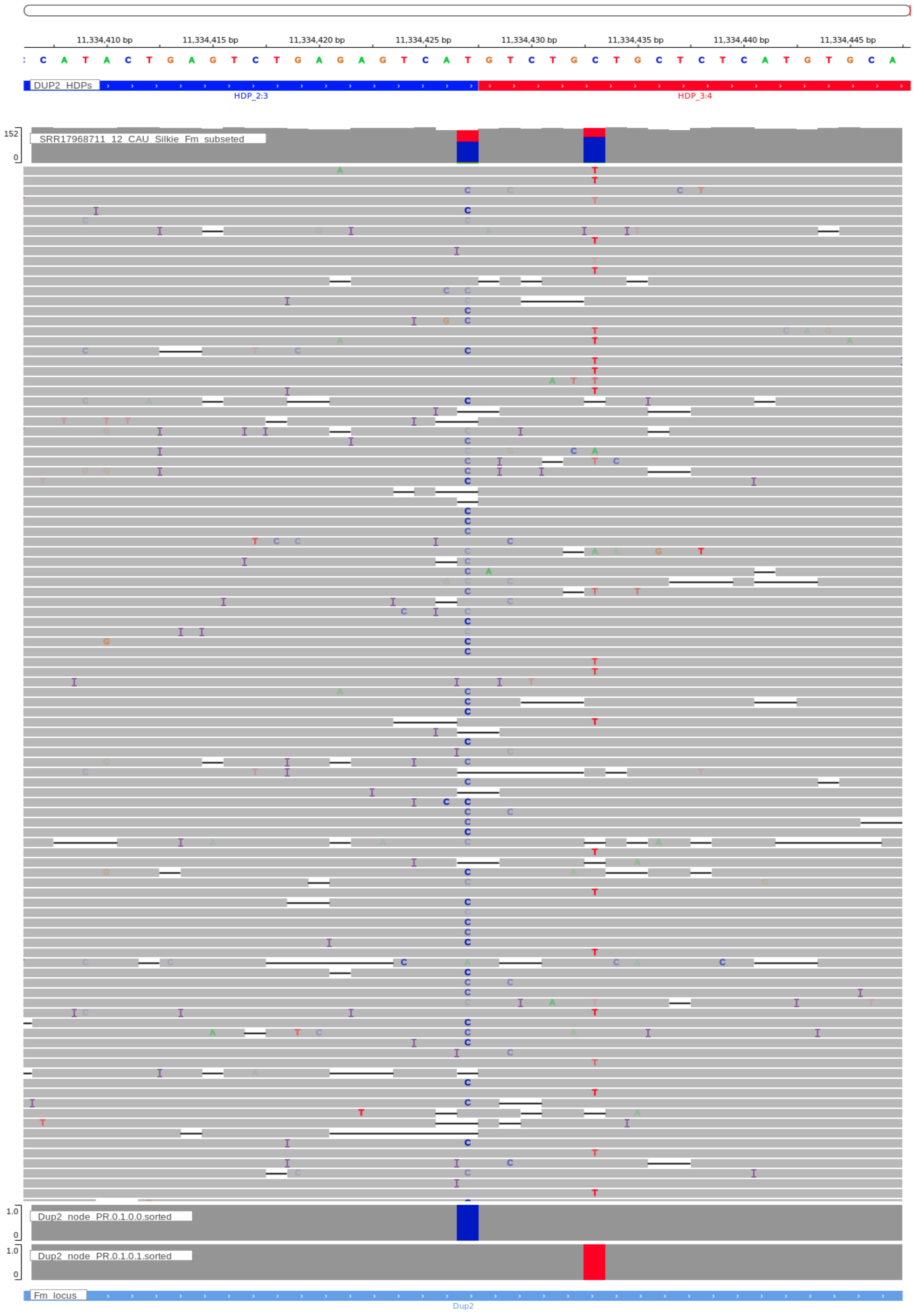


**Supplementary Figure. S21a:** The Dup2 edges PR.0.1.0.0 and PR.0.1.0.1 assembled in Shasta at position 11334427 (third HDP of Dup2). Dup2 region differs between the two assembled edges.

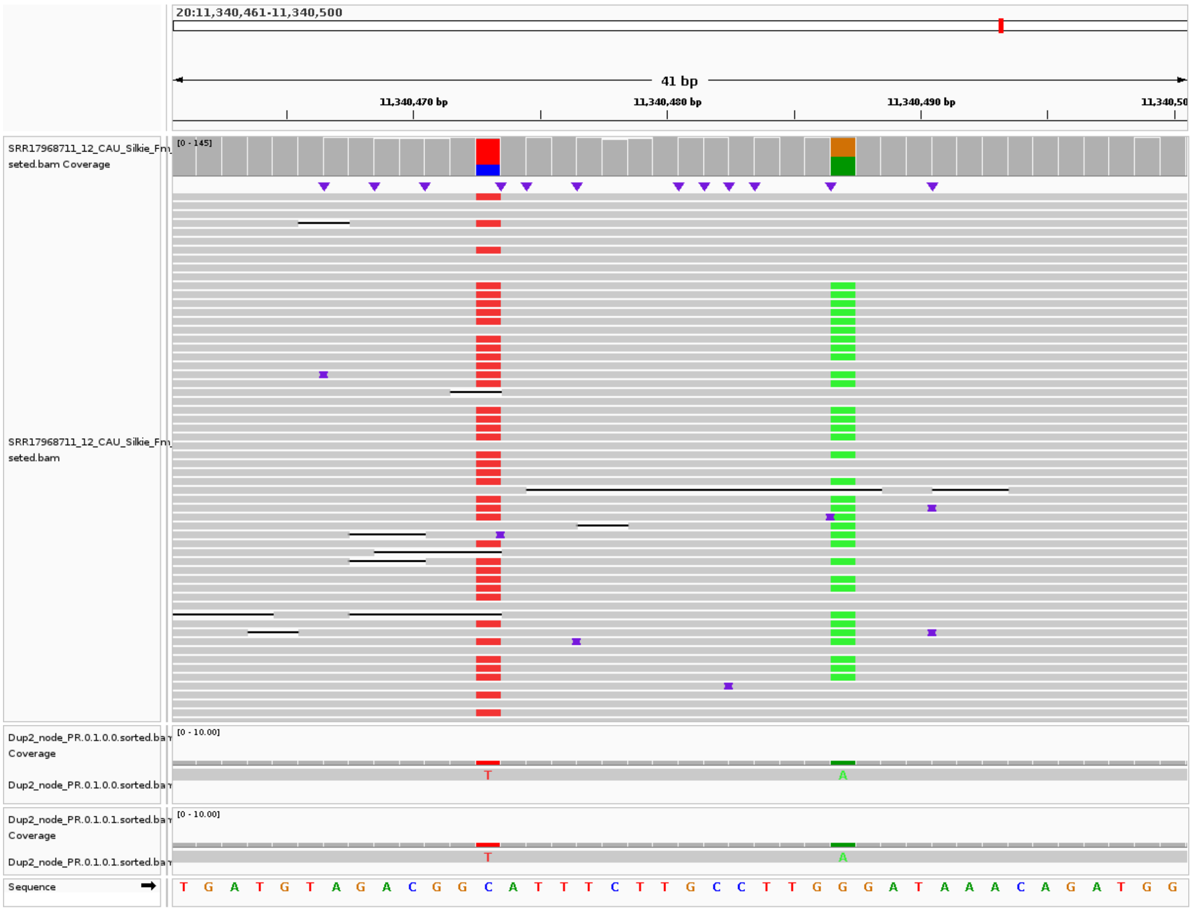


**Supplementary Figure. S22:** IGV screenshot of the Dup2 edges assembled in Shasta, showing a collapse. Despite the comparable proportion of the two haplotypes in the raw reads at position 11340487, Shasta assembled only one haplotype.

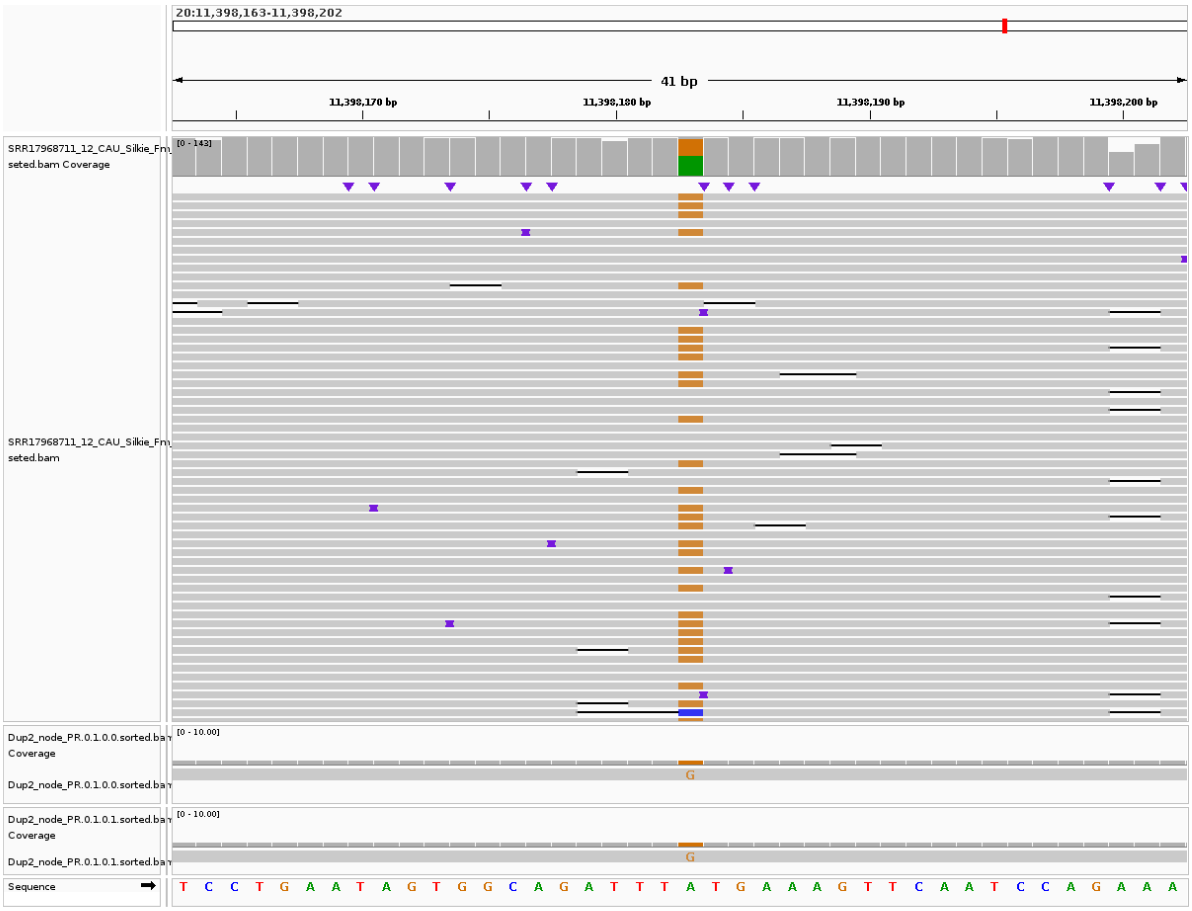


**Supplementary Figure. S23:** IGV screenshot of the Dup2 edges assembled in Shasta, showing a collapse. Despite the comparable proportion of the two haplotypes in the raw reads at position 11398183, Shasta assembled only one haplotype.

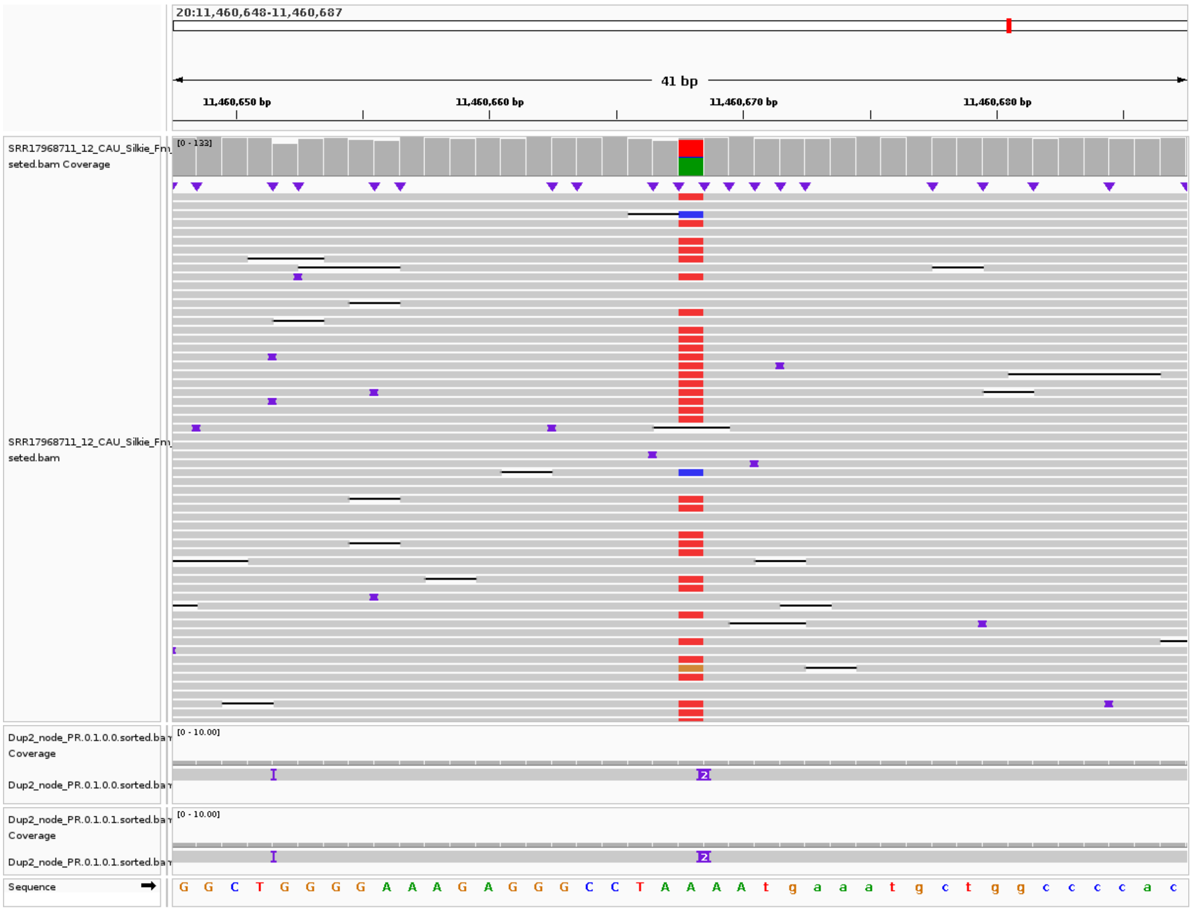


**Supplementary Figure. S24:** IGV screenshot of the Dup2 edges assembled in Shasta, showing a collapse. Despite the comparable proportion of the two haplotypes in the raw reads at position 11460668, Shasta assembled only one haplotype.

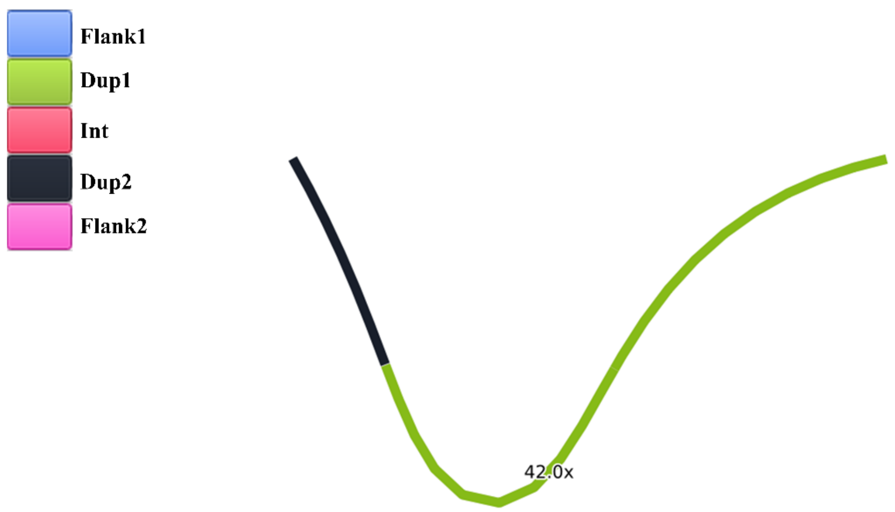


**Supplementary Figure. S25:** The de novo assembly of the Dup1 region, using Raven, does not resolve the two haplotypes. Edges are color-coded based on BLAST-based annotation. The graph has been visualized using Bandage.

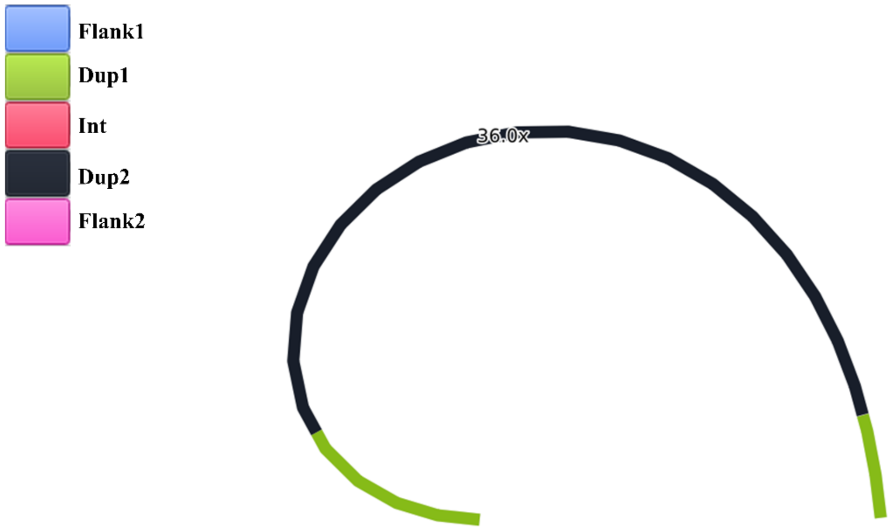


**Supplementary Figure. S26:** The de novo assembly of the Dup2 region, using Raven, does not resolve the two haplotypes. Edges are color-coded based on BLAST-based annotation. The graph has been visualized using Bandage.

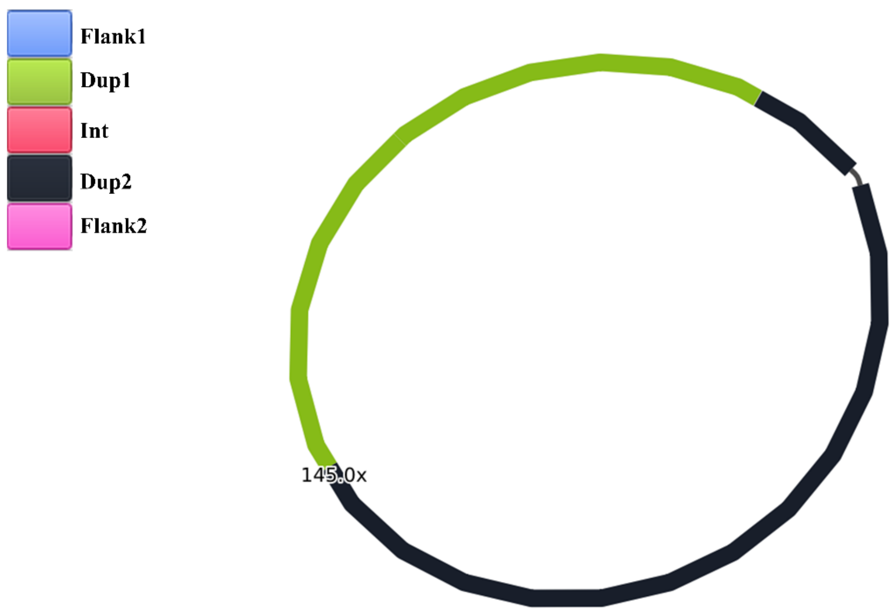


**Supplementary Figure. S27:** The de novo assembly of the Dup1 and Dup2 region together, using Raven, does not resolve the haplotypes of Dup1 and Dup2. Edges are color-coded based on BLAST-based annotation. The graph has been visualized using Bandage.

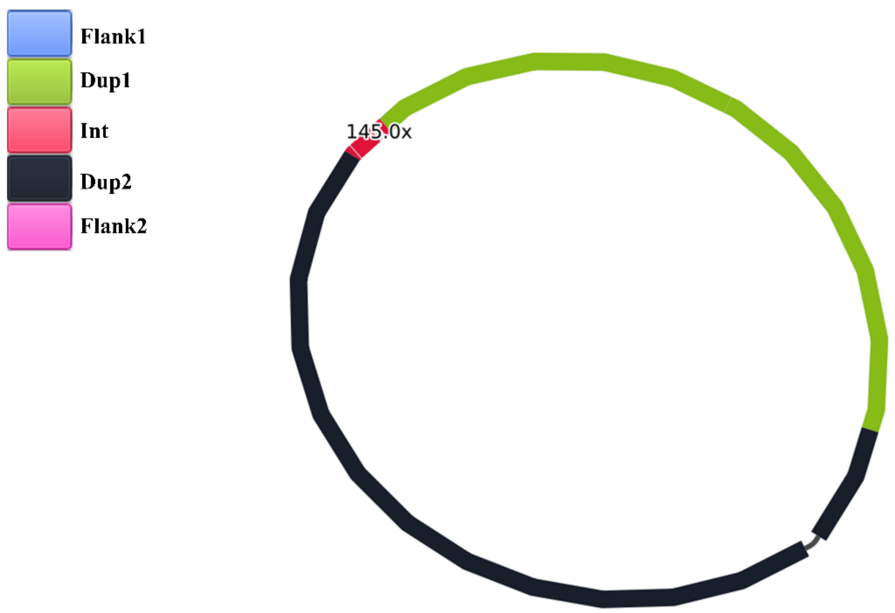


**Supplementary Figure. S28:** The de novo assembly of the Dup1, Dup2, and Int region together, using Raven, does not resolve the haplotypes of Dup1 and Dup2. Edges are color-coded based on BLAST-based annotation. The graph has been visualized using Bandage.

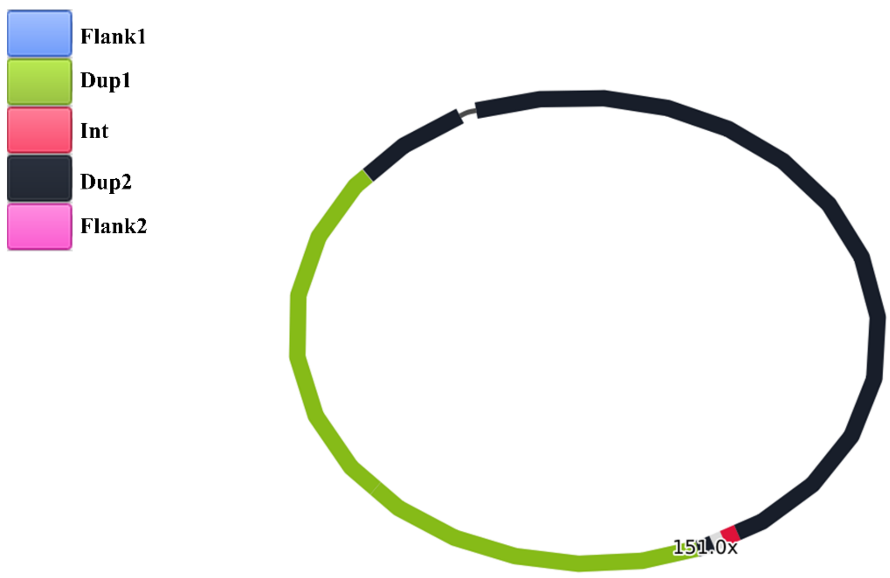


**Supplementary Figure. S29:** The de novo assembly of the entire Fm locus, using Raven, does not resolve the haplotypes of Dup1 and Dup2. Edges are color-coded based on BLAST-based annotation. The graph has been visualized using Bandage.

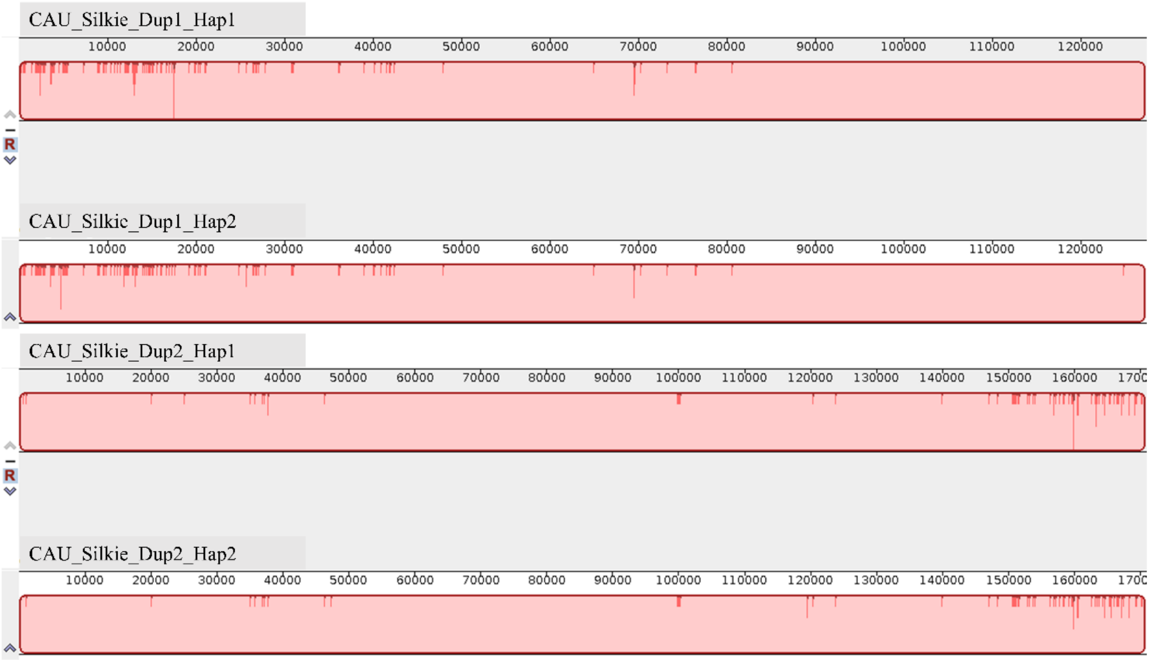


**Supplementary Figure. S30: Alignment of Dup1 and Dup2 Haplotypes assembled in CAU Silkie Genome:** The alignment of Dup1 and Dup2 haplotypes assembled in the CAU Silkie genome was conducted using progressiveMauve. Both haplotypes exhibit a remarkably high sequence similarity, exceeding 99%. Identical sequences are highlighted in pink, while mismatches within a 13-mer window size are indicated in red.

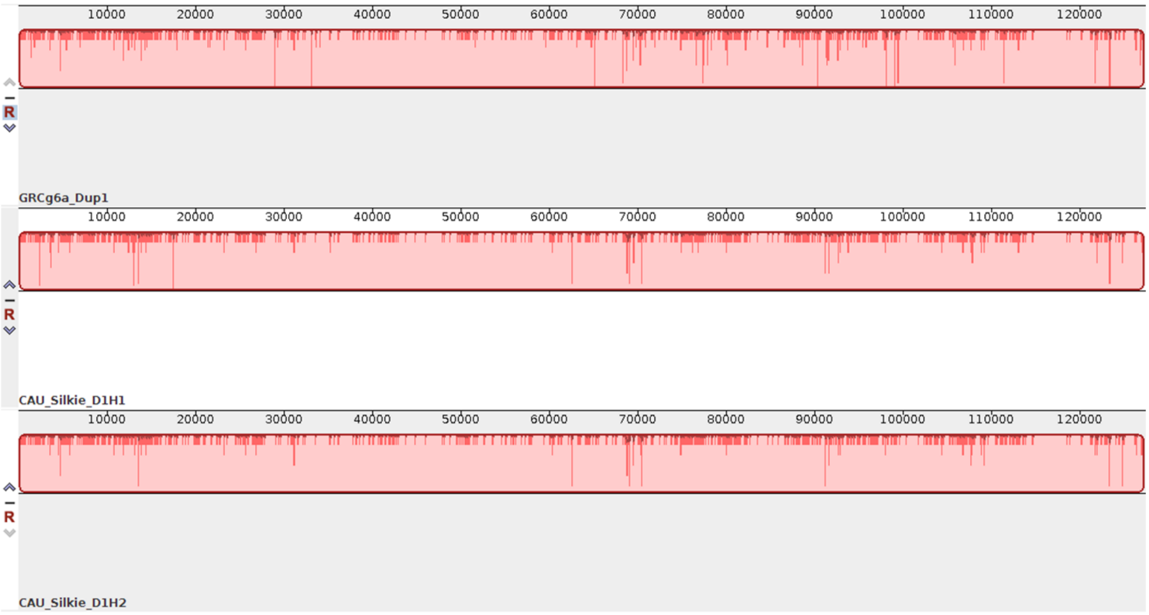


**Supplementary Figure. S31: Alignment of the Dup1 Region from the GRCg6a and CAU Silkie Genome Assemblies:** This figure illustrates the alignment of the Dup1 region, assembled in the GRCg6a genome assembly, with both haplotypes of Dup1 assembled in the CAU Silkie genome. The alignment was performed using progressiveMauve. The comparison reveals a remarkably high sequence similarity, exceeding 99%, between the two haplotypes. Identical sequences are highlighted in pink and mismatches are indicated in red within a 13-mer window size.

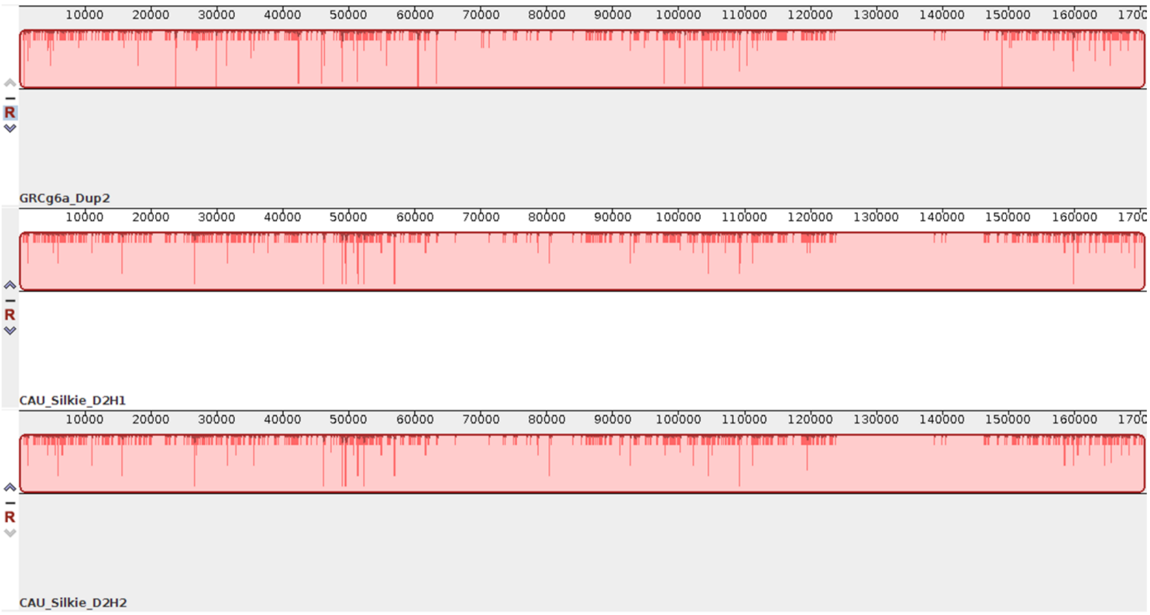


**Supplementary Figure. S32: Alignment of the Dup2 Region from the GRCg6a and CAU Silkie Genome Assemblies:** This figure illustrates the alignment of the Dup2 region, assembled in the GRCg6a genome assembly, with both haplotypes of Dup2 assembled in the CAU Silkie genome. The alignment was performed using progressiveMauve. The comparison reveals a remarkably high sequence similarity, exceeding 99%, between the two haplotypes. Identical sequences are highlighted in pink and mismatches are indicated in red within a 13-mer window size.

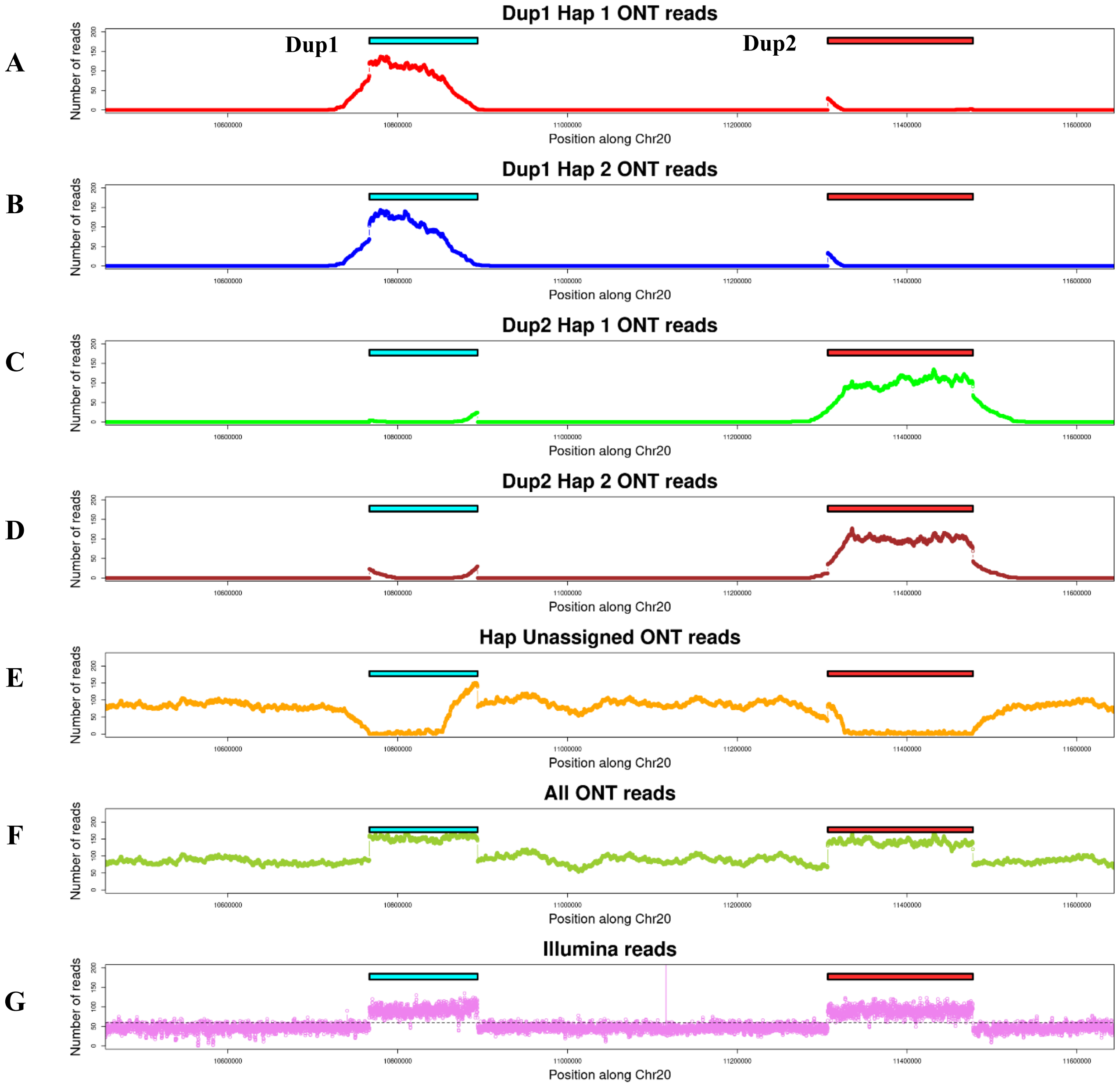


**Supplementary Figure. S33: Read coverage of CAU Silkie ONT and Illumina reads mapped to the GRCg6a genome assembly at *Fm* locus using minimap2 and bedtools coverage: (A)** Dup1 Hap1-specific ONT reads. **(B)** Dup1 Hap2-specific ONT reads. **(C)** Dup2 Hap1-specific ONT reads. **(D)** Dup2 Hap2-specific ONT reads. **(E)** Reads not assigned to any haplotype. **(F)** All ONT reads. **(G)** Illumina reads. The read coverage of Illumina and all ONT reads along the Fm loci is consistent, showing a sharp increase in coverage at the Dup1 and Dup2 regions. The read coverage of haplotype-specific ONT reads (A-D) decreases at the Dup1 end and Dup2 start. However, the total read coverage remains comparable across the duplicated regions. This difference in coverage between haplotype-assigned reads and the total coverage is reflected by an increase in unassigned reads (E) at the Dup1 end and Dup2 start. The observed drop in coverage for haplotype-specific reads in these regions is attributed to a lack of Haplotype Defining Positions (HDPs).

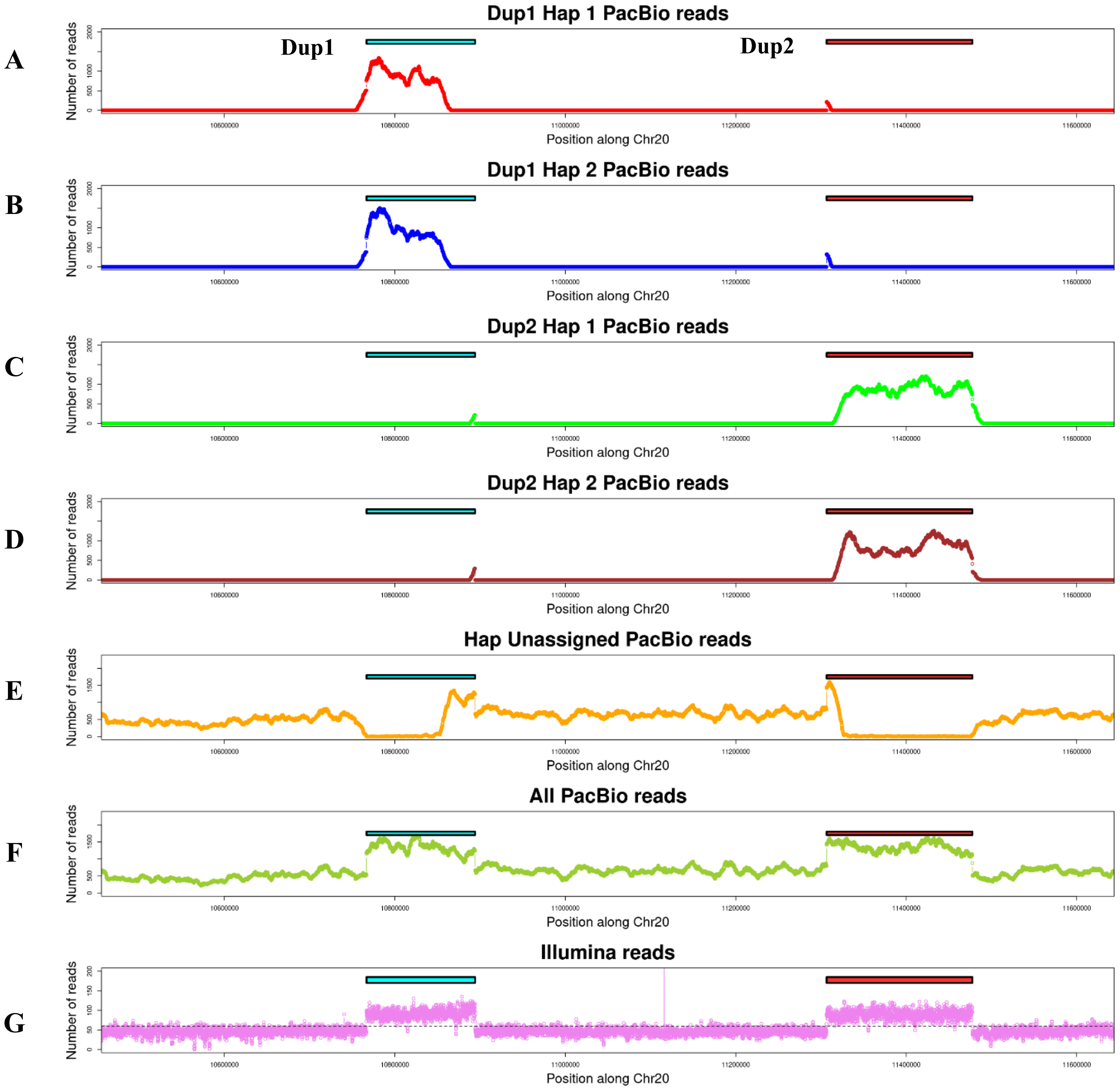


**Supplementary Figure. S34: Read coverage of CAU Silkie PacBio and Illumina reads mapped to the GRCg6a genome assembly at *Fm* locus using minimap2 and bedtools coverage: (A)** Dup1 Hap1-specific PacBio reads. **(B)** Dup1 Hap2-specific PacBio reads. **(C)** Dup2 Hap1-specific PacBio reads. **(D)** Dup2 Hap2-specific PacBio reads. **(E)** Reads not assigned to any haplotype. **(F)** All PacBio reads. **(G)** Illumina reads. The read coverage of Illumina and all PacBio reads along the Fm loci is consistent, showing a sharp increase in coverage at the Dup1 and Dup2 regions. The read coverage of haplotype-specific PacBio reads (A-D) decreases at the Dup1 end and Dup2 start. However, the total read coverage remains comparable across the duplicated regions. This difference in coverage between haplotype-assigned reads and the total coverage is reflected by an increase in unassigned reads (E) at the Dup1 end and Dup2 start. The observed drop in coverage for haplotype-specific reads in these regions is attributed to a lack of Haplotype Defining Positions (HDPs).

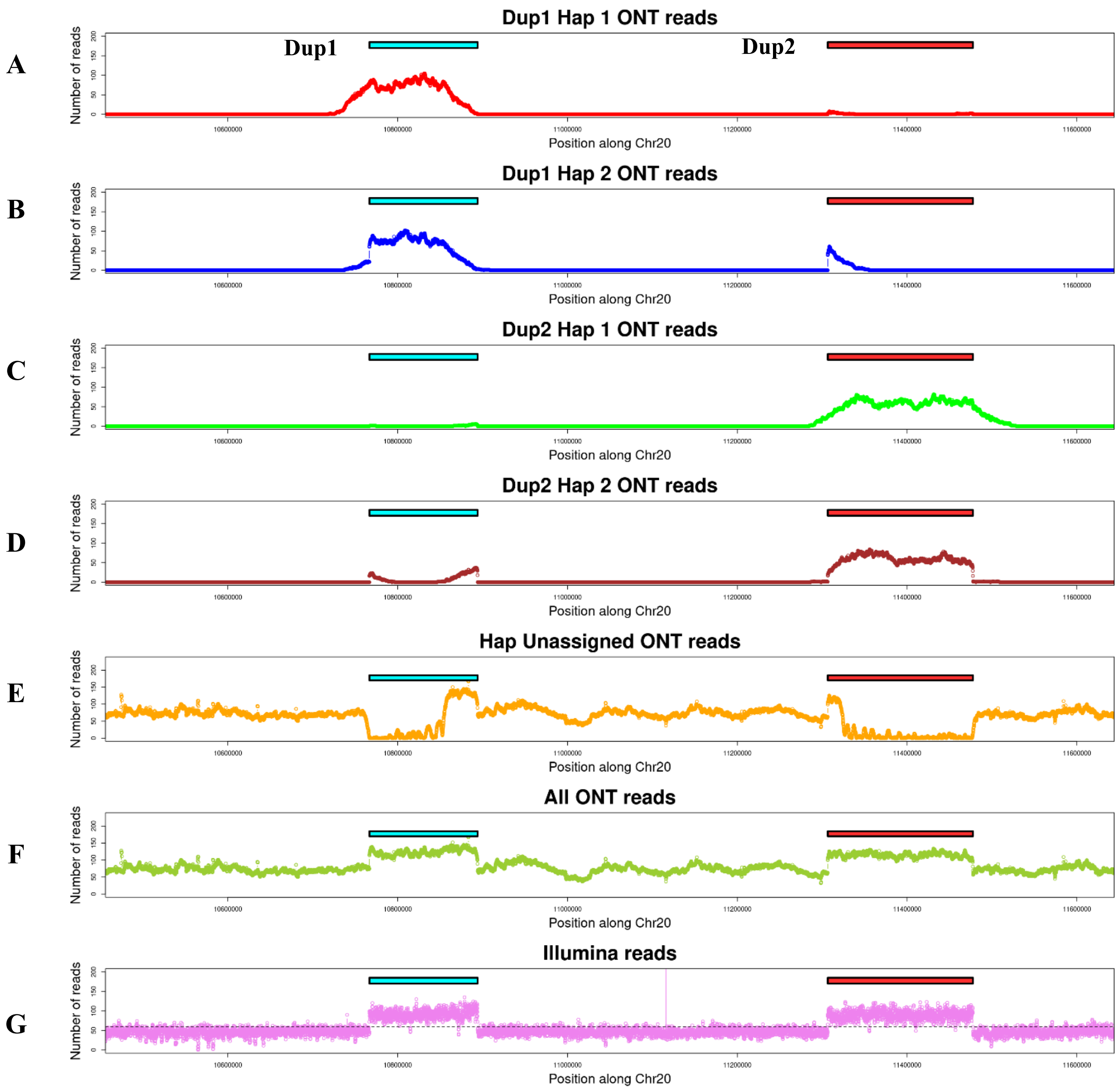


**Supplementary Figure. S35: Read coverage of ONT and Illumina reads mapped to the GRCg6a genome assembly at *Fm* locus using bwa and bedtools coverage: (A)** Dup1 Hap1-specific ONT reads. **(B)** Dup1 Hap2-specific ONT reads. **(C)** Dup2 Hap1-specific ONT reads. **(D)** Dup2 Hap2-specific ONT reads. **(E)** Reads not assigned to any haplotype. **(F)** All ONT reads. **(G)** Illumina reads. The read coverage of Illumina and all ONT reads along the Fm loci is consistent, showing a sharp increase in coverage at the Dup1 and Dup2 regions. The read coverage of haplotype-specific ONT reads (A-D) decreases at the Dup1 end and Dup2 start. However, the total read coverage remains comparable across the duplicated regions. This difference in coverage between haplotype-assigned reads and the total coverage is reflected by an increase in unassigned reads (E) at the Dup1 end and Dup2 start. The observed drop in coverage for haplotype-specific reads in these regions is attributed to a lack of Haplotype Defining Positions (HDPs).

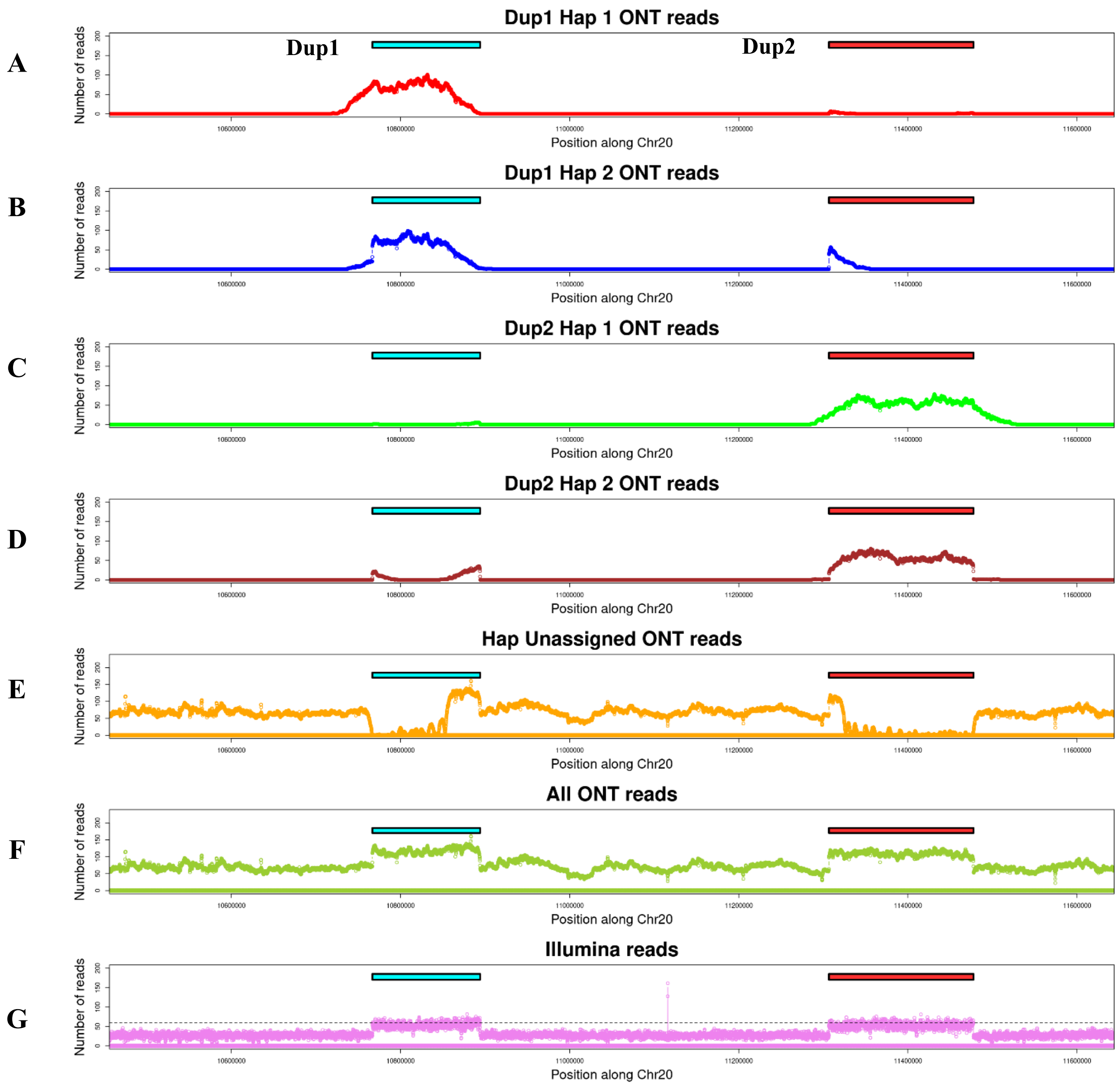


**Supplementary Figure. S36: Read coverage of ONT and Illumina reads mapped to the GRCg6a genome assembly at *Fm* locus using bwa and MosDepth coverage: (A)** Dup1 Hap1-specific ONT reads. **(B)** Dup1 Hap2-specific ONT reads. **(C)** Dup2 Hap1-specific ONT reads. **(D)** Dup2 Hap2-specific ONT reads. **(E)** Reads not assigned to any haplotype. **(F)** All ONT reads. **(G)** Illumina reads. The read coverage of Illumina and all ONT reads along the Fm loci is consistent, showing a sharp increase in coverage at the Dup1 and Dup2 regions. The read coverage of haplotype-specific ONT reads (A-D) decreases at the Dup1 end and Dup2 start. However, the total read coverage remains comparable across the duplicated regions. This difference in coverage between haplotype-assigned reads and the total coverage is reflected by an increase in unassigned reads (E) at the Dup1 end and Dup2 start. The observed drop in coverage for haplotype-specific reads in these regions is attributed to a lack of Haplotype Defining Positions (HDPs).

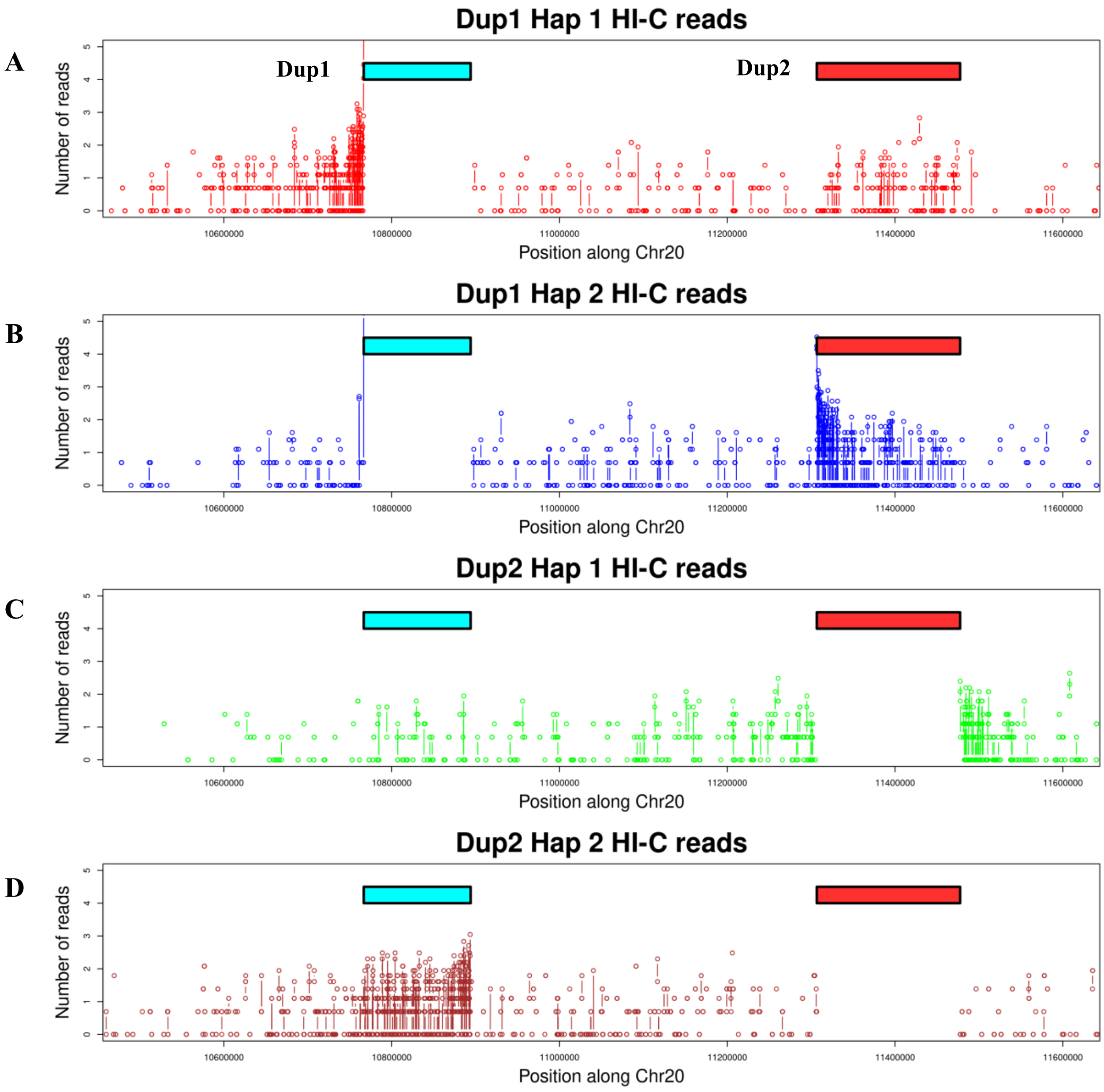


**Supplementary Figure. S36a: Read coverage of CAU Silkie HI-C reads mapped to the GRCg6a genome assembly at *Fm* locus using bwa and bedtools coverage: (A)** Dup1 Hap1-specific HI-C reads. **(B)** Dup1 Hap2-specific HI-C reads. **(C)** Dup2 Hap1-specific HI-C reads. **(D)** Dup2 Hap2-specific HI-C reads. The read coverage for Dup1 and Dup2 region is not shown for a better visualization of mapping of mate pair reads. The mate pair reads of Dup1 Hap1 showing an increase in coverage at the flank1 regions while the mate pair reads of Dup1 Hap2 show an increase in coverage Dup2 region. The mate pair reads of Dup2 Hap1 showing an increase in coverage at the Flank2 and Int regions while the mate pair reads of Dup2 Hap2 show an increase in coverage Dup1 region.

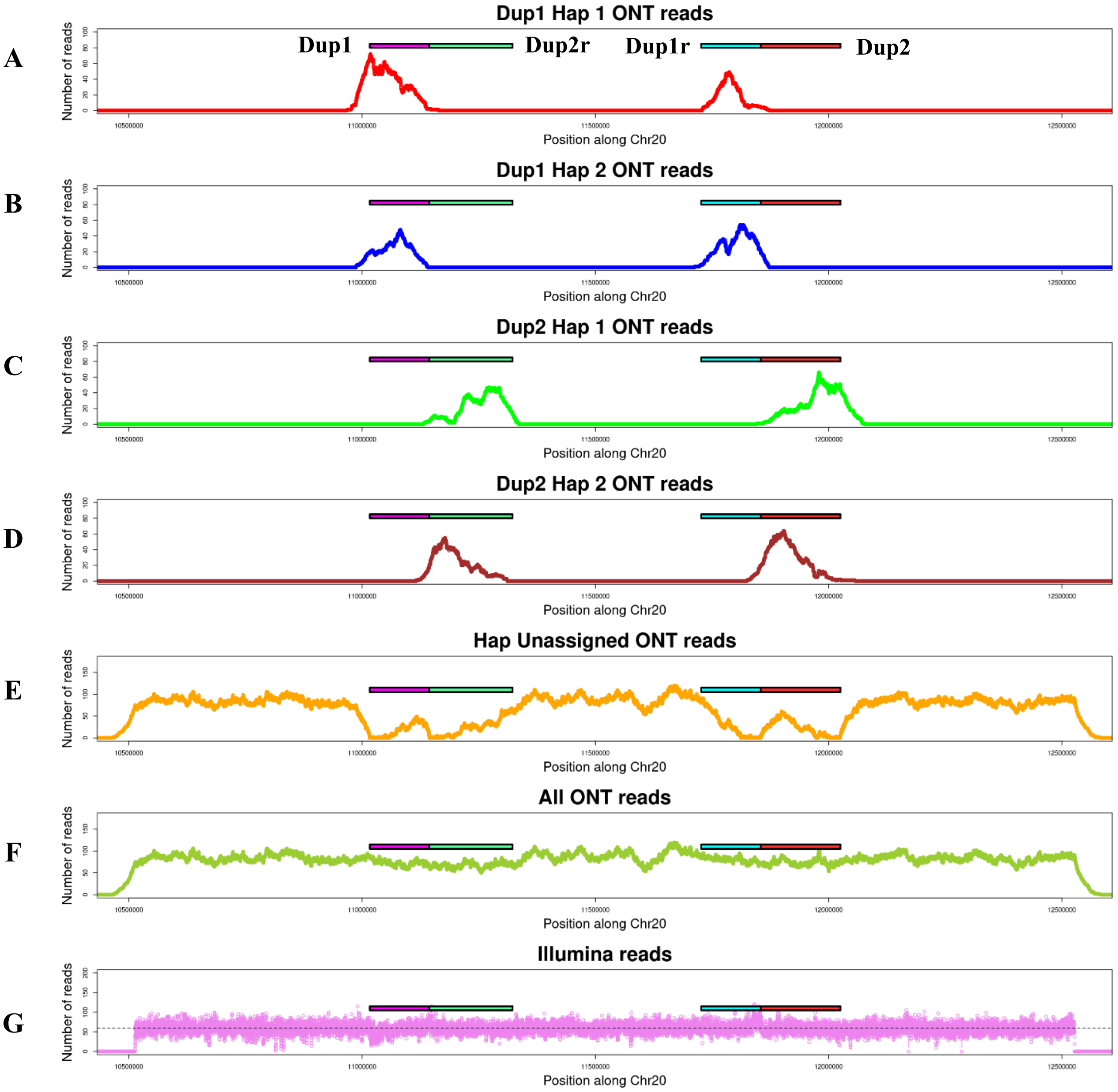


**Supplementary Figure. S37: Read coverage analysis of ONT and Illumina reads mapped to the CAU Silkie genome assembly at the *Fm* locus, performed using minimap2 and bedtools coverage:** The figure includes the following panels: **(A)** Coverage of Dup1 Hap1-specific ONT reads, **(B)** Coverage of Dup1 Hap2-specific ONT reads, **(C)** Coverage of Dup2 Hap1-specific ONT reads, **(D)** Coverage of Dup2 Hap2-specific ONT reads, **(E)** Coverage of reads not assigned to any haplotype, **(F)** Coverage of all ONT reads combined, and **(G)** Coverage of Illumina reads. The coverage profiles for Illumina and all ONT reads (F and G) are consistent across the Fm locus. However, the haplotype-specific ONT reads (A-D) exhibit reduced coverage at the end of Dup1, the start of Dup2r, as well as the end of Dup1r and start of Dup2. This difference in coverage suggests a mosaic assembly in CAU silkie in this region which lacks Haplotype Defining Positions (HDPs).

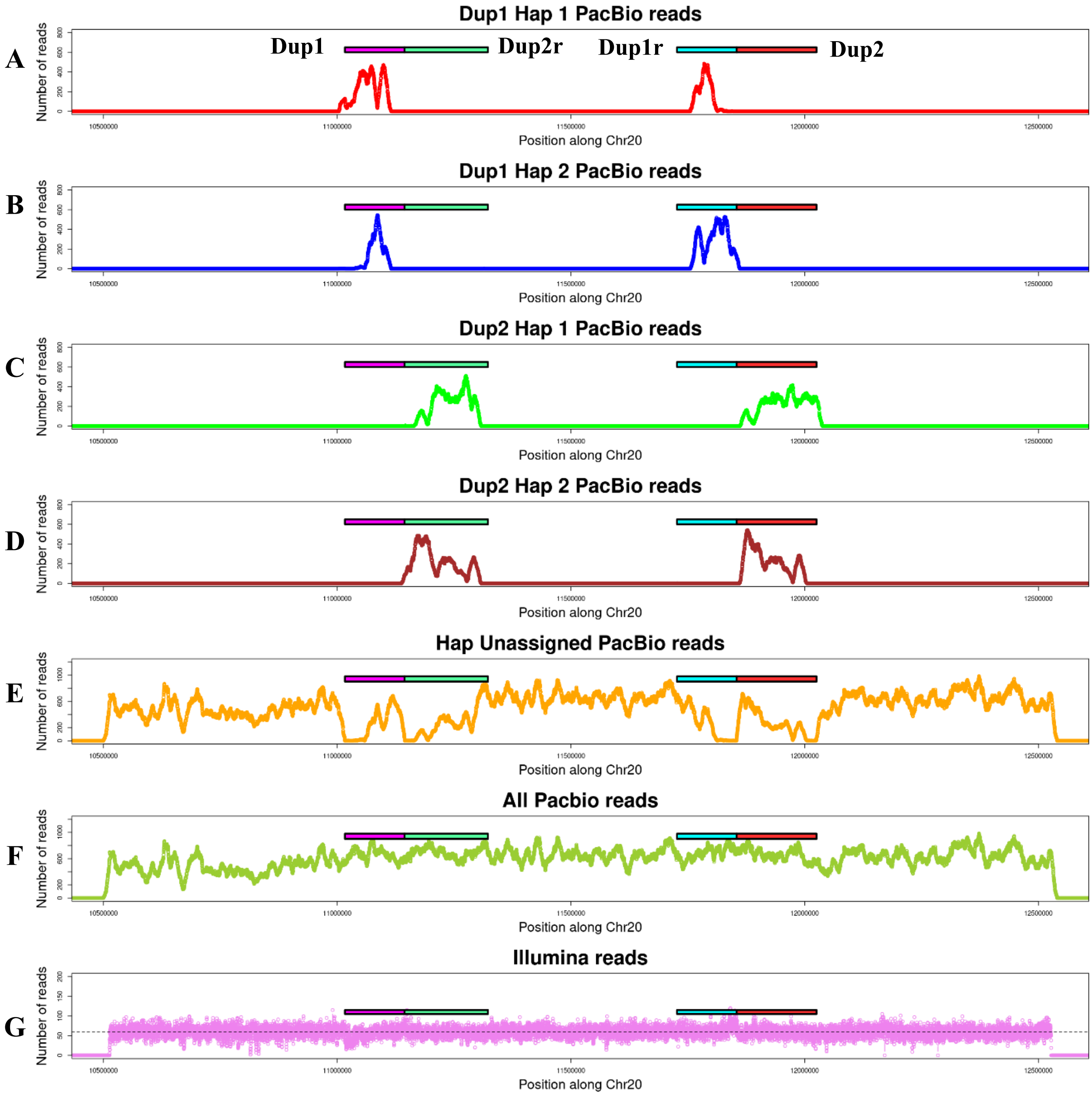


**Supplementary Figure. S38: Read coverage analysis of PacBio and Illumina reads mapped to the CAU Silkie genome assembly at the *Fm* locus, performed using minimap2 and bedtools coverage:** The figure includes the following panels: **(A)** Coverage of Dup1 Hap1-specific PacBio reads, **(B)** Coverage of Dup1 Hap2-specific PacBio reads, **(C)** Coverage of Dup2 Hap1-specific PacBio reads, **(D)** Coverage of Dup2 Hap2-specific PacBio reads, **(E)** Coverage of reads not assigned to any haplotype, **(F)** Coverage of all PacBio reads combined, and **(G)** Coverage of Illumina reads. The coverage profiles for Illumina and all PacBio reads (F and G) are consistent across the Fm locus. However, the haplotype-specific PacBio reads (A-D) exhibit reduced coverage at the end of Dup1, the start of Dup2r, as well as the end of Dup1r and the start of Dup2. This difference in coverage suggests a mosaic assembly in CAU silkie in this region which lacks Haplotype Defining Positions (HDPs).

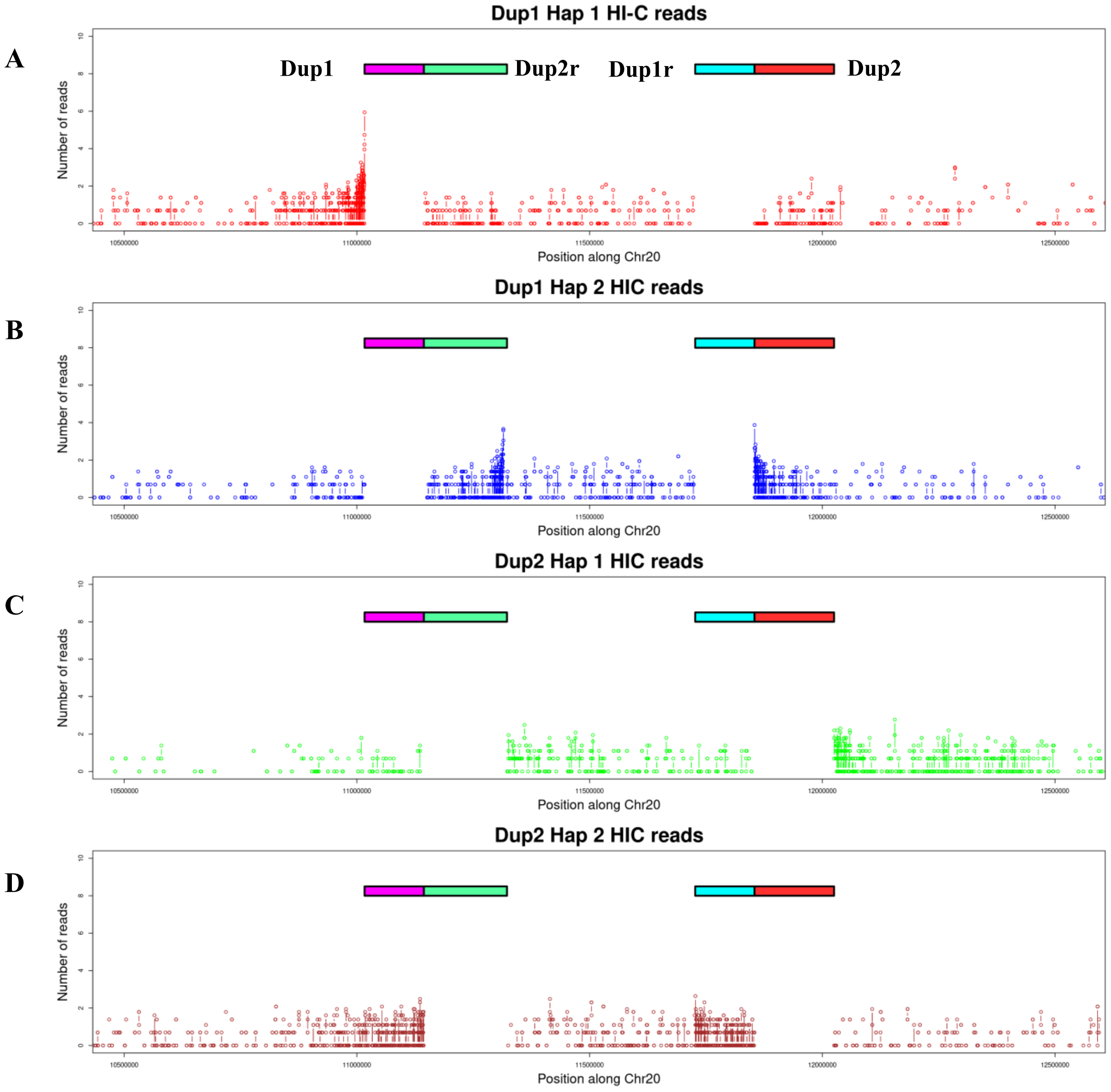


**Supplementary Figure. S38a: Read coverage of CAU Silkie HI-C reads mapped to the CAU Silkie genome assembly at *Fm* locus using bwa and bedtools coverage: (A)** Dup1 Hap1-specific HI-C reads. **(B)** Dup1 Hap2-specific HI-C reads. **(C)** Dup2 Hap1-specific HI-C reads. **(D)** Dup2 Hap2-specific HI-C reads. The read coverage for Dup1 and Dup2 region is not shown for a better visualization of mapping of mate pair reads. The mate pair reads of Dup1 Hap1 showing an increase in coverage at the flank1 regions while the mate pair reads of Dup1 Hap2 show an increase in coverage at start region of Dup2r and Dup2. The mate pair reads of Dup2 Hap1 showing an increase in coverage at the Flank2 and Int regions compared to Dup1r. The mate pair reads of Dup2 Hap2 show an increase in coverage at Dup1 and Dup1r region at same level.

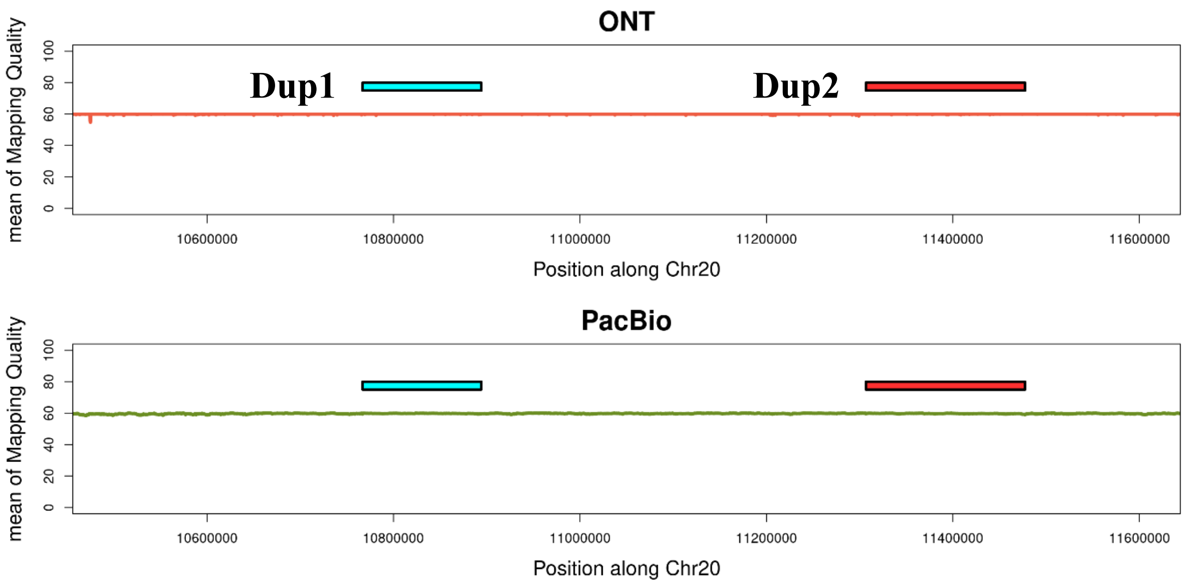


**Supplementary Figure. S39: Mean mapping quality at the *Fm* locus for ONT and PacBio long-reads mapped to the GRCg6a genome assembly:** This figure shows the mean mapping quality scores for ONT and PacBio long-reads aligned to the *Fm* locus in the GRCg6a genome assembly. The mapping quality scores reflect the reliability of read alignments and provide an overview of the reliability and accuracy of read mappings in this region of the GRCg6a genome assembly.

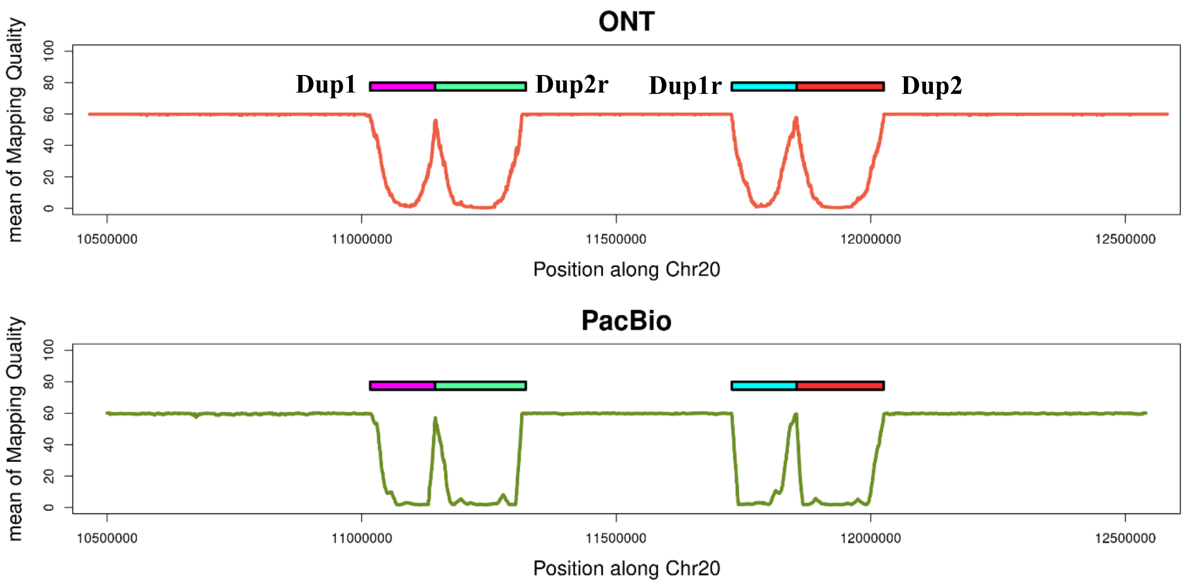


**Supplementary Figure. S40: Mean mapping quality at the Fm locus for ONT and PacBio long-reads mapped to the CAU Silkie genome assembly:** This figure shows the mean mapping quality scores for ONT and PacBio long-reads aligned to the Fm locus in the CAU Silkie genome assembly. The mapping quality is notably lower in duplicated regions, attributed to the multimapping of the same reads to both haplotypes.

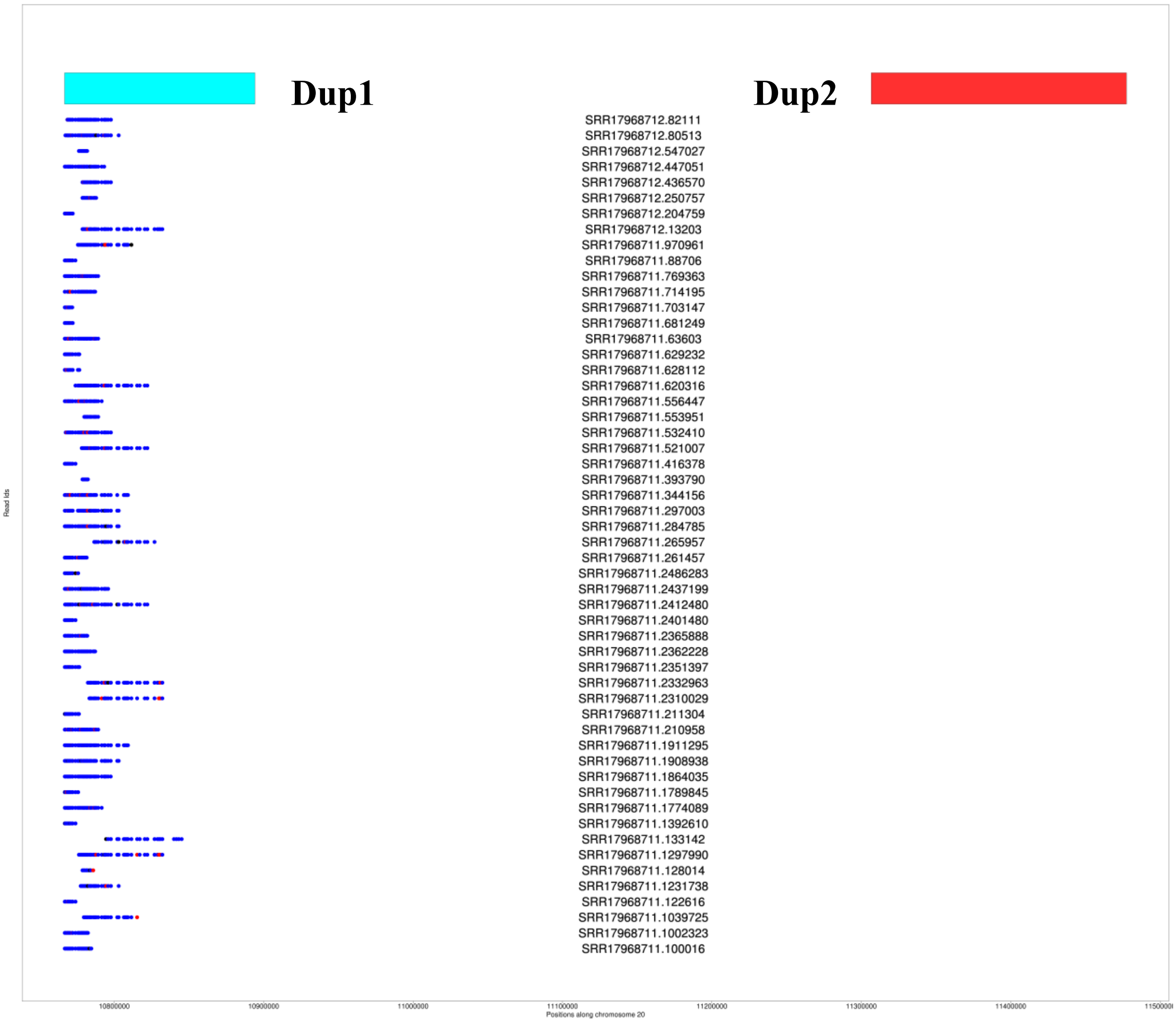


**Supplementary Figure. S41: Individual Read-Based Support for Each HDP in Dup1 Hap1 CAU Silkie ONT:** In this figure, blue indicates that the base in a particular read matches the haplotype-specific base, while red denotes a mismatch. The tiling path of reads comprehensively covers the approximate Dup1 region along haplotype-consistent paths.

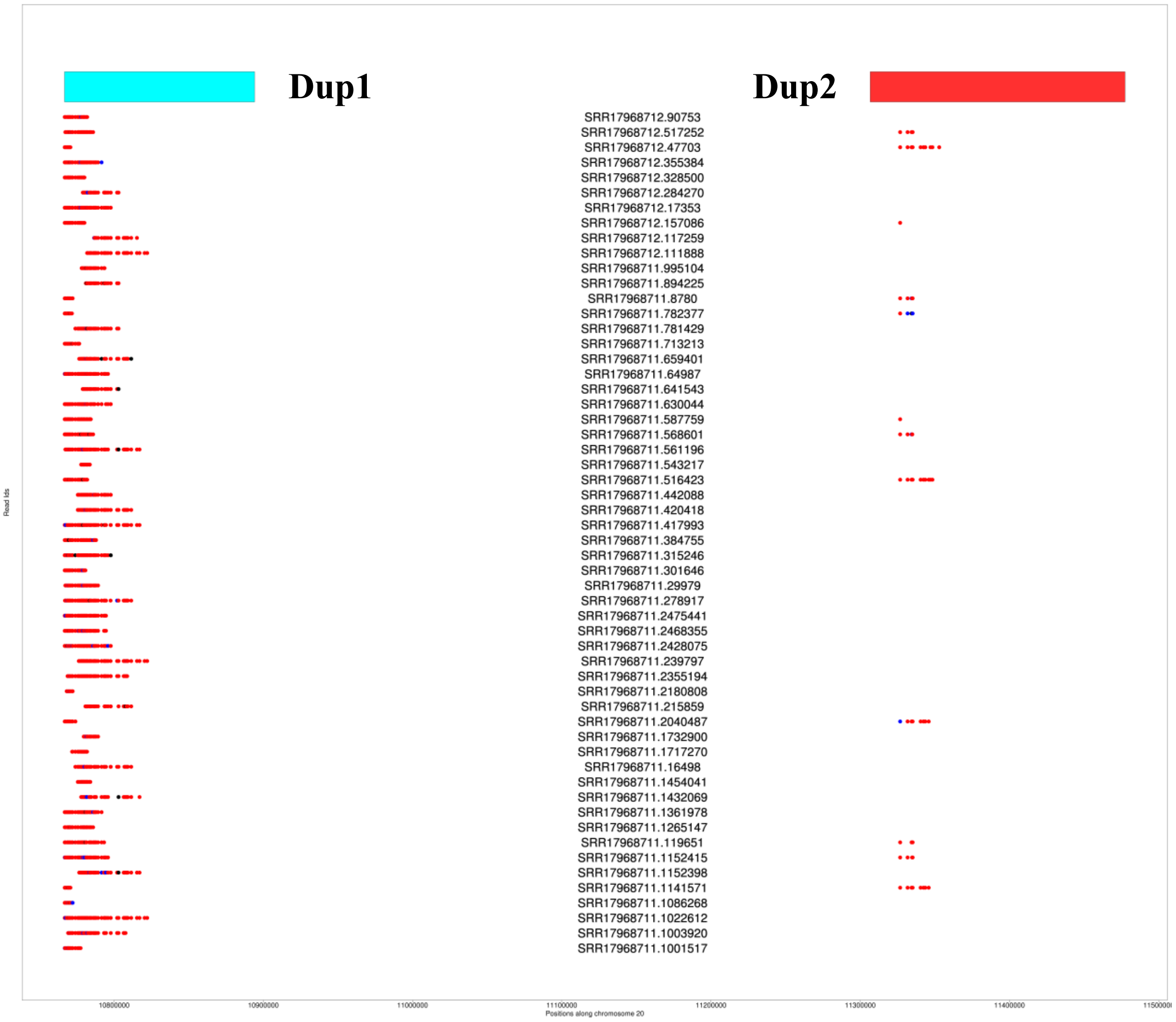


**Supplementary Figure. S42: Individual Read-Based Support for Each HDP in Dup1 Hap2 CAU Silkie ONT:** In this figure, red indicates that the base in a particular read matches the haplotype-specific base, while blue denotes a mismatch. The tiling path of reads comprehensively covers the approximate Dup1 region along haplotype-consistent paths.

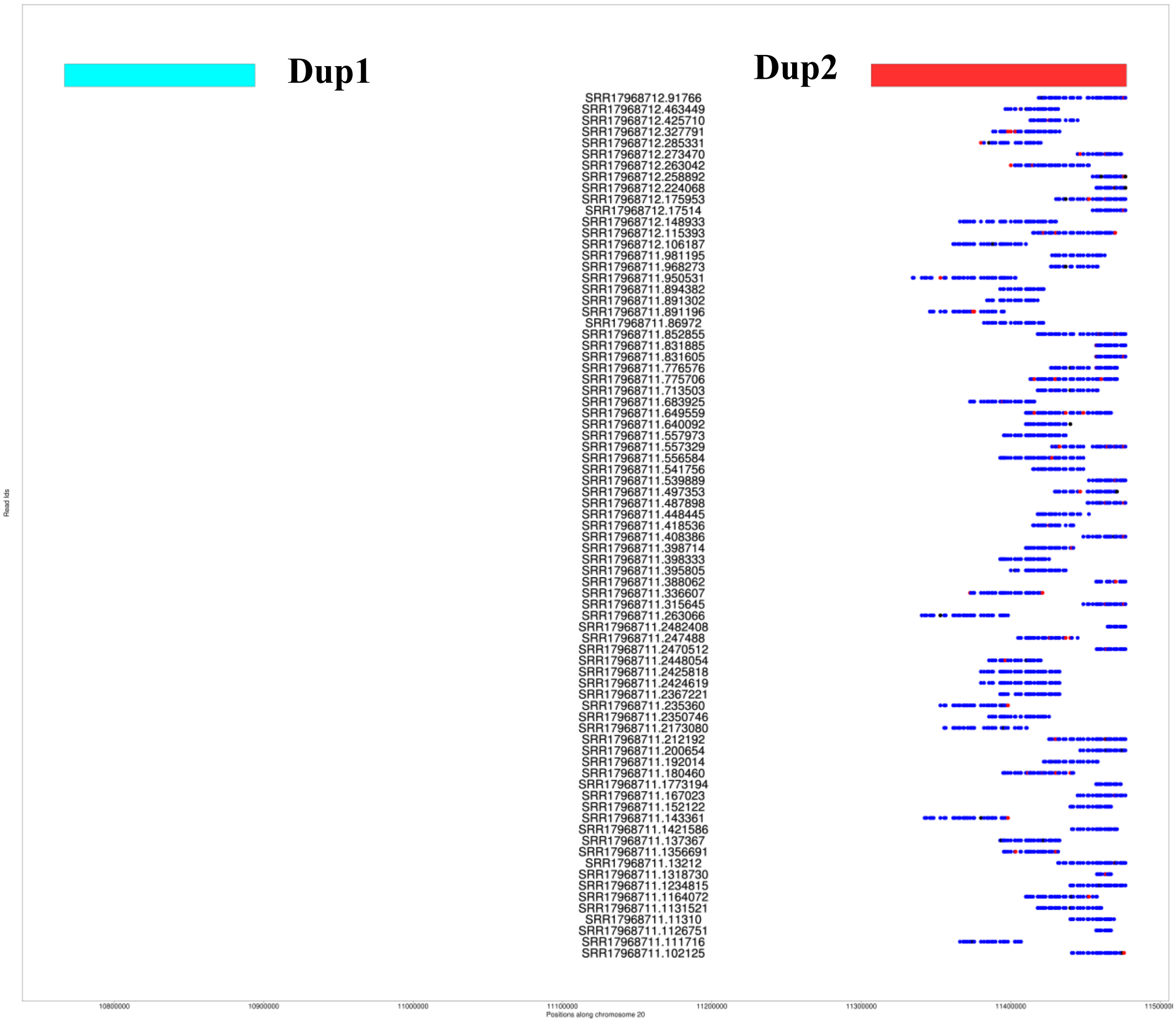


**Supplementary Figure. S43: Individual Read-Based Support for Each HDP in Dup2 Hap1 CAU Silkie ONT:** In this figure, blue indicates that the base in a particular read matches the haplotype-specific base, while red denotes a mismatch. The tiling path of reads comprehensively covers the approximate Dup2 region along haplotype-consistent paths.

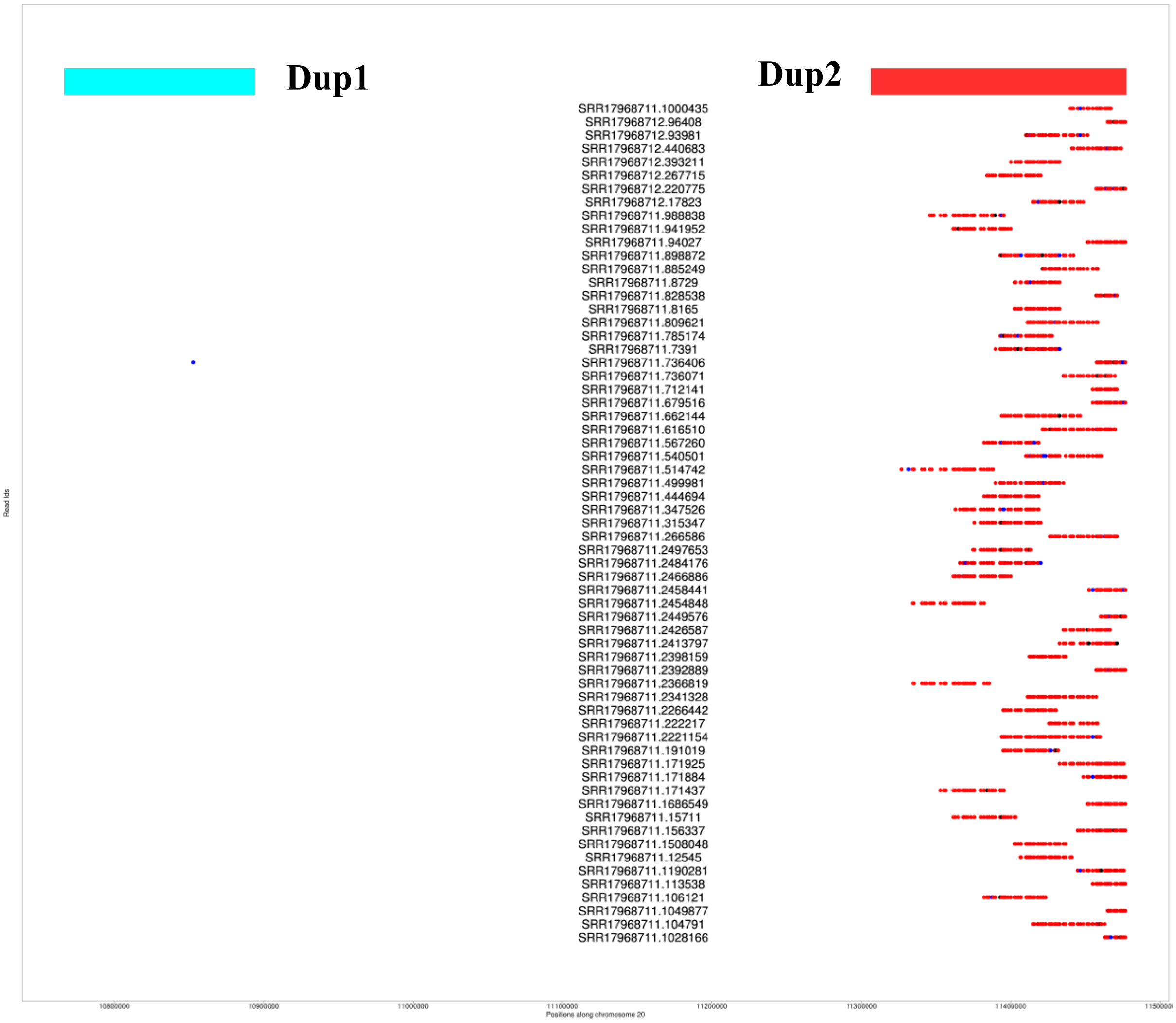


**Supplementary Figure. S44: Individual Read-Based Support for Each HDP in Dup2 Hap2 CAU Silkie ONT:** In this figure, red indicates that the base in a particular read matches the haplotype-specific base, while blue denotes a mismatch. The tiling path of reads comprehensively covers the approximate Dup2 region along haplotype-consistent paths.

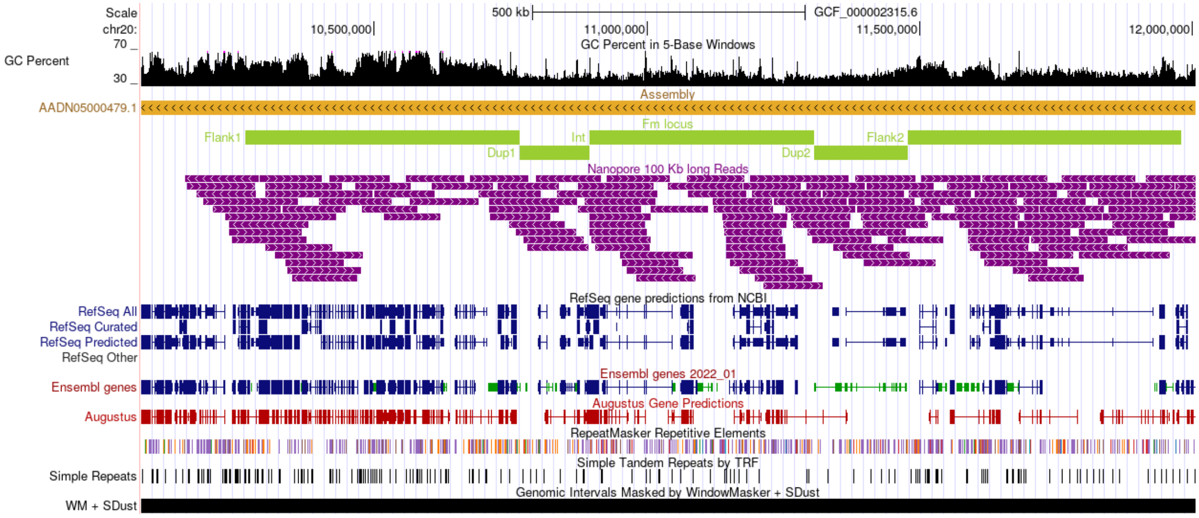


**Supplementary Figure. S45: Verification of chicken genome assembly (GRCg6a) using Oxford Nanopore Technologies (ONT) long reads:** This figure presents verification of GRCg6a genome assembly using long reads from the huxu and rooster breeds, aligned with the Fm locus. The screenshot from the UCSC Genome Browser shows the alignment of reads longer than 100 kb to the chicken genome at the *Fm* locus. To enhance the clarity of the figure, the nanopore reads have been compressed vertically. The consistent tiling path of these long reads across the Fm locus indicates robust support for the high quality of the genome assembly.

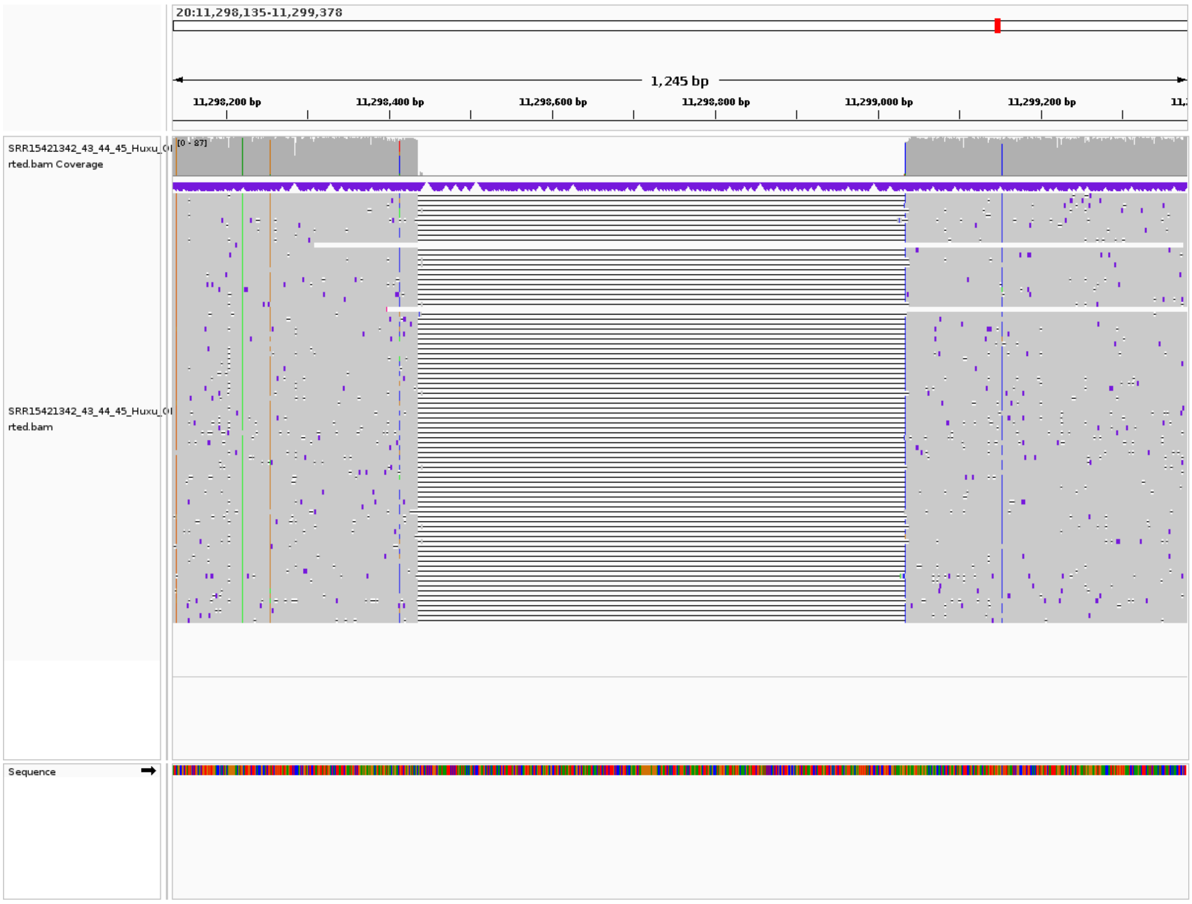


**Supplementary Figure. S46:** A Huxu breed-specific 598 bp deletion in the Int region. ONT reads from Huxu chicken was mapped to the GRCg6a genome assembly to identify this breed-specific deletion.

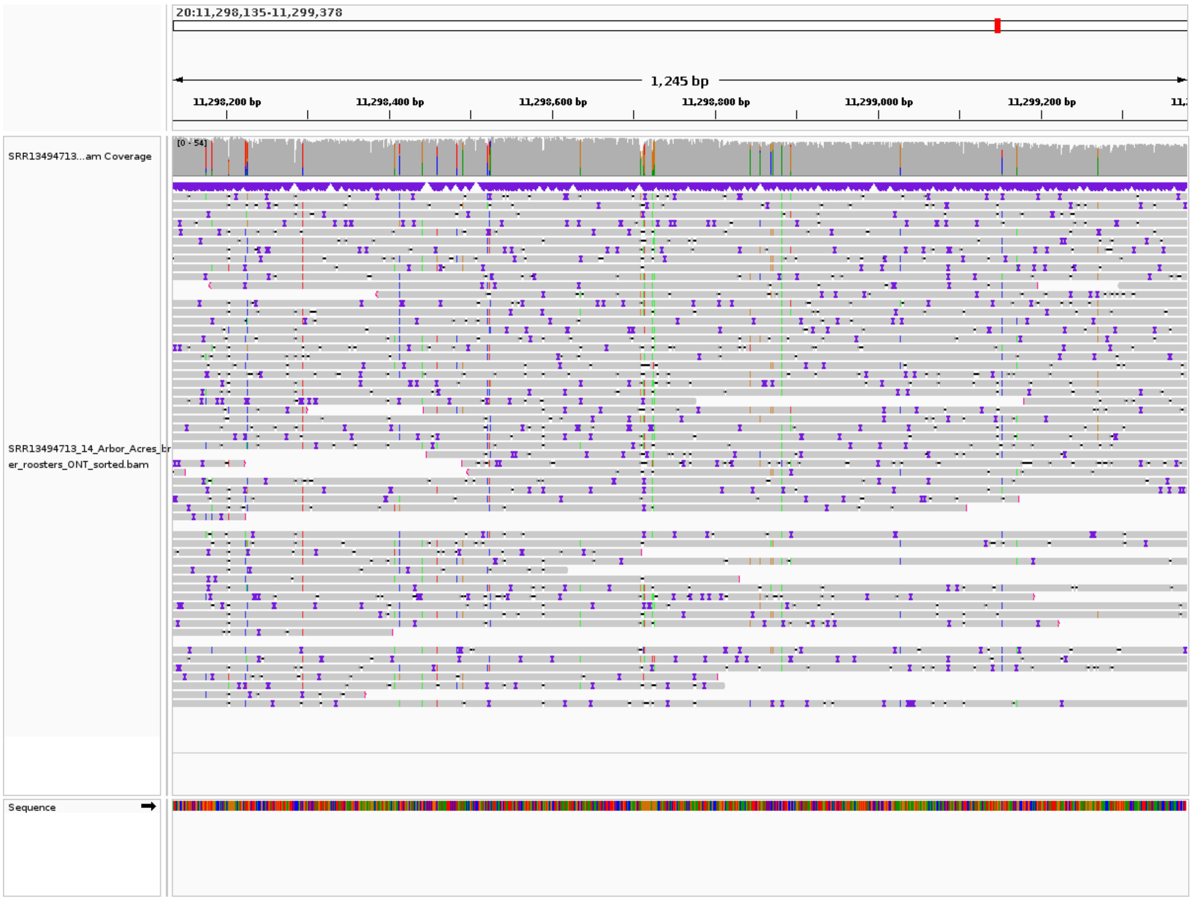


**Supplementary Figure. S47:** IGV screenshot of the Rooster chicken ONT long-reads mapped to the GRCg6a assembly. This image illustrates the absence of a ~598 bp deletion that is specific to the Huxu breed, thereby validating the deletion's exclusivity to Huxu. The screenshot provides visual evidence from IGV confirming that the deletion is not present in the GRCg6a genome assembly.

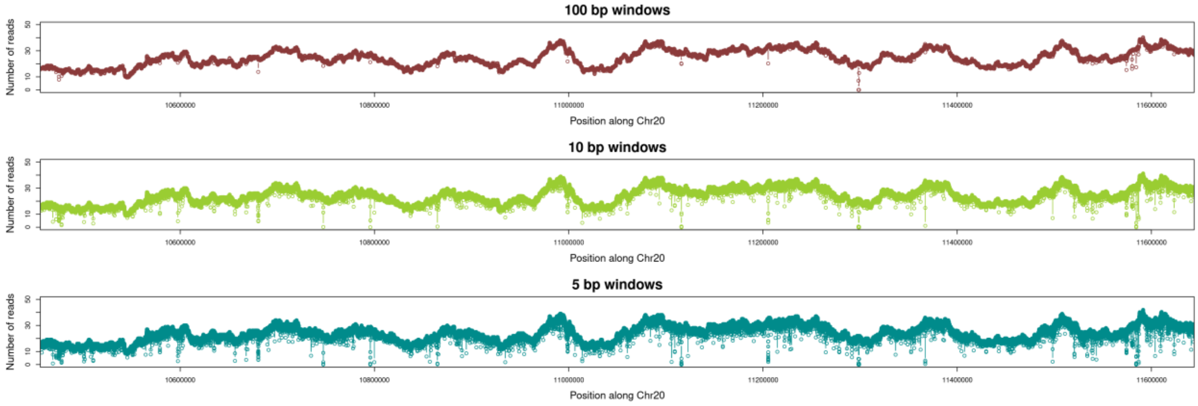


**Supplementary Figure. S48:** Read coverage in 100bp, 10bp, and 5bp windows along the entire length of the *Fm* locus in ONT long-reads of Huxu breed chicken mapped to the GRCg6a chicken genome assembly.

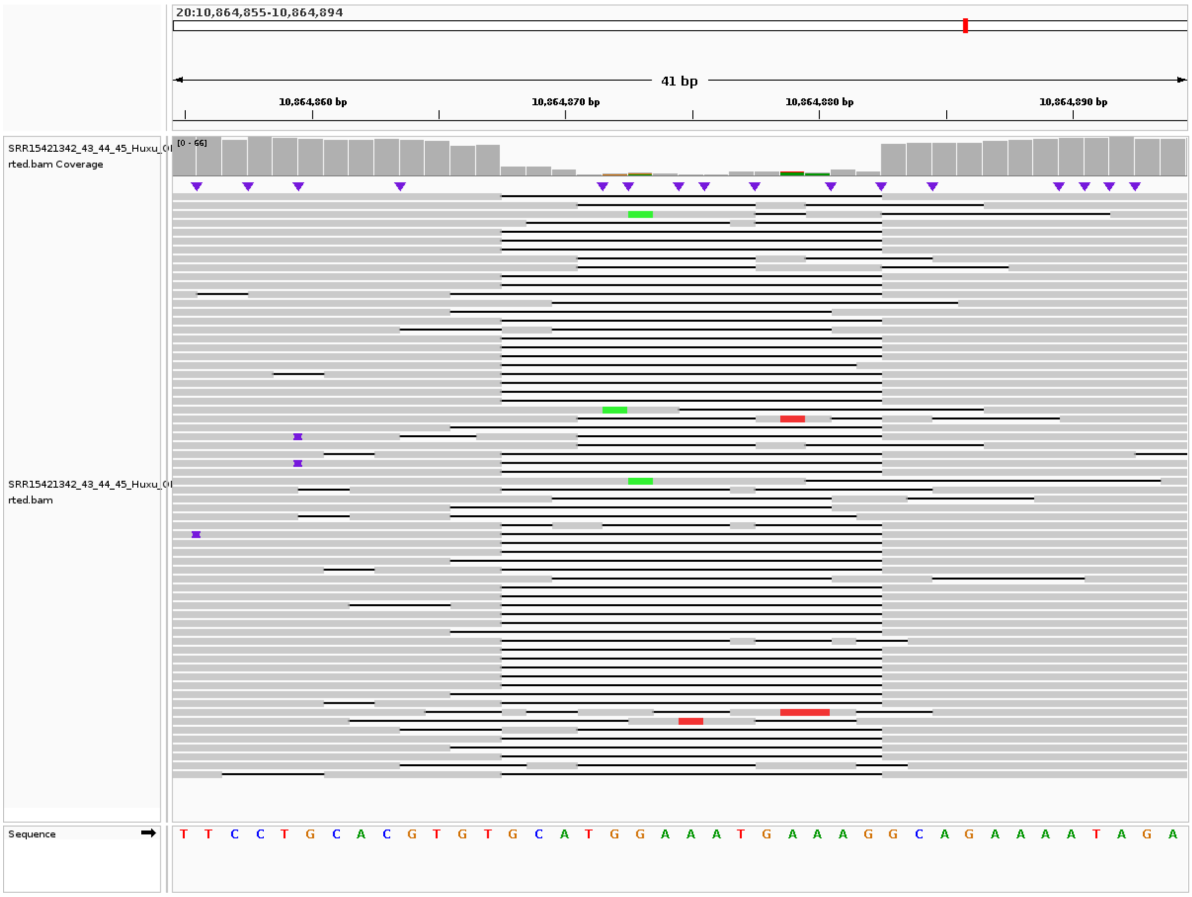


**Supplementary Figure. S49:** A Huxu breed-specific 15 bp deletion in the Dup1 region. ONT reads from Huxu chicken was mapped to the GRCg6a genome assembly to identify this breed-specific deletion.

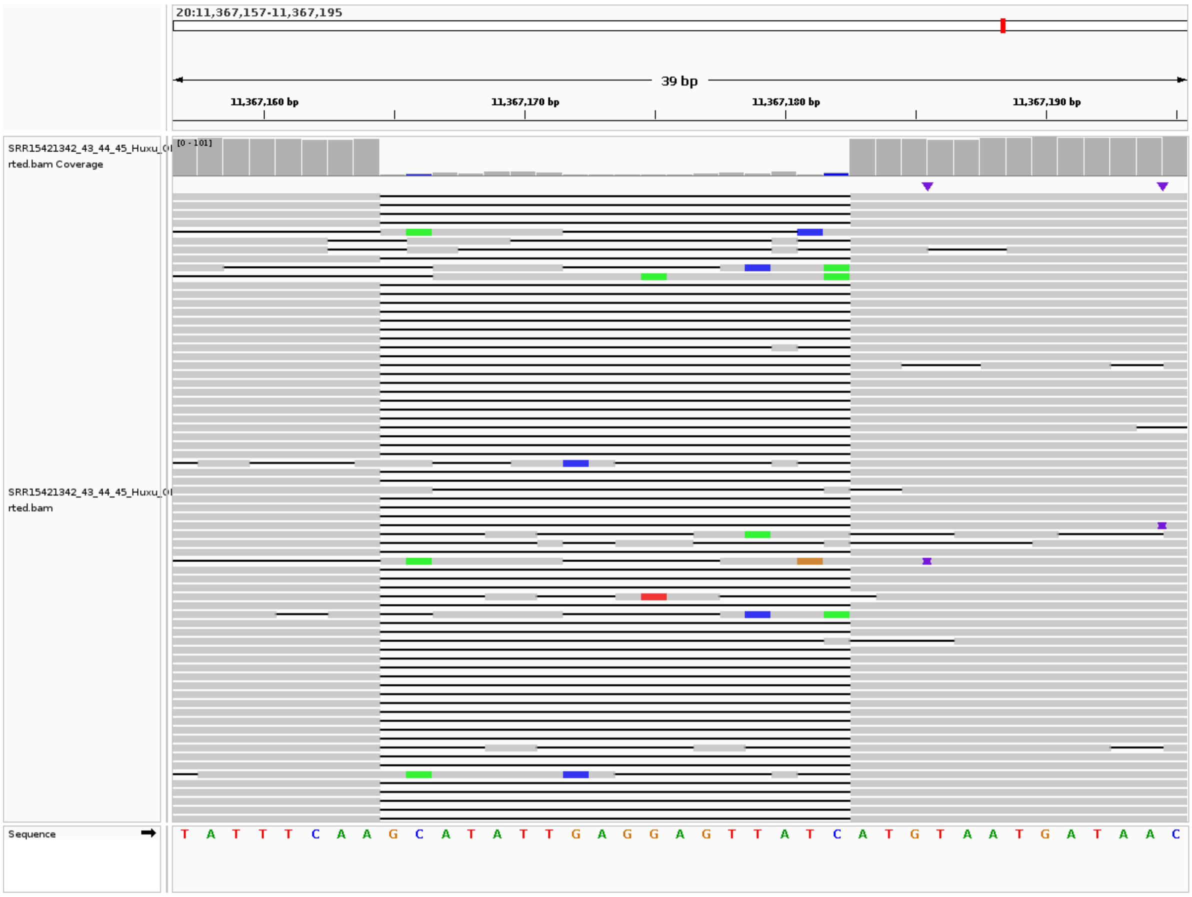


**Supplementary Figure. S50:** A Huxu breed-specific 18 bp deletion in the Dup2 region. ONT reads from Huxu chicken was mapped to the GRCg6a genome assembly to identify this breed-specific deletion.

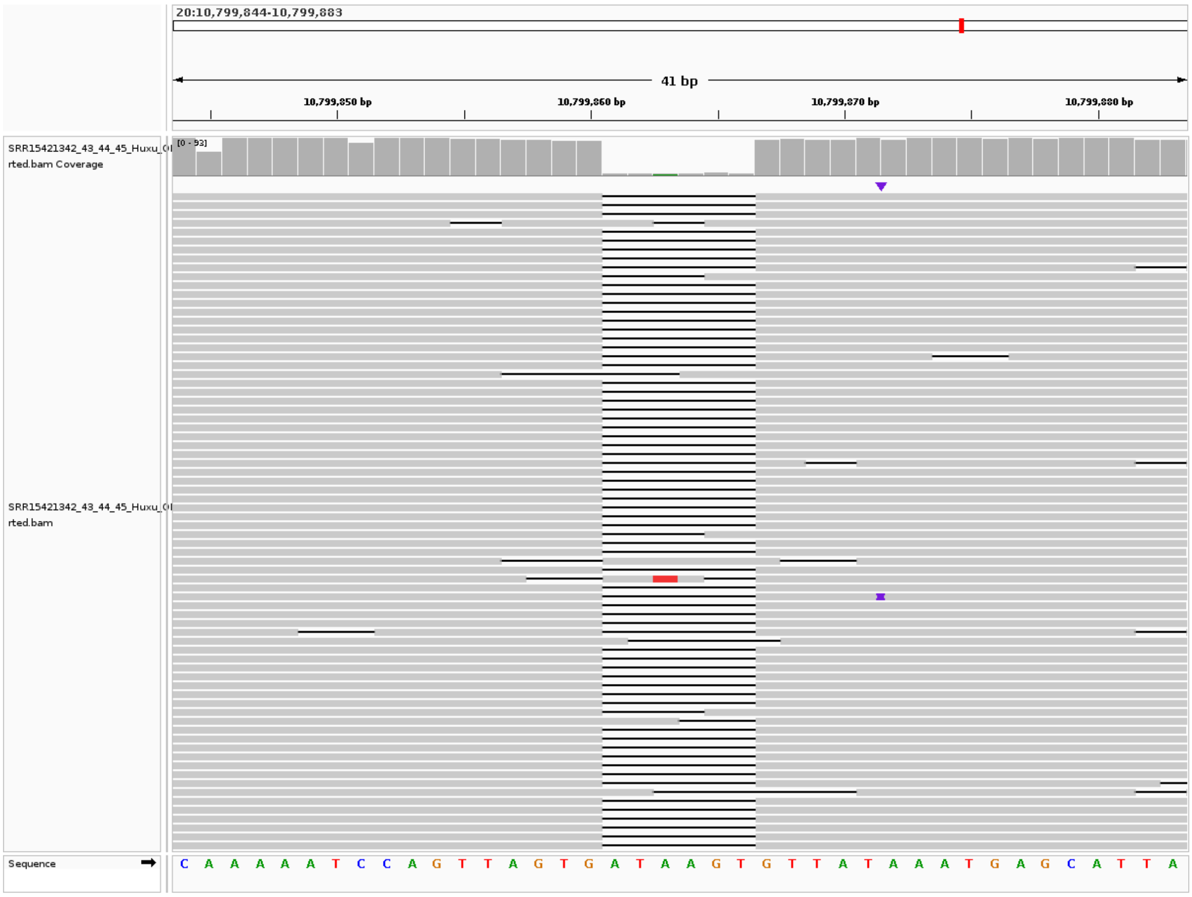


**Supplementary Figure. S51:** A Huxu breed-specific 6 bp deletion in the Dup1 region. ONT reads from Huxu chicken was mapped to the GRCg6a genome assembly to identify this breed-specific deletion.

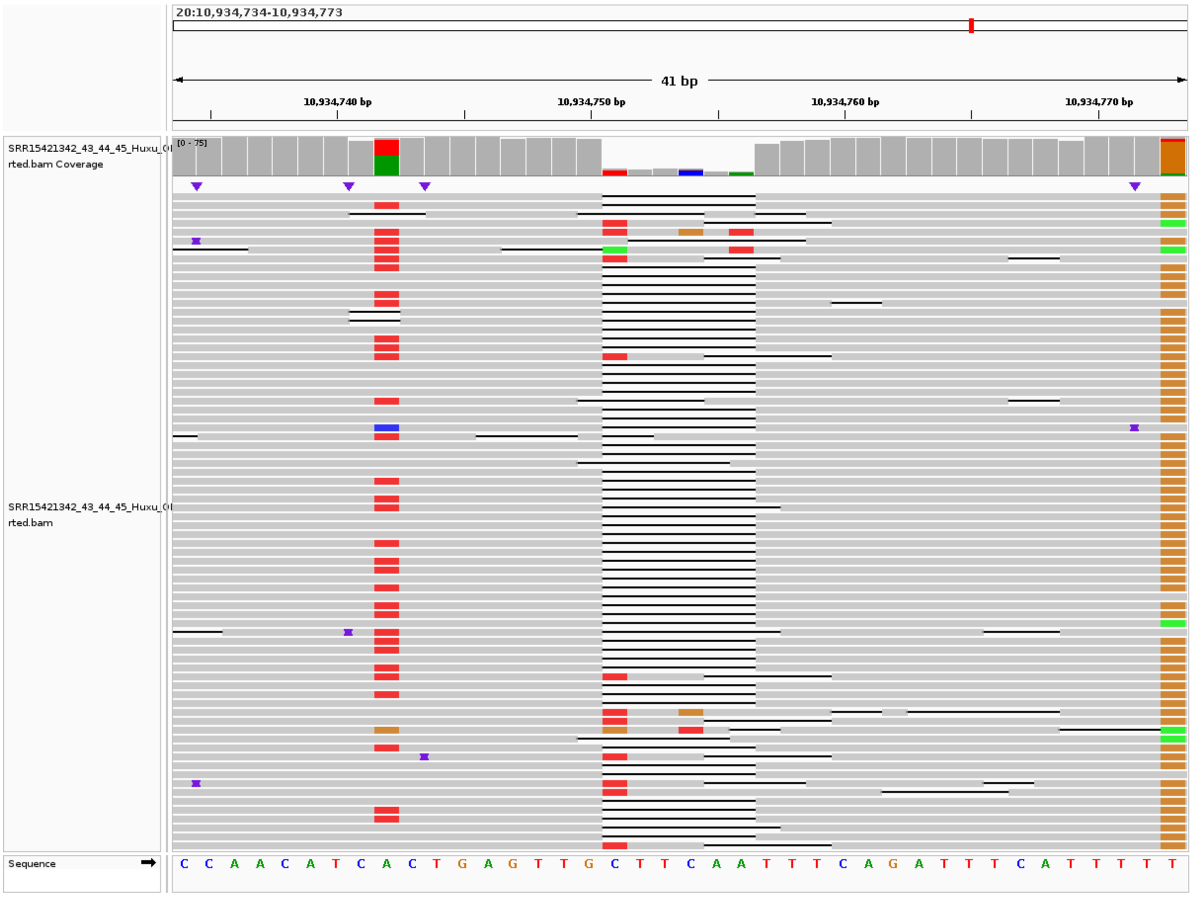


**Supplementary Figure. S52:** A Huxu breed-specific 6 bp deletion in the Int region. ONT reads from Huxu chicken was mapped to the GRCg6a genome assembly to identify this breed-specific deletion.

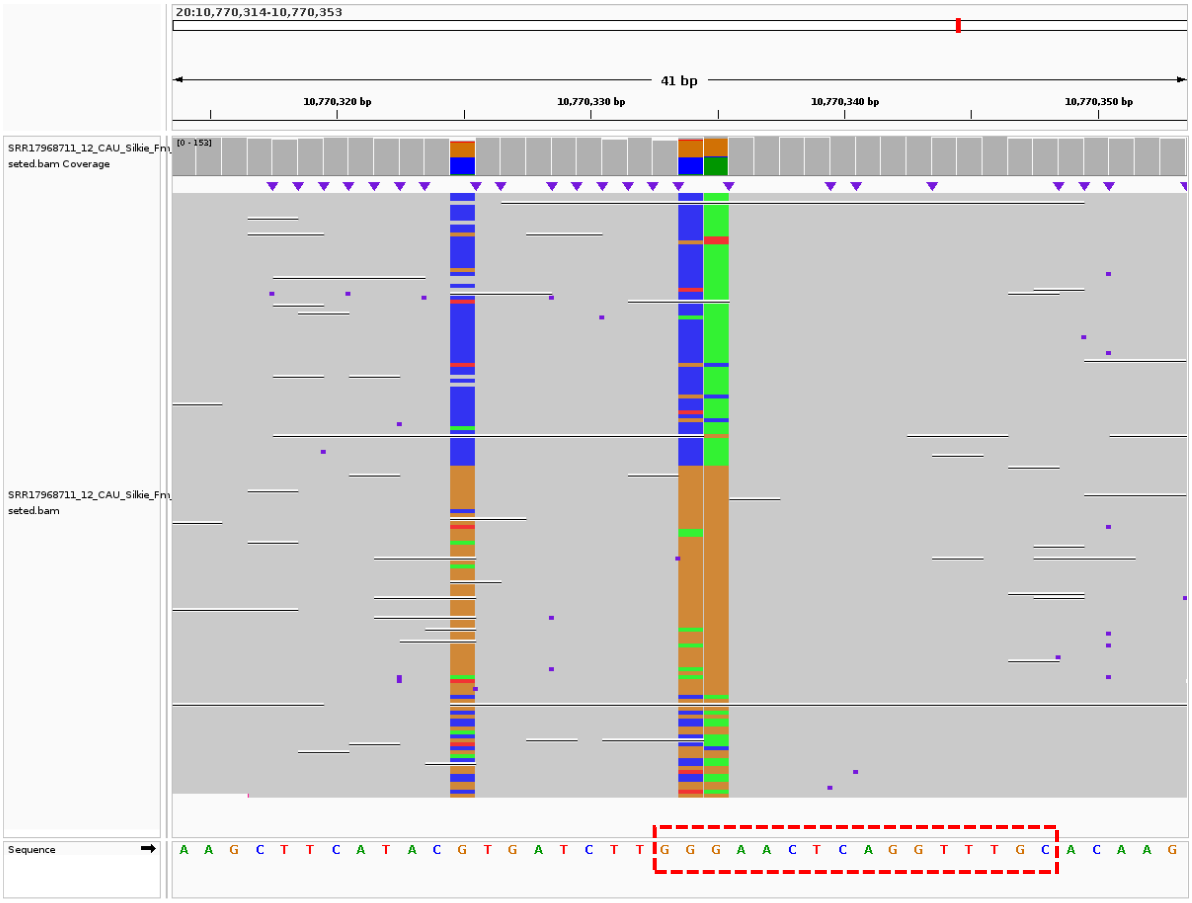


**Supplementary Figure S53:** IGV screenshot displaying Dup1 HDPs 10770334 and 10770335 present within a homopolymer run of G (3 bases) in a 10-mer window, highlighted by a red dotted rectangular box. The visualization shows comparable read support for both haplotypes, indicating that these HDPs are unlikely due to sequencing errors.

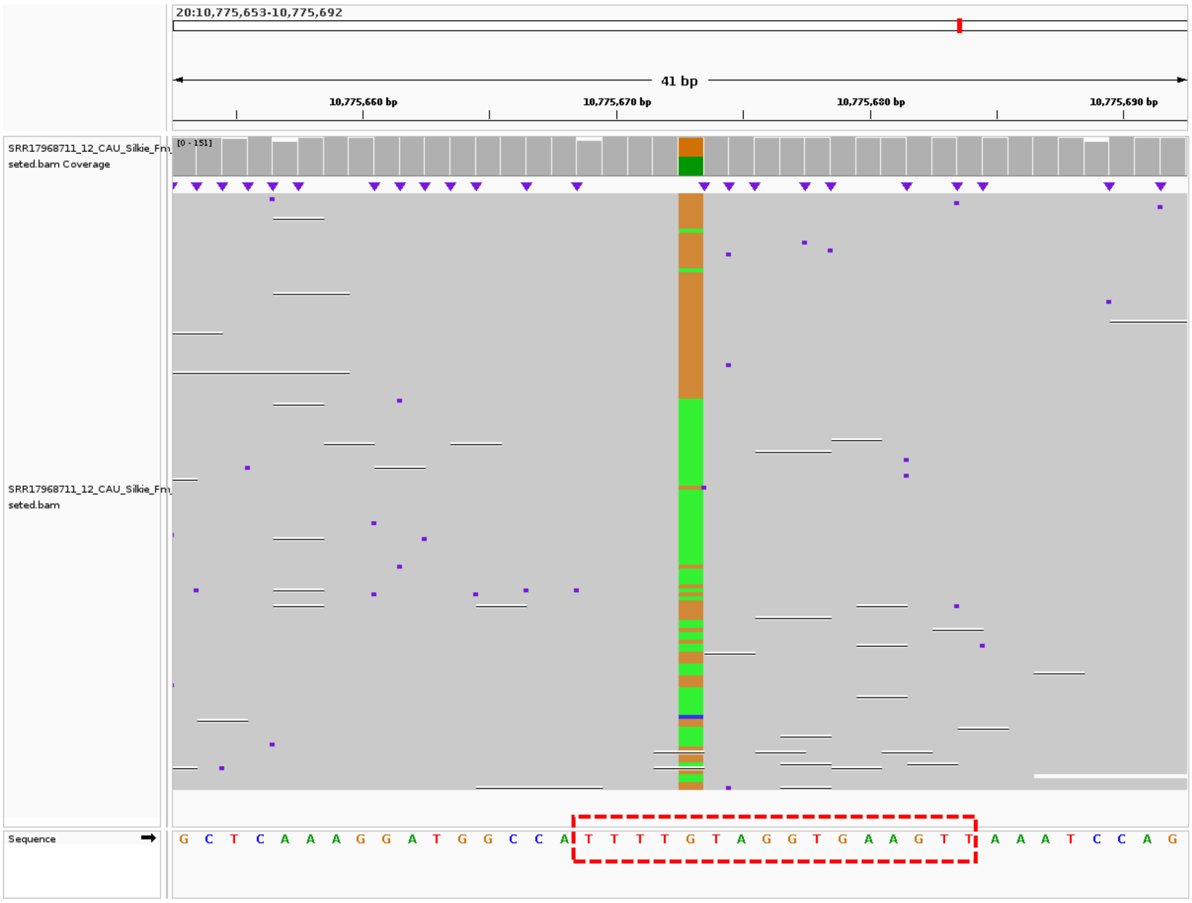


**Supplementary Figure S54:** IGV screenshot displaying Dup1 HDP 10775673 present within a homopolymer run of T (4 bases) in a 10-mer window, highlighted by a red dotted rectangular box. The visualization shows comparable read support for both haplotypes, indicating that the HDP is unlikely due to sequencing errors.

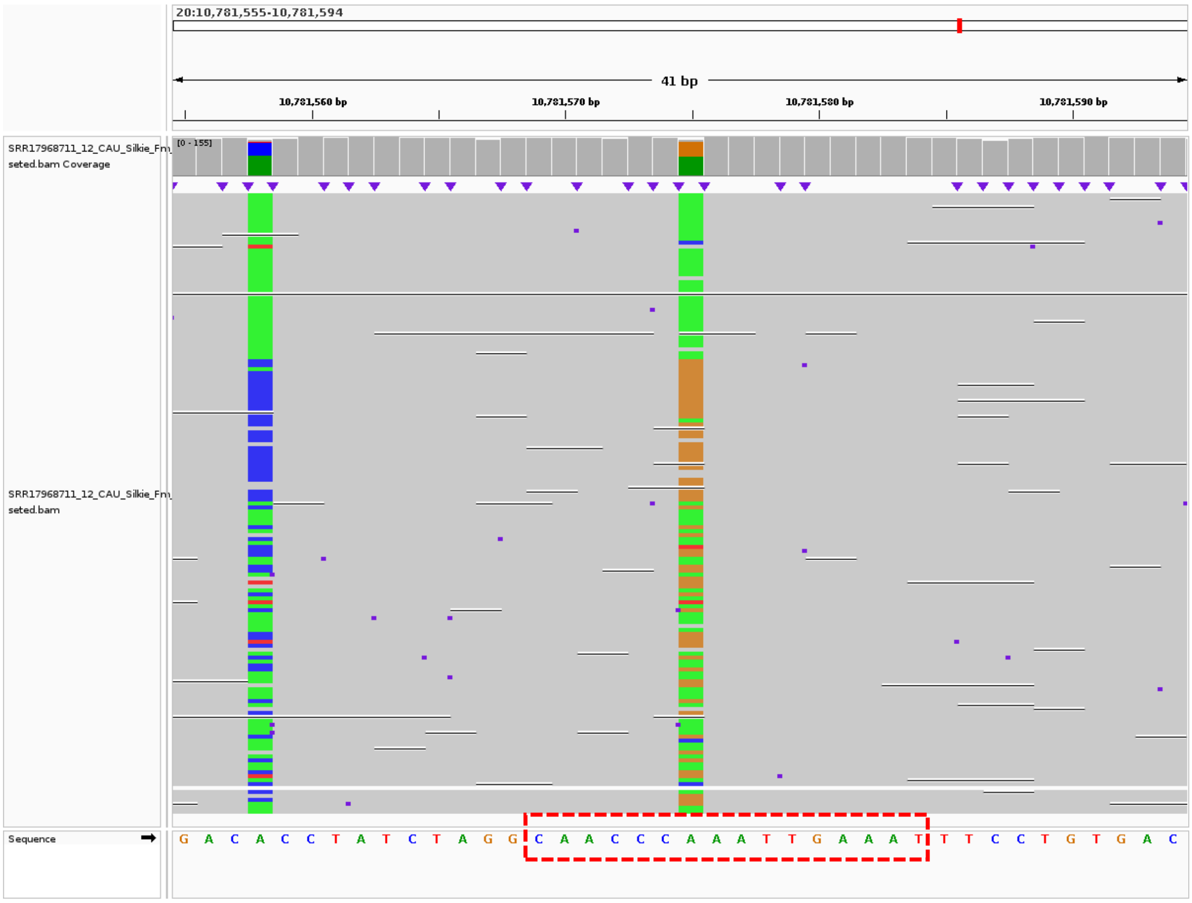


**Supplementary Figure S55:** IGV screenshot displaying Dup1 HDP 10781575 present within a homopolymer run of C (3 bases) in a 10-mer window, highlighted by a red dotted rectangular box. The visualization shows comparable read support for both haplotypes, indicating that the HDP is unlikely due to sequencing errors.

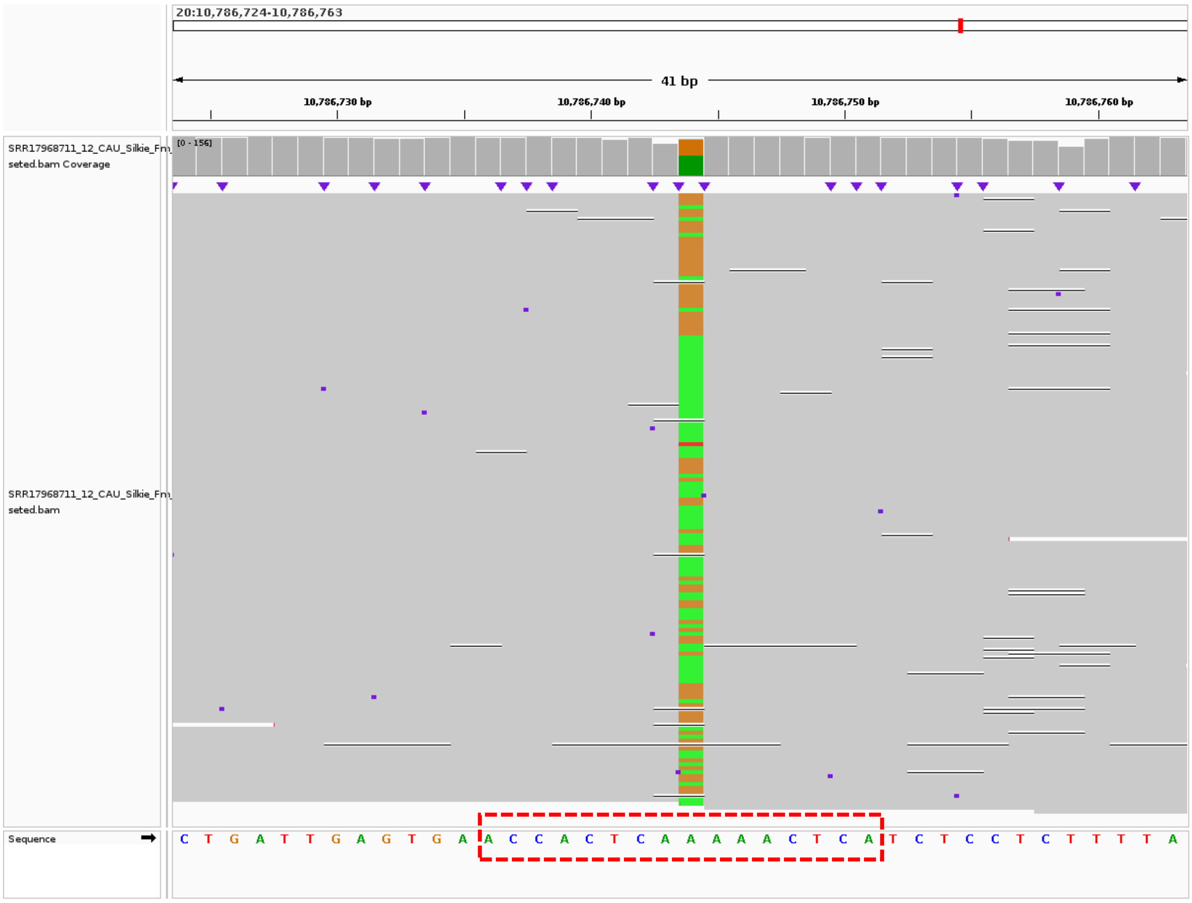


**Supplementary Figure S56:** IGV screenshot displaying Dup1 HDP 10786744 present within a homopolymer run of A (5 bases) in a 10-mer window, highlighted by a red dotted rectangular box. The visualization shows comparable read support for both haplotypes, indicating that the HDP is unlikely due to sequencing errors.

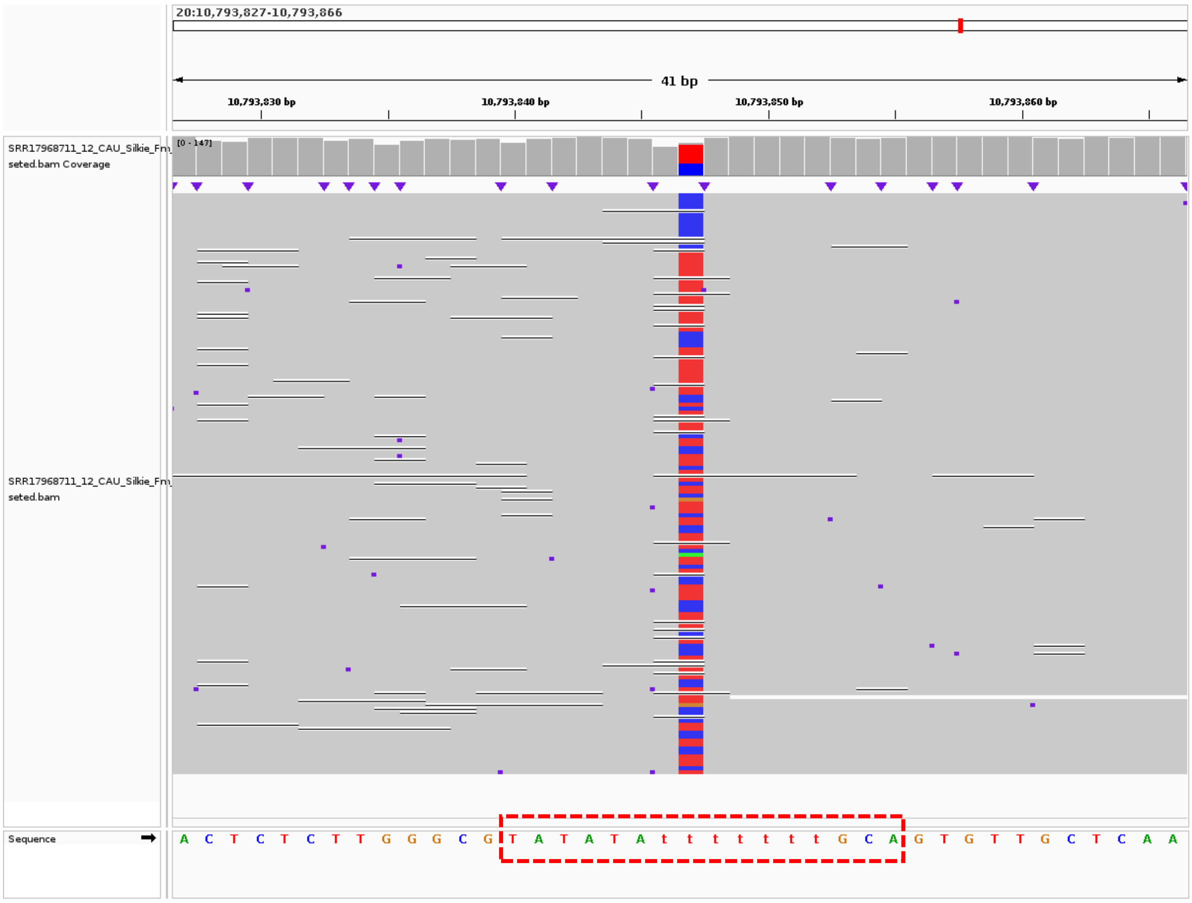


**Supplementary Figure S57:** IGV screenshot displaying Dup1 HDP 10793847 present within a homopolymer run of T (7 bases) in a 10-mer window, highlighted by a red dotted rectangular box. The visualization shows comparable read support for both haplotypes, indicating that the HDP is unlikely due to sequencing errors.

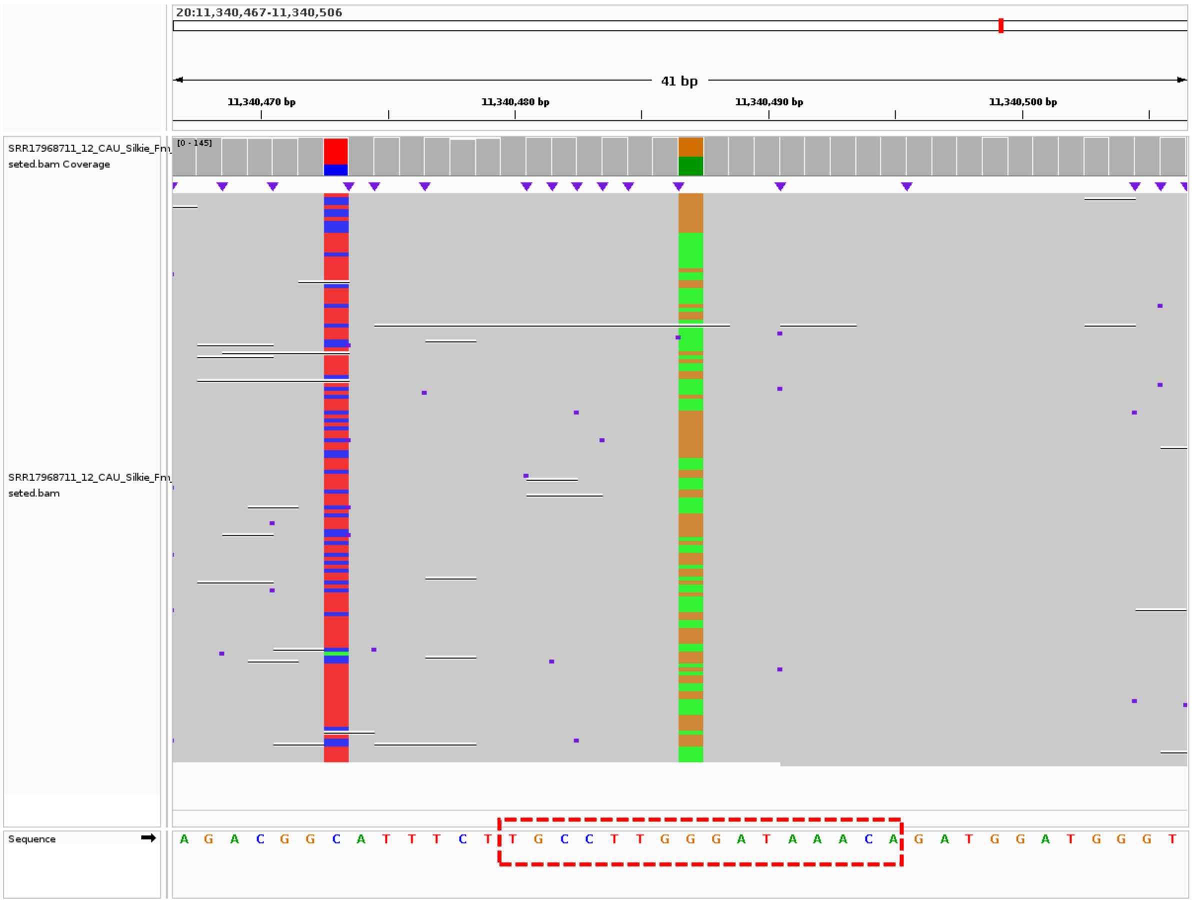


**Supplementary Figure S58:** IGV screenshot displaying Dup2 HDP 11340487 present within a homopolymer run of G (3 bases) in a 10-mer window, highlighted by a red dotted rectangular box. The visualization shows comparable read support for both haplotypes, indicating that the HDP is unlikely due to sequencing errors.

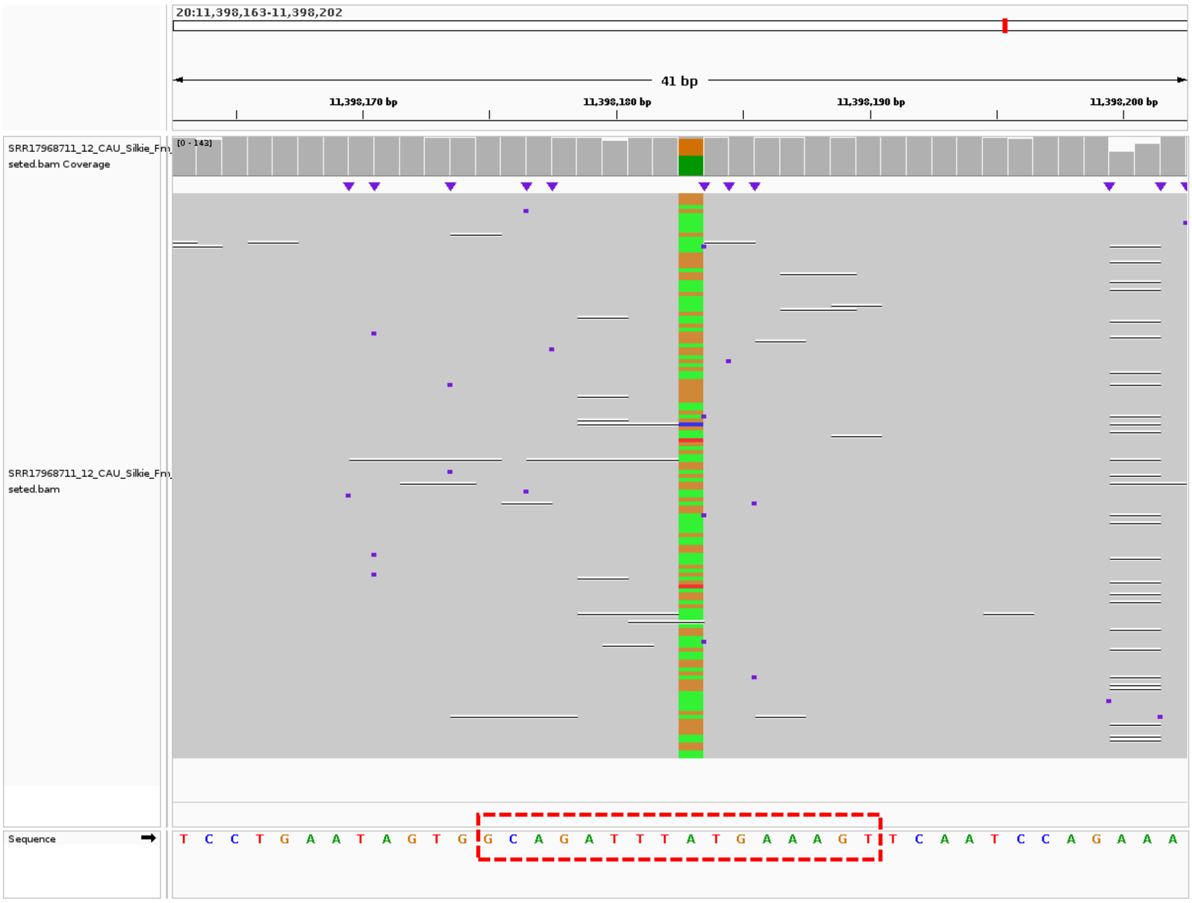


**Supplementary Figure S59:** IGV screenshot displaying Dup2 HDP 11398183 present within a homopolymer run of T (3 bases) in a 10-mer window, highlighted by a red dotted rectangular box. The visualization shows comparable read support for both haplotypes, indicating that the HDP is unlikely due to sequencing errors.

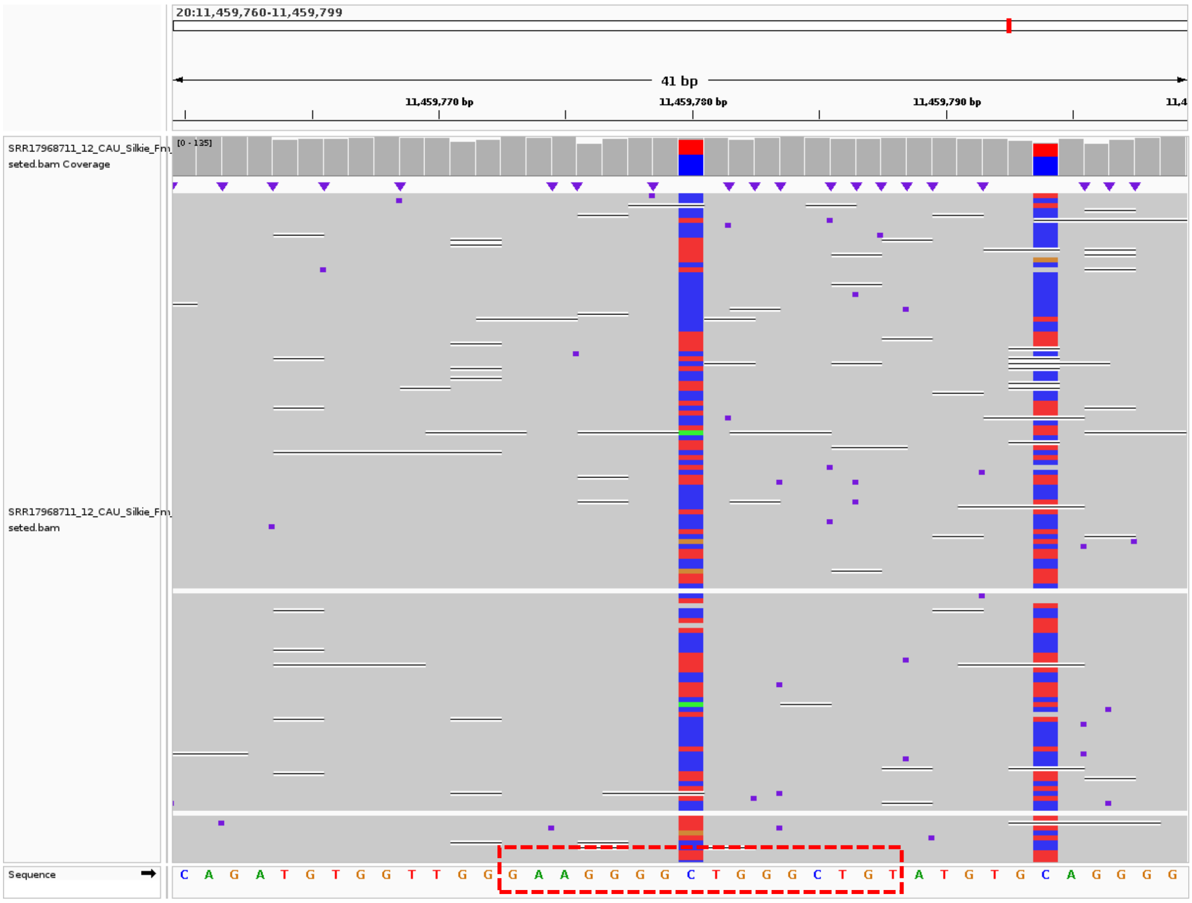


**Supplementary Figure S60:** IGV screenshot displaying Dup2 HDP 11459780 present within a homopolymer run of G (4 bases) in a 10-mer window, highlighted by a red dotted rectangular box. The visualization shows comparable read support for both haplotypes, indicating that the HDP is unlikely due to sequencing errors.

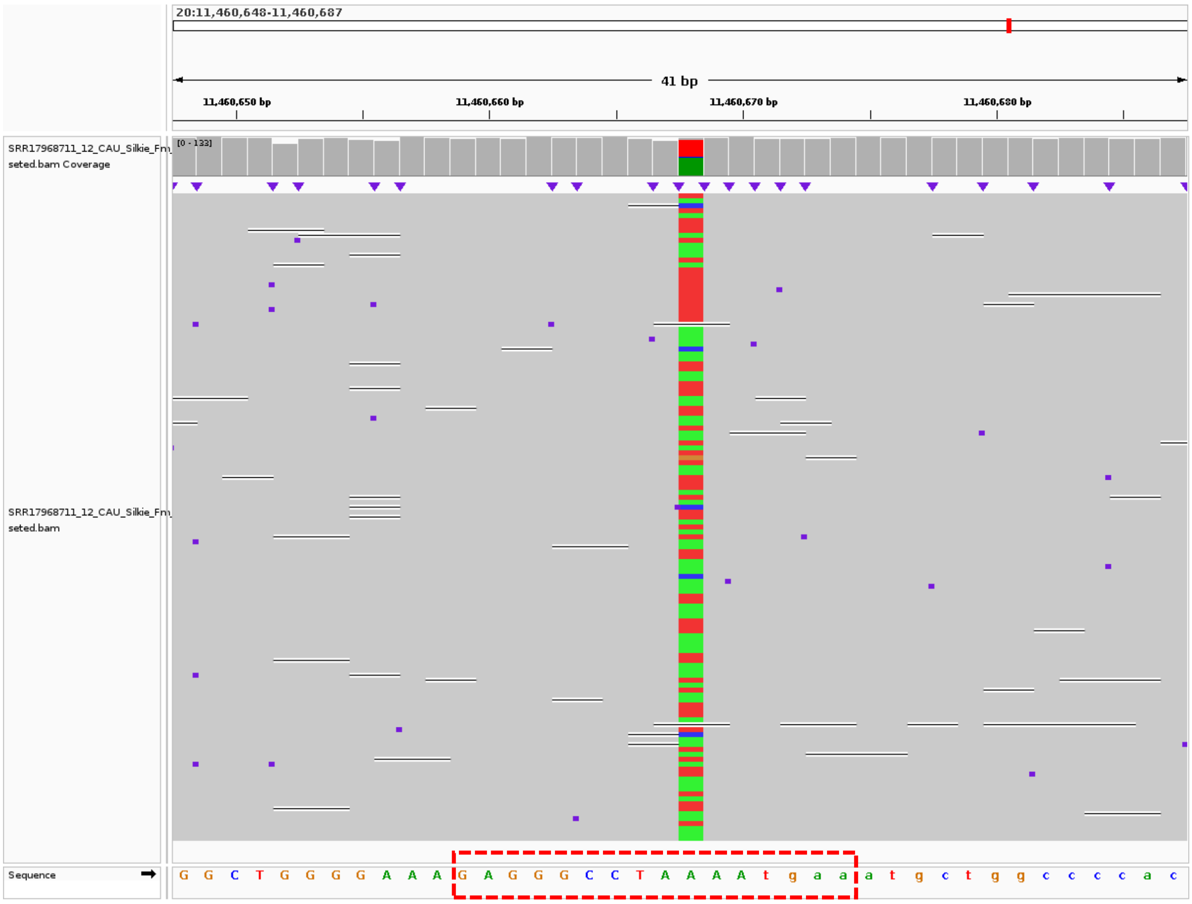


**Supplementary Figure S61:** IGV screenshot displaying Dup2 HDP 11460668 present within a homopolymer run of A (4 bases) in a 10-mer window, highlighted by a red dotted rectangular box. The visualization shows comparable read support for both haplotypes, indicating that the HDP is unlikely due to sequencing errors.

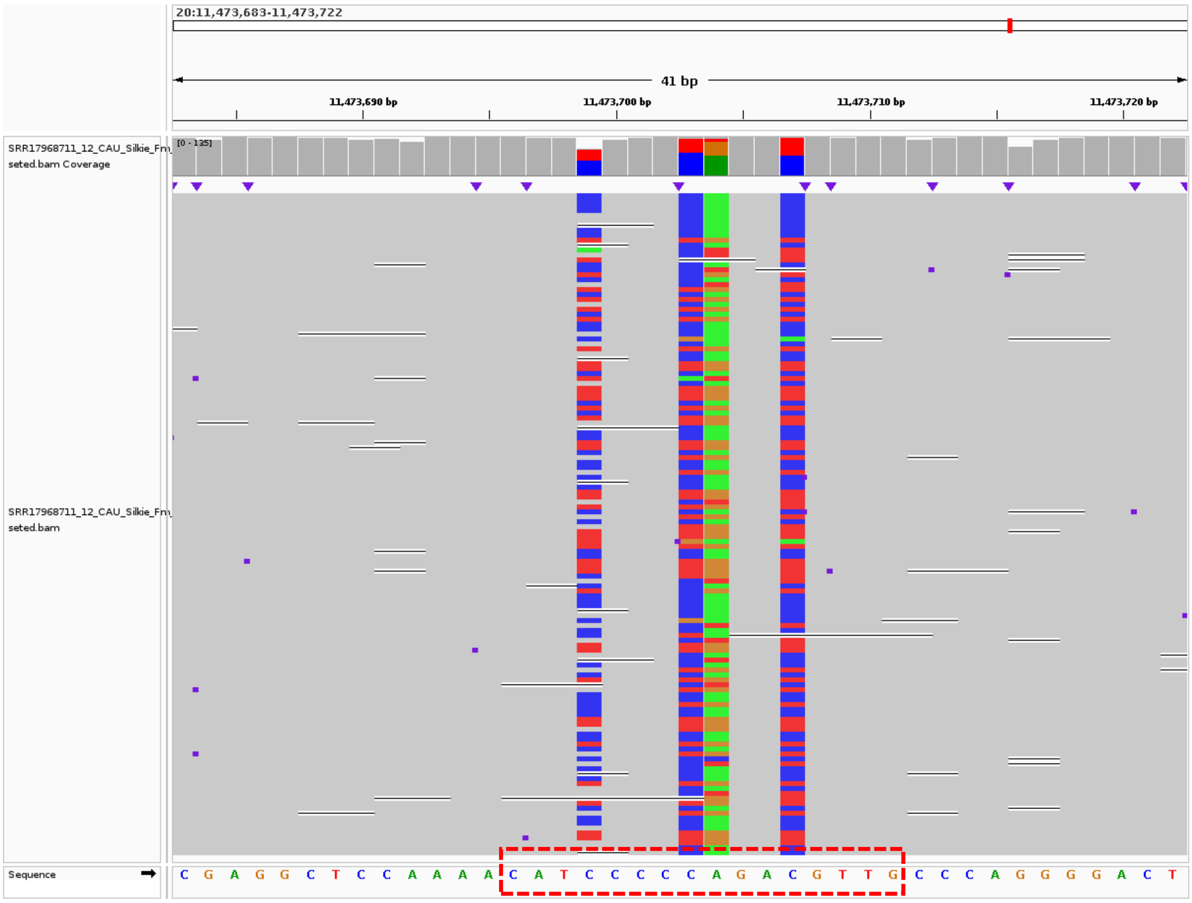


**Supplementary Figure S62:** IGV screenshot displaying Dup2 HDPs 11473699 and 11473703 present within a homopolymer run of C (5 bases) in a 10-mer window, highlighted by a red dotted rectangular box. The visualization shows comparable read support for both haplotypes, indicating that these HDPs are unlikely due to sequencing errors.

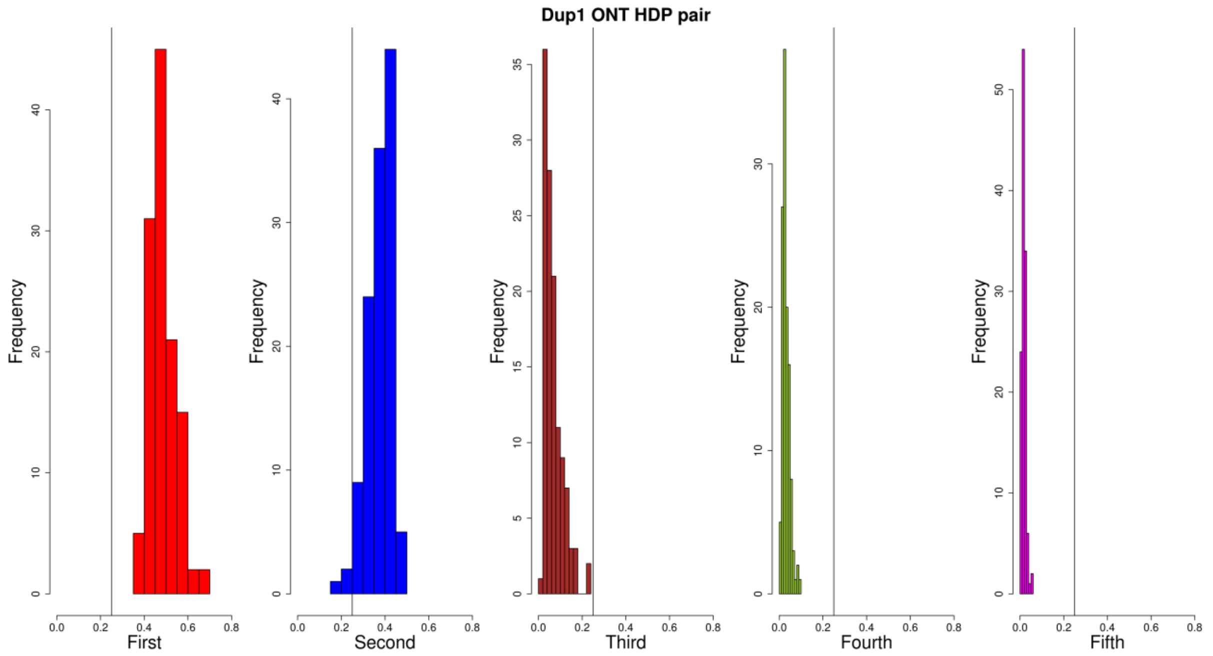


**Supplementary Figure. S63:** Histogram of the ratio of read pair count between HDPs for the Dup1 region, based on CAU Silkie ONT long-reads mapped to the GRCg6a genome assembly. The ratio for the first and second pairs is much higher than for the remaining three pairs, based on a defined threshold of 0.25. This suggests that the majority of HDPs have two alleles.

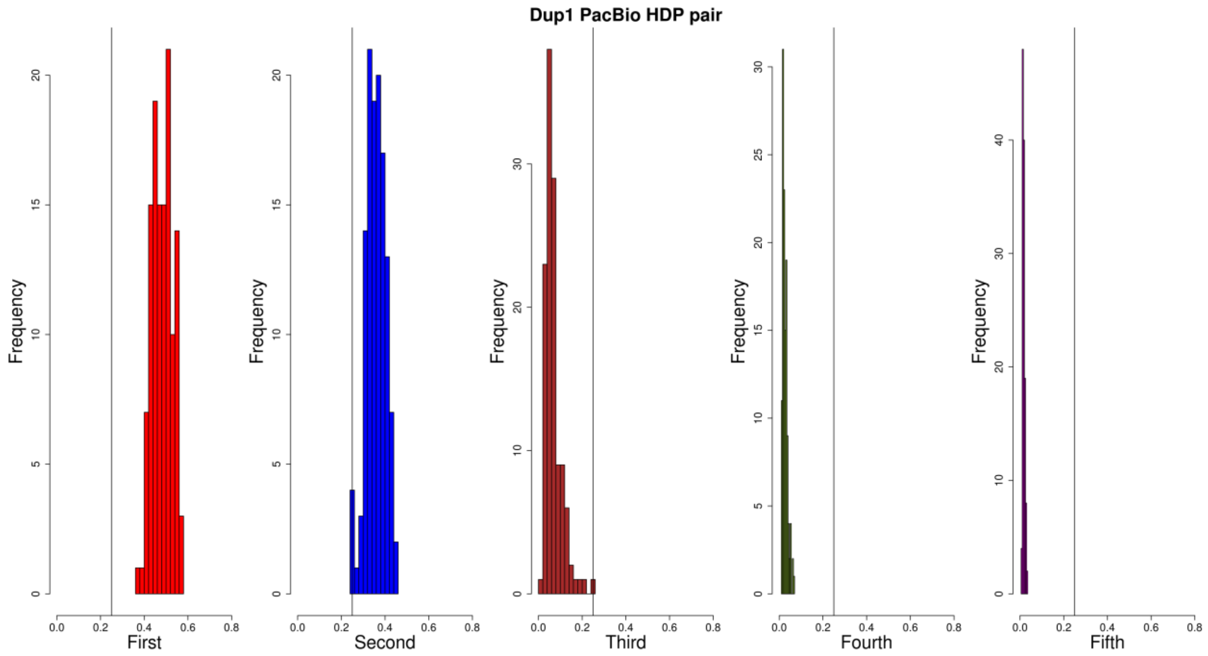


**Supplementary Figure. S64:** Histogram of read count fractions between HDPs for the Dup1 region, based on CAU Silkie PacBio long-reads mapped to the GRCg6a genome assembly. The read counts for the first and second pairs are much higher than for the remaining three pairs, based on a defined threshold of 0.25. This suggests that the majority of HDPs have two alleles.

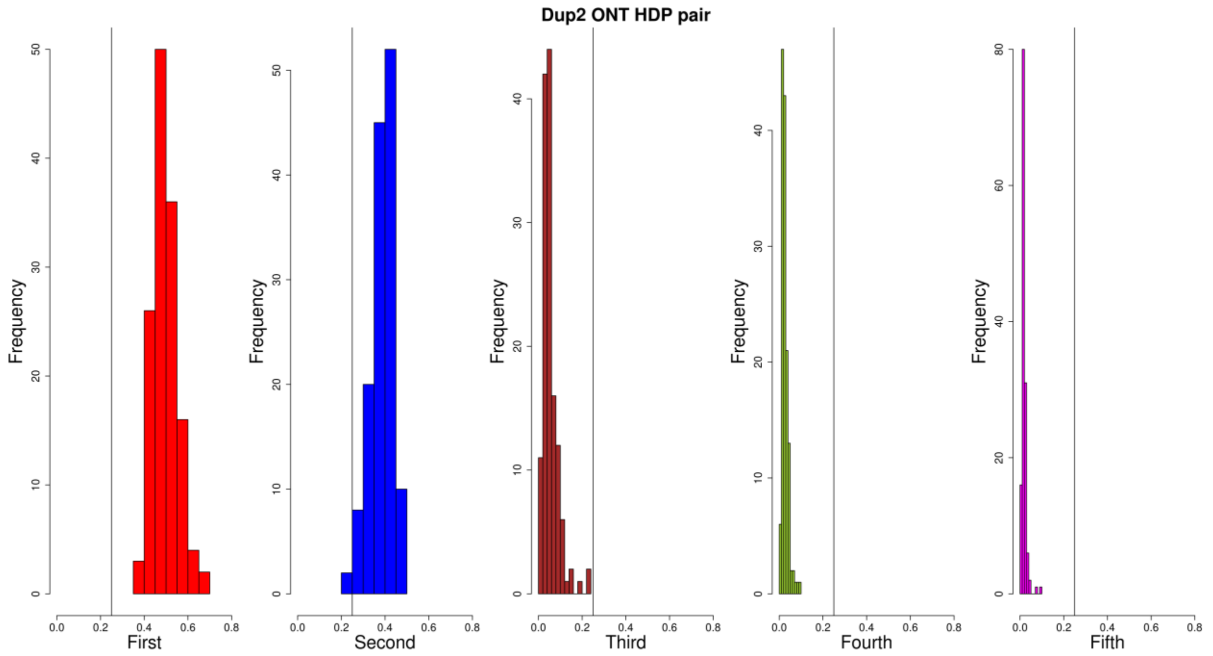


**Supplementary Figure. S65:** Histogram of read count fractions between HDPs for the Dup2 region, based on CAU Silkie ONT long-reads mapped to the GRCg6a genome assembly. The read counts for the first and second pairs are much higher than for the remaining three pairs, based on a defined threshold of 0.25. This suggests that the majority of HDPs have two alleles.

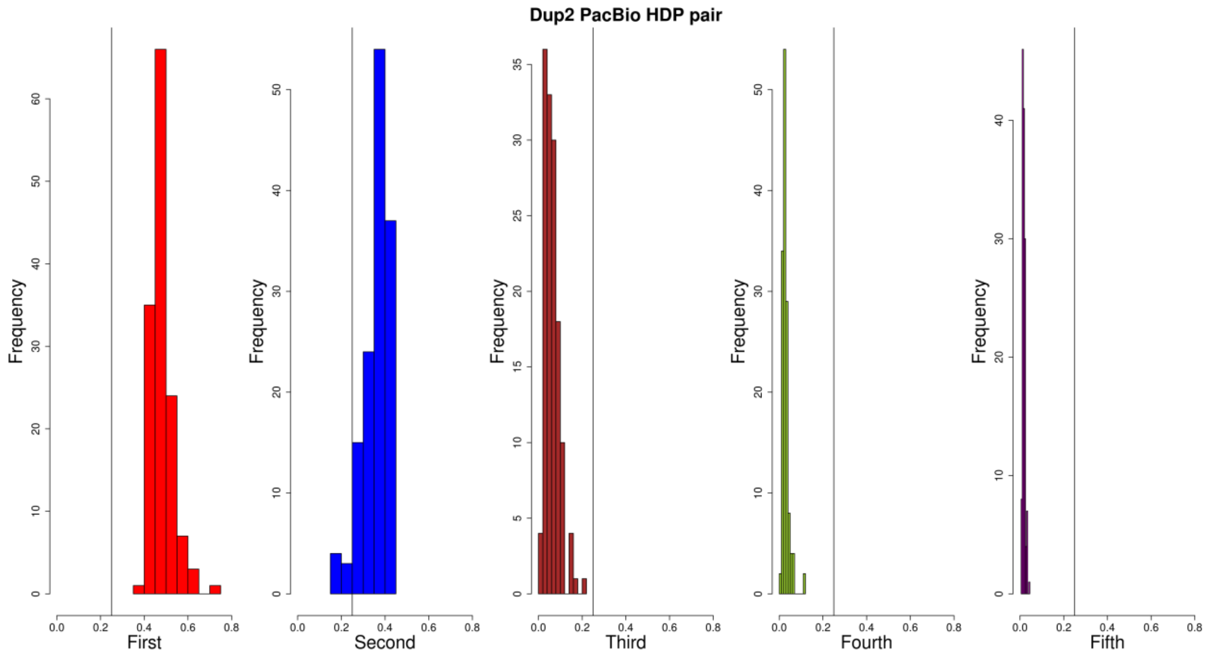


**Supplementary Figure. S66:** Histogram of read count fractions between HDPs for the Dup2 region, based on CAU Silkie PacBio long-reads mapped to the GRCg6a genome assembly. The read counts for the first and second pairs are much higher than for the remaining three pairs, based on a defined threshold of 0.25. This suggests that the majority of HDPs have two alleles.

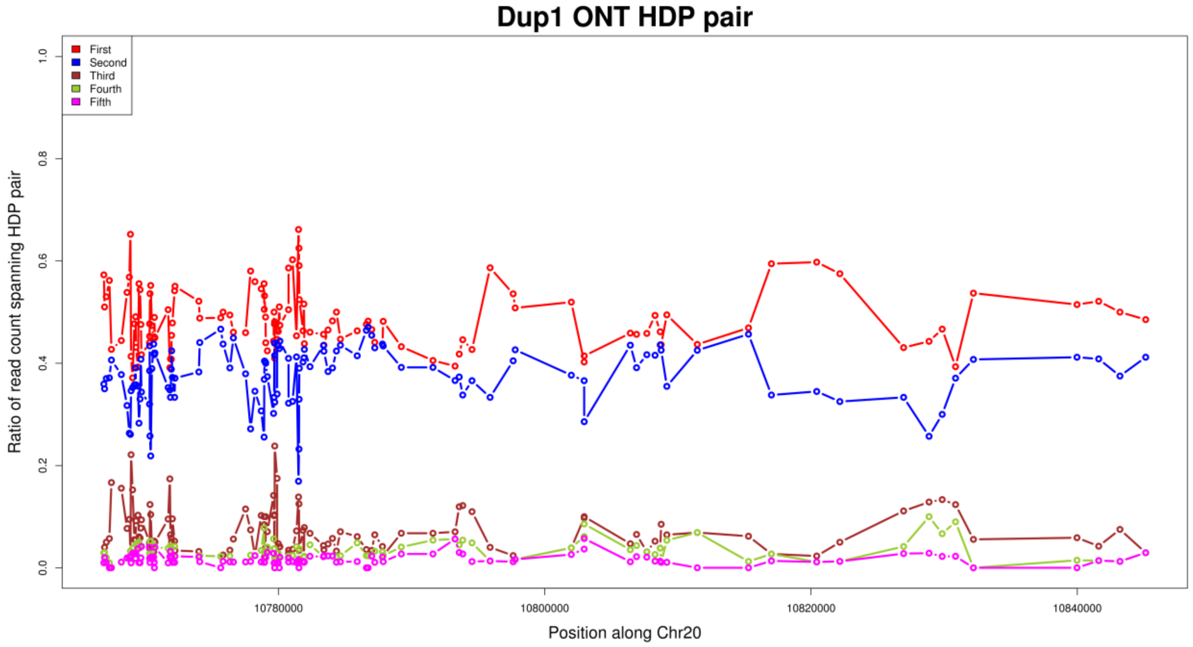


**Supplementary Figure. S67:** Ratio of read pair count between two HDPs for the Dup1 region, based on CAU Silkie ONT long-reads mapped to the GRCg6a genome assembly. The ratio for the first and second pairs is much higher than for the remaining three pairs, suggesting that most HDPs have two alleles. Some HDPs show an increased count for the third pair, which is attributed to within-haplotype polymorphisms.

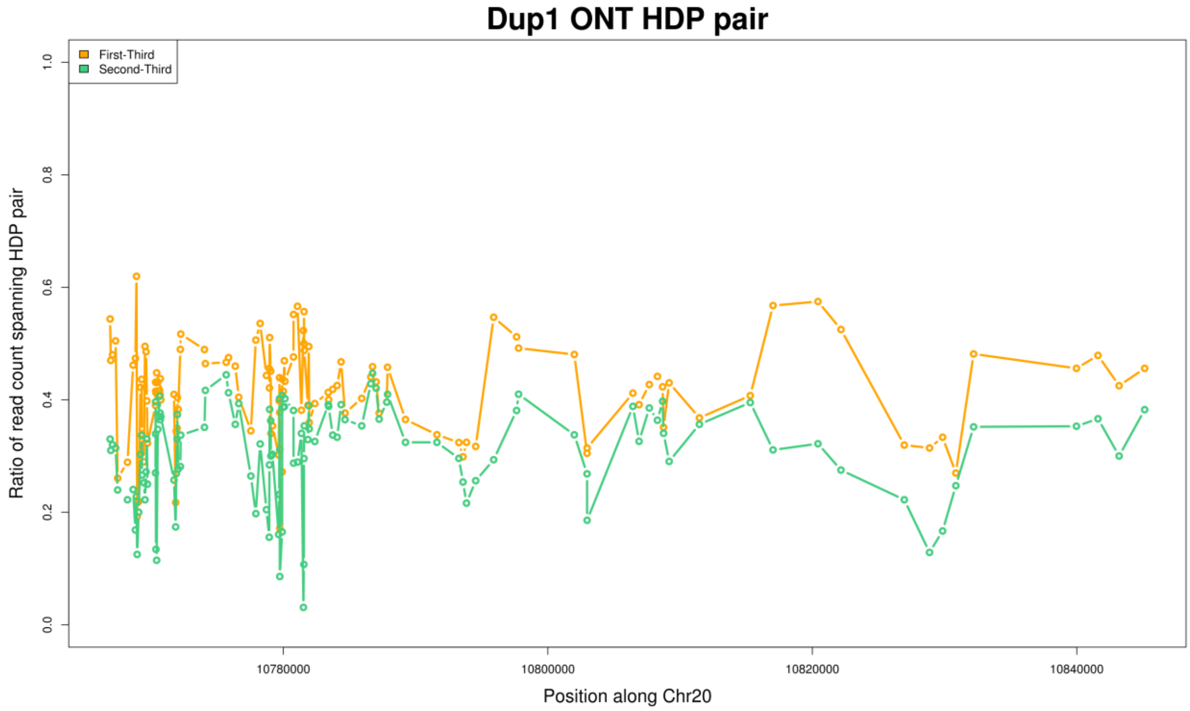


**Supplementary Figure. S68:** Ratio of read pair count for first and second pairs between two HDPs for the Dup1 region after subtracting the count of the third pair. The ratio for the first and second pairs is still comparable.

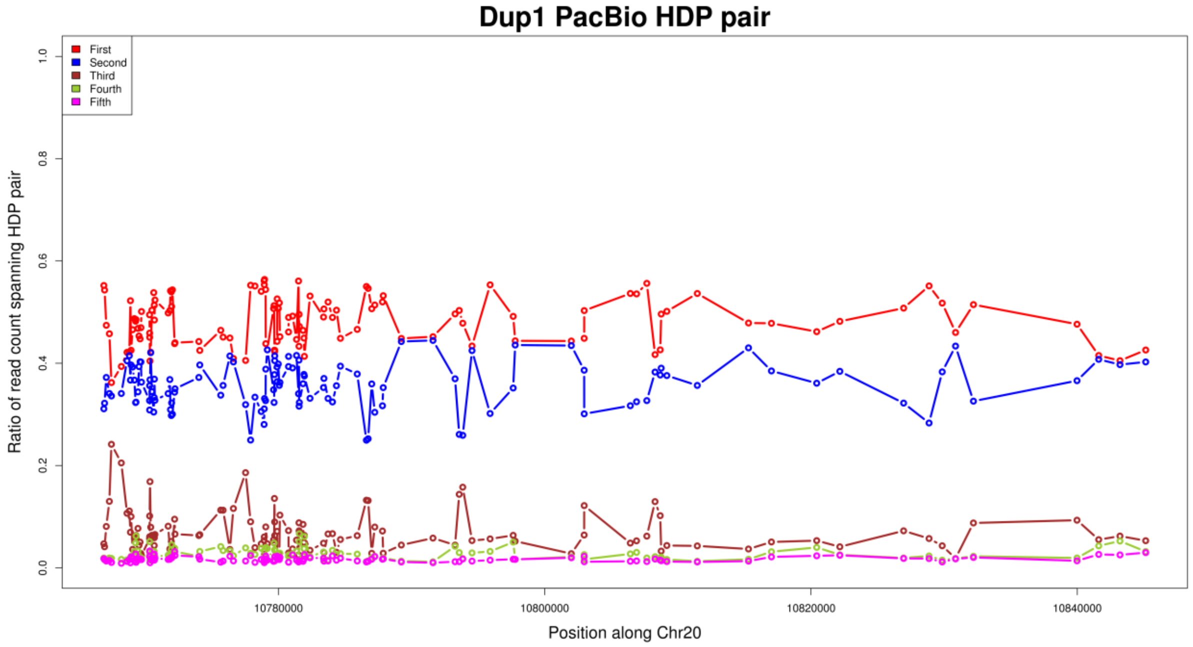


**Supplementary Figure. S69:** Ratio of read pair count between two HDPs for the Dup1 region, based on CAU Silkie PacBio long-reads mapped to the GRCg6a genome assembly. The ratio for the first and second pairs is much higher than for the remaining three pairs, suggesting that most HDPs have two alleles. Some HDPs show an increased count for the third pair, which is attributed to within-haplotype polymorphisms.

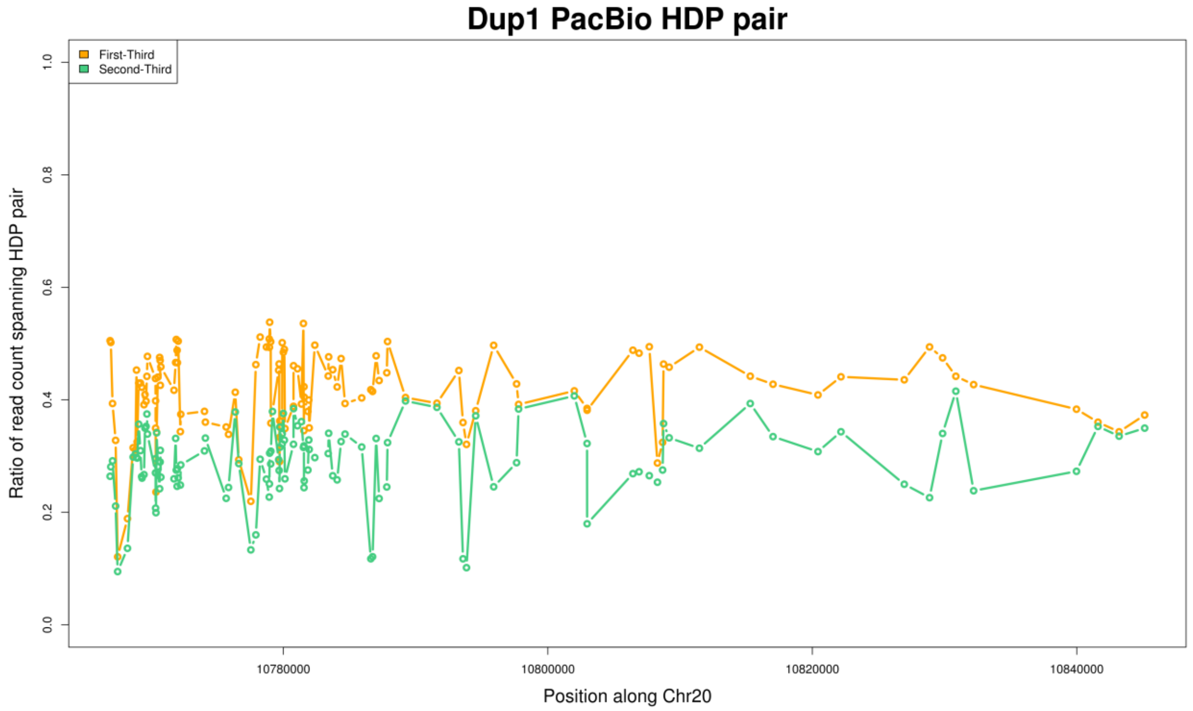


**Supplementary Figure. S70:** Ratio of read pair count for first and second pairs between two HDPs for the Dup1 region after subtracting the count of the third pair. The ratio for the first and second pairs is still comparable.

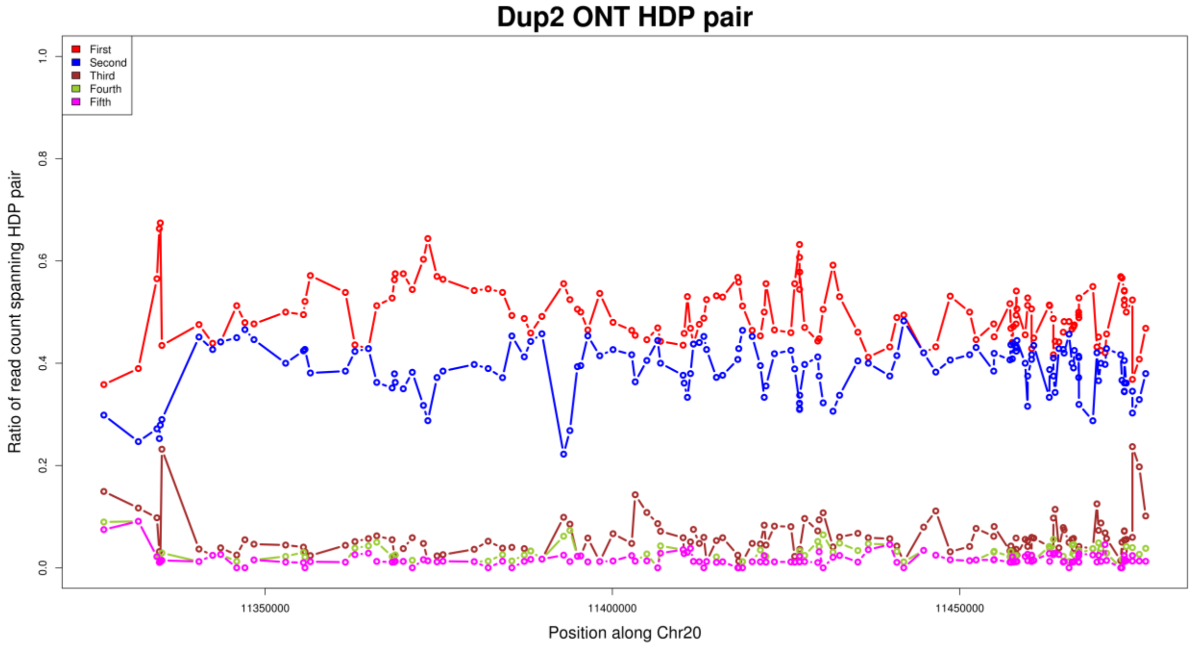


**Supplementary Figure. S71:** Ratio of read pair count between two HDPs for the Dup2 region, based on CAU Silkie ONT long-reads mapped to the GRCg6a genome assembly. The read counts for the first and second pairs are much higher than for the remaining three pairs, suggesting that the majority of HDPs have two alleles. Some HDPs show an increased count for the third pair, which is attributed to within-haplotype polymorphisms.

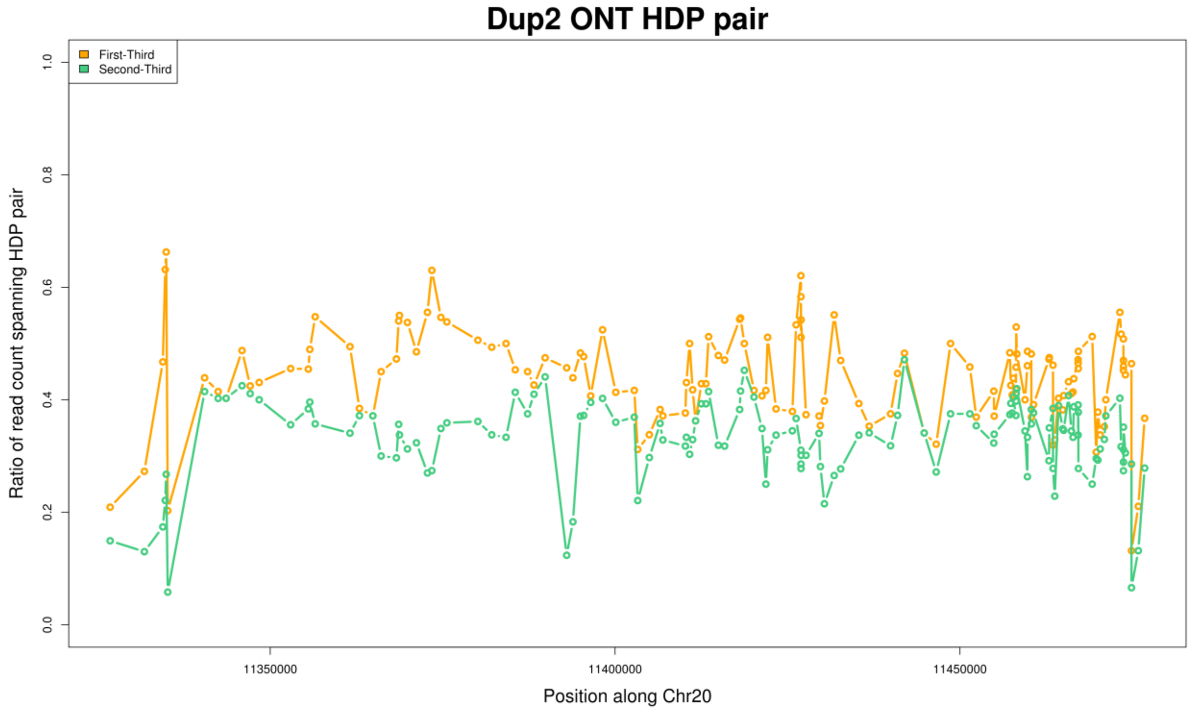


**Supplementary Figure. S72:** Ratio of read pair count for first and second pairs between two HDPs for the Dup2 region after subtracting the count of the third pair. The ratio for the first and second pairs is still comparable.

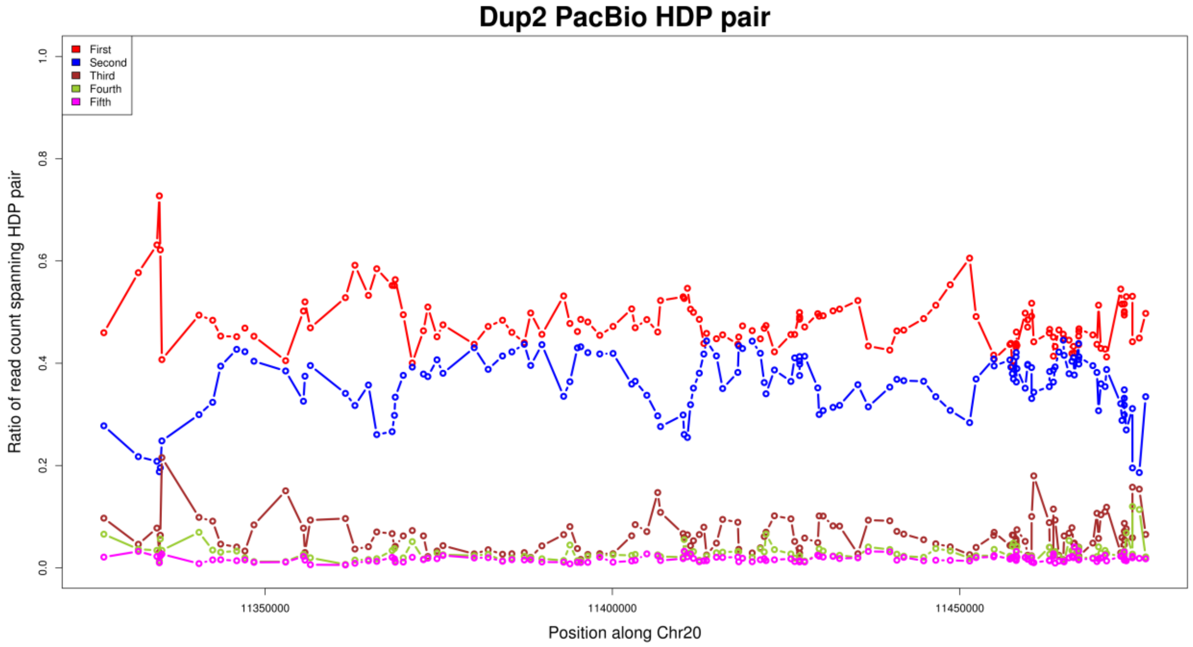


**Supplementary Figure. S73:** Ratio of read pair count between two HDPs for the Dup2 region, based on CAU Silkie PacBio long-reads mapped to the GRCg6a genome assembly. The read counts for the first and second pairs are much higher than for the remaining three pairs, suggesting that the majority of HDPs have two alleles. Some HDPs show an increased count for the third pair, which is attributed to within-haplotype polymorphisms.

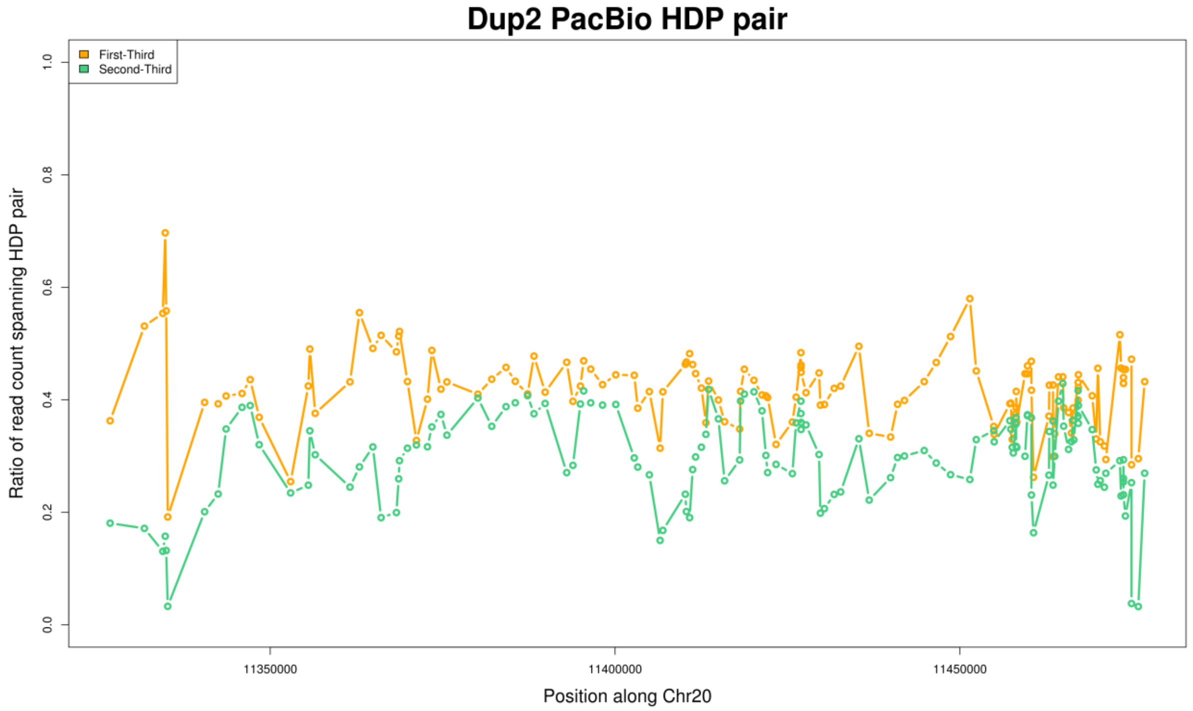


**Supplementary Figure. S74:** Ratio of read pair count for first and second pairs between two HDPs for the Dup2 region after subtracting the count of the third pair. The ratio for the first and second pairs is still comparable.

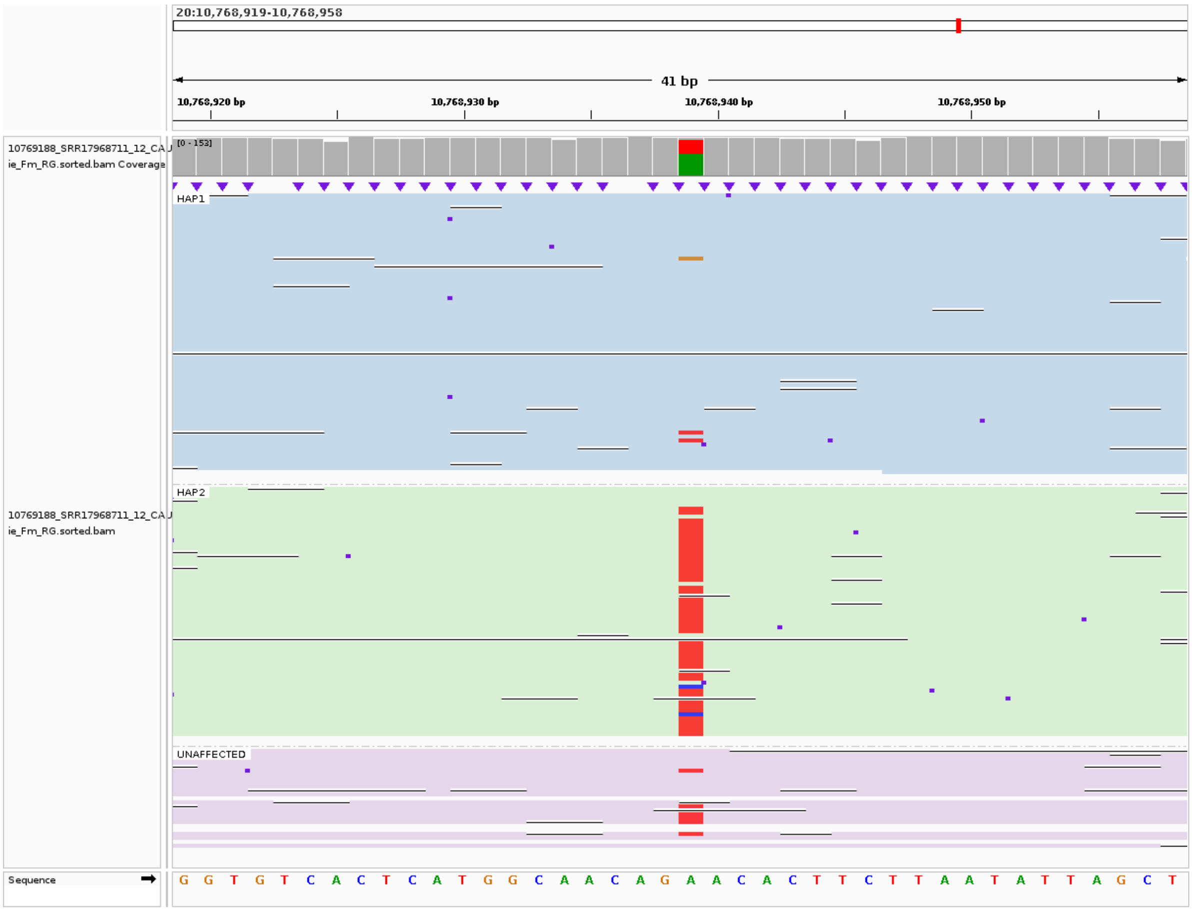


**Supplementary Figure. S75:** The IGV screenshot shows Dup1 HDP 10768939 with a within-haplotype polymorphism in the CAU Silkie ONT long-read data, mapped to the GRCg6a genome assembly. The light blue panel represents Dup1 haplotype1, the light green panel represents Dup1 haplotype2, and the light pink panel represents the unassigned reads.

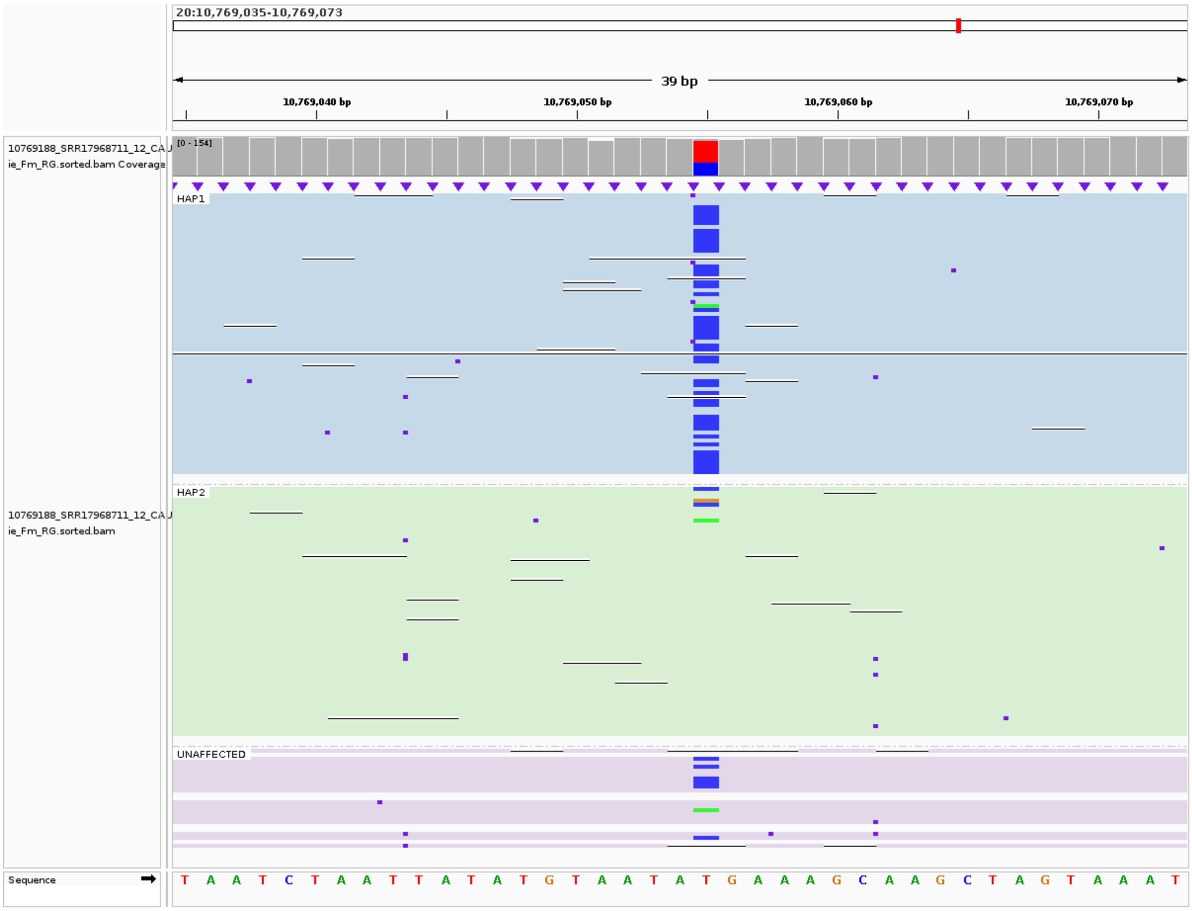


**Supplementary Figure. S76:** The IGV screenshot shows Dup1 HDP 10769055 with a within-haplotype polymorphism in the CAU Silkie ONT long-read data, mapped to the GRCg6a genome assembly. The light blue panel represents Dup1 haplotype1, the light green panel represents Dup1 haplotype2, and the light pink panel represents the unassignedreads.

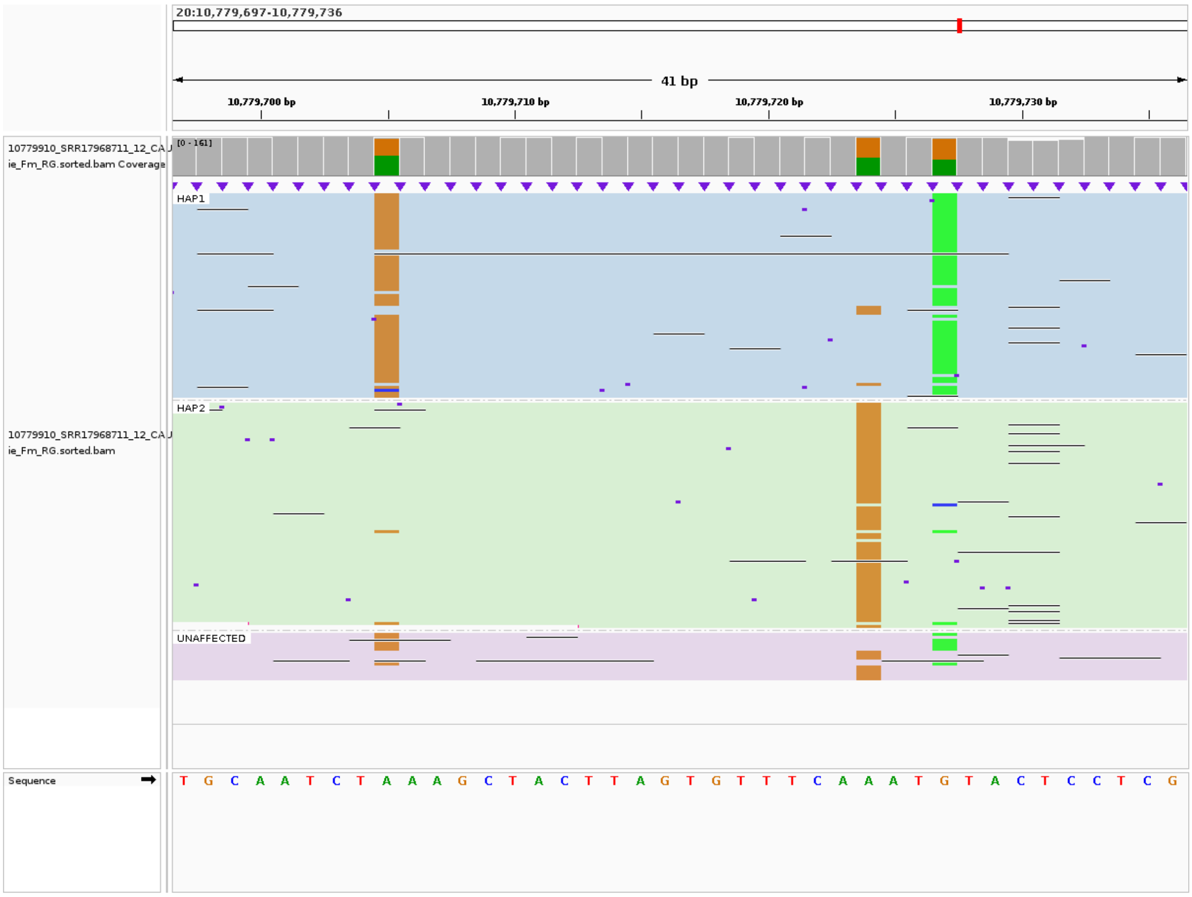


**Supplementary Figure. S77:** The IGV screenshot shows Dup1 HDPs 10779705, 10779724, and 10779727 with a within-haplotype polymorphism in the CAU Silkie ONT long-read data, mapped to the GRCg6a genome assembly. The light blue panel represents Dup1 haplotype1, the light green panel represents Dup1 haplotype2, and the light pink panel represents the unassigned reads.

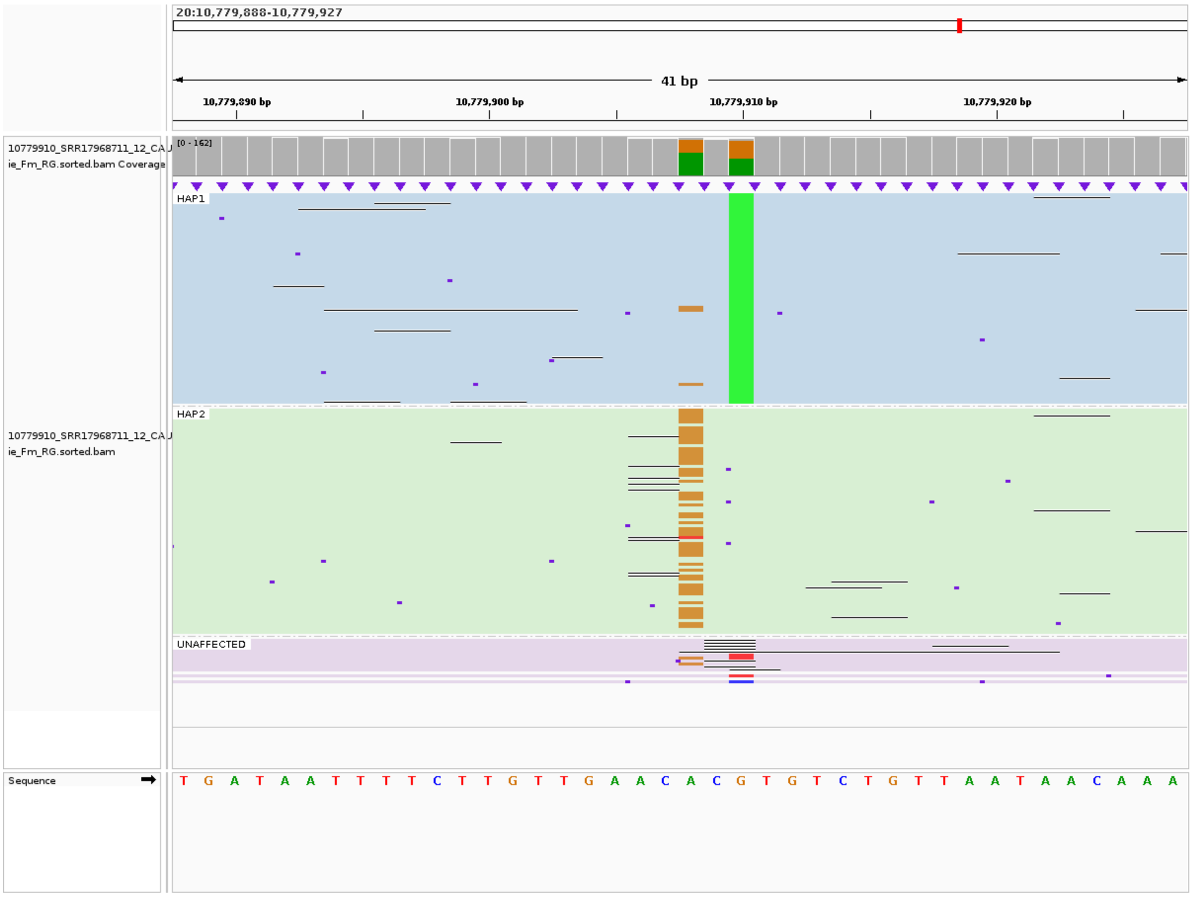


**Supplementary Figure. S78:** The IGV screenshot shows Dup1 HDP 10779908 with a within-haplotype polymorphism in the CAU Silkie ONT long-read data, mapped to the GRCg6a genome assembly. The light blue panel represents Dup1 haplotype1, the light green panel represents Dup1 haplotype2, and the light pink panel represents the unassigned reads.

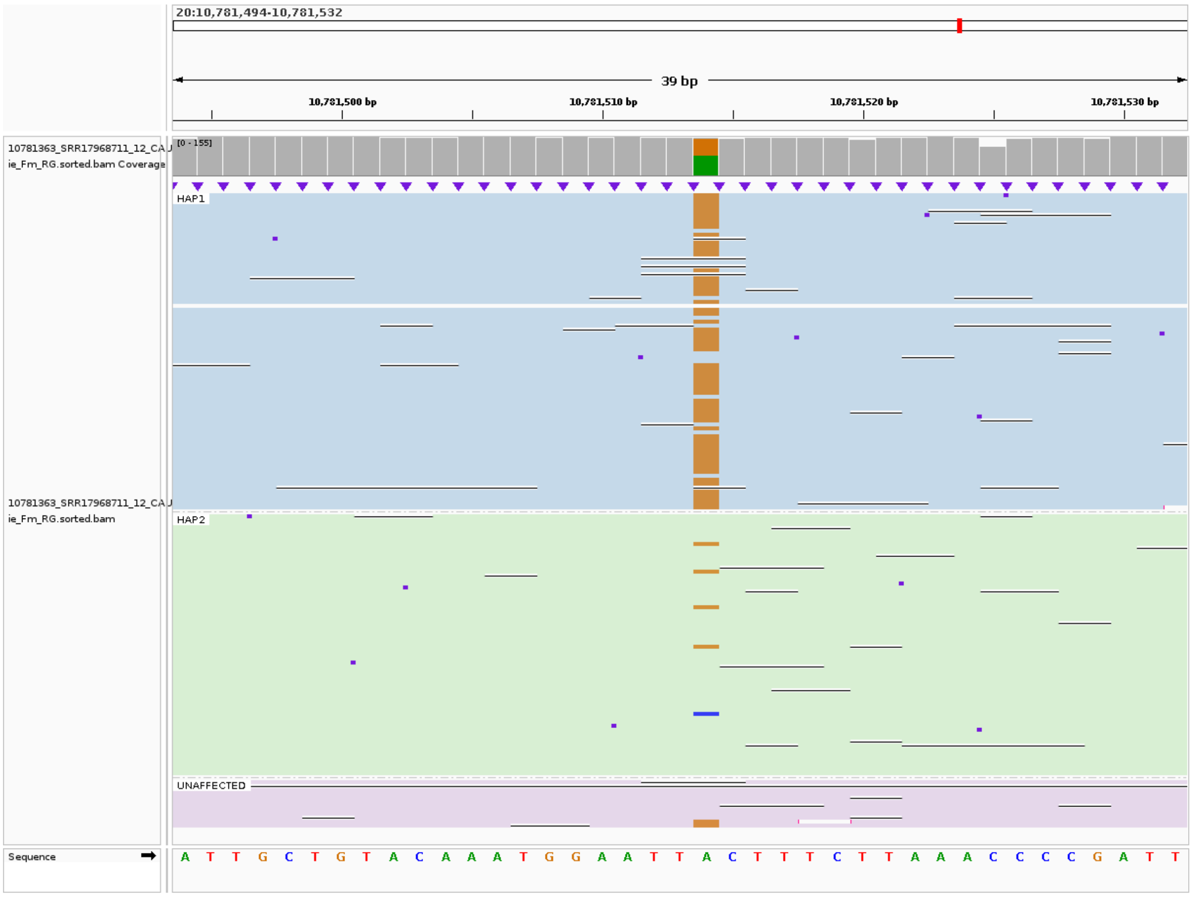


**Supplementary Figure. S79:** The IGV screenshot shows Dup1 HDP 10781514 with a within-haplotype polymorphism in the CAU Silkie ONT long-read data, mapped to the GRCg6a genome assembly. The light blue panel represents Dup1 haplotype1, the light green panel represents Dup1 haplotype2, and the light pink panel represents the unassigned reads.

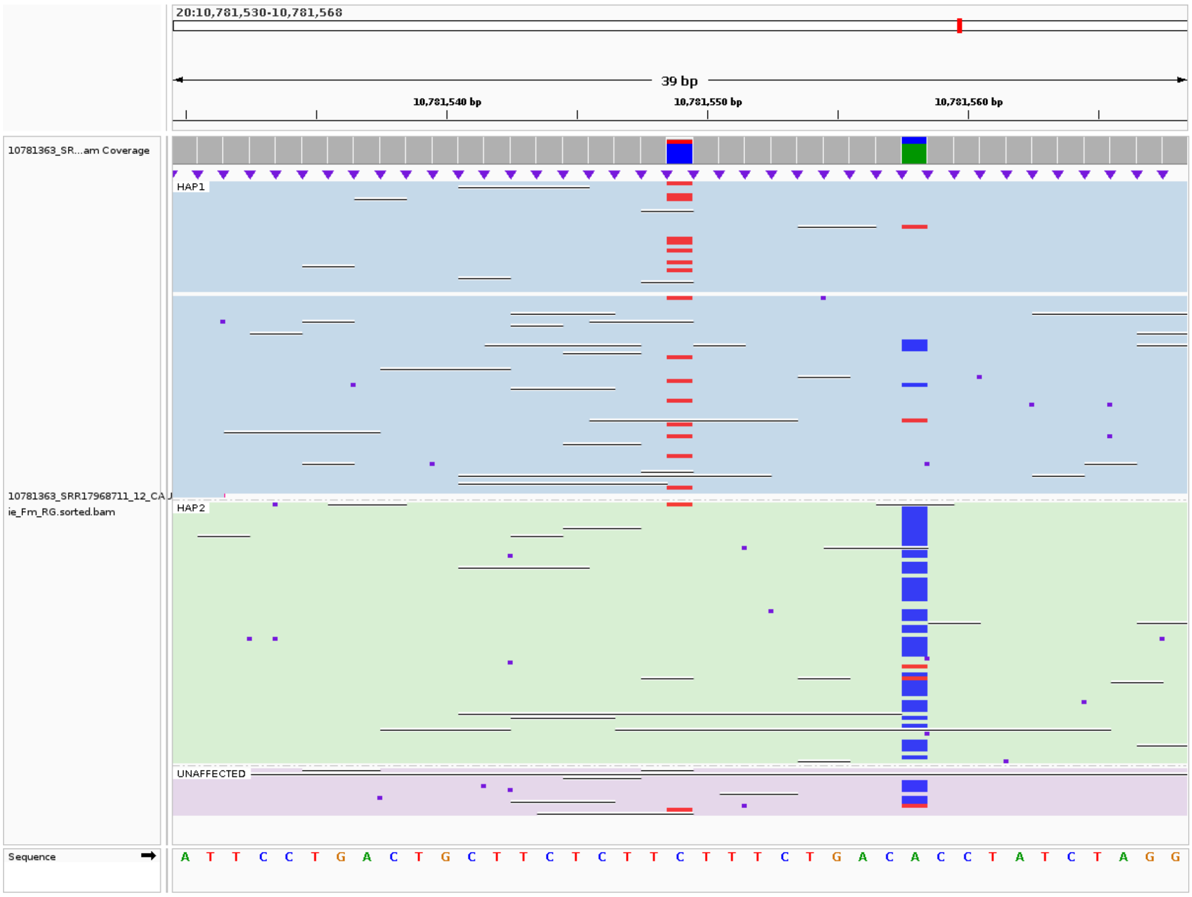


**Supplementary Figure. S80:** The IGV screenshot shows Dup1 HDPs 10781549 and 10781558 with a within-haplotype polymorphism in the CAU Silkie ONT long-read data, mapped to the GRCg6a genome assembly. The light blue panel represents Dup1 haplotype1, the light green panel represents Dup1 haplotype2, and the light pink panel represents the unassigned reads.

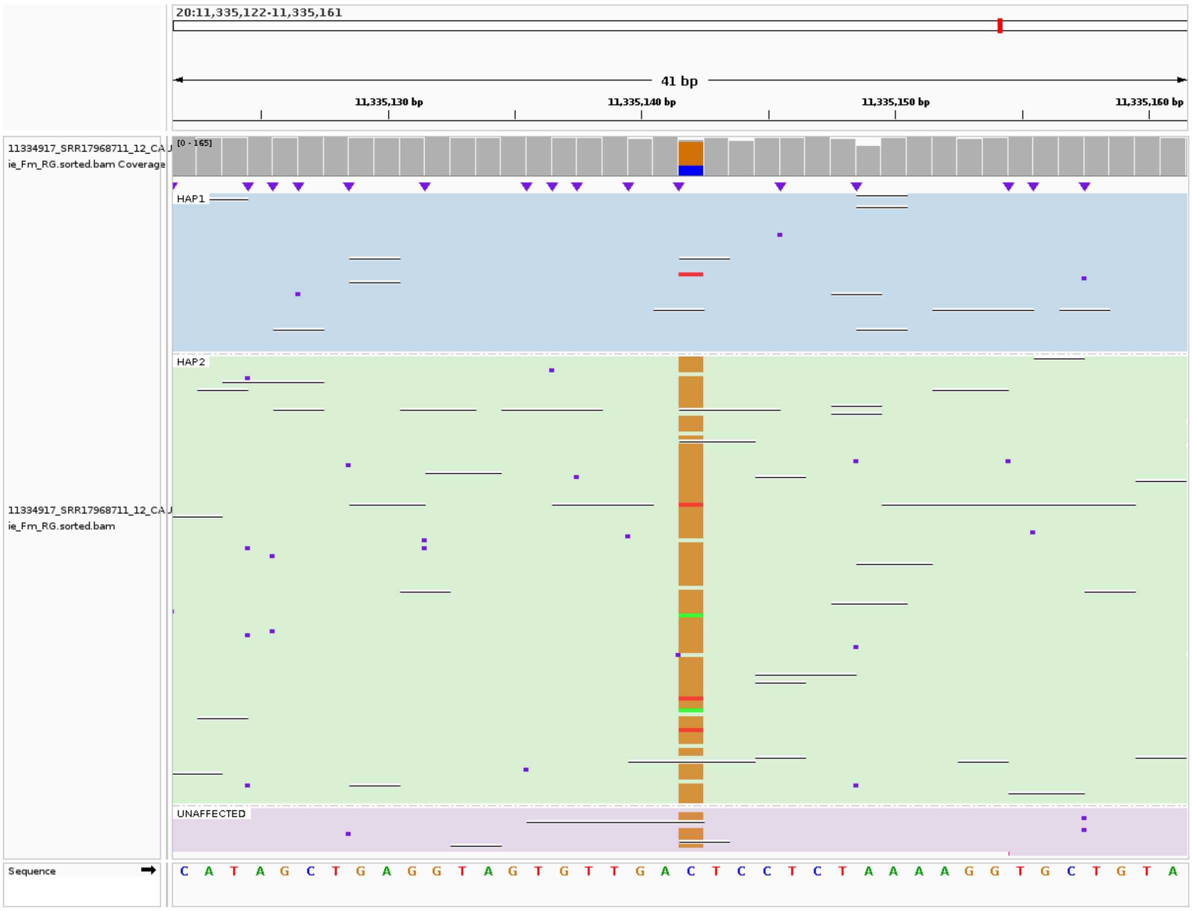


**Supplementary Figure. S81:** The IGV screenshot shows Dup2 HDP 11335142 with a within-haplotype polymorphism in the CAU Silkie ONT long-read data, mapped to the GRCg6a genome assembly. The light blue panel represents Dup1 haplotype1, the light green panel represents Dup1 haplotype2, and the light pink panel represents the unassigned reads.

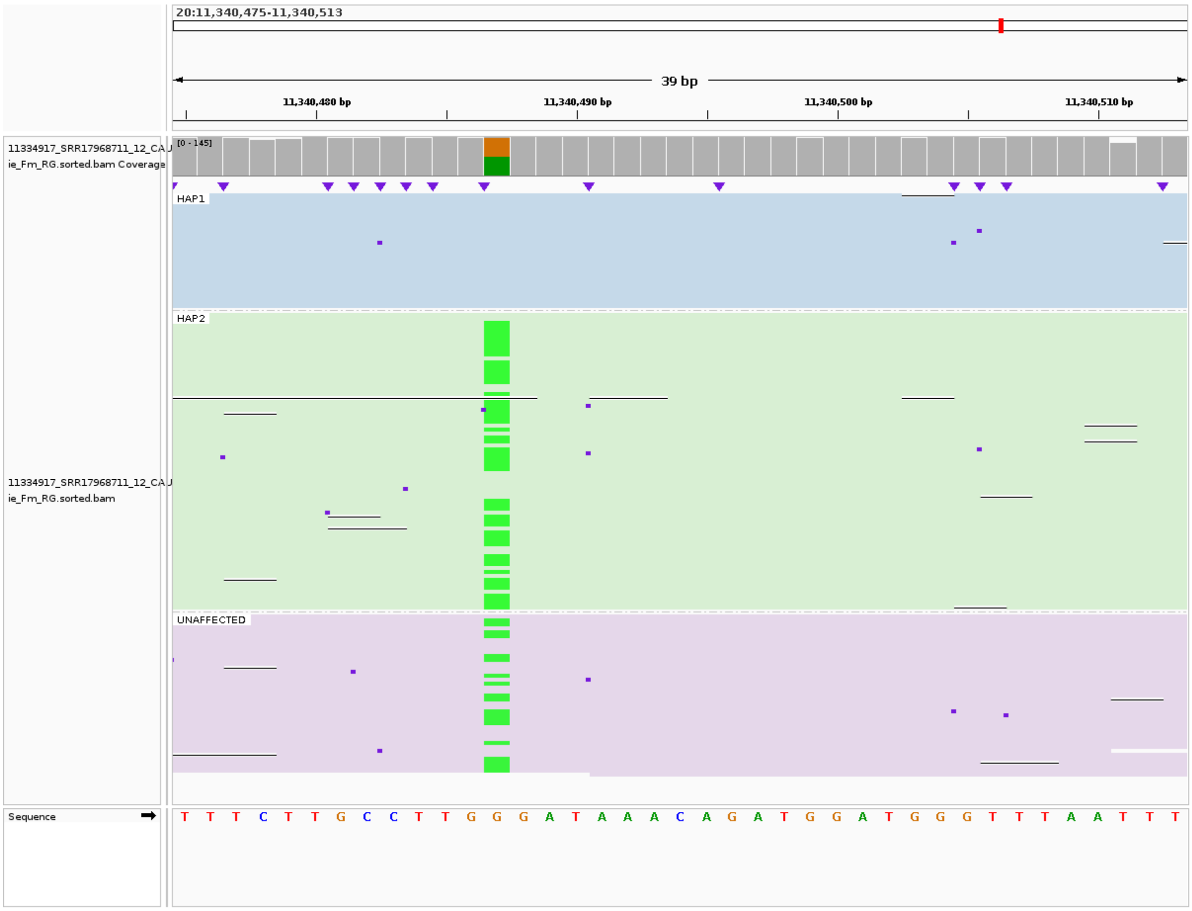


**Supplementary Figure. S82:** The IGV screenshot shows Dup2 HDP 11340487 with a within-haplotype polymorphism in the CAU Silkie ONT long-read data, mapped to the GRCg6a genome assembly. The light blue panel represents Dup1 haplotype1, the light green panel represents Dup1 haplotype2, and the light pink panel represents the unassigned reads.

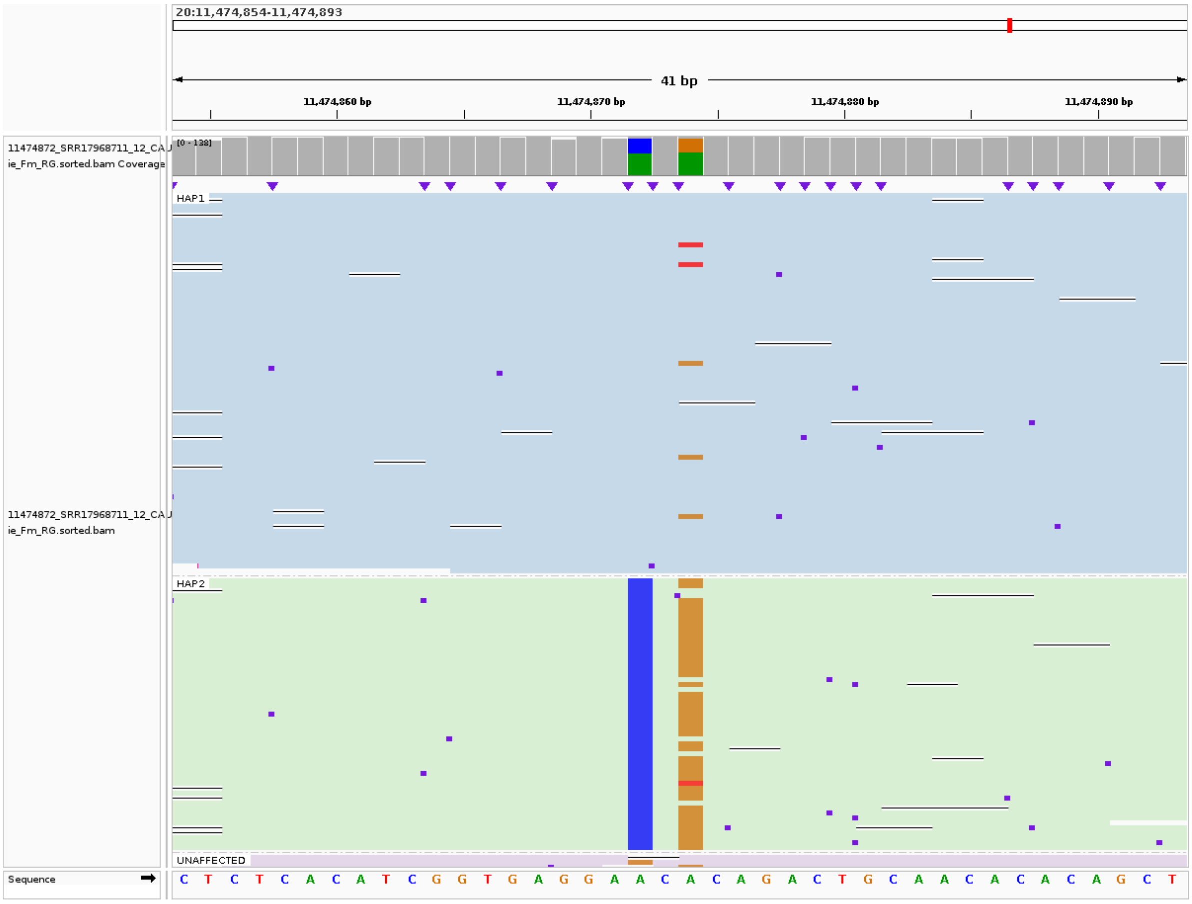


**Supplementary Figure. S83:** The IGV screenshot shows Dup2 HDP 11474874 with a within-haplotype polymorphism in the CAU Silkie ONT long-read data, mapped to the GRCg6a genome assembly. The light blue panel represents Dup1 haplotype1, the light green panel represents Dup1 haplotype2, and the light pink panel represents the unassigned reads.

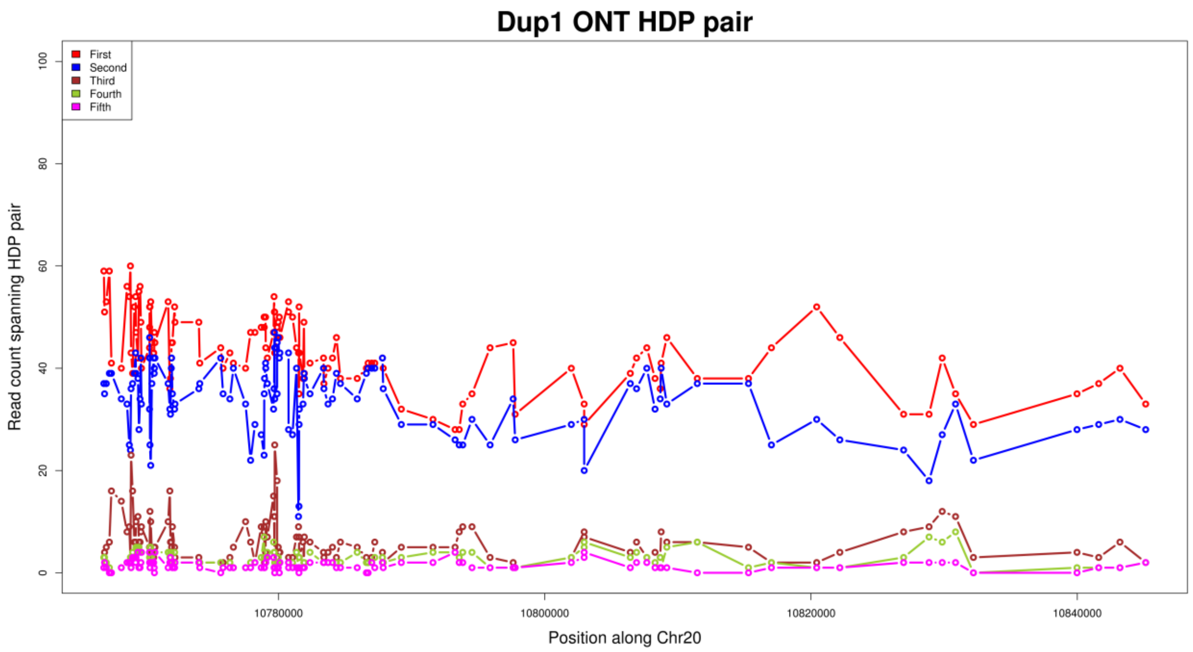


**Supplementary Figure. S84:** Read pair count between two HDPs for the Dup1 region, based on CAU Silkie ONT long-reads mapped to the GRCg6a genome assembly. The read counts for the first and second pairs are much higher than for the remaining three pairs, suggesting that the majority of HDPs have two alleles. Some HDPs show an increased count for the third pair, which is attributed to within-haplotype polymorphisms.

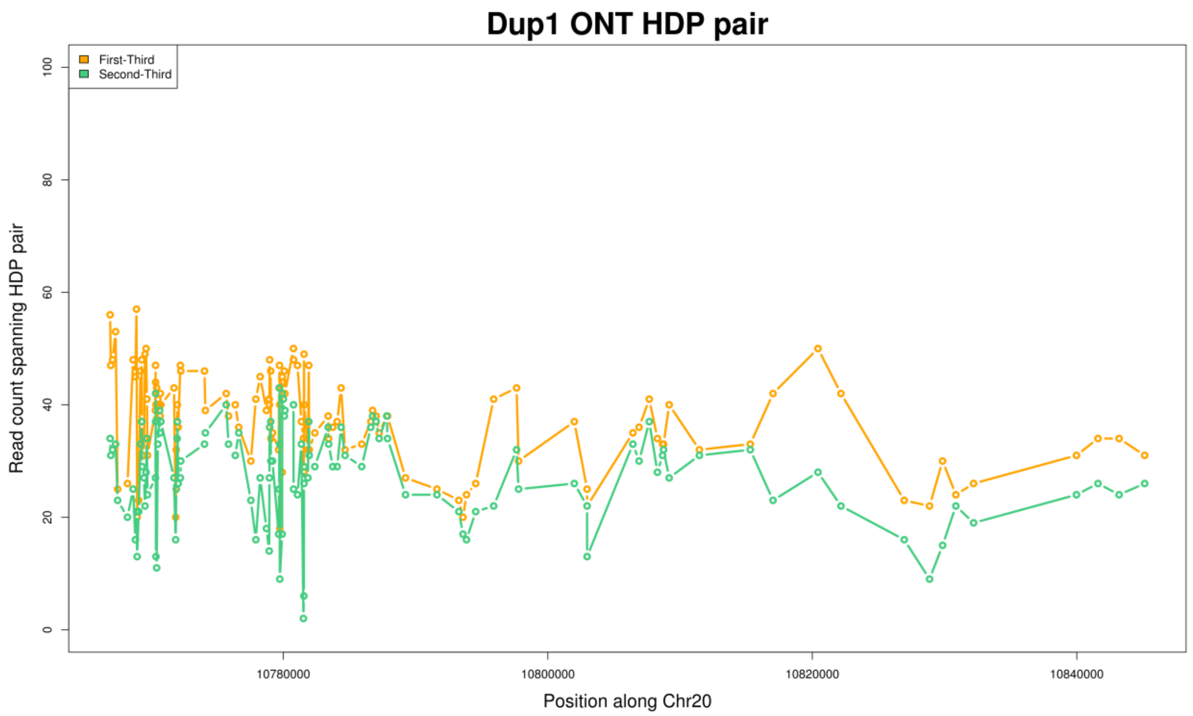


**Supplementary Figure. S85:** Read pair count for the first and second pairs between two HDPs for the Dup1 region after subtracting the count of the third pair. The ratio for the first and second pairs is still comparable.

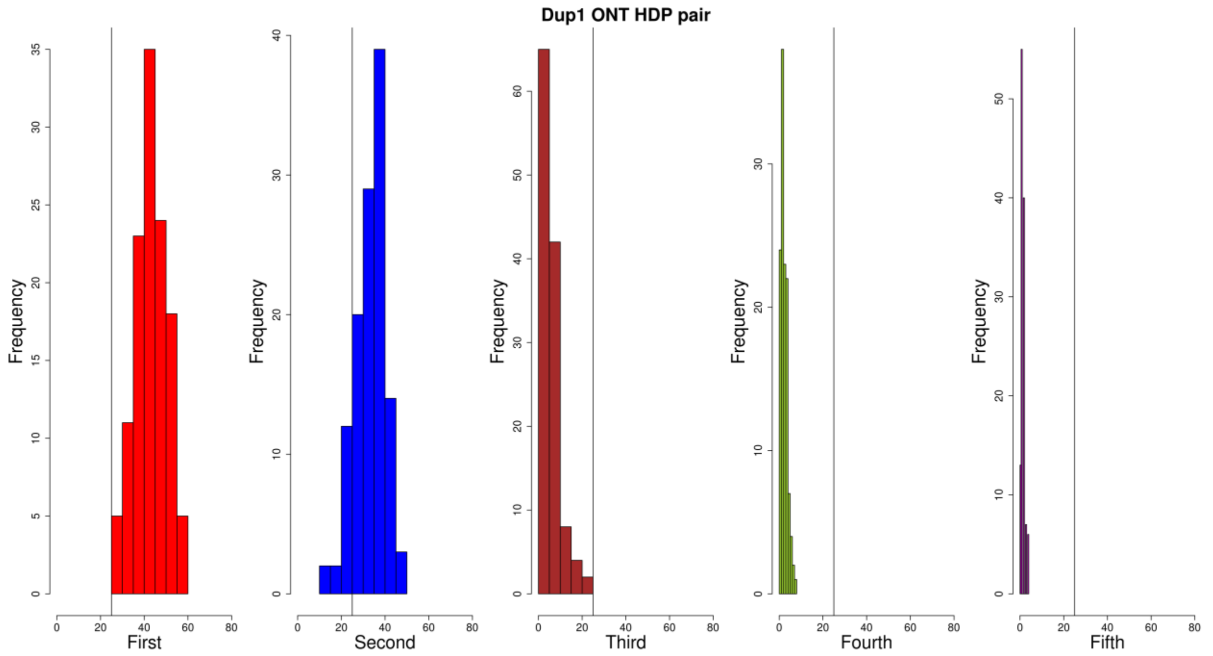


**Supplementary Figure. S86:** Histogram of the read pair count between HDPs for the Dup1 region, based on CAU Silkie ONT long-reads mapped to the GRCg6a genome assembly. The ratio for the first and second pairs is much higher than for the remaining three pairs, based on a defined threshold of 0.25. This suggests that the majority of HDPs have two alleles.

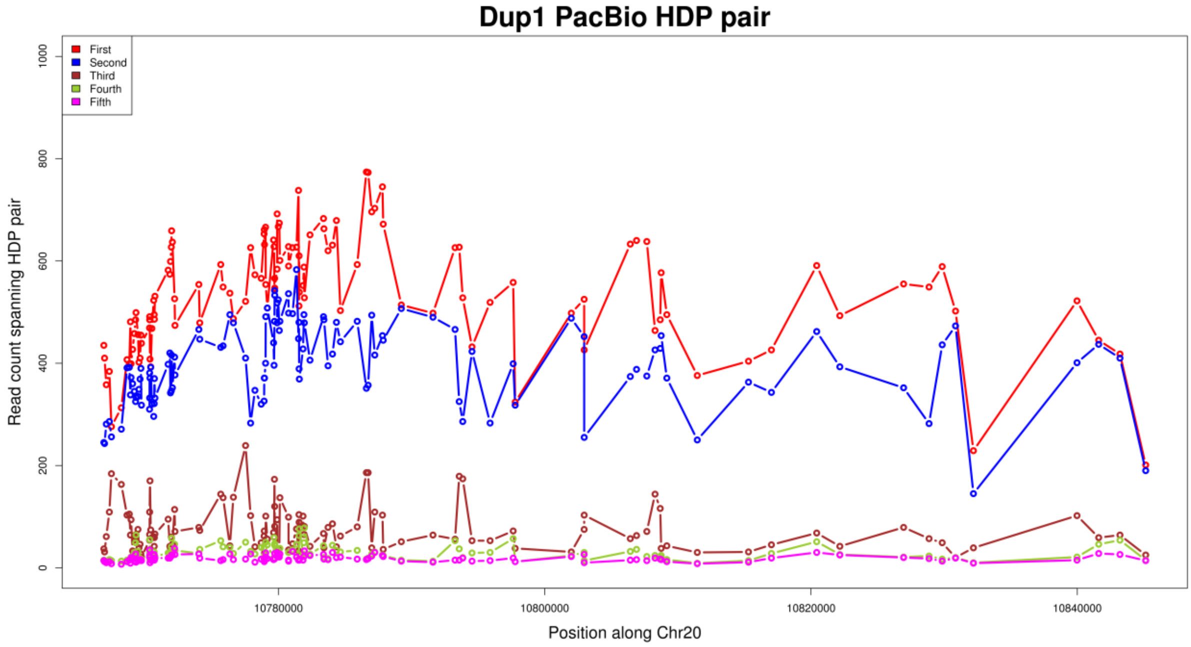


**Supplementary Figure. S87:** Read pair count between two HDPs for the Dup1 region, based on CAU Silkie PacBio long-reads mapped to the GRCg6a genome assembly. The read counts for the first and second pairs are much higher than for the remaining three pairs, suggesting that the majority of HDPs have two alleles. Some HDPs show an increased count for the third pair, which is attributed to within-haplotype polymorphisms.

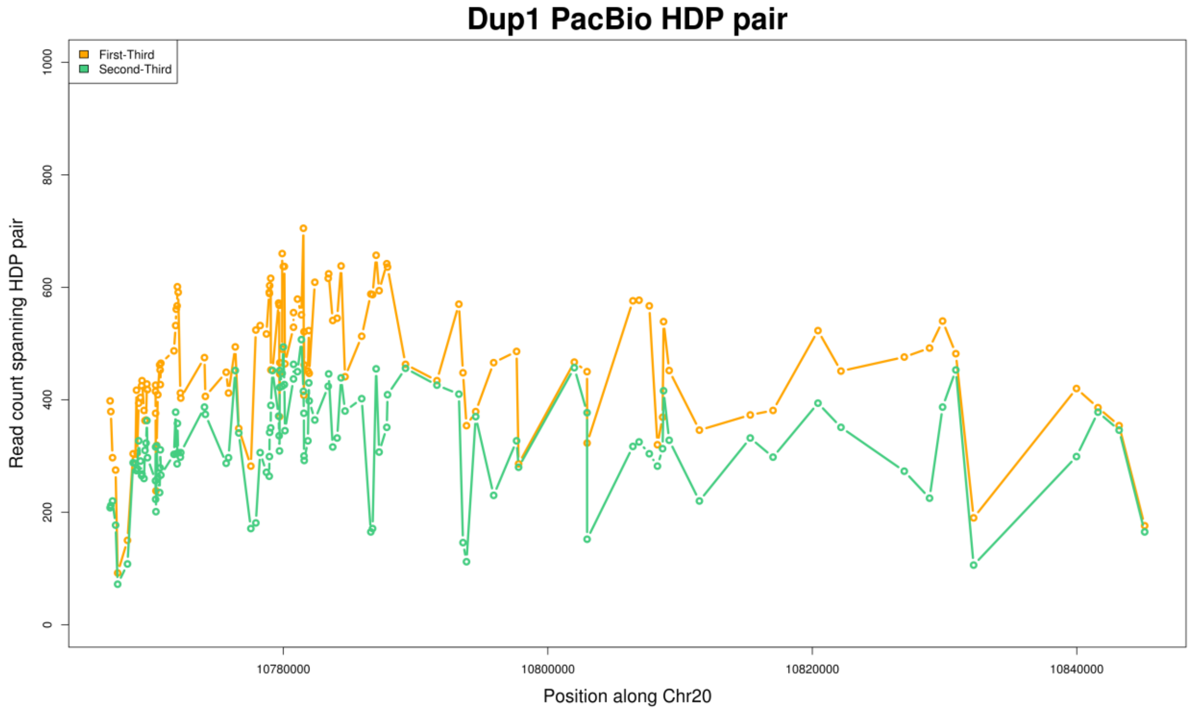


**Supplementary Figure. S88:** Read pair count for the first and second pairs between two HDPs for the Dup1 region after subtracting the count of the third pair. The ratio for the first and second pairs is still comparable.

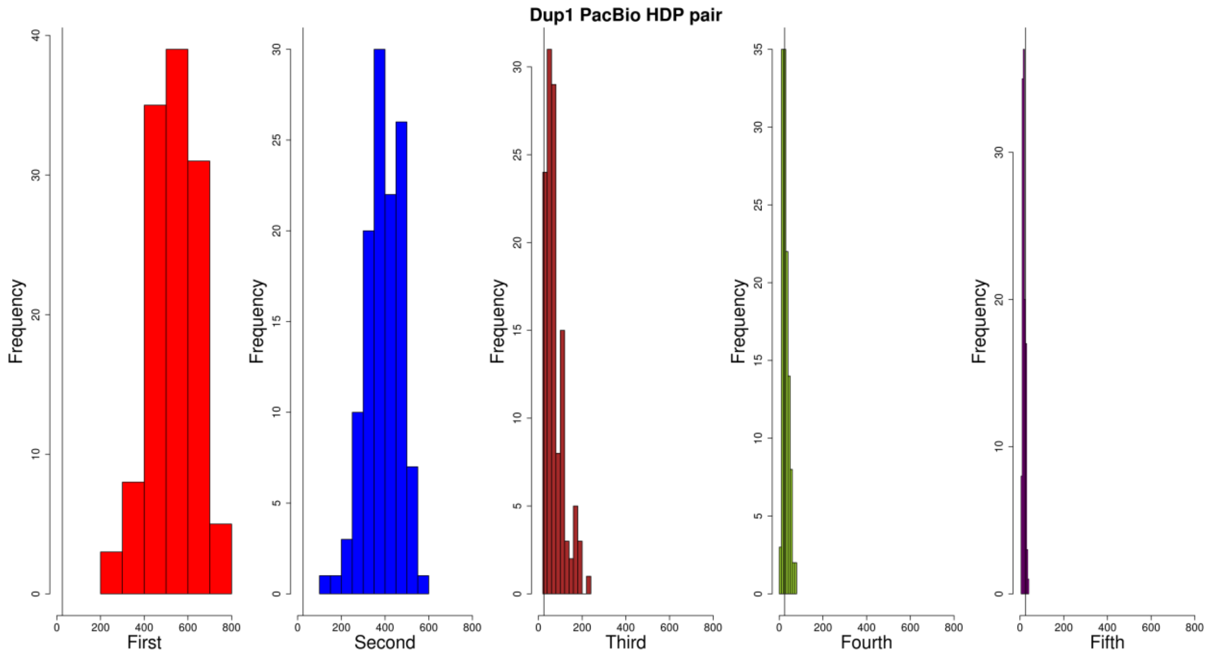


**Supplementary Figure. S89:** Histogram of the read pair count between HDPs for the Dup1 region, based on CAU Silkie PacBio long-reads mapped to the GRCg6a genome assembly. The ratio for the first and second pairs is much higher than for the remaining three pairs, based on a defined threshold of 0.25. This suggests that the majority of HDPs have two alleles.

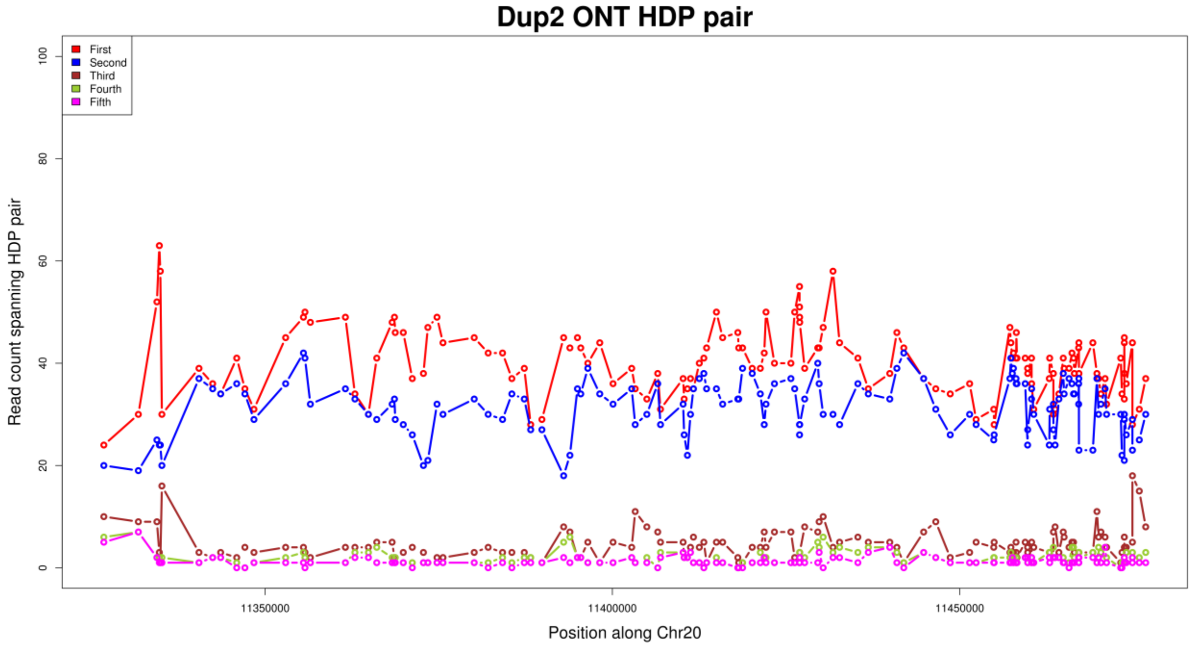


**Supplementary Figure. S90:** Read pair count between two HDPs for the Dup2 region, based on CAU Silkie ONT long-reads mapped to the GRCg6a genome assembly. The read counts for the first and second pairs are much higher than for the remaining three pairs, suggesting that the majority of HDPs have two alleles. Some HDPs show an increased count for the third pair, which is attributed to within-haplotype polymorphisms.

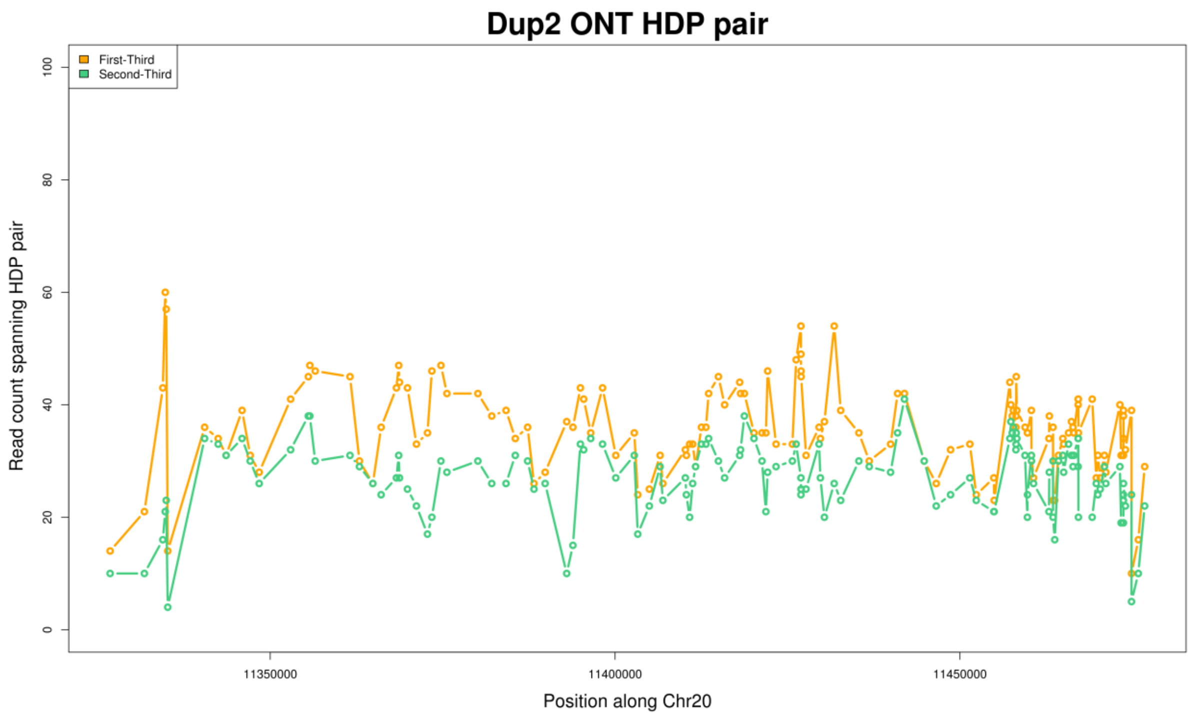


**Supplementary Figure. S91:** Read pair count for the first and second pairs between two HDPs for the Dup2 region after subtracting the count of the third pair. The ratio for the first and second pairs is still comparable.

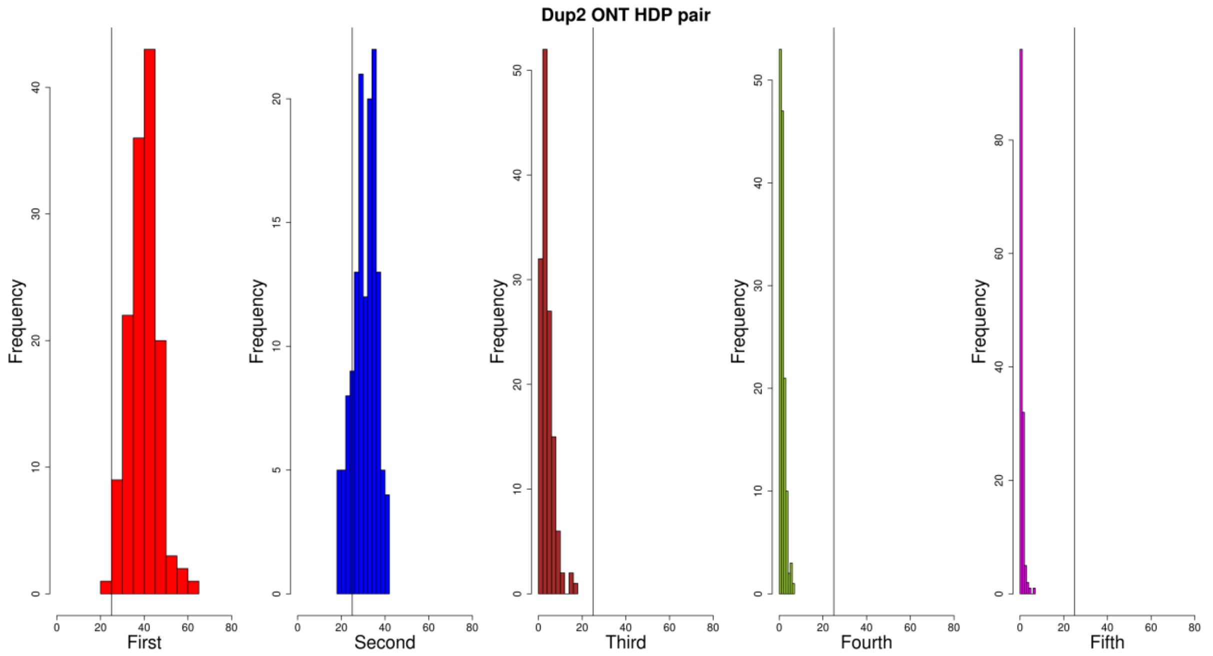


**Supplementary Figure. S92:** Histogram of the read pair count between HDPs for the Dup2 region, based on CAU Silkie ONT long-reads mapped to the GRCg6a genome assembly. The ratio for the first and second pairs is much higher than for the remaining three pairs, based on a defined threshold of 0.25. This suggests that the majority of HDPs have two alleles.

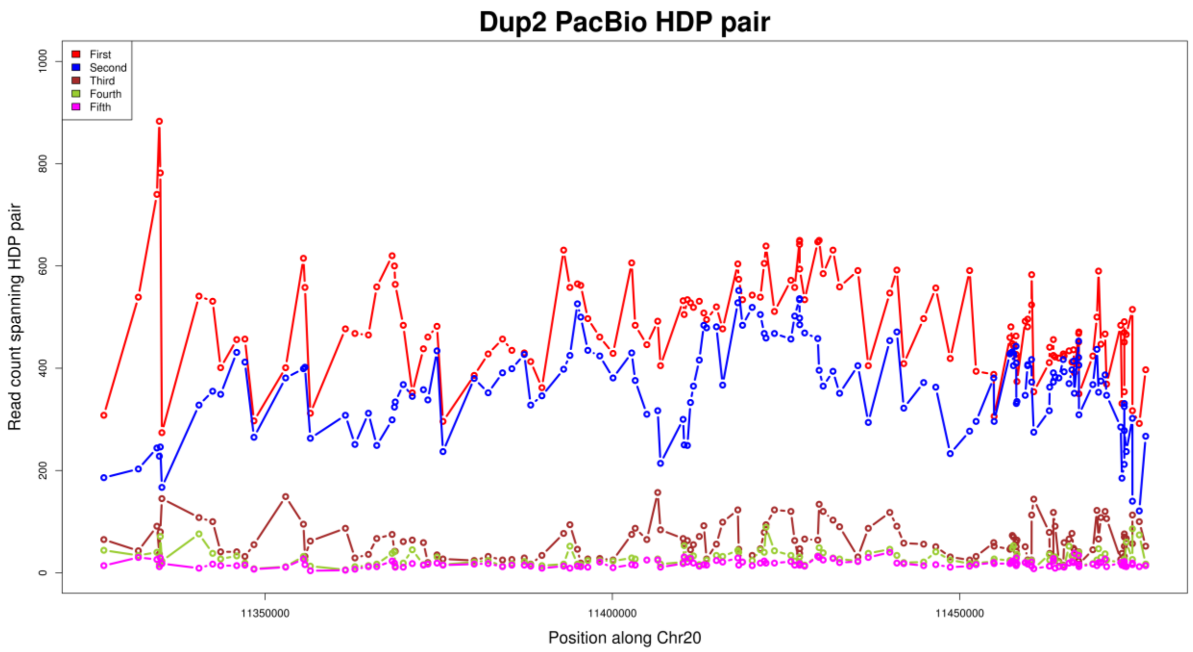


**Supplementary Figure. S93:** Read pair count between two HDPs for the Dup2 region, based on CAU Silkie PacBio long-reads mapped to the GRCg6a genome assembly. The read counts for the first and second pairs are much higher than for the remaining three pairs, suggesting that the majority of HDPs have two alleles. Some HDPs show an increased count for the third pair, which is attributed to within-haplotype polymorphisms.

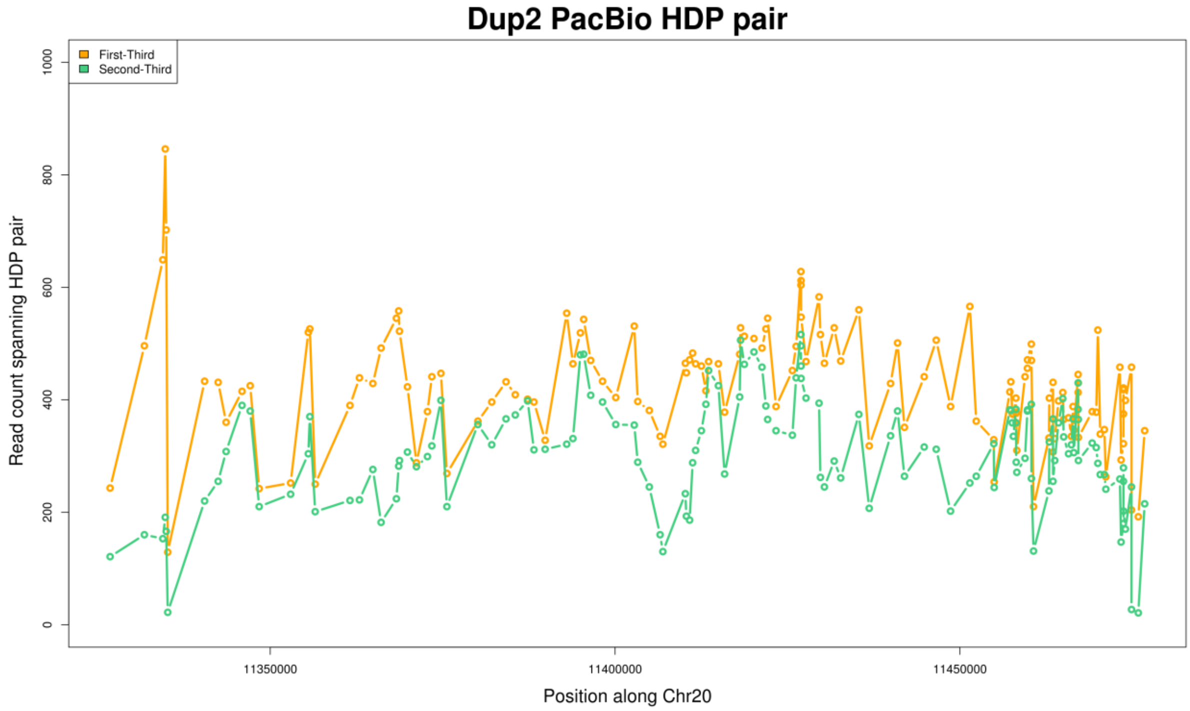


**Supplementary Figure. S94:** Read pair count for the first and second pairs between two HDPs for the Dup2 region after subtracting the count of the third pair. The ratio for the first and second pairs is still comparable.

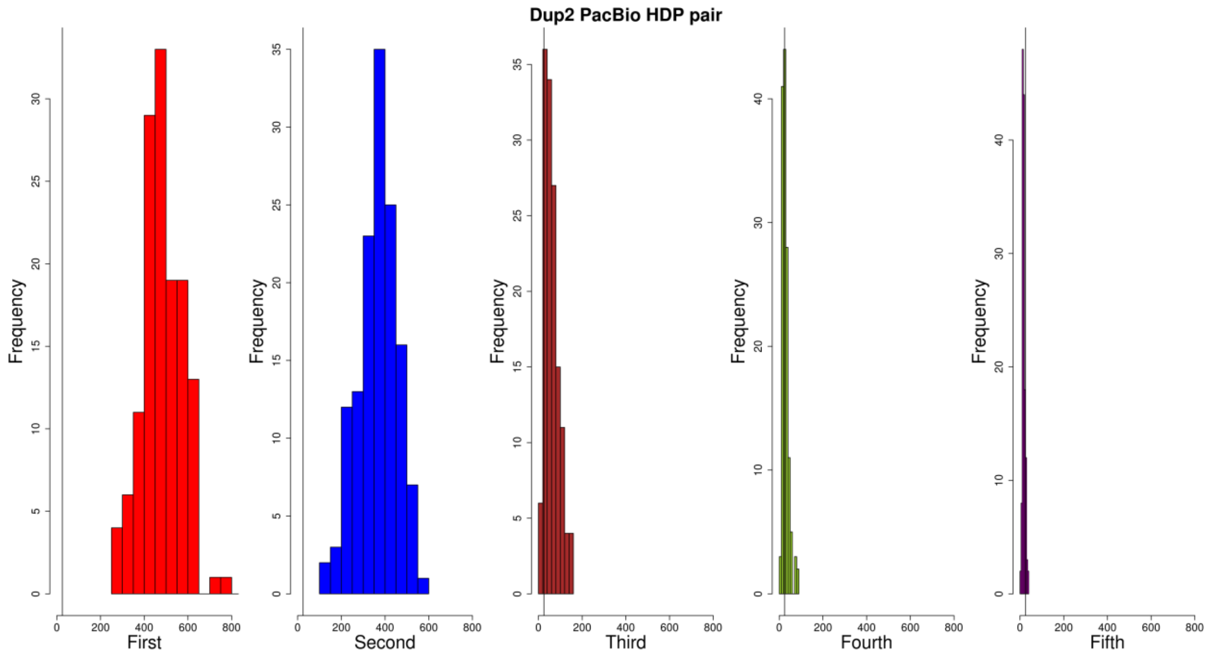


**Supplementary Figure. S95:** Histogram of the read pair count between HDPs for the Dup2 region, based on CAU Silkie PacBio long-reads mapped to the GRCg6a genome assembly. The ratio for the first and second pairs is much higher than for the remaining three pairs, based on a defined threshold of 0.25. This suggests that the majority of HDPs have two alleles.

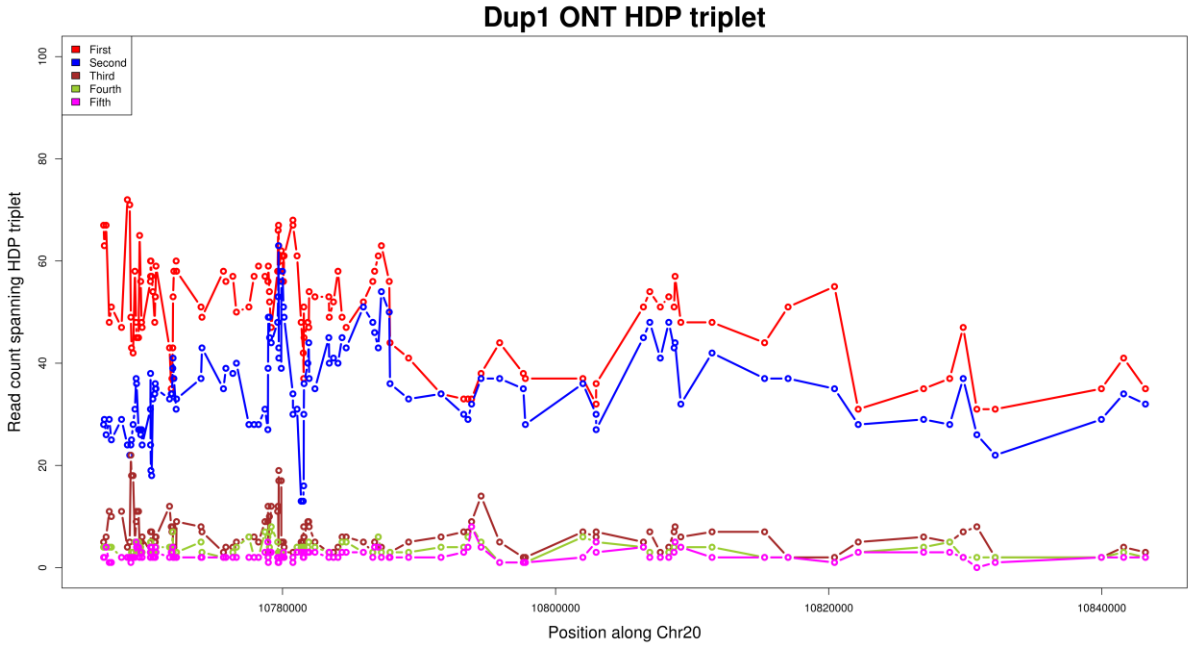


**Supplementary Figure. S96:** Read pair count between HDP triplet for the Dup1 region, based on CAU Silkie ONT long-reads mapped to the GRCg6a genome assembly. The read counts for the first and second pairs are much higher than for the remaining three pairs, suggesting that the majority of HDPs have two alleles. Some HDPs show an increased count for the third pair, which is attributed to within-haplotype polymorphisms.

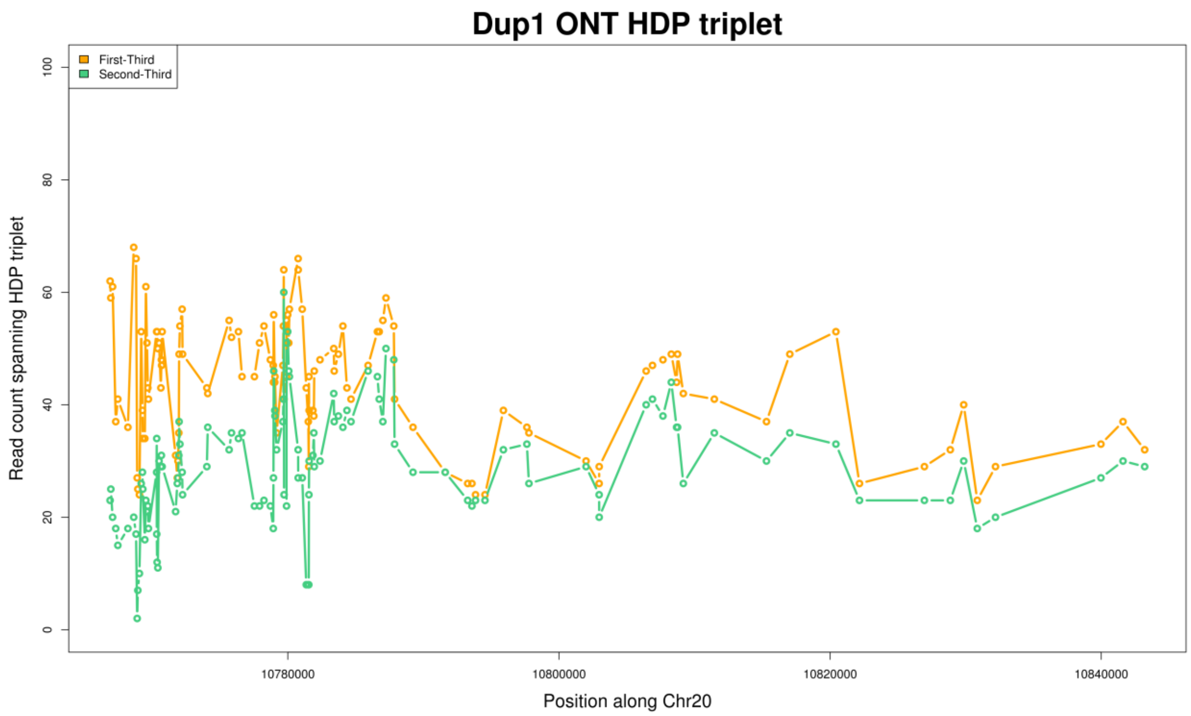


**Supplementary Figure. S97:** Read pair count for the first and second pairs between HDP triplet for the Dup1 region after subtracting the count of the third pair. The ratio for the first and second pairs is still comparable.

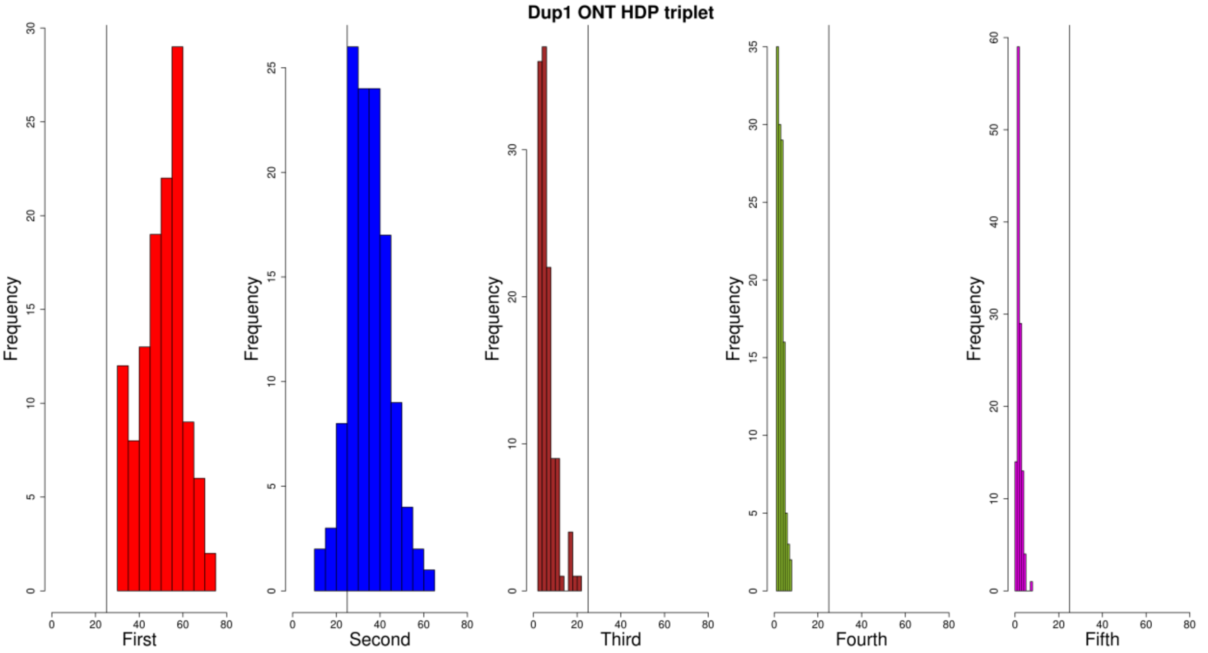


**Supplementary Figure. S98:** Histogram of the read pair count between HDP triplet for the Dup1 region, based on CAU Silkie ONT long-reads mapped to the GRCg6a genome assembly. The ratio for the first and second pairs is much higher than for the remaining three pairs, based on a defined threshold of 0.25. This suggests that the majority of HDPs have two alleles.

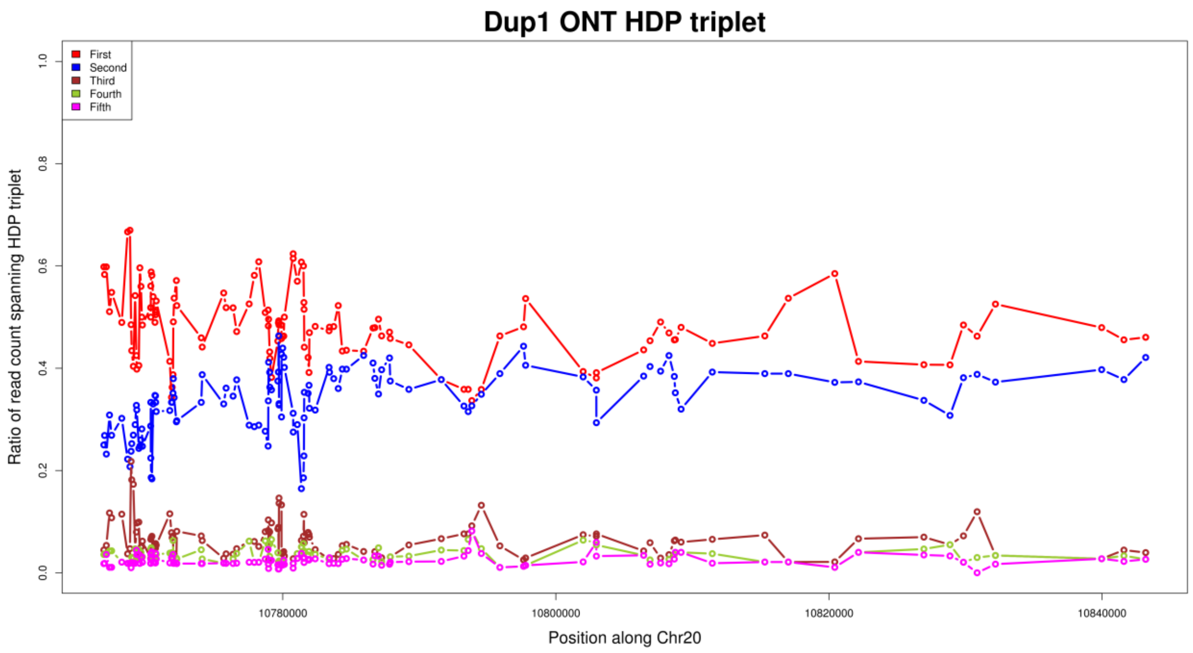


**Supplementary Figure. S99:** Ratio of read pair count between HDP triplet for the Dup1 region, based on CAU Silkie ONT long-reads mapped to the GRCg6a genome assembly. The read counts for the first and second pairs are much higher than for the remaining three pairs, suggesting that the majority of HDPs have two alleles. Some HDPs show an increased count for the third pair, which is attributed to within-haplotype polymorphisms.

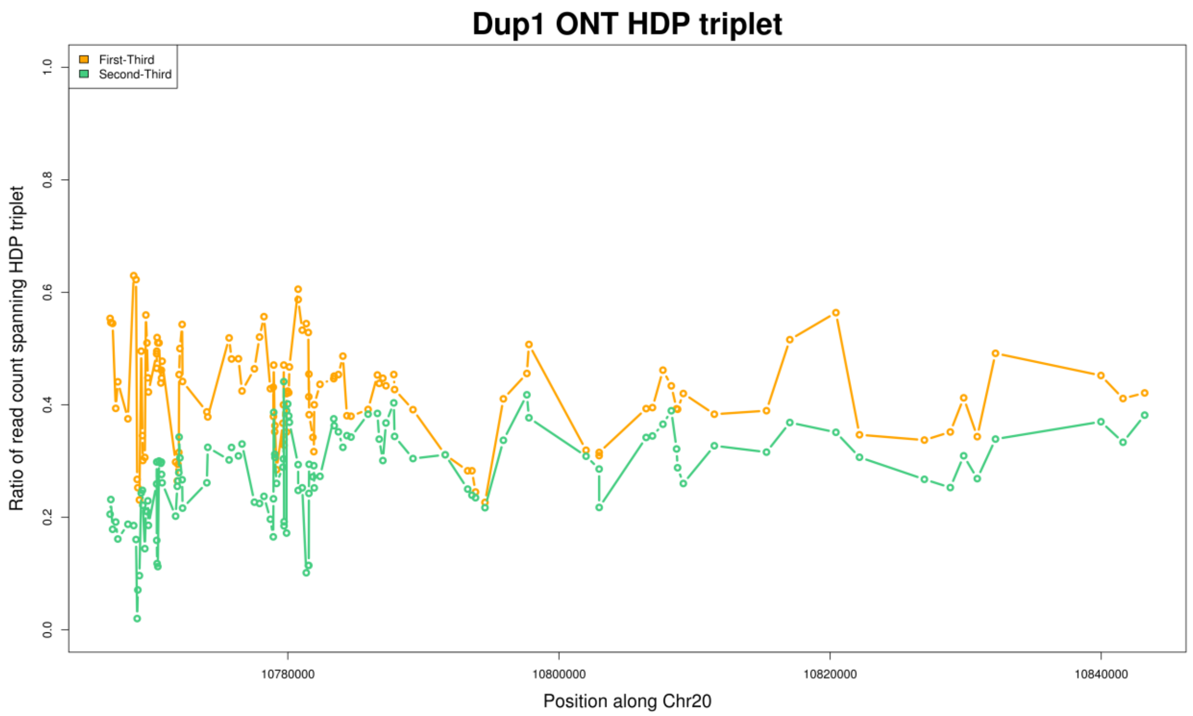


**Supplementary Figure. S100:** Ratio of read pair count for the first and second pairs between HDP triplet for the Dup1 region after subtracting the count of the third pair. The ratio for the first and second pairs is still comparable.

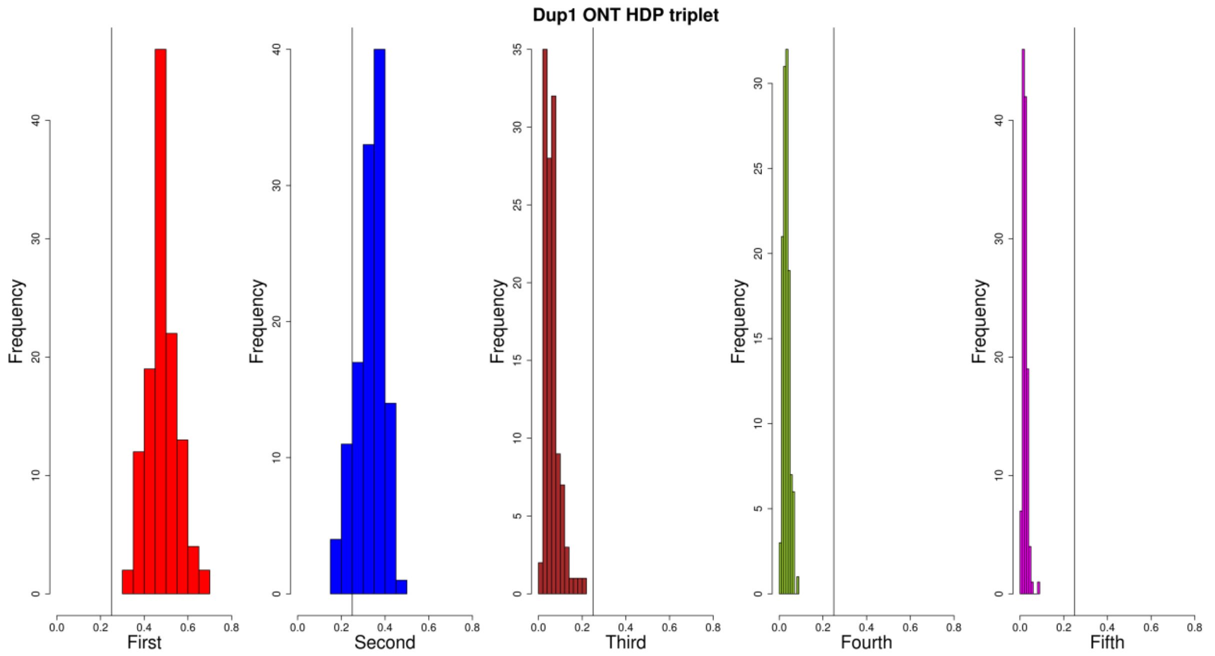


**Supplementary Figure. S101:** Histogram of the ratio of read pair count between HDP triplet for the Dup1 region, based on CAU Silkie ONT long-reads mapped to the GRCg6a genome assembly. The ratio for the first and second pairs is much higher than for the remaining three pairs, based on a defined threshold of 0.25. This suggests that the majority of HDPs have two alleles.

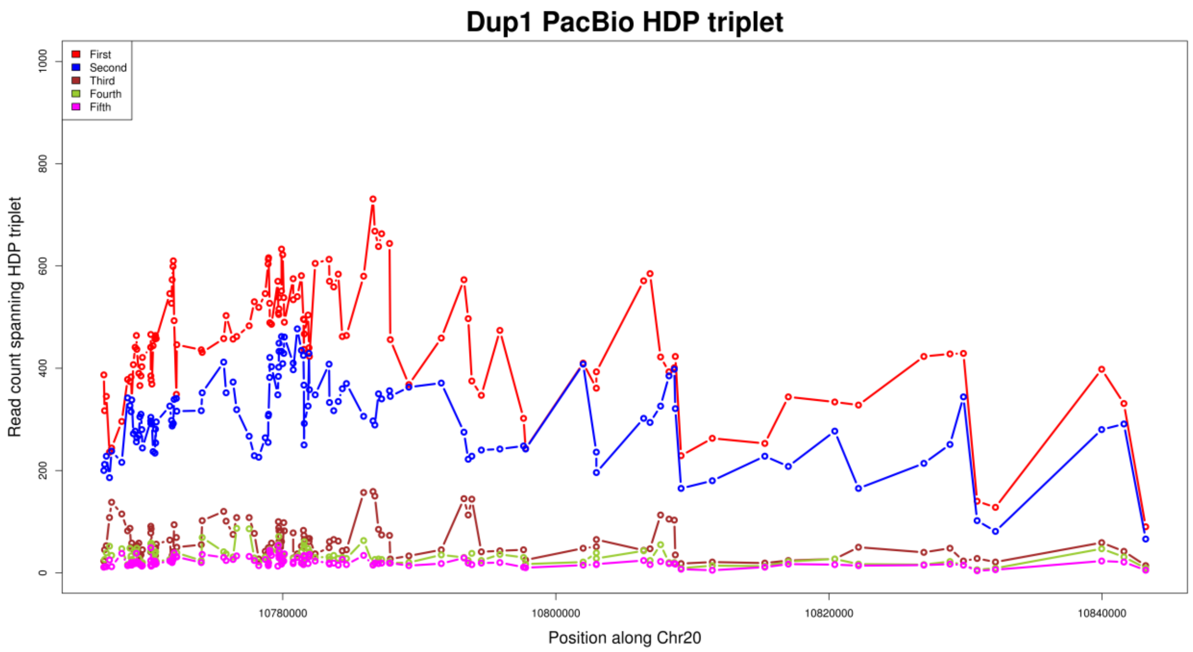


**Supplementary Figure. S102:** Read pair count between HDP triplet for the Dup1 region, based on CAU Silkie PacBio long-reads mapped to the GRCg6a genome assembly. The read counts for the first and second pairs are much higher than for the remaining three pairs, suggesting that the majority of HDPs have two alleles. Some HDPs show an increased count for the third pair, which is attributed to within-haplotype polymorphisms.

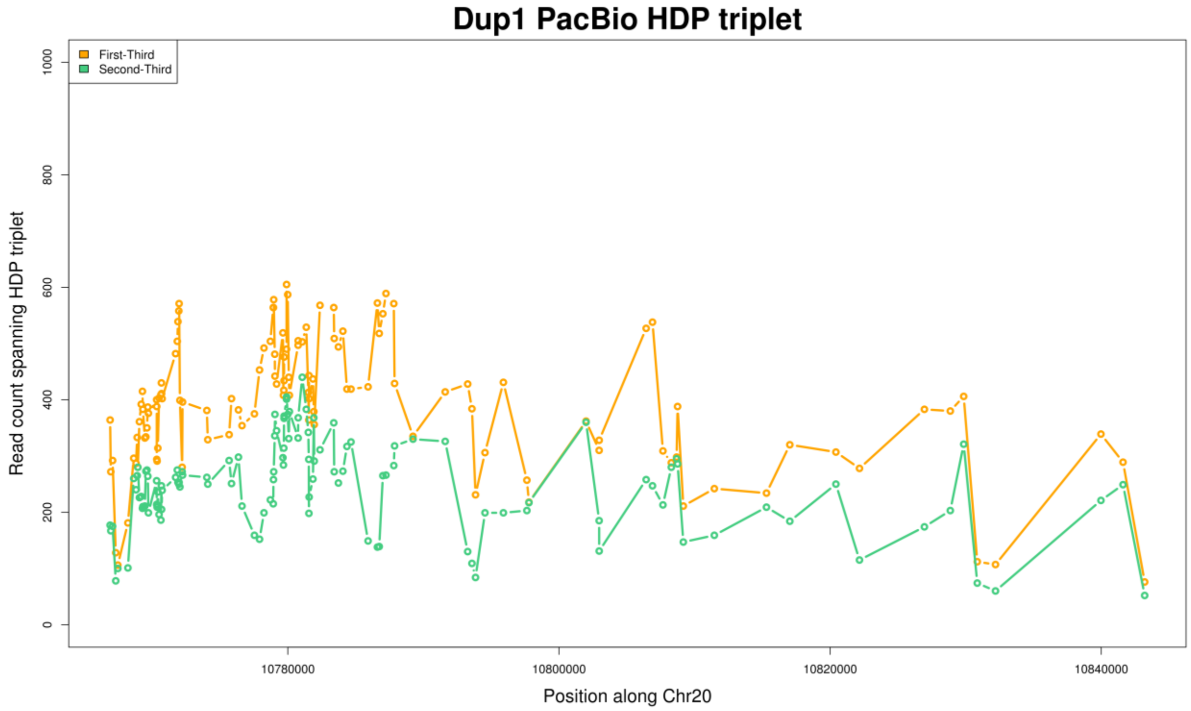


**Supplementary Figure. S103:** Read pair count for the first and second pairs between HDP triplet for the Dup1 region after subtracting the count of the third pair. The ratio for the first and second pairs is still comparable.

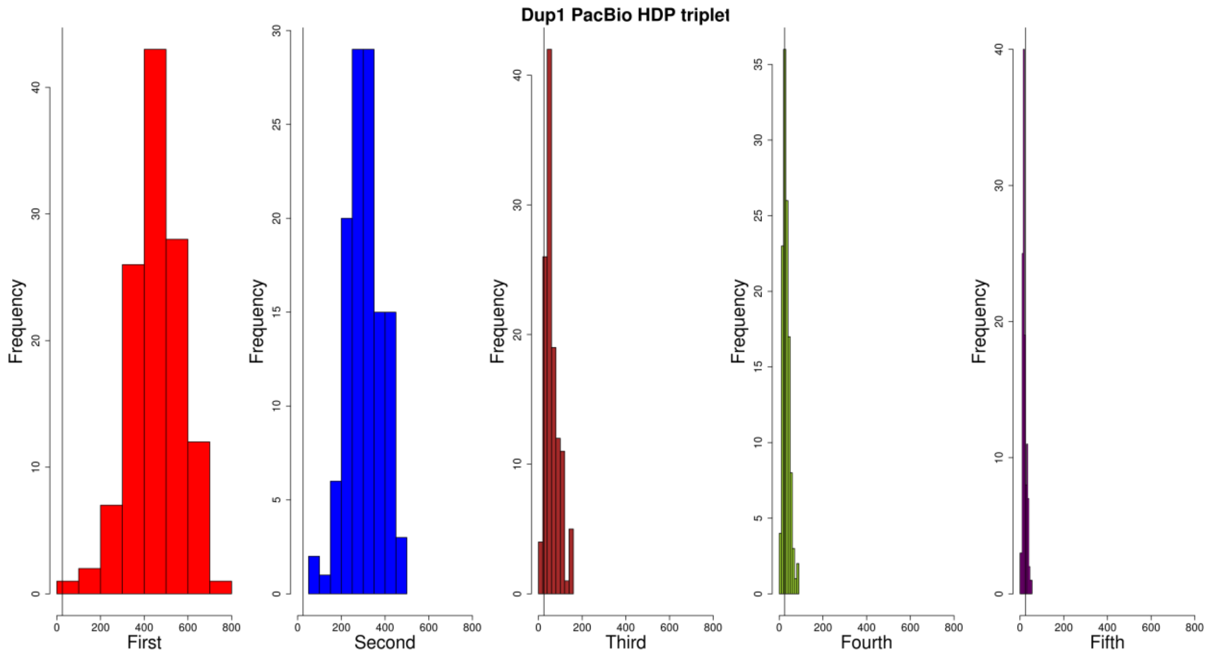


**Supplementary Figure. S104:** Histogram of read pair count between HDP triplet for the Dup1 region, based on CAU Silkie PacBio long-reads mapped to the GRCg6a genome assembly. The ratio for the first and second pairs is much higher than for the remaining three pairs, based on a defined threshold of 0.25. This suggests that the majority of HDPs have two alleles.

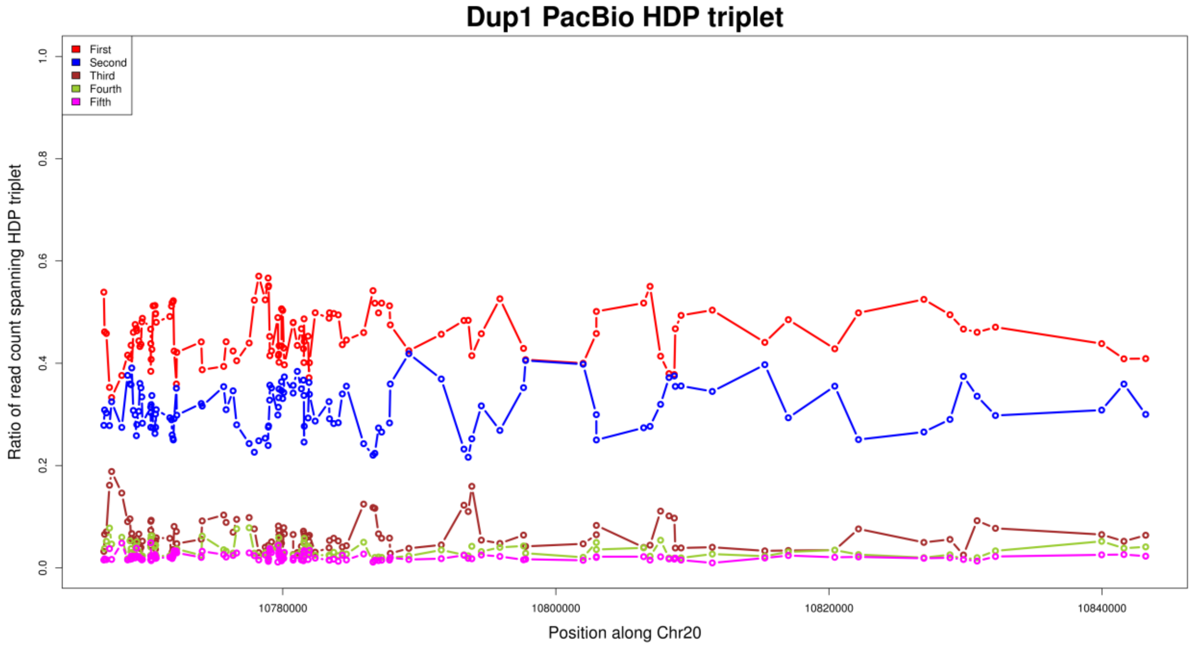


**Supplementary Figure. S105:** Ratio of read pair count between HDP triplet for the Dup1 region, based on CAU Silkie PacBio long-reads mapped to the GRCg6a genome assembly. The read counts for the first and second pairs are much higher than for the remaining three pairs, suggesting that the majority of HDPs have two alleles. Some HDPs show an increased count for the third pair, which is attributed to within-haplotype polymorphisms.

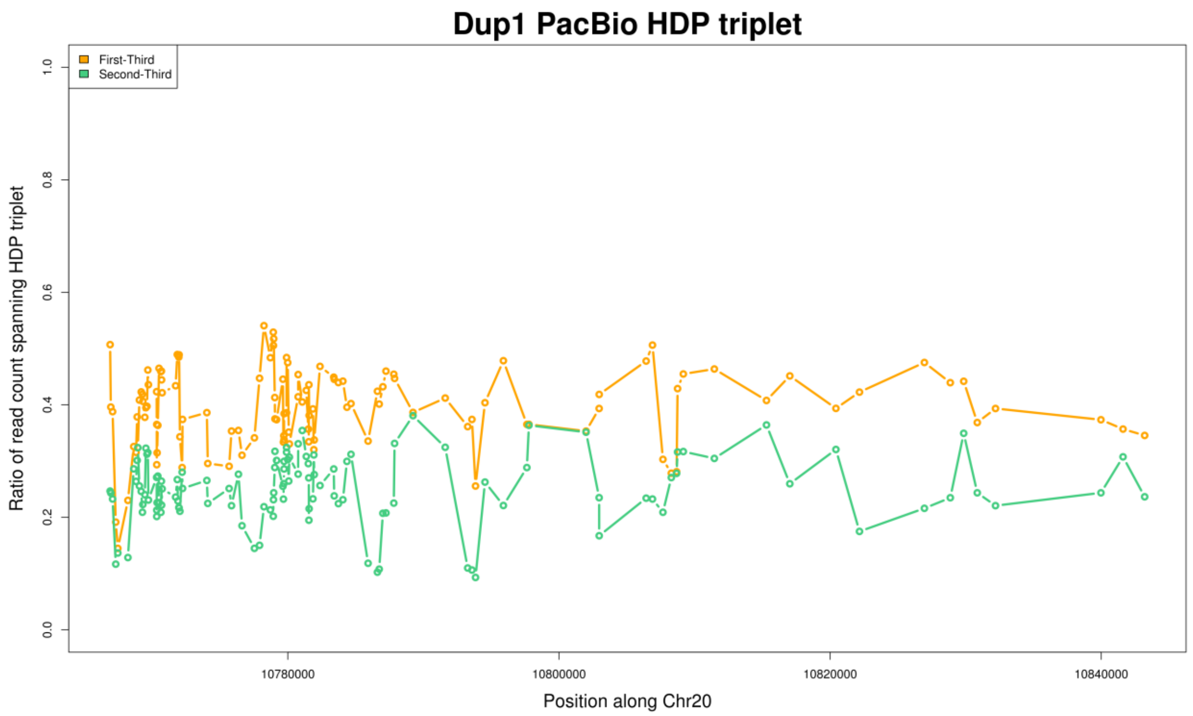


**Supplementary Figure. S106:** Ratio of read pair count for the first and second pairs between HDP triplet for the Dup1 region after subtracting the count of the third pair. The ratio for the first and second pairs is still comparable.

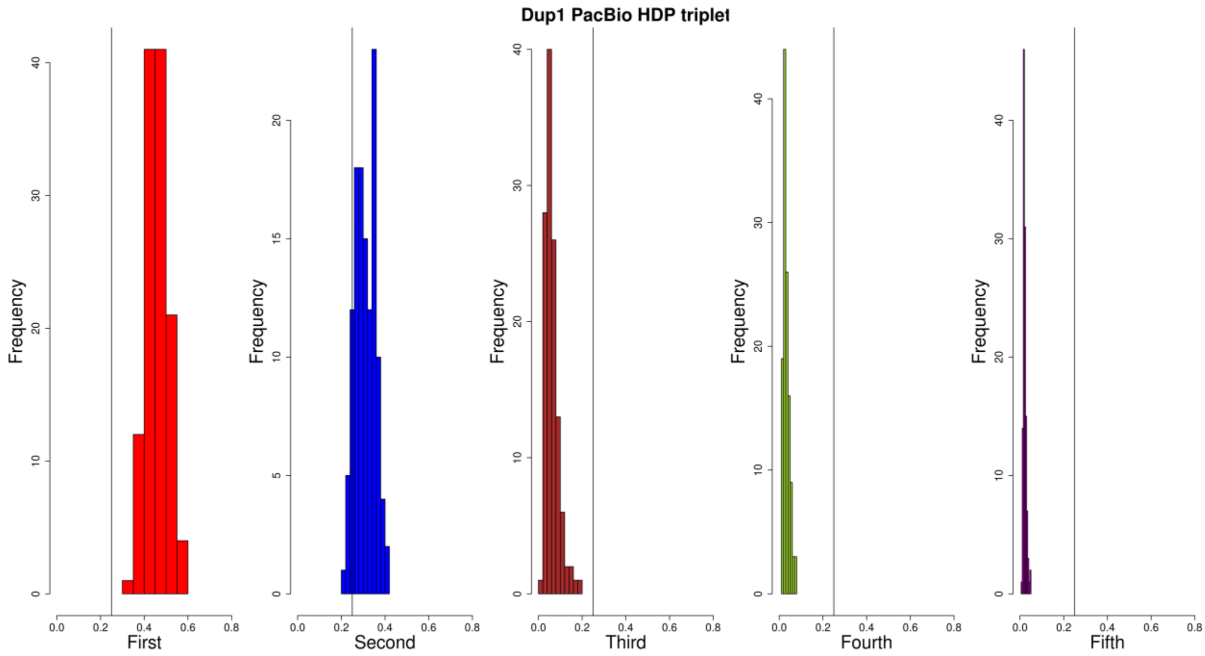


**Supplementary Figure. S107:** Histogram of the ratio of read pair count between HDP triplet for the Dup1 region, based on CAU Silkie PacBio long-reads mapped to the GRCg6a genome assembly. The ratio for the first and second pairs is much higher than for the remaining three pairs, based on a defined threshold of 0.25. This suggests that the majority of HDPs have two alleles.

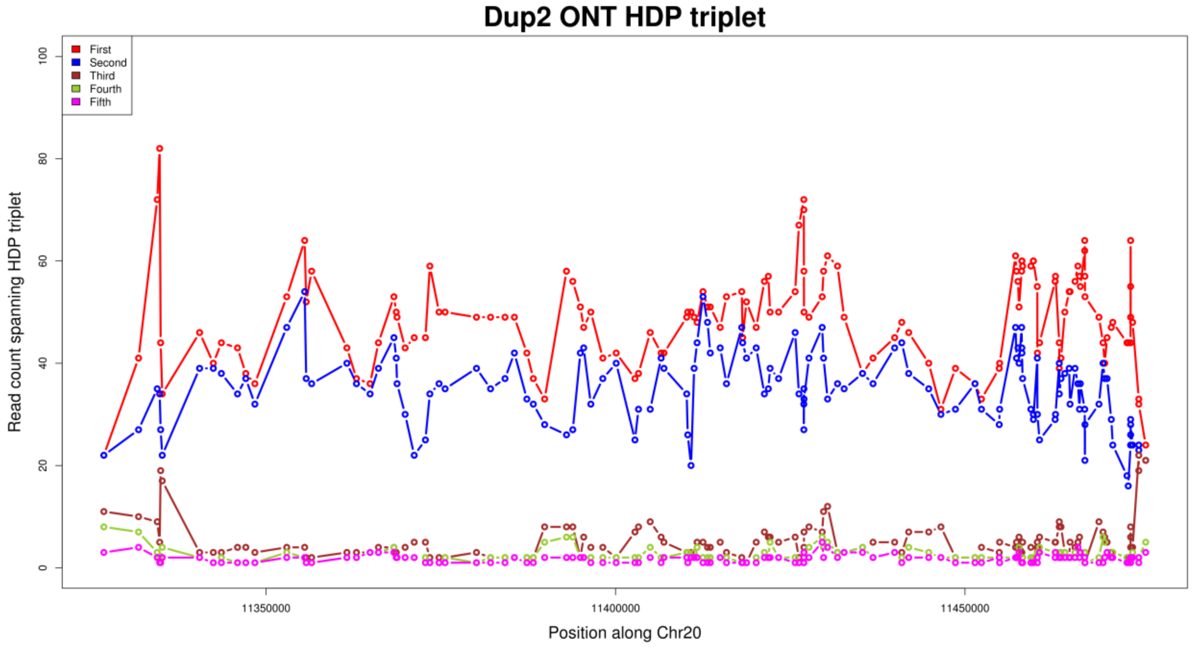


**Supplementary Figure. S108:** Read pair count between HDP triplet for the Dup2 region, based on CAU Silkie ONT long-reads mapped to the GRCg6a genome assembly. The read counts for the first and second pairs are much higher than for the remaining three pairs, suggesting that the majority of HDPs have two alleles. Some HDPs show an increased count for the third pair, which is attributed to within-haplotype polymorphisms.

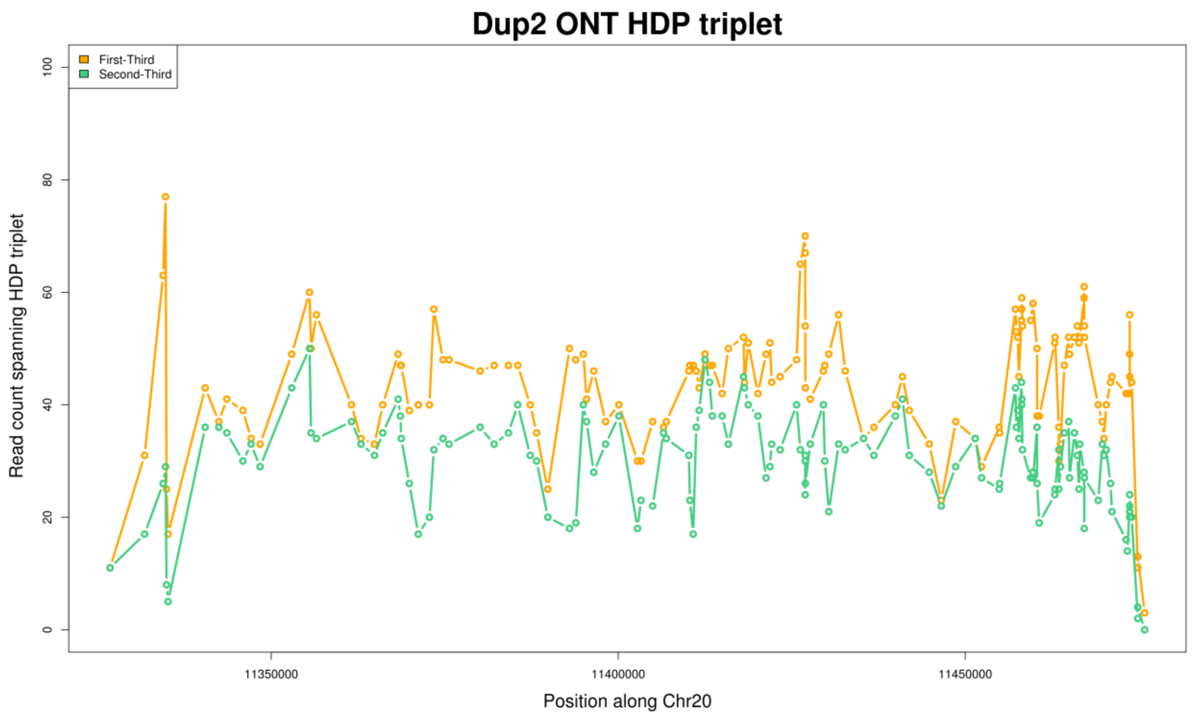


**Supplementary Figure. S109:** Read pair count for the first and second pairs between HDP triplet for the Dup2 region after subtracting the count of the third pair. The ratio for the first and second pairs is still comparable.

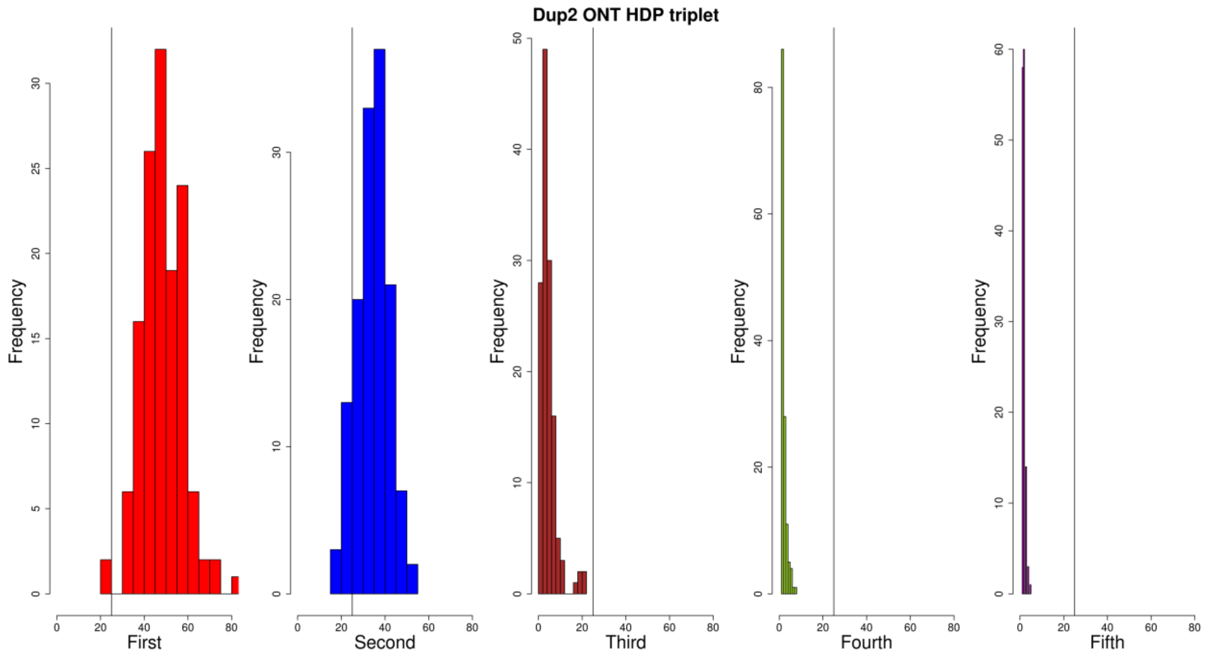


**Supplementary Figure. S110:** Histogram of the read pair count between HDP triplet for the Dup2 region, based on CAU Silkie ONT long-reads mapped to the GRCg6a genome assembly. The ratio for the first and second pairs is much higher than for the remaining three pairs, based on a defined threshold of 0.25. This suggests that the majority of HDPs have two alleles.

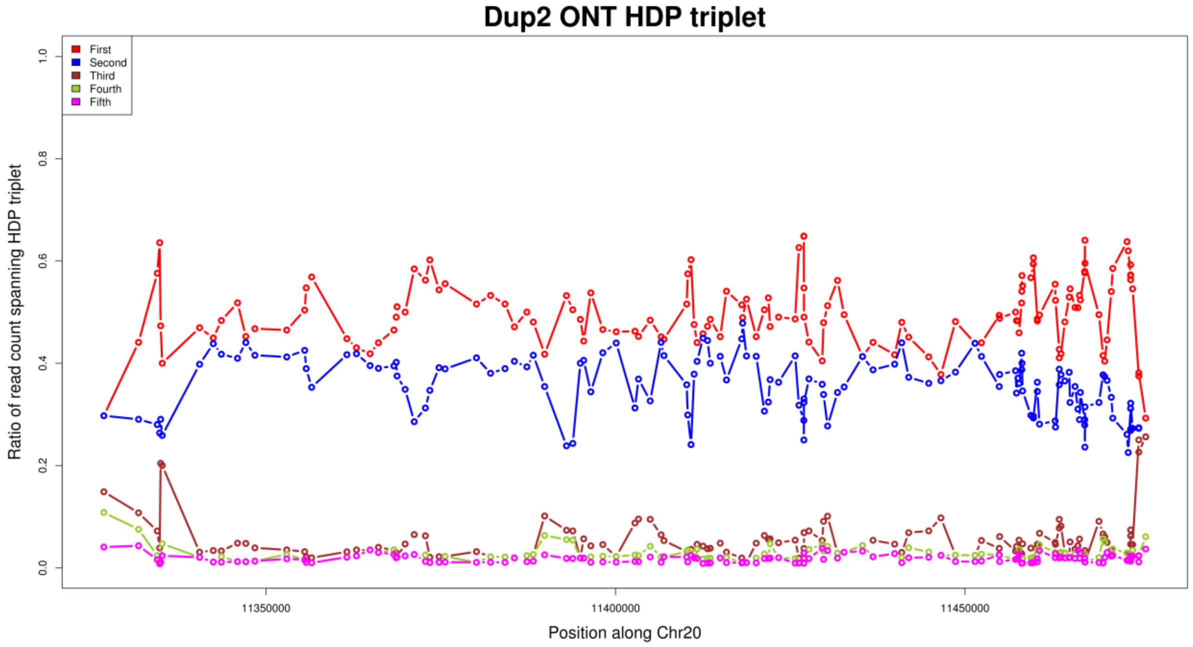


**Supplementary Figure. S111:** Ratio of read pair count between HDP triplet for the Dup2 region, based on CAU Silkie ONT long-reads mapped to the GRCg6a genome assembly. The read counts for the first and second pairs are much higher than for the remaining three pairs, suggesting that the majority of HDPs have two alleles. Some HDPs show an increased count for the third pair, which is attributed to within-haplotype polymorphisms.

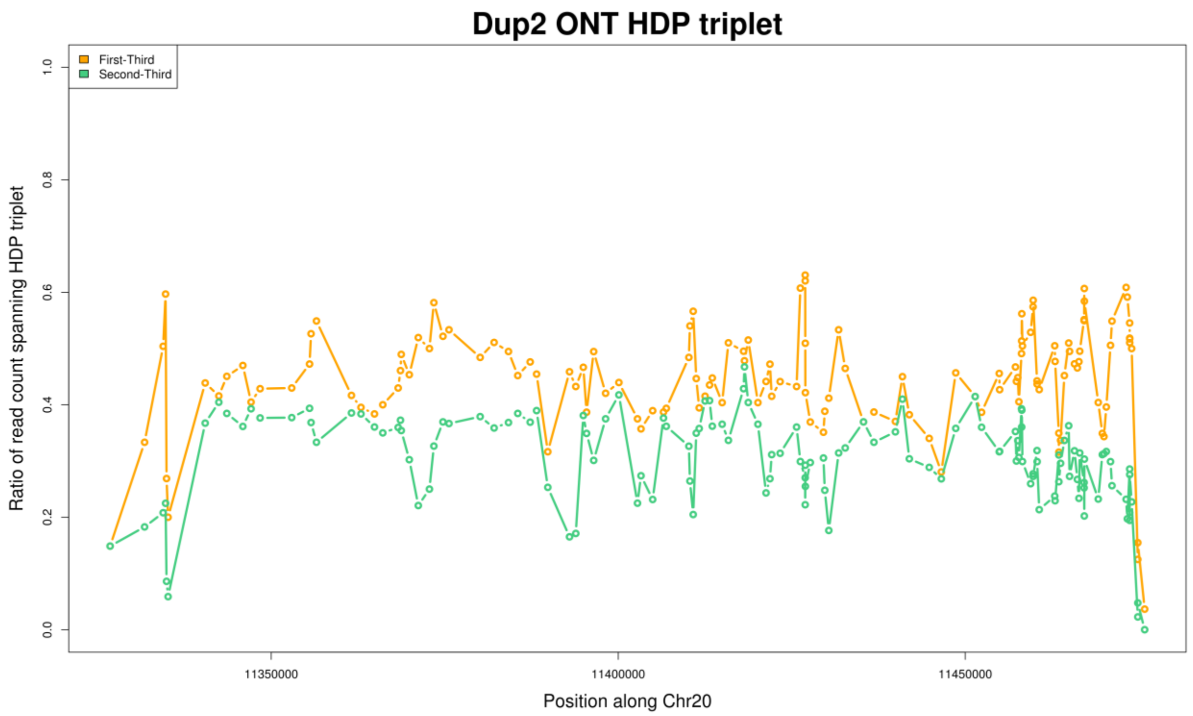


**Supplementary Figure. S112:** Ratio of read pair count for the first and second pairs between HDP triplet for the Dup2 region after subtracting the count of the third pair. The ratio for the first and second pairs is still comparable.

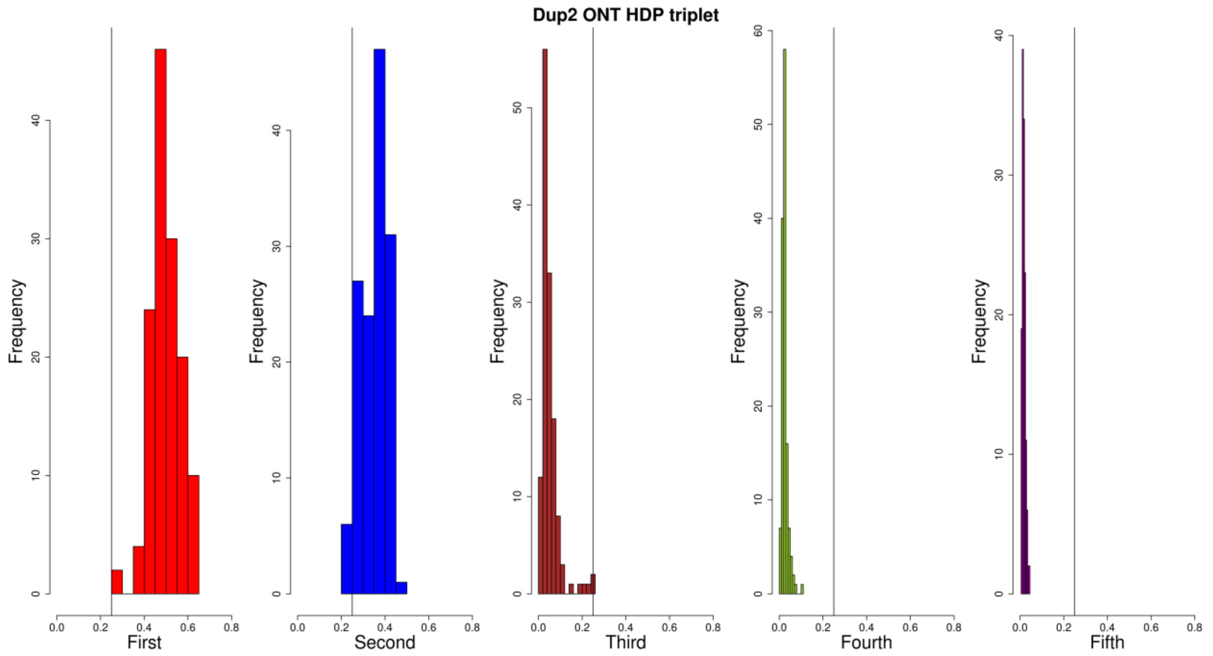


**Supplementary Figure. S113:** Histogram of the ratio of read pair count between HDP triplet for the Dup2 region, based on CAU Silkie ONT long-reads mapped to the GRCg6a genome assembly. The ratio for the first and second pairs is much higher than for the remaining three pairs, based on a defined threshold of 0.25. This suggests that the majority of HDPs have two alleles.

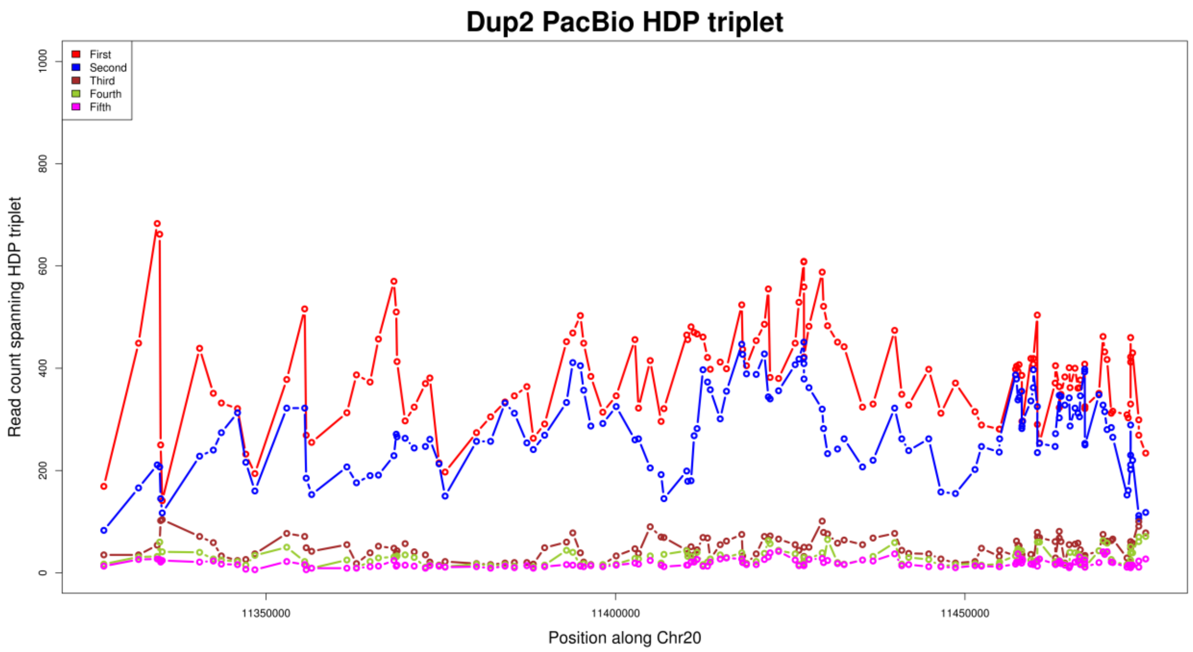


**Supplementary Figure. S114:** Read pair count between HDP triplet for the Dup2 region, based on CAU Silkie PacBio long-reads mapped to the GRCg6a genome assembly. The read counts for the first and second pairs are much higher than for the remaining three pairs, suggesting that the majority of HDPs have two alleles. Some HDPs show an increased count for the third pair, which is attributed to within-haplotype polymorphisms.

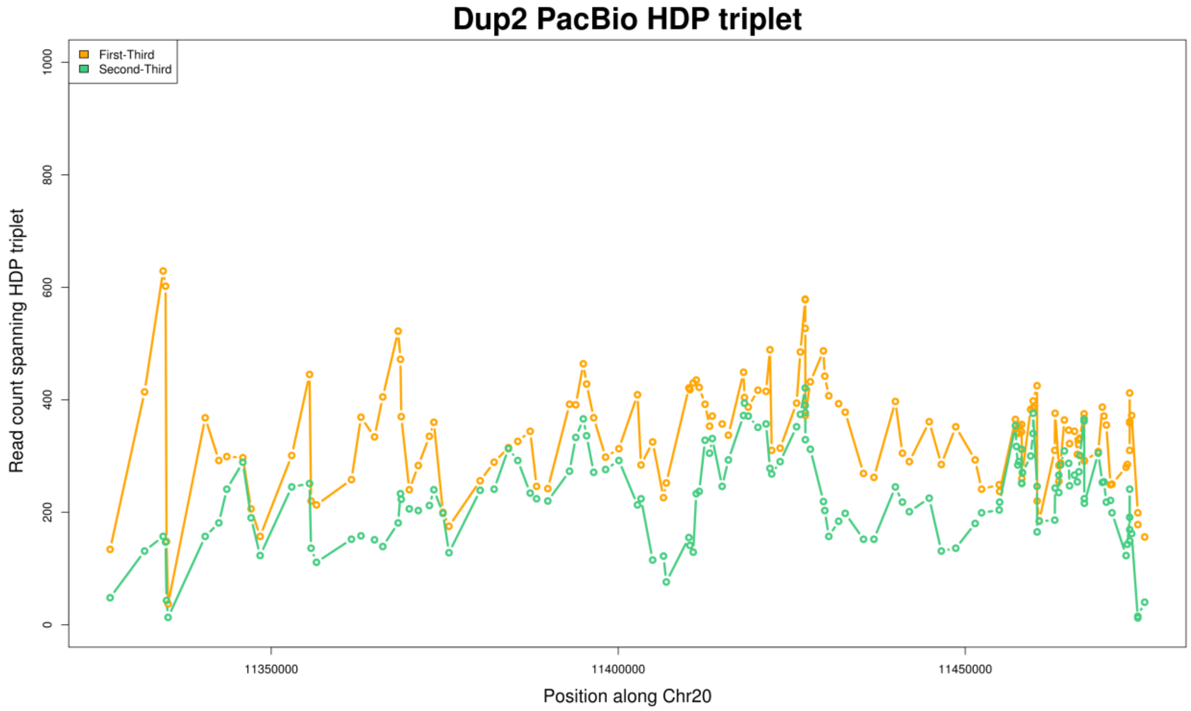


**Supplementary Figure. S115:** Read pair count for the first and second pairs between HDP triplet for the Dup2 region after subtracting the count of the third pair. The ratio for the first and second pairs is still comparable.

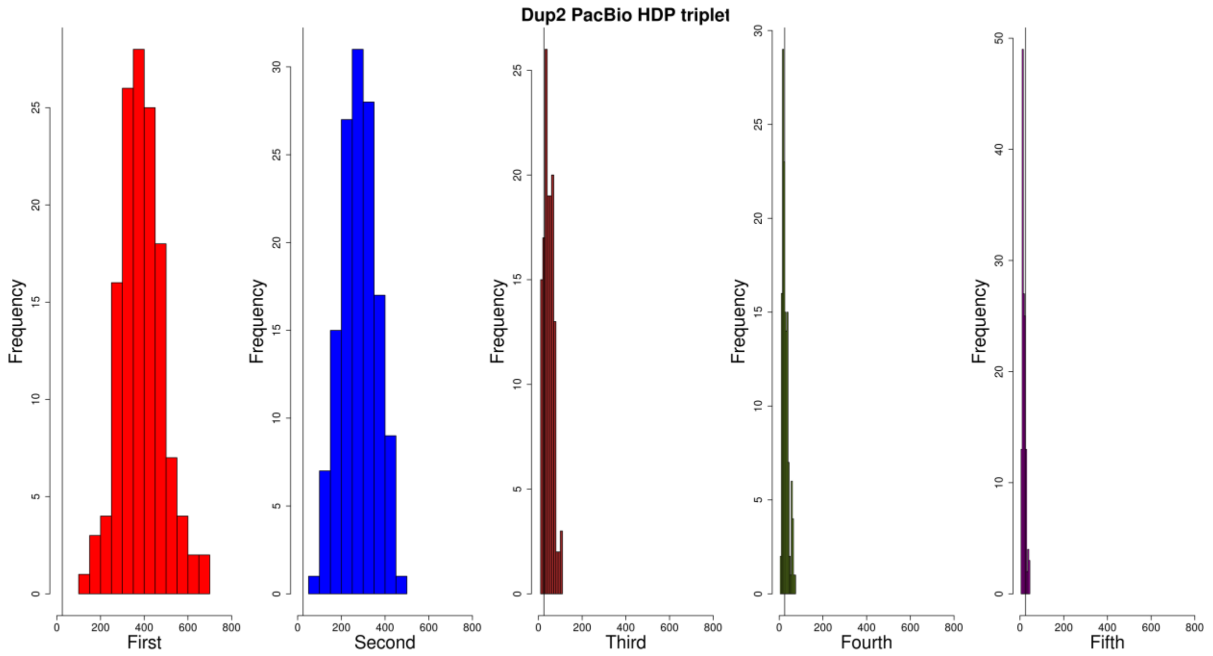


**Supplementary Figure. S116:** Histogram of the read pair count between HDP triplet for the Dup2 region, based on CAU Silkie PacBio long-reads mapped to the GRCg6a genome assembly. The ratio for the first and second pairs is much higher than for the remaining three pairs, based on a defined threshold of 0.25. This suggests that the majority of HDPs have two alleles.

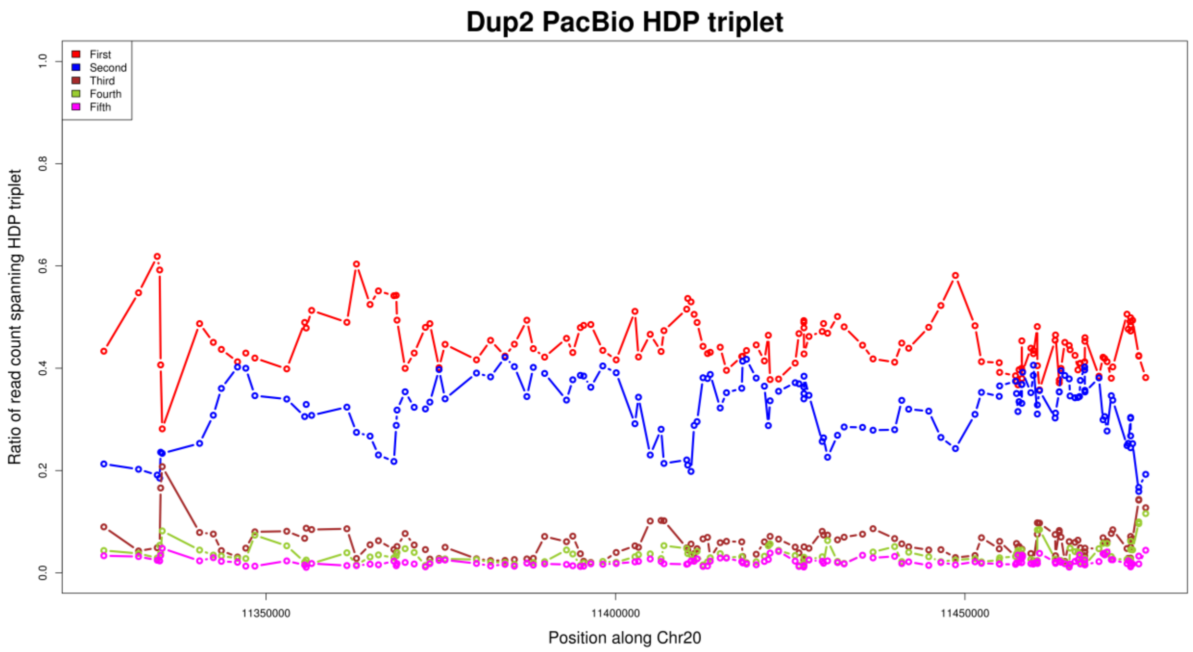


**Supplementary Figure. S117:** Ratio of read pair count between HDP triplet for the Dup2 region, based on CAU Silkie PacBio long-reads mapped to the GRCg6a genome assembly. The read counts for the first and second pairs are much higher than for the remaining three pairs, suggesting that the majority of HDPs have two alleles. Some HDPs show an increased count for the third pair, which is attributed to within-haplotype polymorphisms.

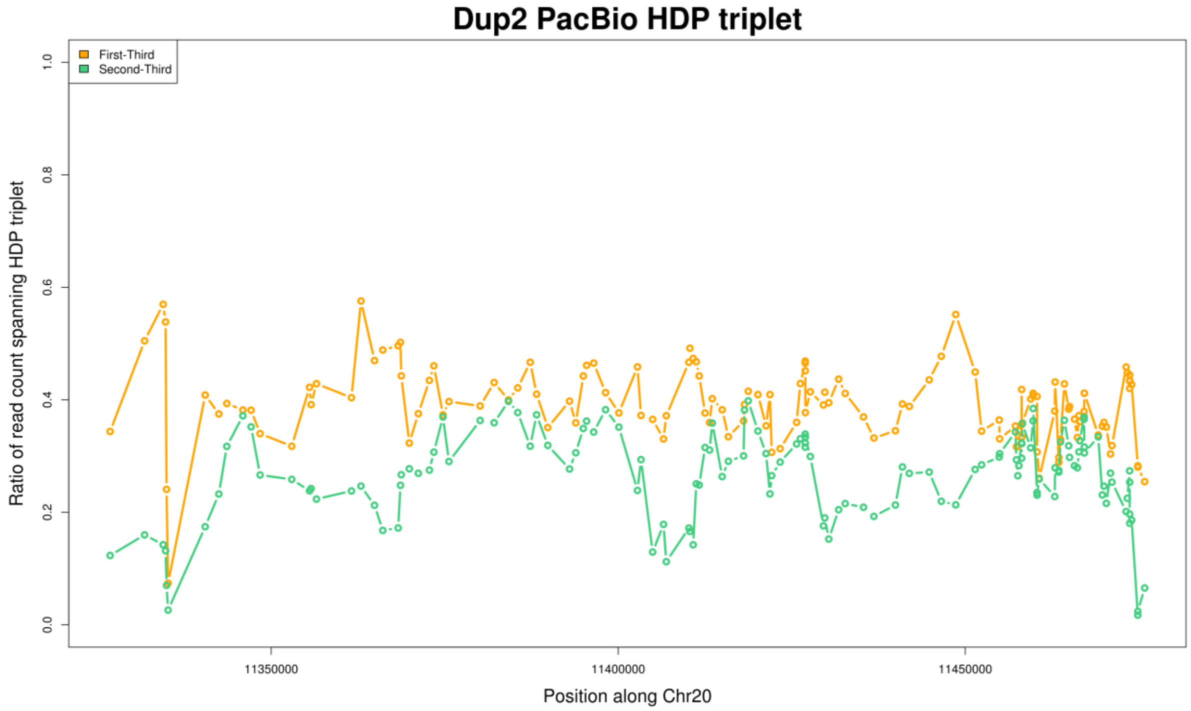


**Supplementary Figure. S118:** Ratio of read pair count for the first and second pairs between HDP triplet for the Dup2 region after subtracting the count of the third pair. The ratio for the first and second pairs is still comparable.

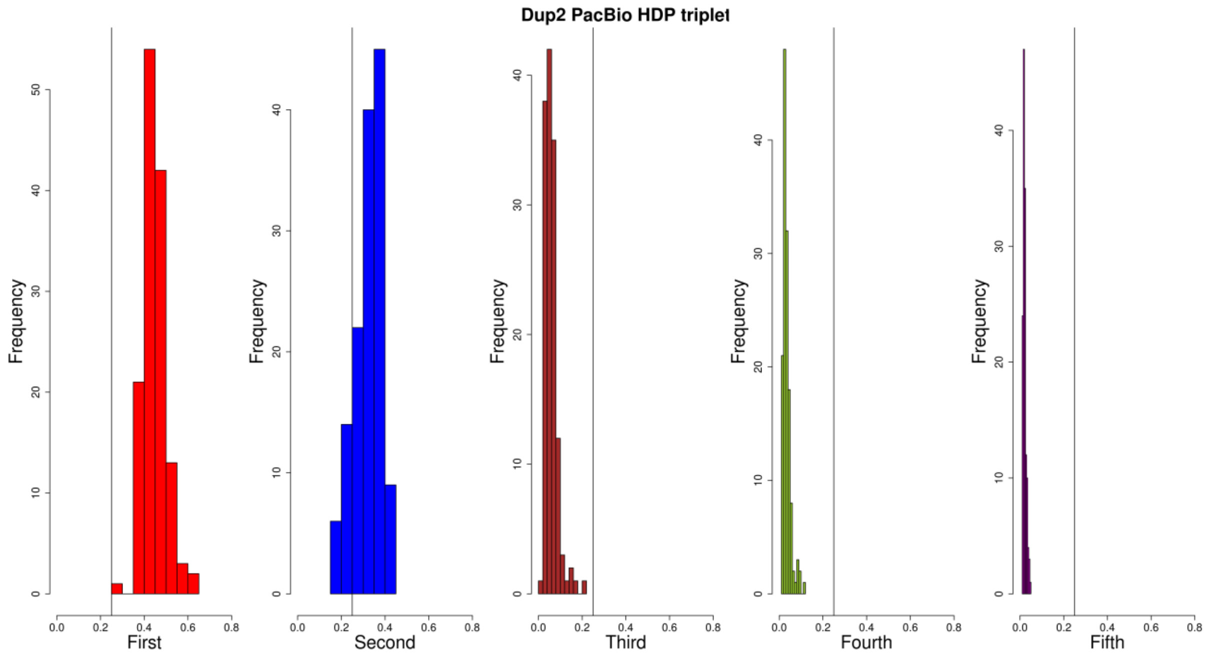


**Supplementary Figure. S119:** Histogram of the ratio of read pair count between HDP triplet for the Dup2 region, based on CAU Silkie PacBio long-reads mapped to the GRCg6a genome assembly. The ratio for the first and second pairs is much higher than for the remaining three pairs, based on a defined threshold of 0.25. This suggests that the majority of HDPs have two alleles.

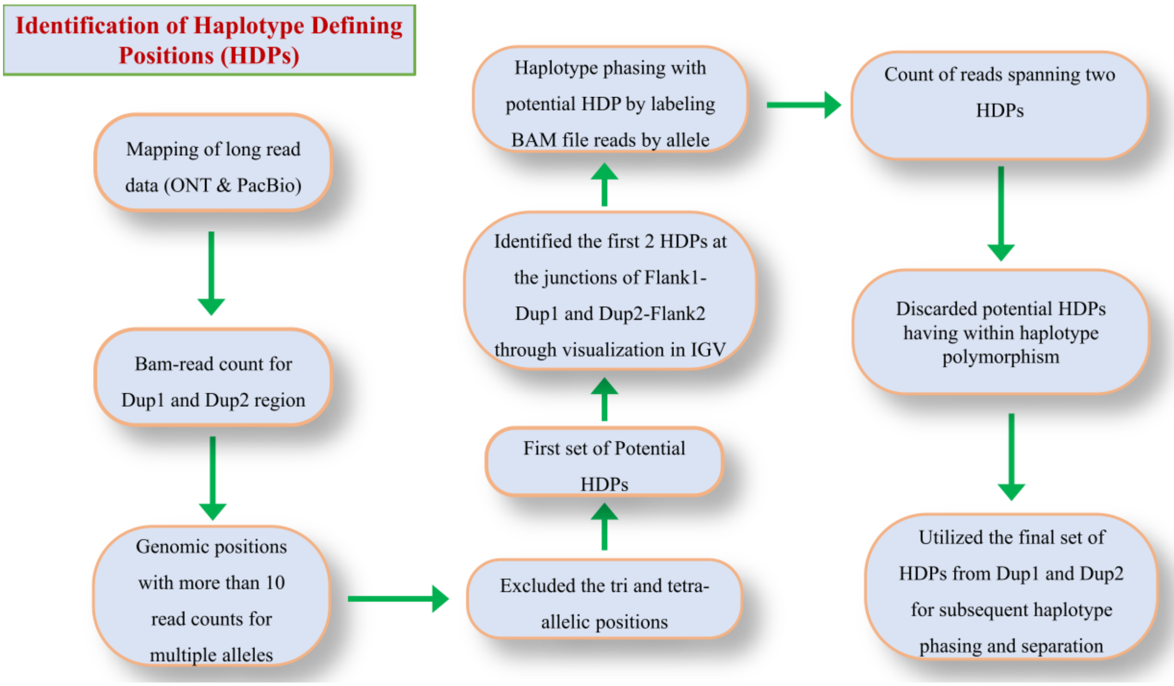


**Supplementary Figure. S120:** A comprehensive flowchart detailing the step-by-step methodology employed for the identification of Haplotype Defining Positions (HDPs) in ONT & PacBio long-reads of CAU Silkie mapped to GRCg6a genome assembly. The flowchart outlines the various stages of the process, from initial read mapping to final HDP identification, ensuring clarity and reproducibility of the methodology employed in this study.

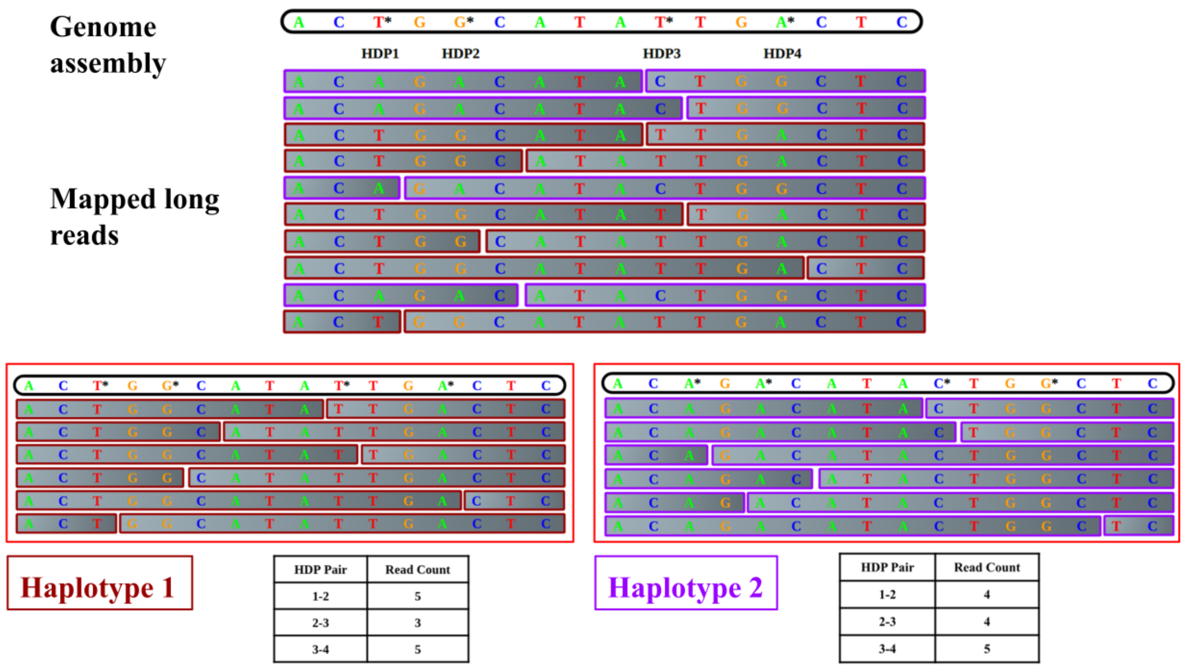


Haplotype 1

| HDP Pair | Read Count |
|---|---|
| 1-2 | 5 |
| 2-3 | 3 |
| 3-4 | 5 |

Haplotype 2

| HDP Pair | Read Count |
|---|---|
| 1-2 | 4 |
| 2-3 | 4 |
| 3-4 | 5 |

**Supplementary Figure. S121:** A graphical example of long-read-based haplotype phasing can be used to identify haplotypes. Long reads are mapped to the genome assembly. Genomic bases marked with an asterisk (*) represent Haplotype Defining Positions (HDPs). Based on these HDPs, Haplotype 1 and Haplotype 2 are separated. The read count for each HDP pair indicates the number of reads containing that specific HDP pair. Sequentially, we calculated the read count for each HDP pair across the region of interest.

**Alignment of CAU_Silkie long-reads to the CAU Silkie genome assembly at *Fm* locus**

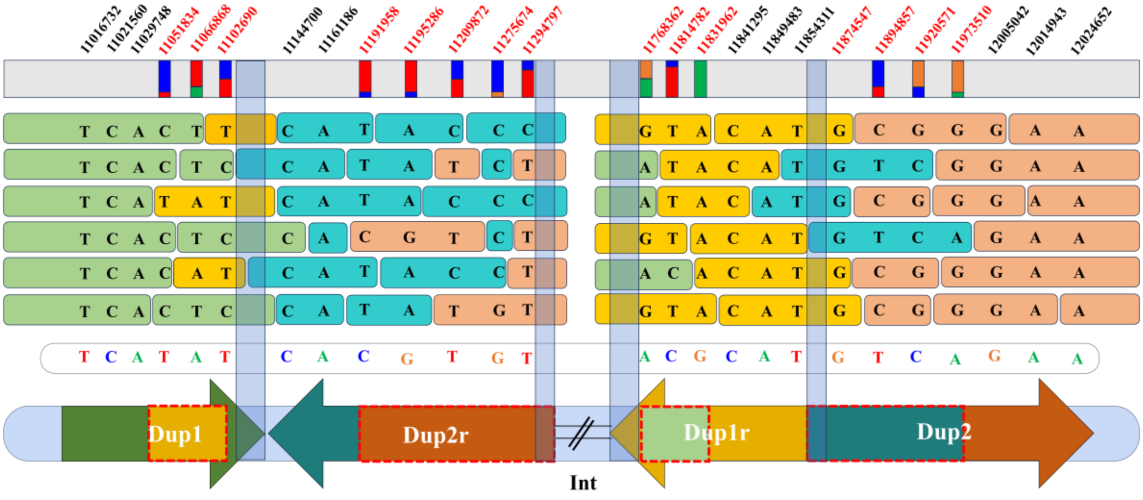


**Supplementary Figure. S122: Identification of mosaic assembly in the CAU Silkie genome:** This panel displays reads spanning the *Fm* locus. Reads are color-coded as follows: Dup1 (light green), Dup2r (teal), Dup1r (yellow), and Dup2 (light orange). Due to the mosaic assembly at the end of Dup1 and the start of Dup2r, as well as the end of Dup1r and the start of Dup2, reads from other haplotypes are misaligned to these regions highlighted in red dotted outlines. Nucleotide positions denote haplotype-defining positions (HDPs) within the Dup1 and Dup2 regions, which are critical for distinguishing the different haplotypes. HDPs in black represent the genomic regions where the CAU Silkie genome assembly is supported by haplotype-specific long-reads. HDPs highlighted in red indicate genomic coordinates where the CAU Silkie genome assembly is mosaic, leading to assembly errors. The light blue transparent rectangular box represents the HDP poor regions.

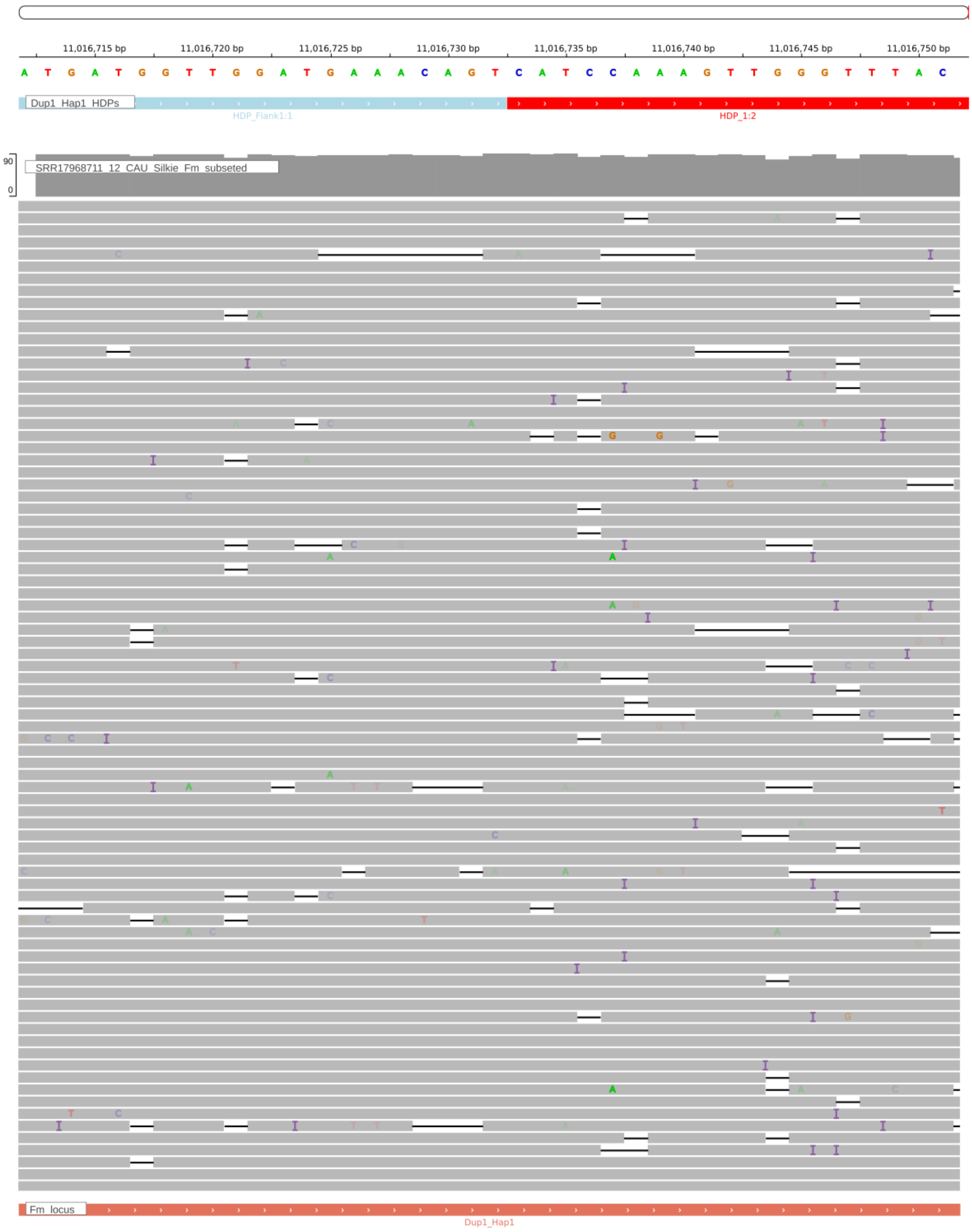


**Supplementary Figure S123:** The IGV screenshot displays Dup1 HDP 11016732 correctly assembled in the CAU Silkie genome, supported by raw reads from a single haplotype mapped to the genome.

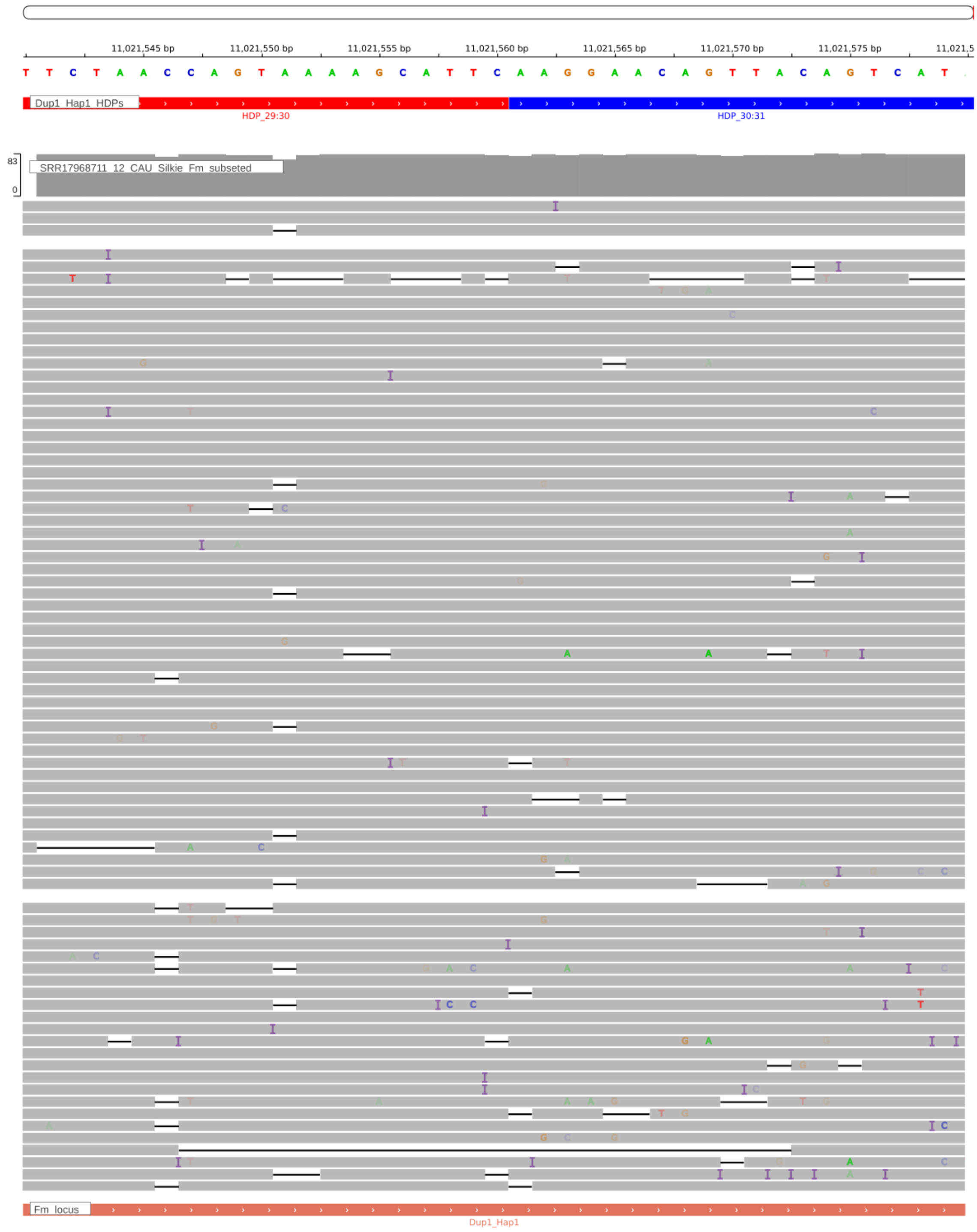


**Supplementary Figure. S124:** The IGV screenshot displays Dup1 HDP 11021560 correctly assembled in the CAU Silkie genome, supported by raw reads from a single haplotype mapped to the genome.

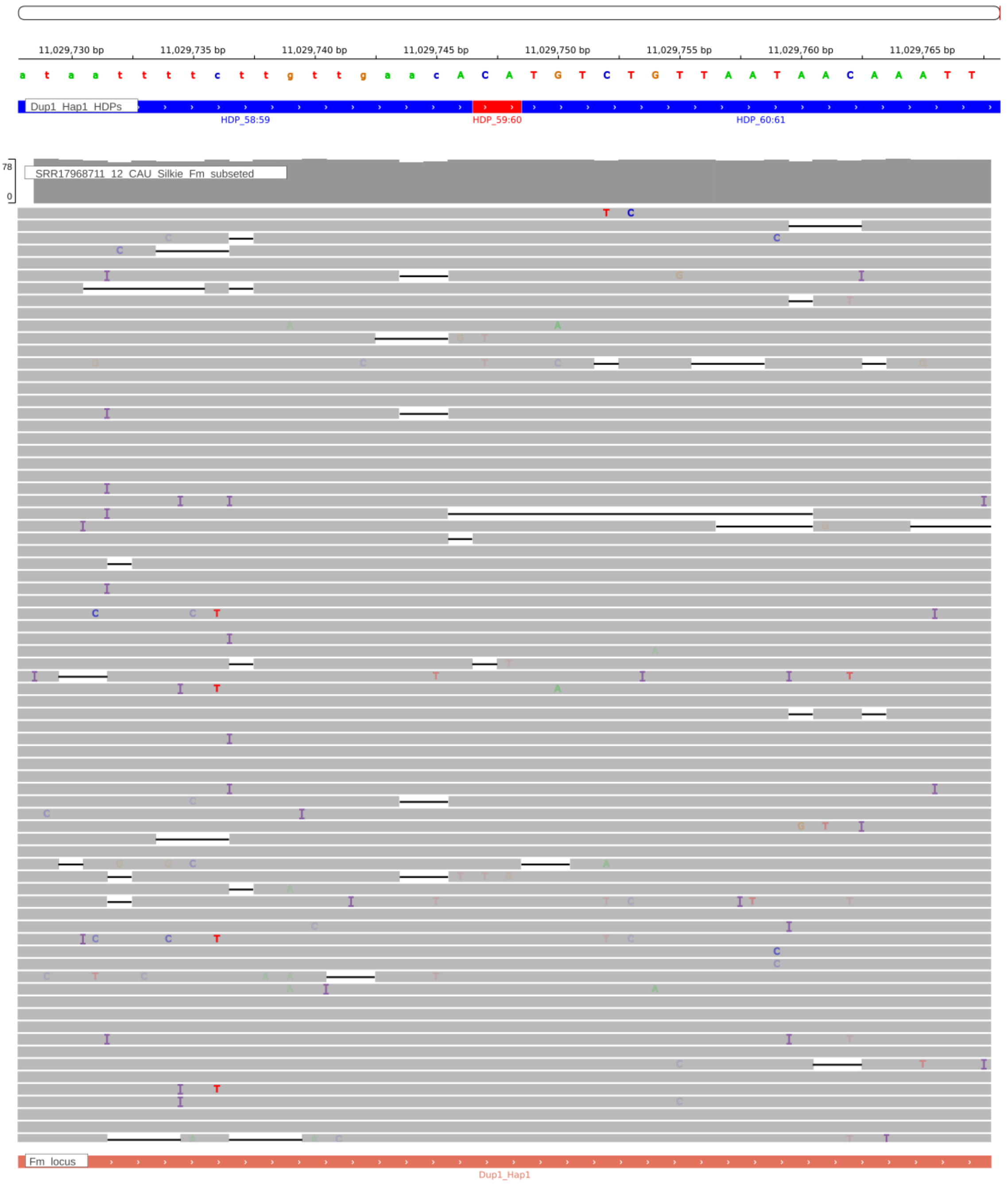


**Supplementary Figure. S124:** The IGV screenshot displays Dup1 HDP 11029748 correctly assembled in the CAU Silkie genome, supported by raw reads from a single haplotype mapped to the genome.

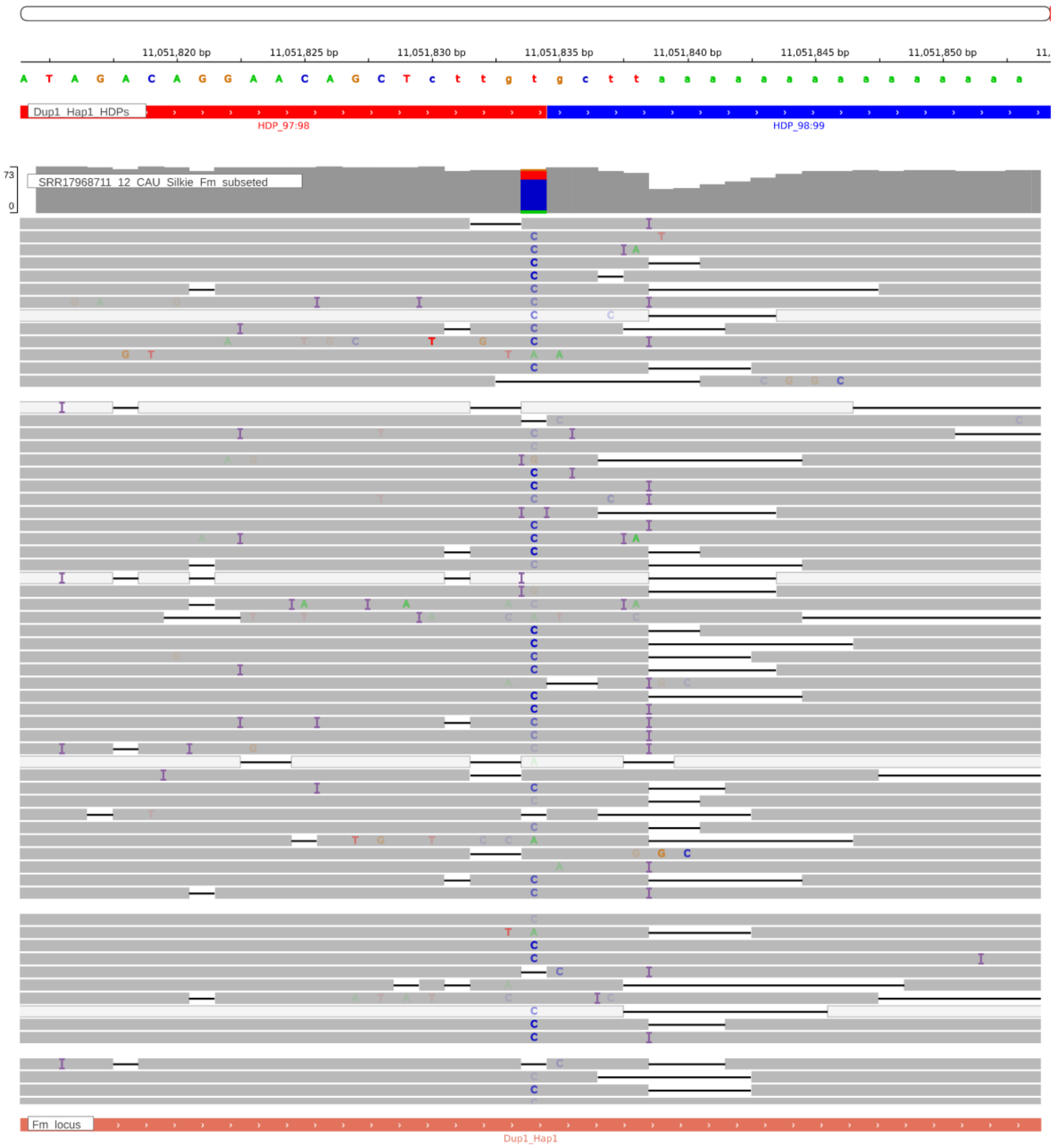


**Supplementary Figure. S126:** Screenshot from IGV showing the HDP 11051834 within the Dup1 region of the CAU Silkie genome. The CAU Silkie ONT long-reads were aligned to the CAU Silkie genome using minimap2, following the settings described by Zhu et al. The alignment reveals that support for the HDP within the genome is notably poor, indicating a mosaic genome assembly due to haplotype switching. The genomic base assembled at the given HDP position actually belongs to the Dup1r haplotype.

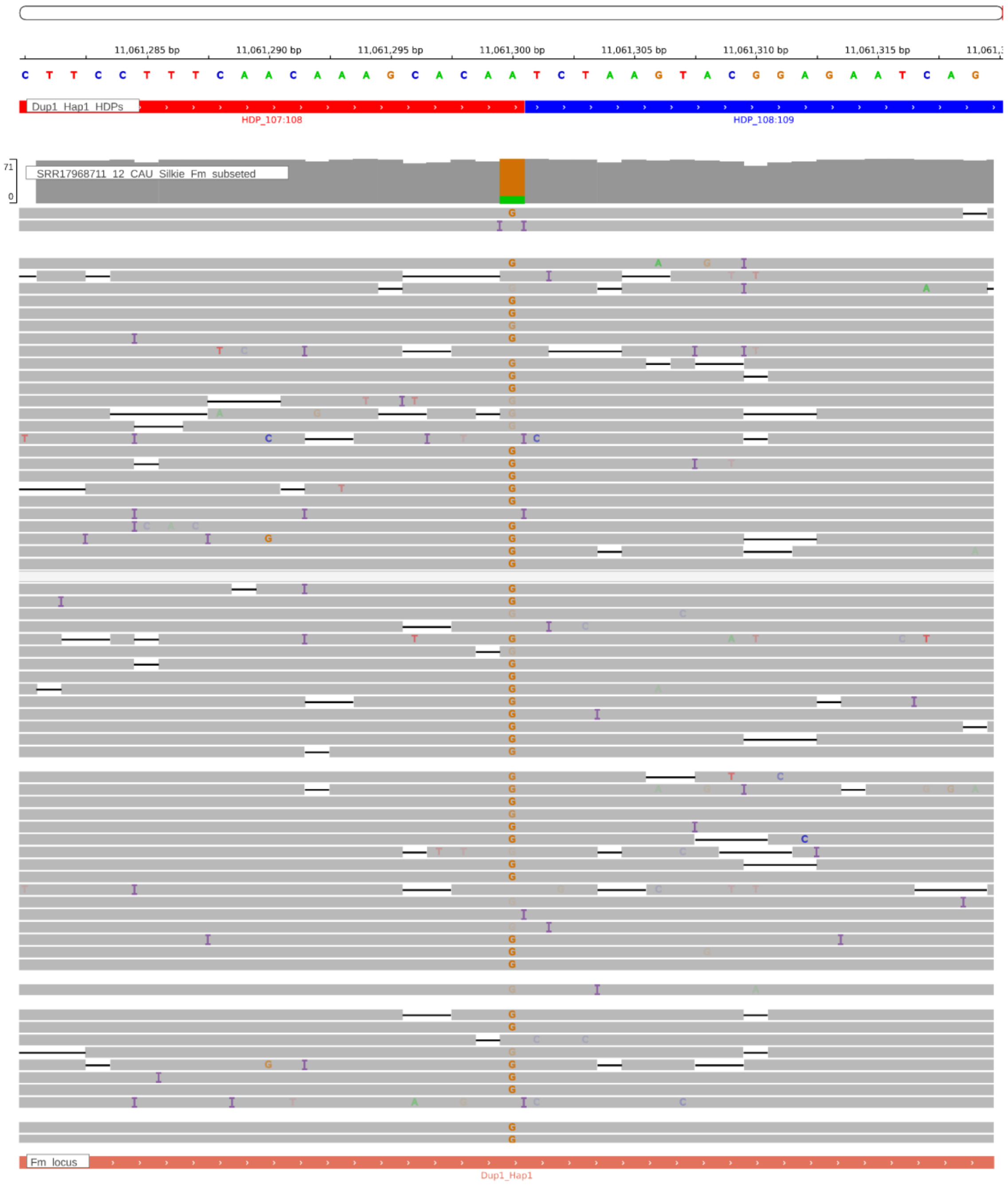


**Supplementary Figure. S127:** Screenshot from IGV showing the HDP 11061300 within the Dup1 region of the CAU Silkie genome. The CAU Silkie ONT long-reads were aligned to the CAU Silkie genome using minimap2, following the settings described by Zhu et al. The alignment reveals that support for the HDP within the genome is notably poor, indicating a mosaic genome assembly due to haplotype switching. The genomic base assembled at the given HDP position actually belongs to the Dup1r haplotype.

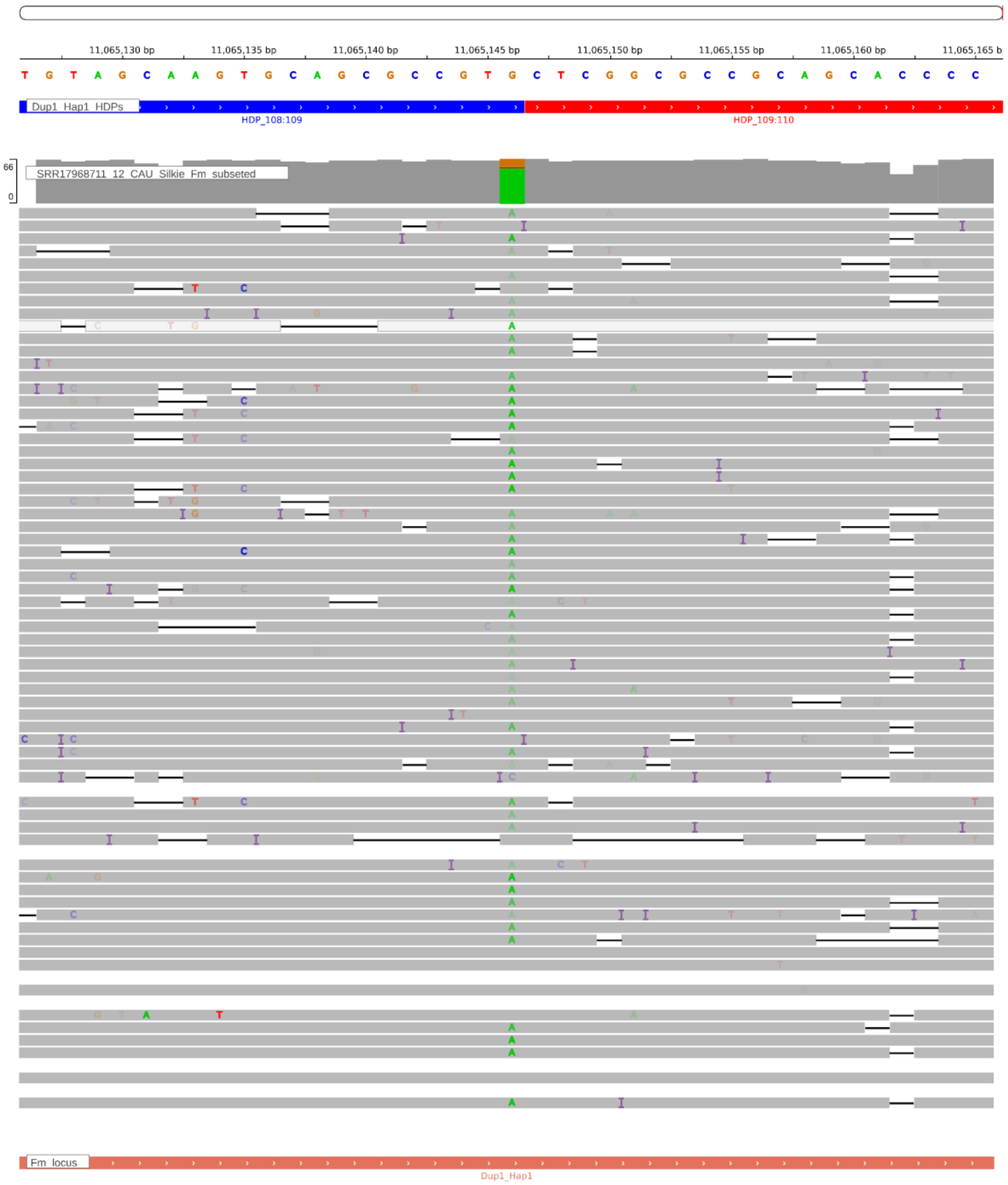


**Supplementary Figure. S128:** Screenshot from IGV showing the HDP 11065146 within the Dup1 region of the CAU Silkie genome. The CAU Silkie ONT long-reads were aligned to the CAU Silkie genome using minimap2, following the settings described by Zhu et al. The alignment reveals that support for the HDP within the genome is notably poor, indicating a mosaic genome assembly due to haplotype switching. The genomic base assembled at the given HDP position actually belongs to the Dup1r haplotype.

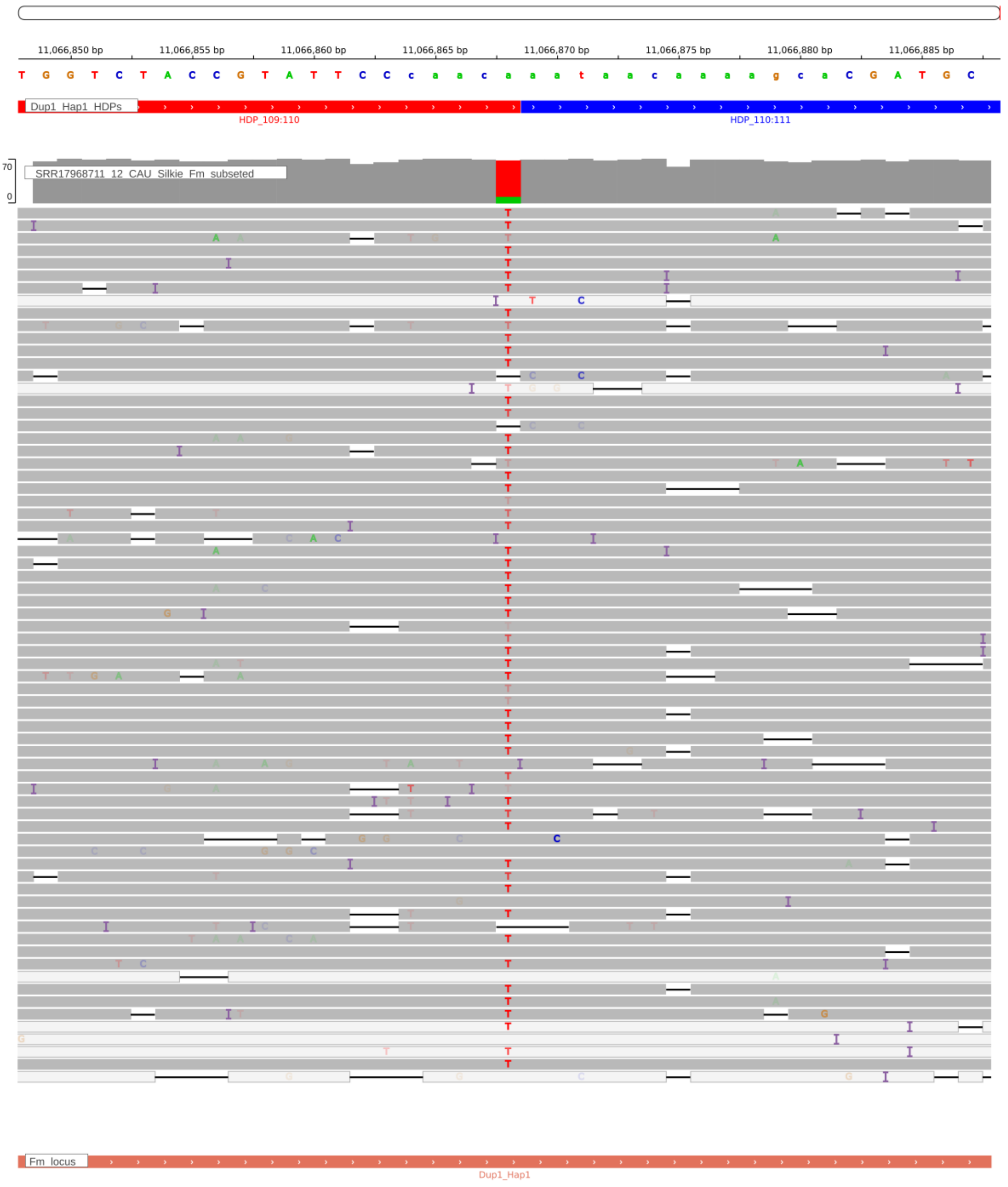


**Supplementary Figure. S129:** Screenshot from IGV showing the HDP 11066868 within the Dup1 region of the CAU Silkie genome. The CAU Silkie ONT long-reads were aligned to the CAU Silkie genome using minimap2, following the settings described by Zhu et al. The alignment reveals that support for the HDP within the genome is notably poor, indicating a mosaic genome assembly due to haplotype switching. The genomic base assembled at the given HDP position actually belongs to the Dup1r haplotype.

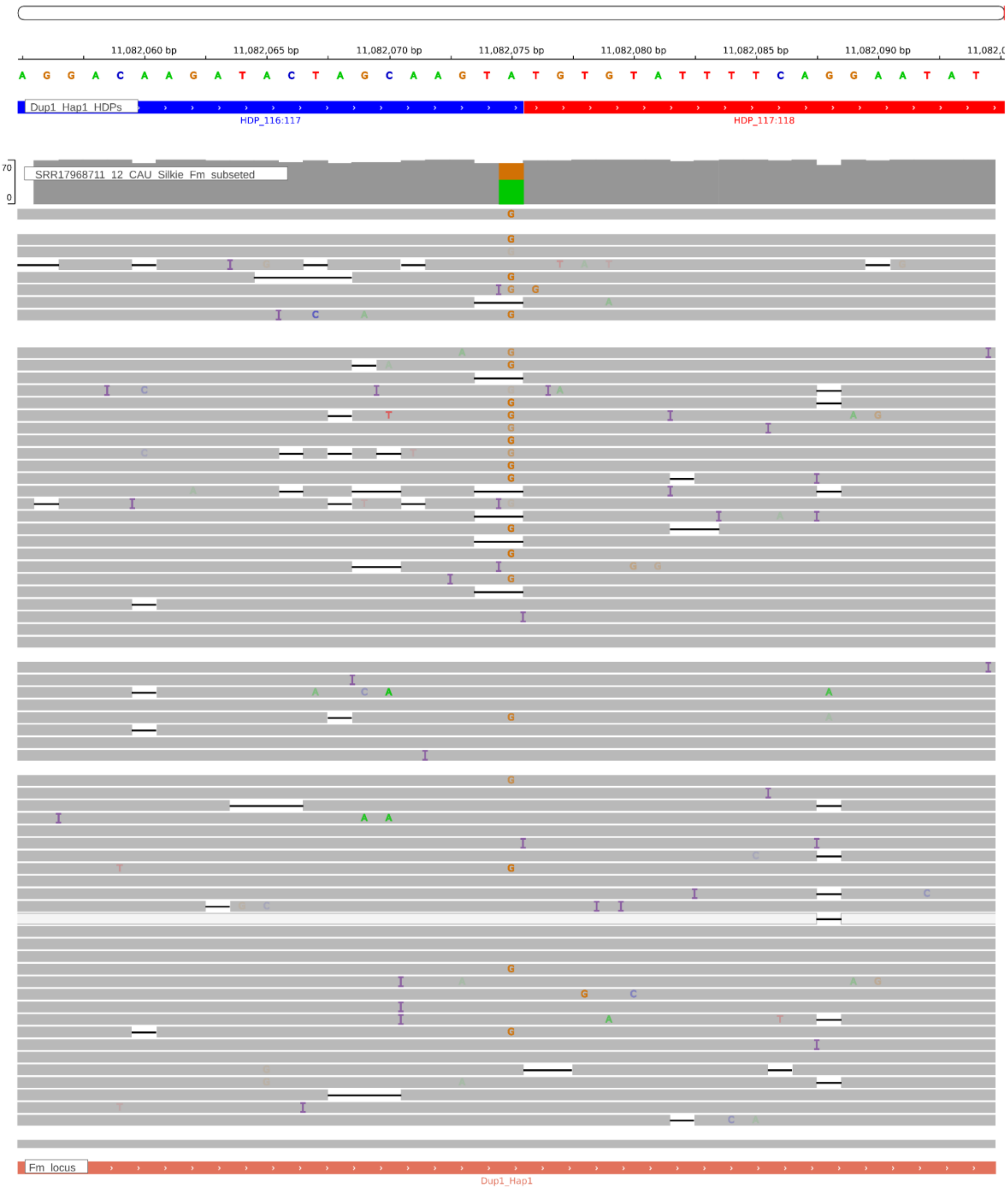


**Supplementary Figure. S130:** Screenshot from IGV showing the HDP 11082075 within the Dup1 region of the CAU Silkie genome. The CAU Silkie ONT long-reads were aligned to the CAU Silkie genome using minimap2, following the settings described by Zhu et al. The alignment reveals that support for the HDP within the genome is notably poor, indicating a mosaic genome assembly due to haplotype switching. The genomic base assembled at the given HDP position actually belongs to the Dup1r haplotype.

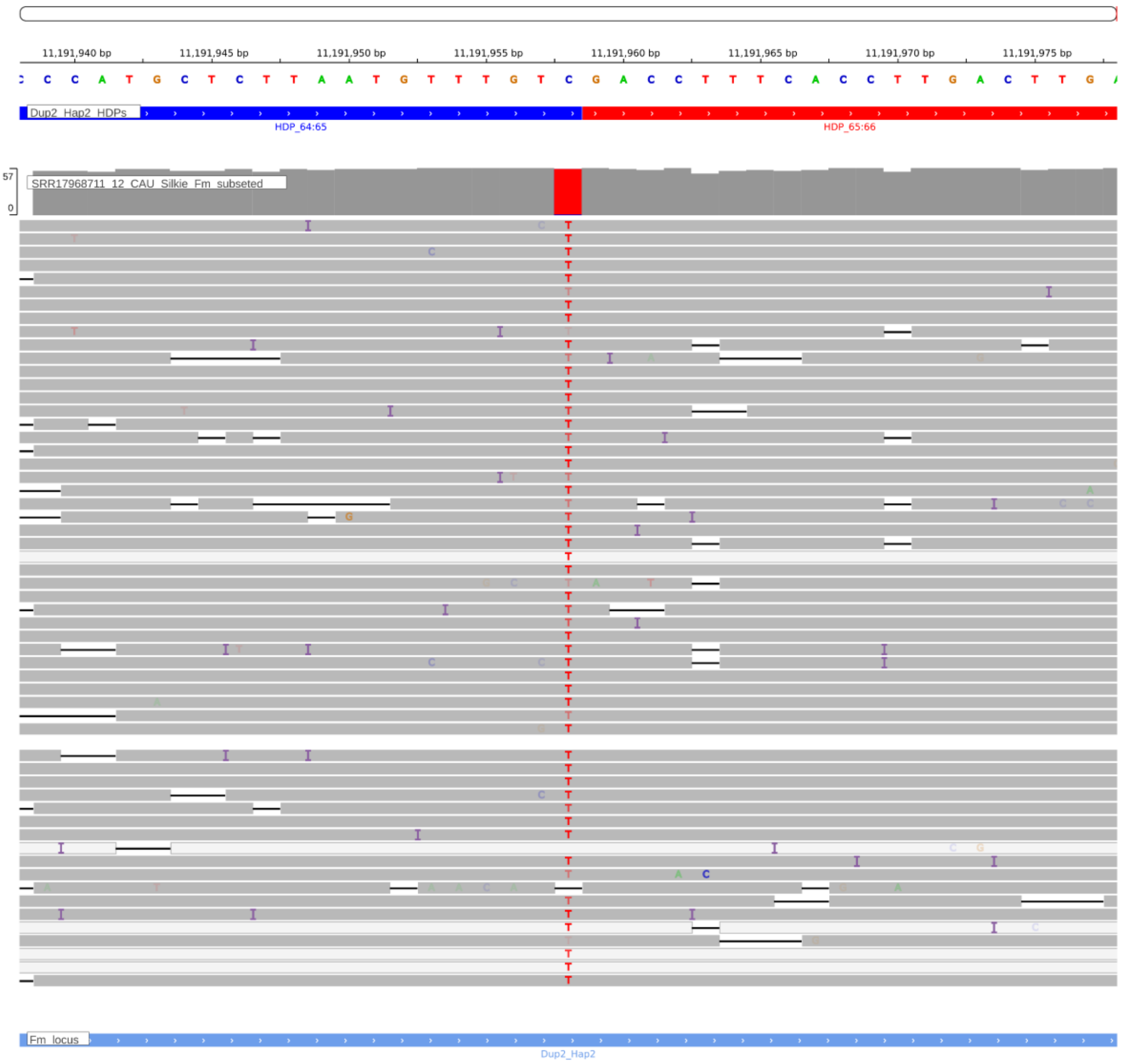


**Supplementary Figure. S131:** Screenshot from IGV showing the HDP 11191958 within the Dup2r region of the CAU Silkie genome. The CAU Silkie ONT long-reads were aligned to the CAU Silkie genome using minimap2, following the settings described by Zhu et al. The alignment reveals that support for the HDP within the genome is notably poor, indicating a mosaic genome assembly due to haplotype switching. The genomic base assembled at the given HDP position actually belongs to the Dup2 haplotype.

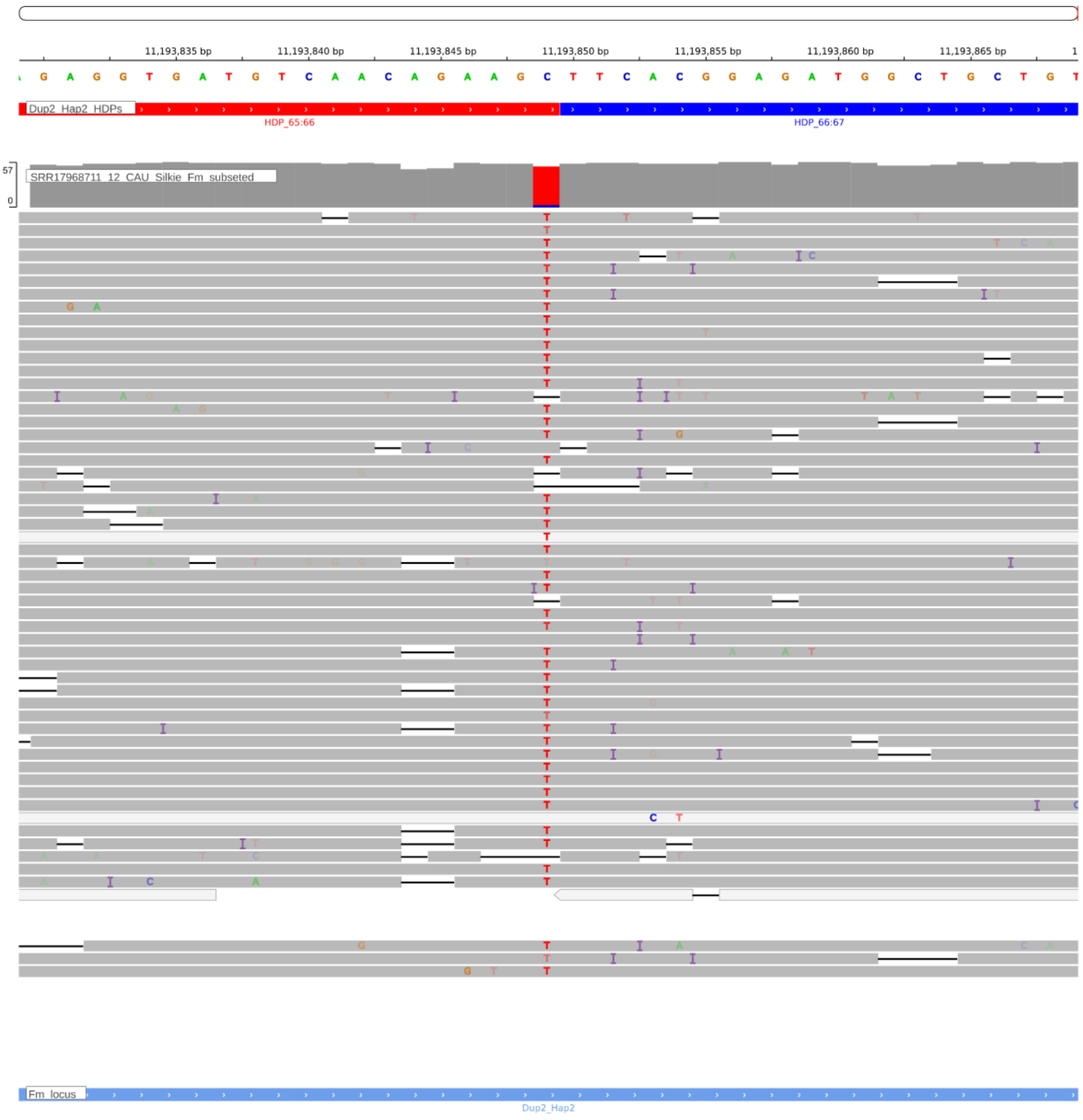


**Supplementary Figure. S132:** Screenshot from IGV showing the HDP 11193849 within the Dup2r region of the CAU Silkie genome. The CAU Silkie ONT long-reads were aligned to the CAU Silkie genome using minimap2, following the settings described by Zhu et al. The alignment reveals that support for the HDP within the genome is notably poor, indicating a mosaic genome assembly due to haplotype switching. The genomic base assembled at the given HDP position actually belongs to the Dup2 haplotype.

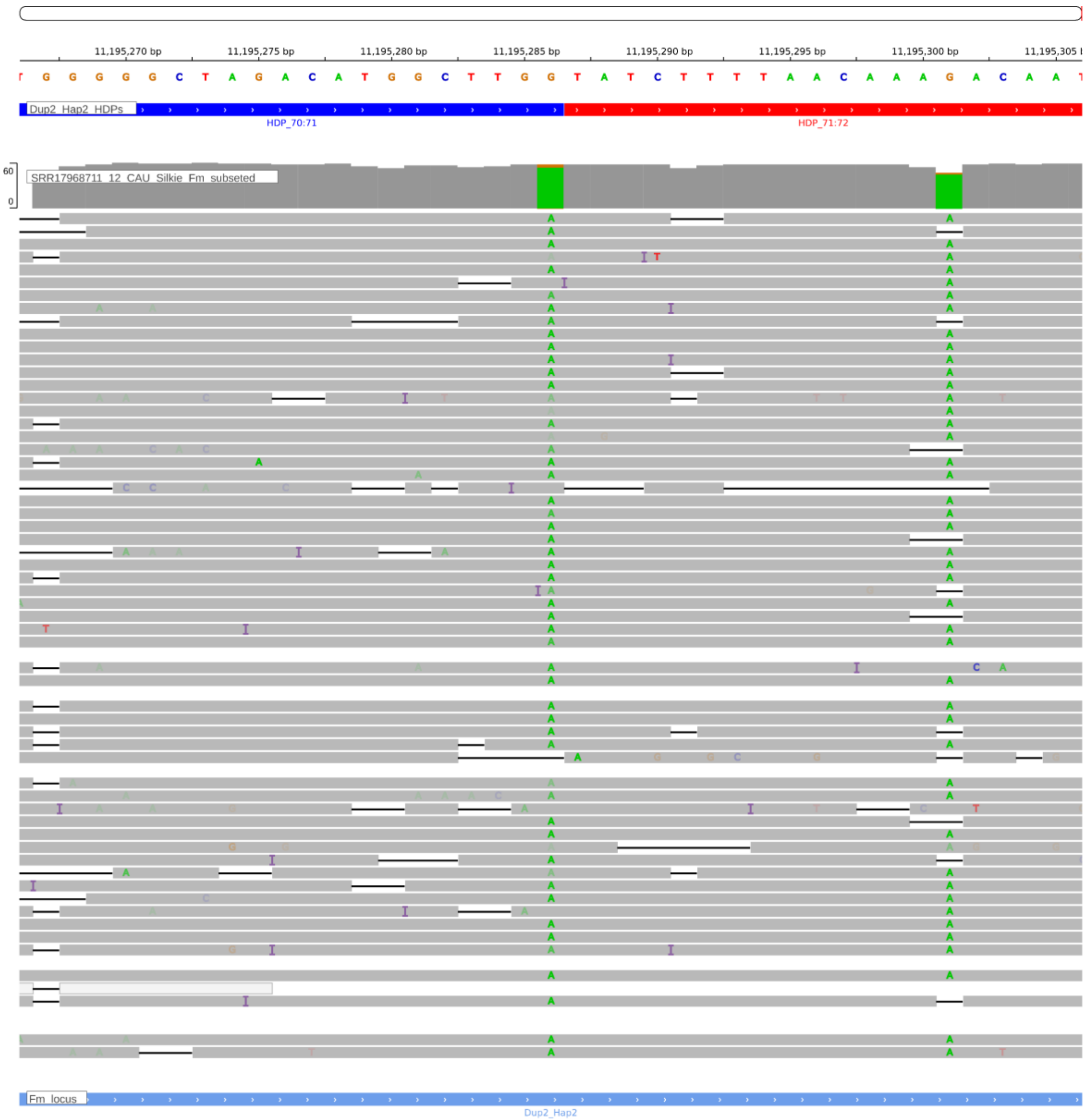


**Supplementary Figure. S133:** Screenshot from IGV showing the HDP 11195286 within the Dup2r region of the CAU Silkie genome. The CAU Silkie ONT long-reads were aligned to the CAU Silkie genome using minimap2, following the settings described by Zhu et al. The alignment reveals that support for the HDP within the genome is notably poor, indicating a mosaic genome assembly due to haplotype switching. The genomic base assembled at the given HDP position actually belongs to the Dup2 haplotype.

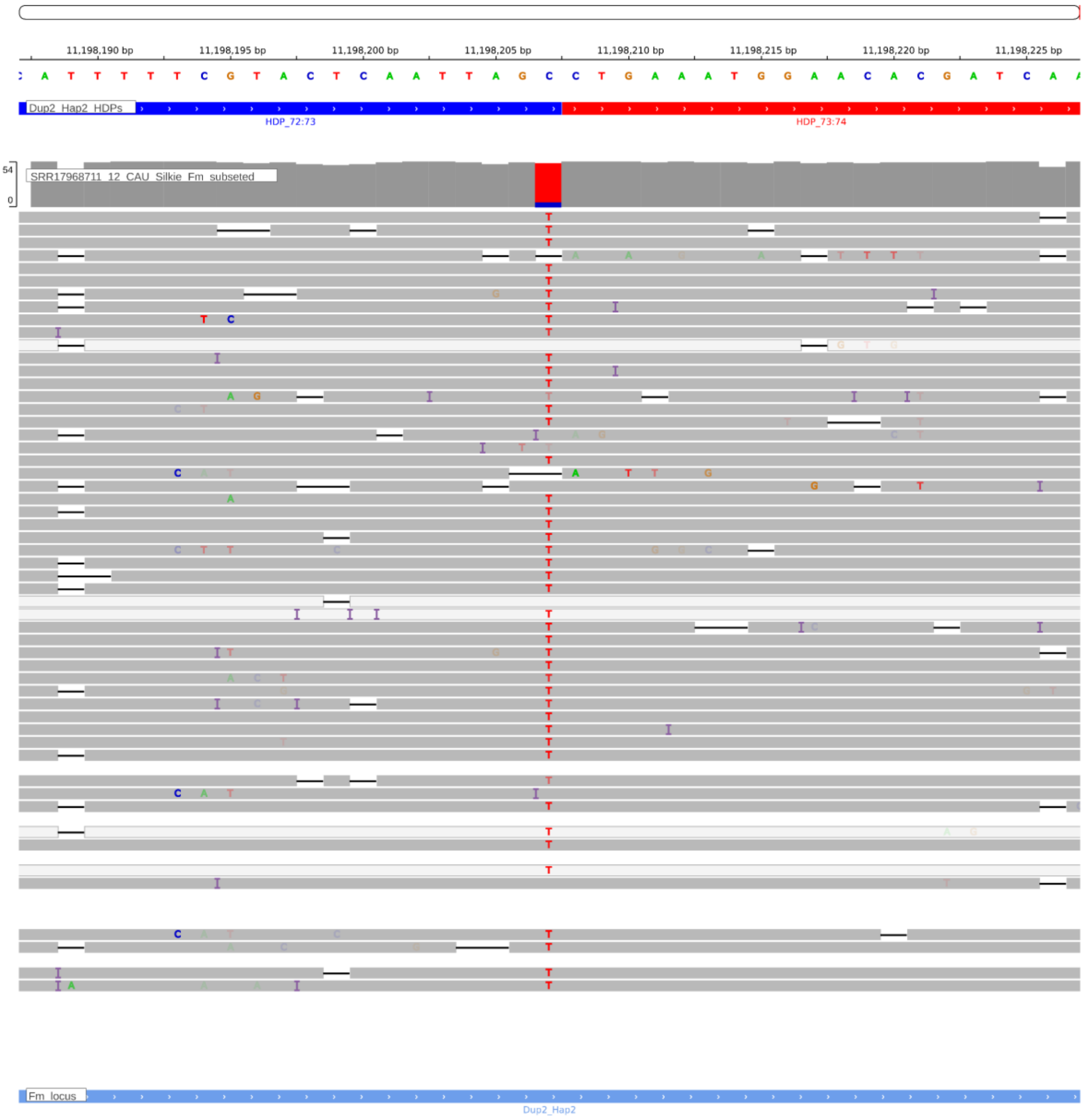


**Supplementary Figure. S134:** Screenshot from IGV showing the HDP 11198207 within the Dup2r region of the CAU Silkie genome. The CAU Silkie ONT long-reads were aligned to the CAU Silkie genome using minimap2, following the settings described by Zhu et al. The alignment reveals that support for the HDP within the genome is notably poor, indicating a mosaic genome assembly due to haplotype switching. The genomic base assembled at the given HDP position actually belongs to the Dup2 haplotype.

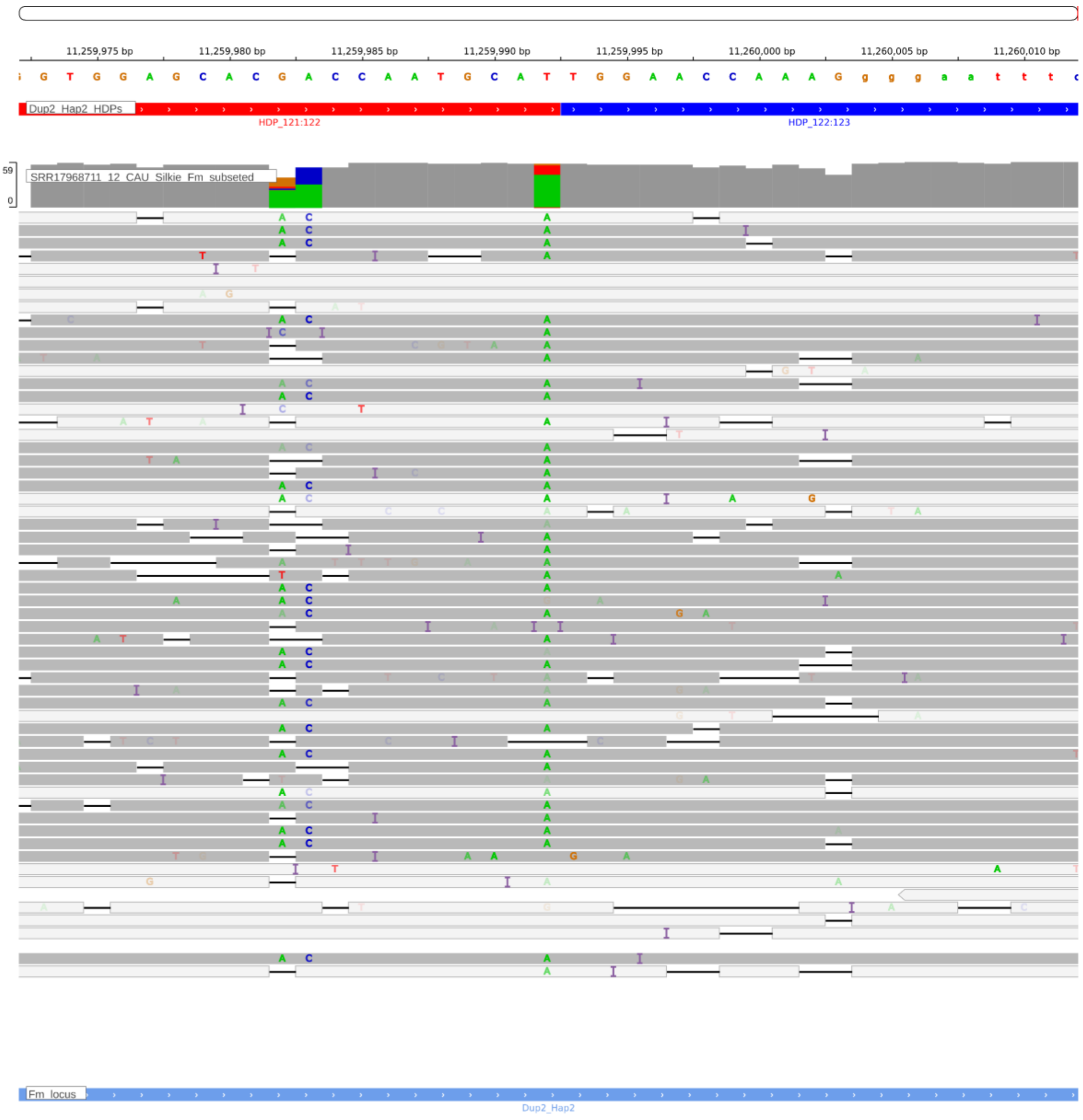


**Supplementary Figure. S135:** Screenshot from IGV showing the HDP 11259992 within the Dup2r region of the CAU Silkie genome. The CAU Silkie ONT long-reads were aligned to the CAU Silkie genome using minimap2, following the settings described by Zhu et al. The alignment reveals that support for the HDP within the genome is notably poor, indicating a mosaic genome assembly due to haplotype switching. The genomic base assembled at the given HDP position actually belongs to the Dup2 haplotype.

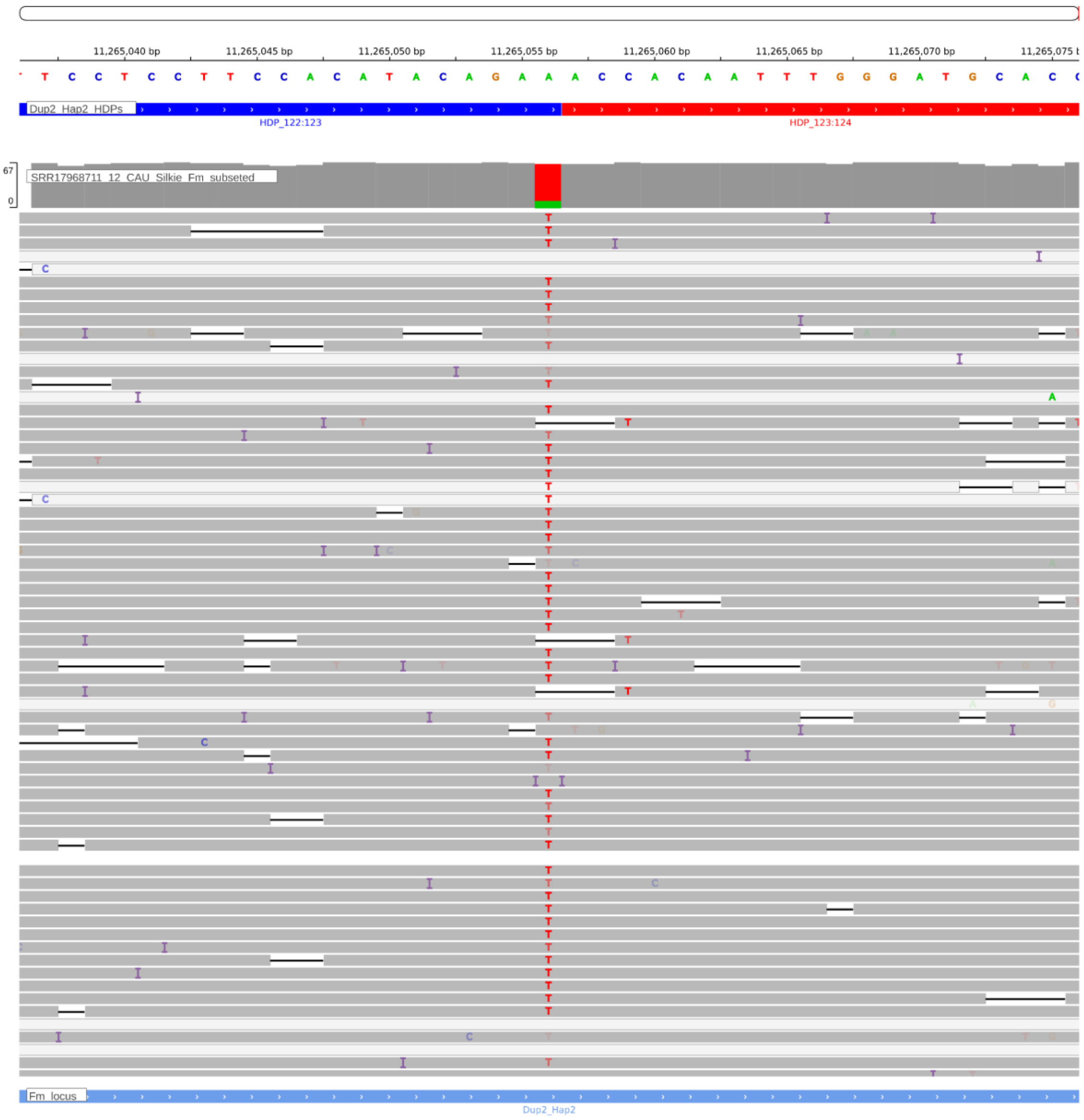


**Supplementary Figure. S136:** Screenshot from IGV showing the HDP 11265056 within the Dup2r region of the CAU Silkie genome. The CAU Silkie ONT long-reads were aligned to the CAU Silkie genome using minimap2, following the settings described by Zhu et al. The alignment reveals that support for the HDP within the genome is notably poor, indicating a mosaic genome assembly due to haplotype switching. The genomic base assembled at the given HDP position actually belongs to the Dup2 haplotype.

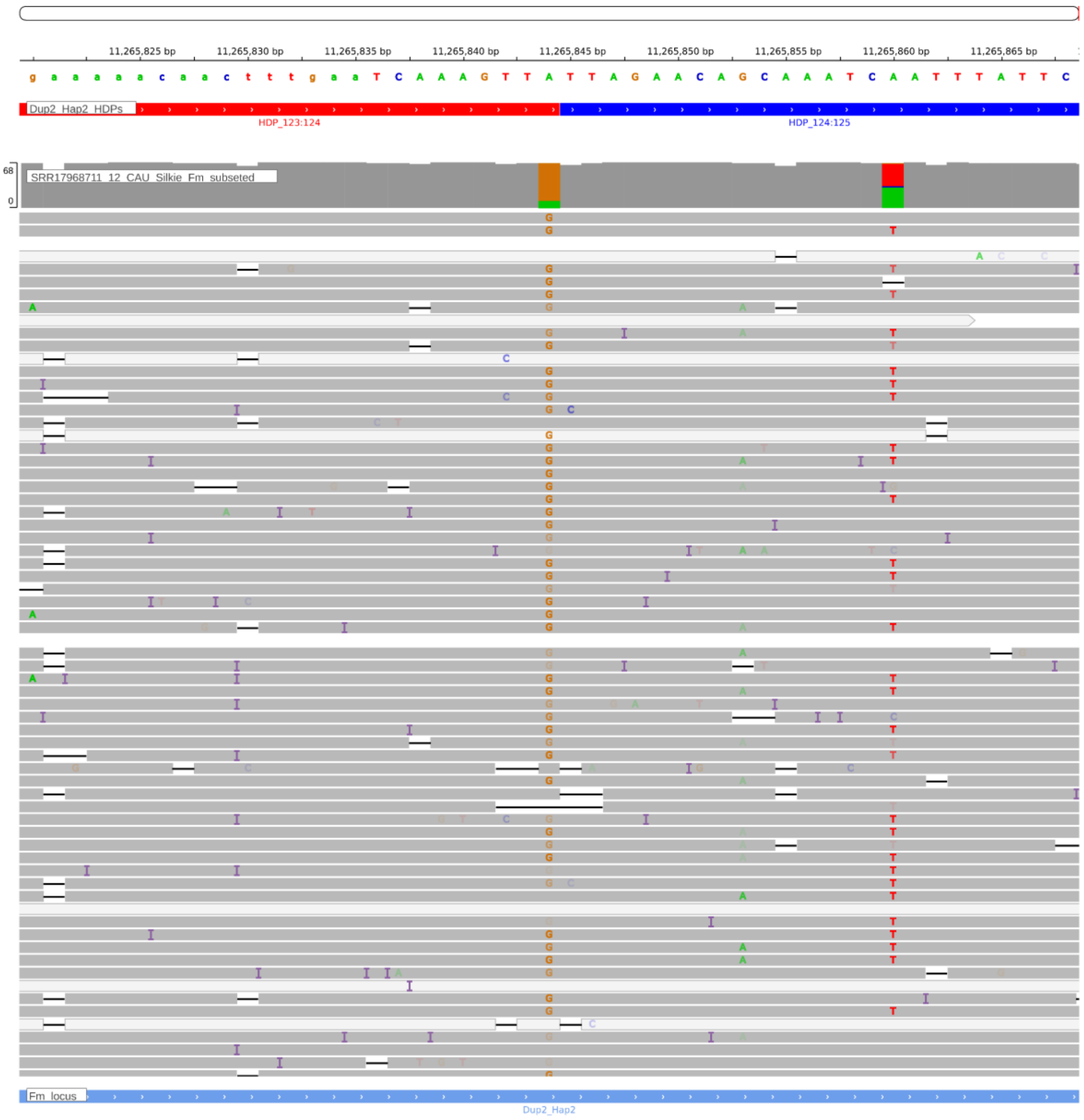


**Supplementary Figure. S137:** Screenshot from IGV showing the HDP 11265844 within the Dup2r region of the CAU Silkie genome. The CAU Silkie ONT long-reads were aligned to the CAU Silkie genome using minimap2, following the settings described by Zhu et al. The alignment reveals that support for the HDP within the genome is notably poor, indicating a mosaic genome assembly due to haplotype switching. The genomic base assembled at the given HDP position actually belongs to the Dup2 haplotype.

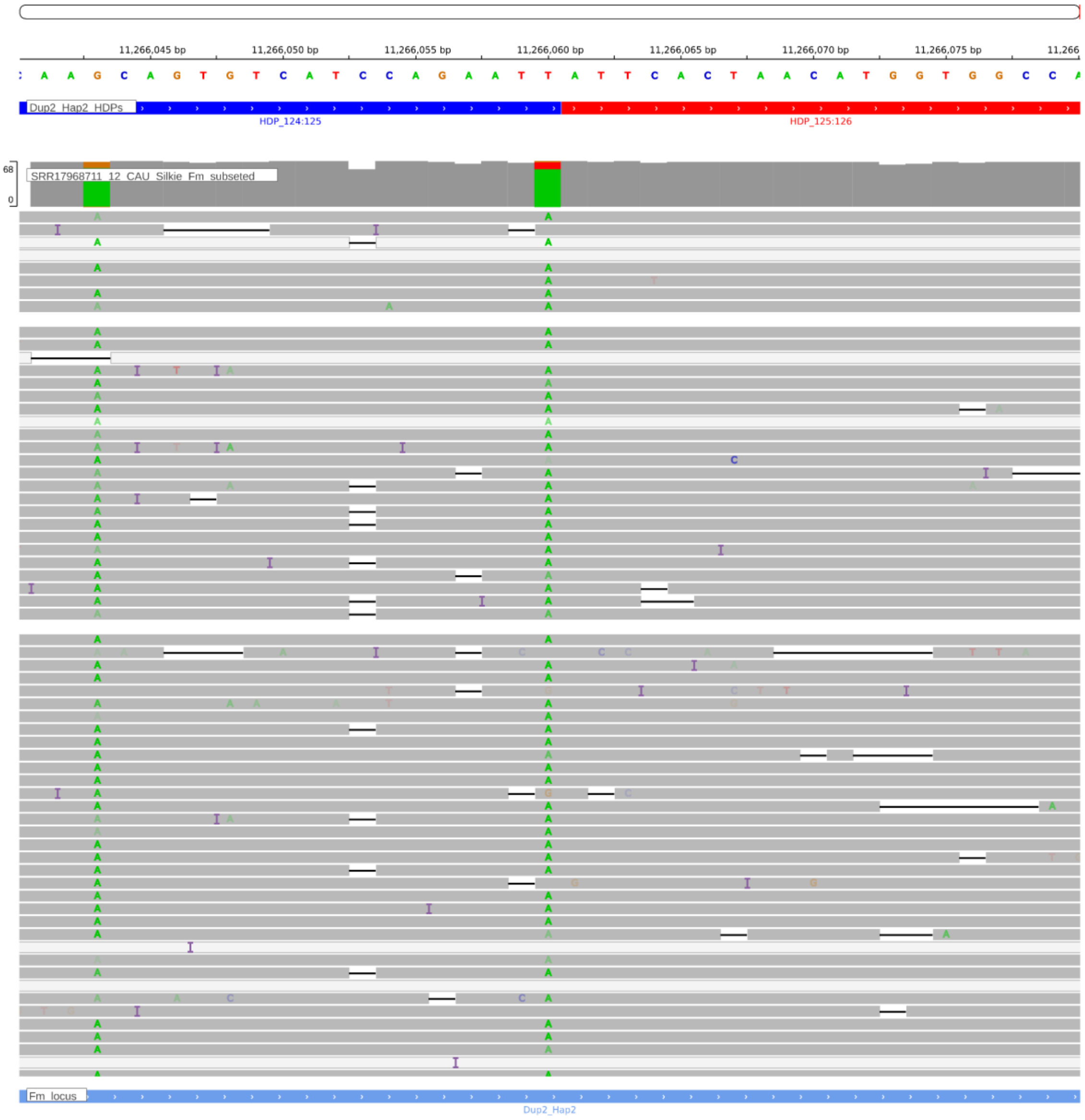


**Supplementary Figure. S138:** Screenshot from IGV showing the HDP 11268625 within the Dup2r region of the CAU Silkie genome. The CAU Silkie ONT long-reads were aligned to the CAU Silkie genome using minimap2, following the settings described by Zhu et al. The alignment reveals that support for the HDP within the genome is notably poor, indicating a mosaic genome assembly due to haplotype switching. The genomic base assembled at the given HDP position actually belongs to the Dup2 haplotype.

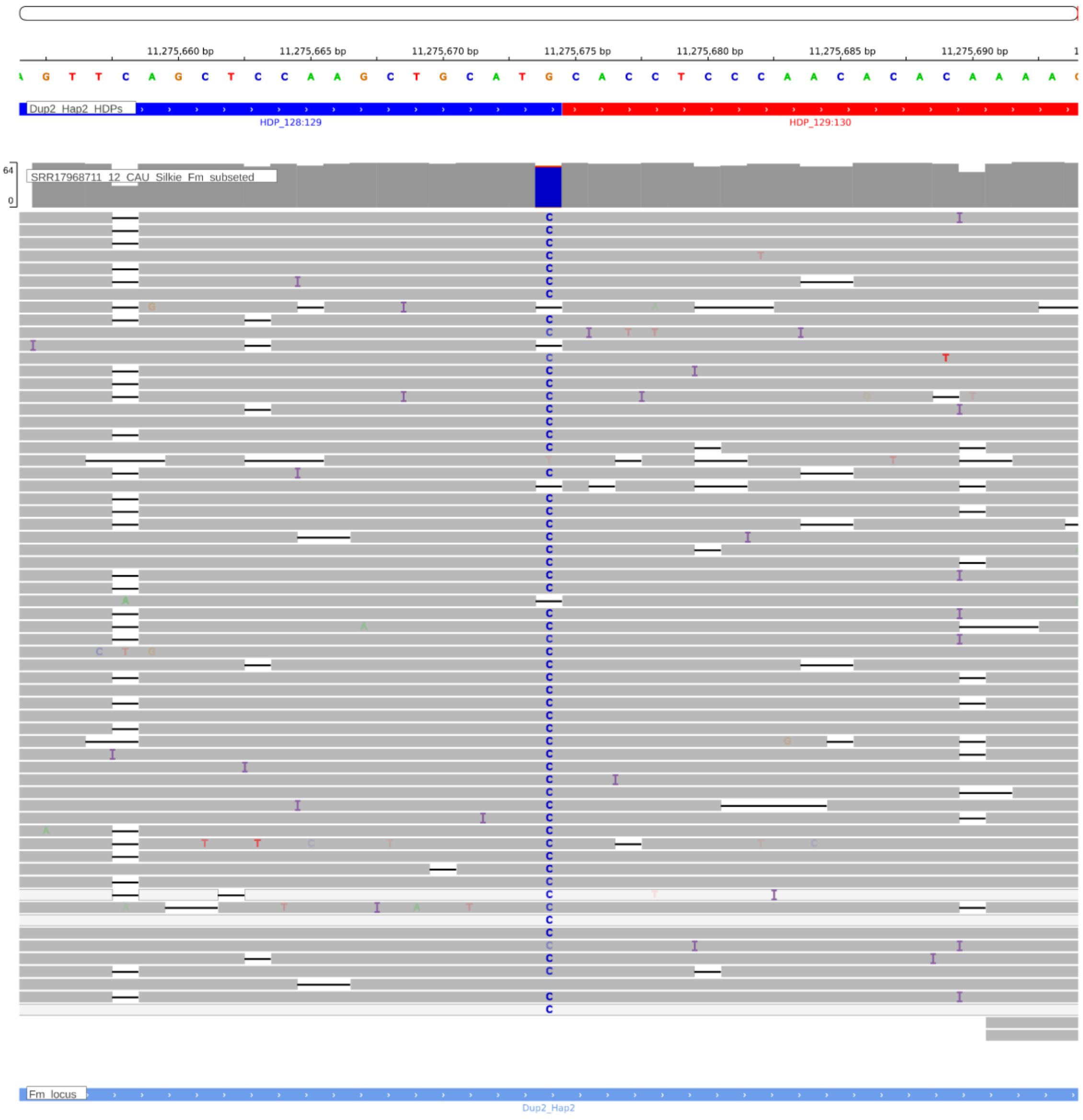


**Supplementary Figure. S139:** Screenshot from IGV showing the HDP 11275674 within the Dup2r region of the CAU Silkie genome. The CAU Silkie ONT long-reads were aligned to the CAU Silkie genome using minimap2, following the settings described by Zhu et al. The alignment reveals that support for the HDP within the genome is notably poor, indicating a mosaic genome assembly due to haplotype switching. The genomic base assembled at the given HDP position actually belongs to the Dup2 haplotype.

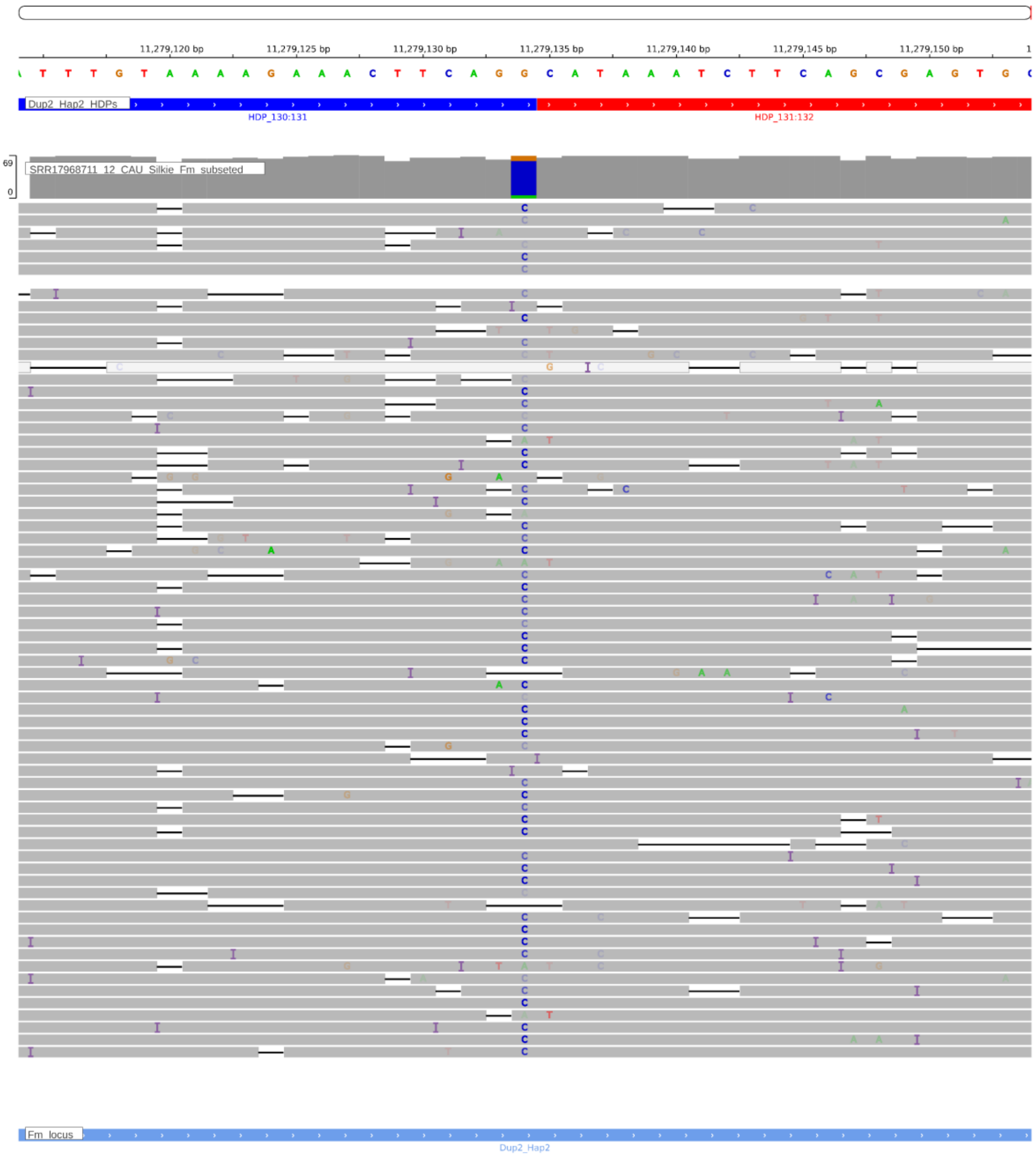


**Supplementary Figure. S140:** Screenshot from IGV showing the HDP 11279134 within the Dup2r region of the CAU Silkie genome. The CAU Silkie ONT long-reads were aligned to the CAU Silkie genome using minimap2, following the settings described by Zhu et al. The alignment reveals that support for the HDP within the genome is notably poor, indicating a mosaic genome assembly due to haplotype switching. The genomic base assembled at the given HDP position actually belongs to the Dup2 haplotype.

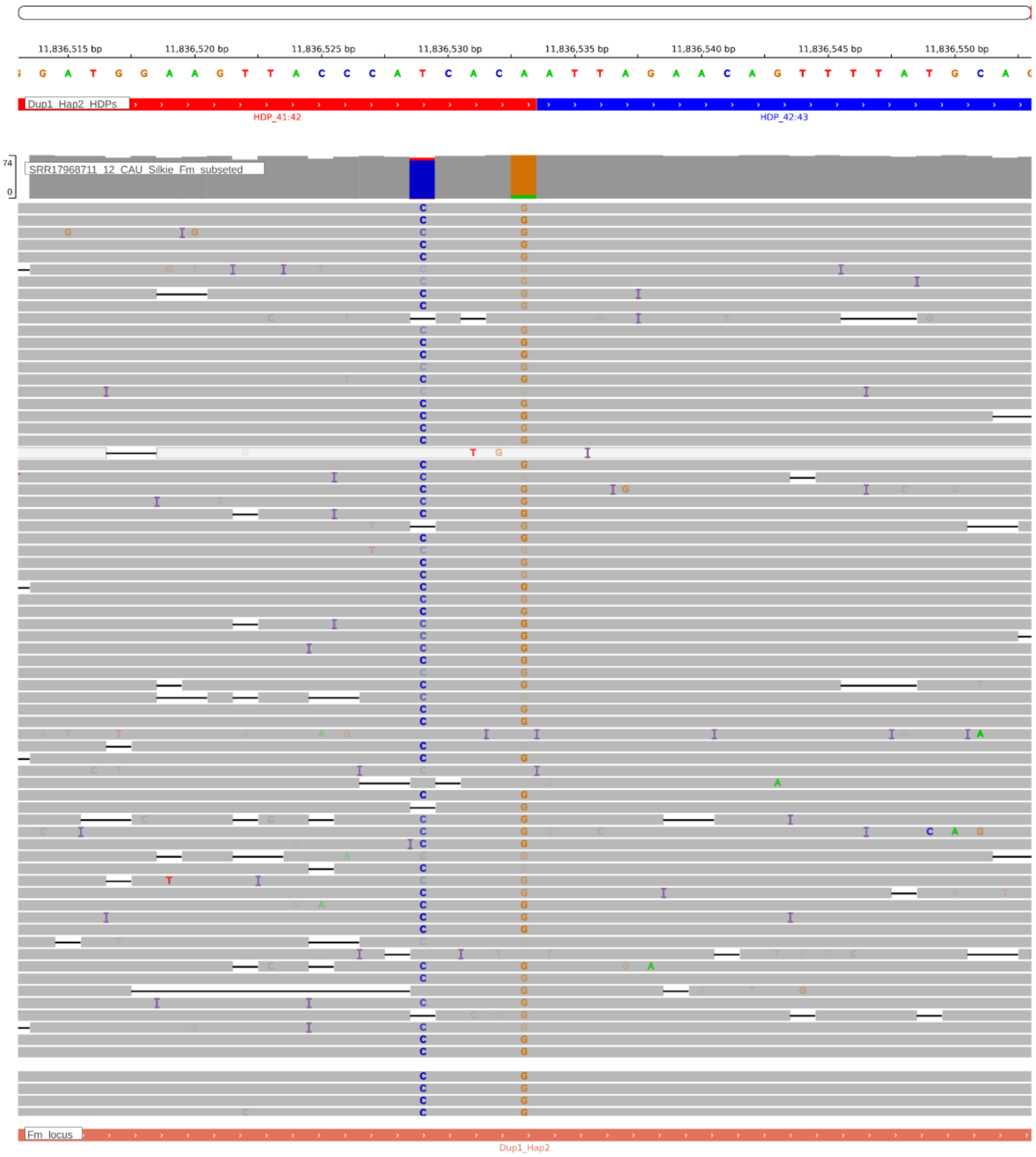


**Supplementary Figure. S141:** Screenshot from IGV showing the HDP 11836533 within the Dup1r region of the CAU Silkie genome. The CAU Silkie ONT long-reads were aligned to the CAU Silkie genome using minimap2, following the settings described by Zhu et al. The alignment reveals that support for the HDP within the genome is notably poor, indicating a mosaic genome assembly due to haplotype switching. The genomic base assembled at the given HDP position actually belongs to the Dup1 haplotype.

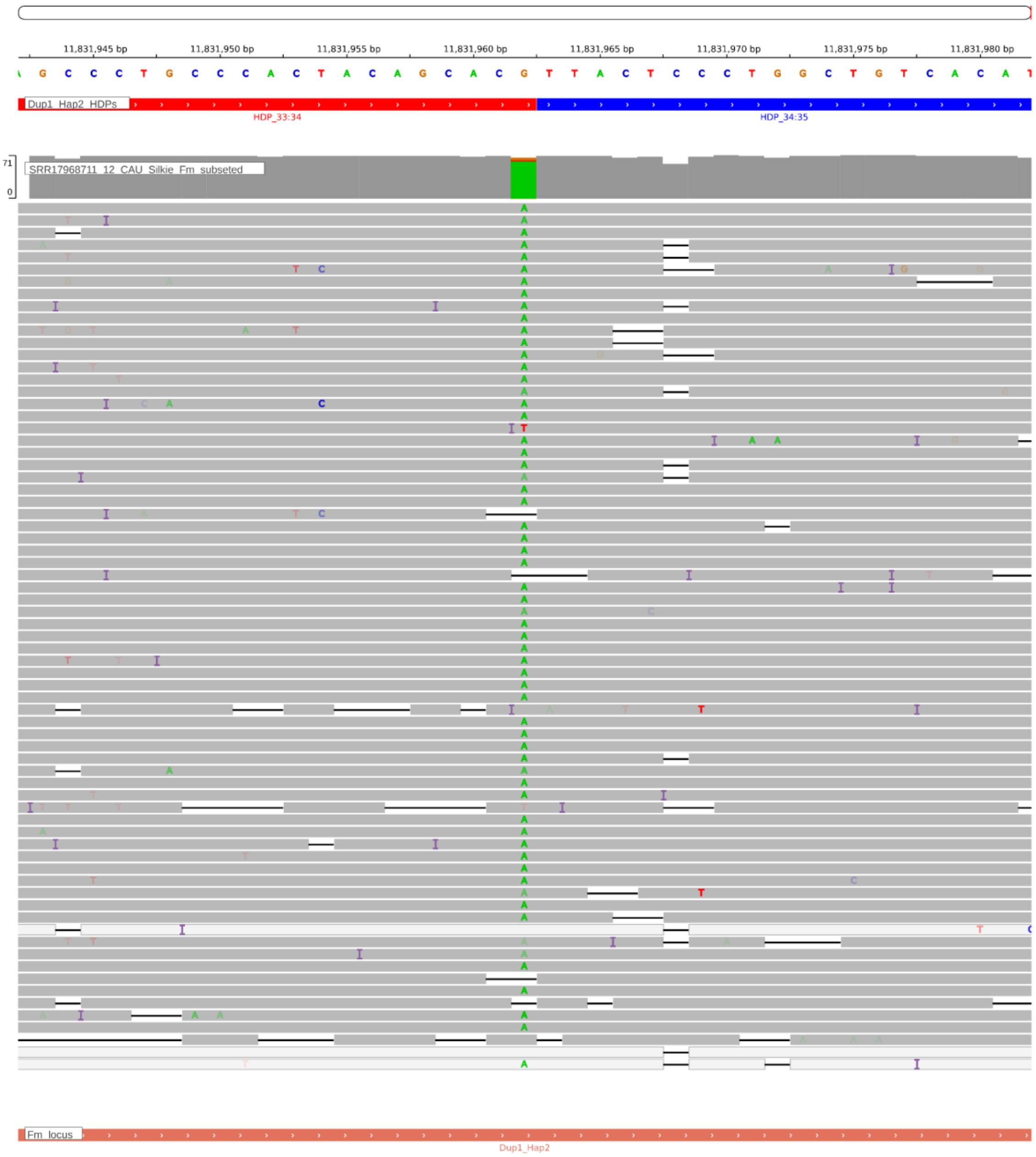


**Supplementary Figure. S142:** Screenshot from IGV showing the HDP 11831962 within the Dup1r region of the CAU Silkie genome. The CAU Silkie ONT long-reads were aligned to the CAU Silkie genome using minimap2, following the settings described by Zhu et al. The alignment reveals that support for the HDP within the genome is notably poor, indicating a mosaic genome assembly due to haplotype switching. The genomic base assembled at the given HDP position actually belongs to the Dup1 haplotype.

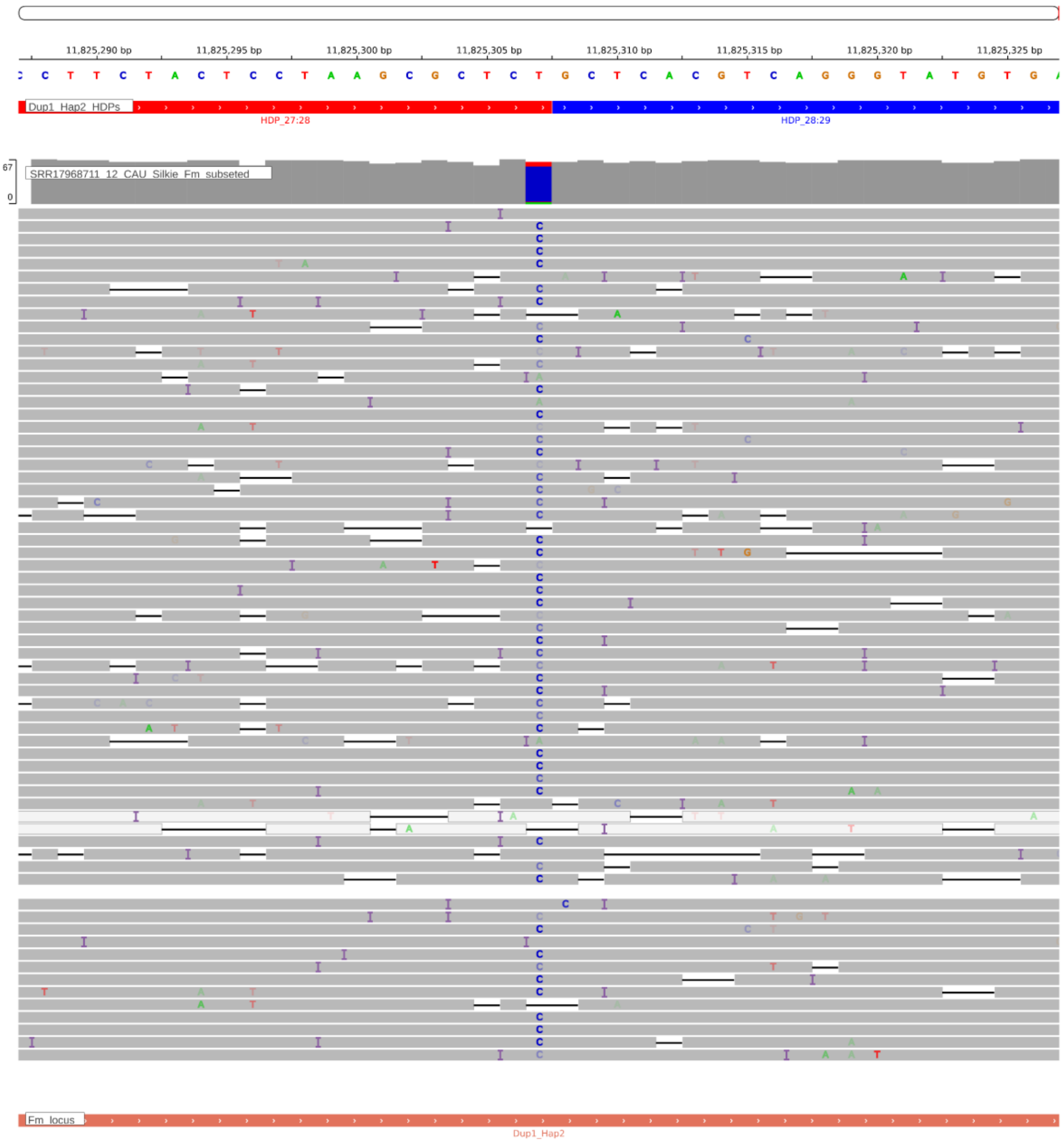


**Supplementary Figure. S143:** Screenshot from IGV showing the HDP 11825307 within the Dup1r region of the CAU Silkie genome. The CAU Silkie ONT long-reads were aligned to the CAU Silkie genome using minimap2, following the settings described by Zhu et al. The alignment reveals that support for the HDP within the genome is notably poor, indicating a mosaic genome assembly due to haplotype switching. The genomic base assembled at the given HDP position actually belongs to the Dup1 haplotype.

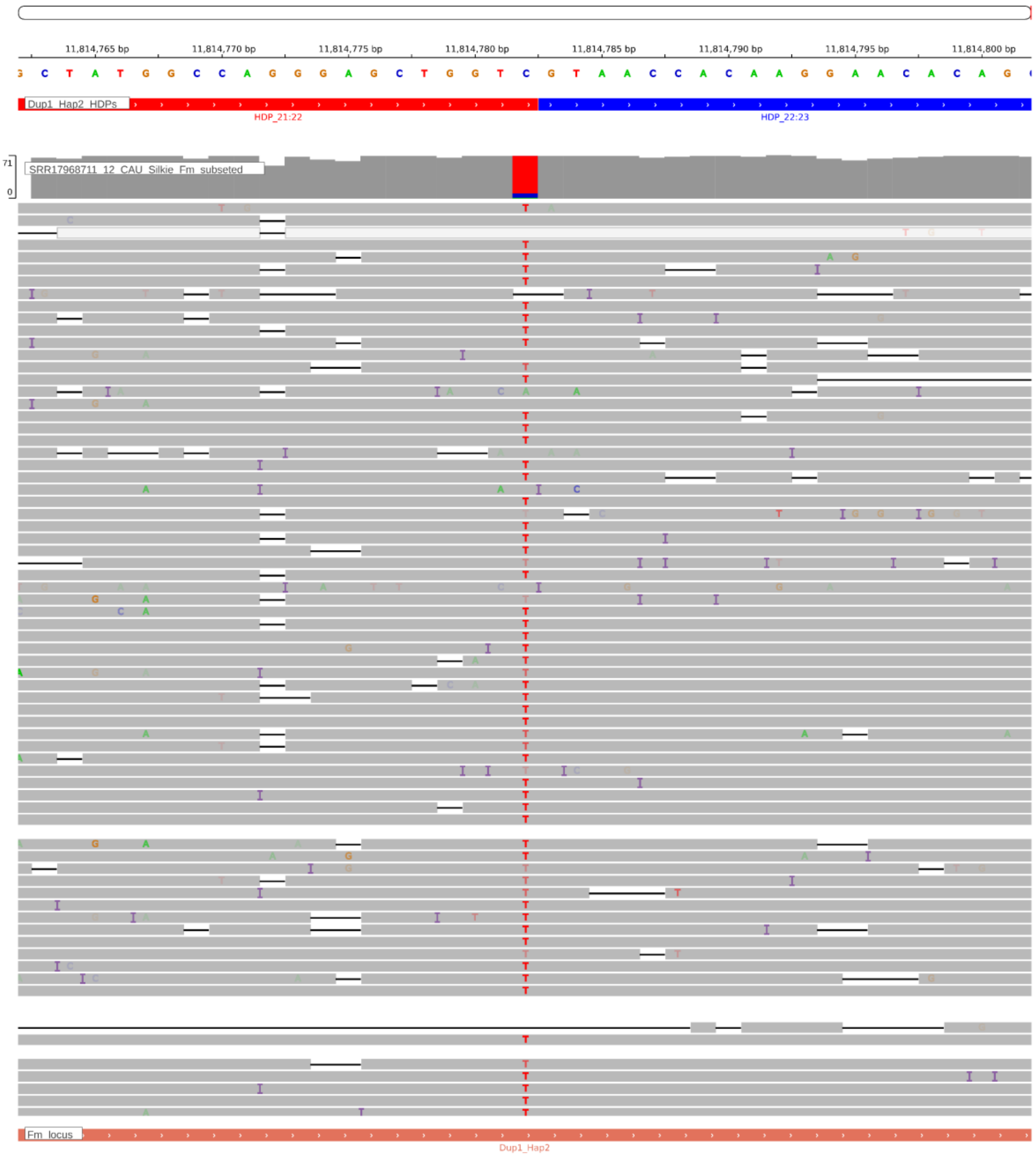


**Supplementary Figure. S144:** Screenshot from IGV showing the HDP 11814782 within the Dup1r region of the CAU Silkie genome. The CAU Silkie ONT long-reads were aligned to the CAU Silkie genome using minimap2, following the settings described by Zhu et al. The alignment reveals that support for the HDP within the genome is notably poor, indicating a mosaic genome assembly due to haplotype switching. The genomic base assembled at the given HDP position actually belongs to the Dup1 haplotype.

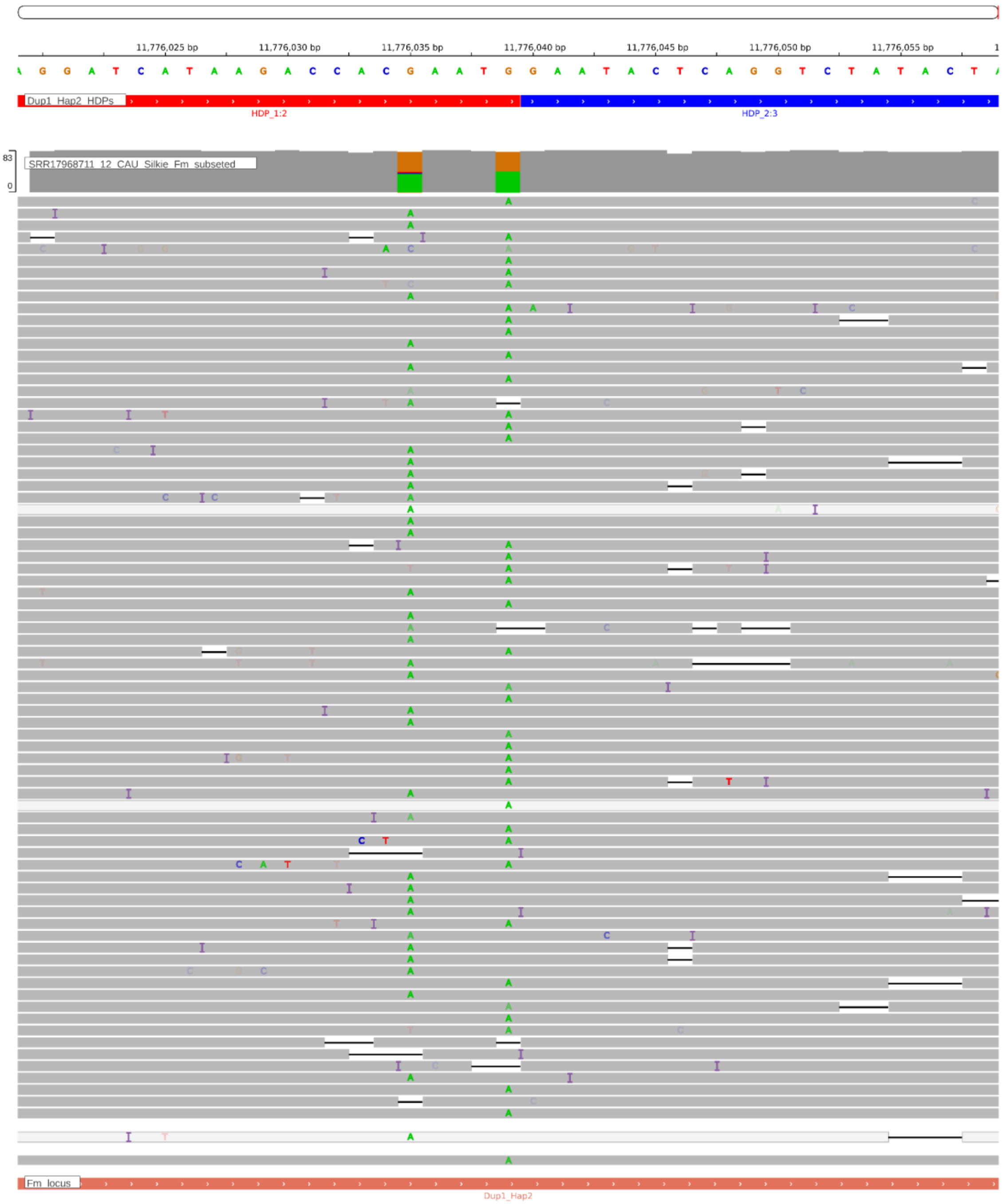


**Supplementary Figure. S145:** Screenshot from IGV showing the HDP 11776039 within the Dup1r region of the CAU Silkie genome. The CAU Silkie ONT long-reads were aligned to the CAU Silkie genome using minimap2, following the settings described by Zhu et al. The alignment reveals that support for the HDP within the genome is notably poor, indicating a mosaic genome assembly due to haplotype switching. The genomic base assembled at the given HDP position actually belongs to the Dup1 haplotype.

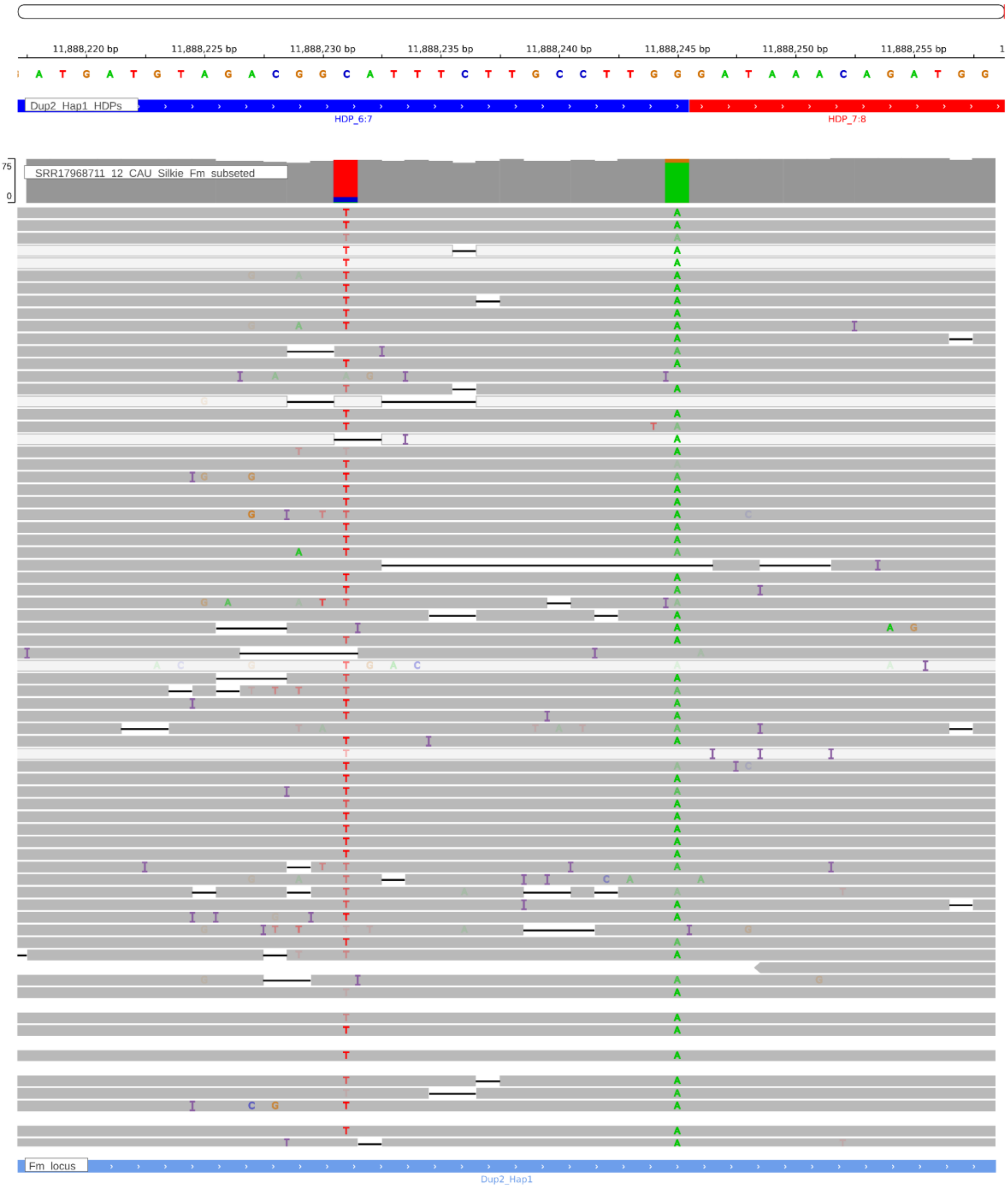


**Supplementary Figure. S146:** Screenshot from IGV showing the HDP 11888245 within the Dup2 region of the CAU Silkie genome. The CAU Silkie ONT long-reads were aligned to the CAU Silkie genome using minimap2, following the settings described by Zhu et al. The alignment reveals that support for the HDP within the genome is notably poor, indicating a mosaic genome assembly due to haplotype switching. The genomic base assembled at the given HDP position actually belongs to the Dup2r haplotype.

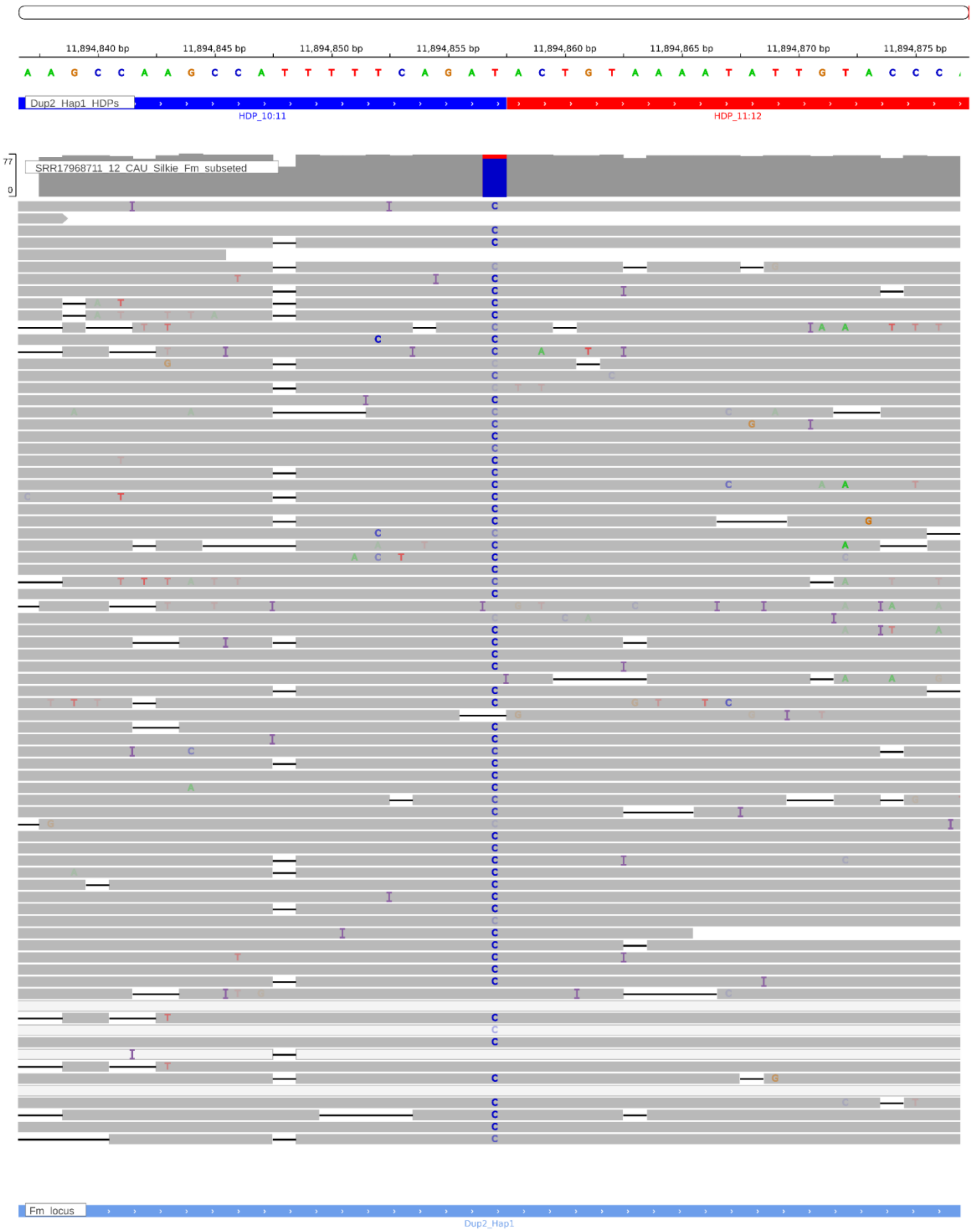


**Supplementary Figure. S147:** Screenshot from IGV showing the HDP 11894857 within the Dup2 region of the CAU Silkie genome. The CAU Silkie ONT long-reads were aligned to the CAU Silkie genome using minimap2, following the settings described by Zhu et al. The alignment reveals that support for the HDP within the genome is notably poor, indicating a mosaic genome assembly due to haplotype switching. The genomic base assembled at the given HDP position actually belongs to the Dup2r haplotype.

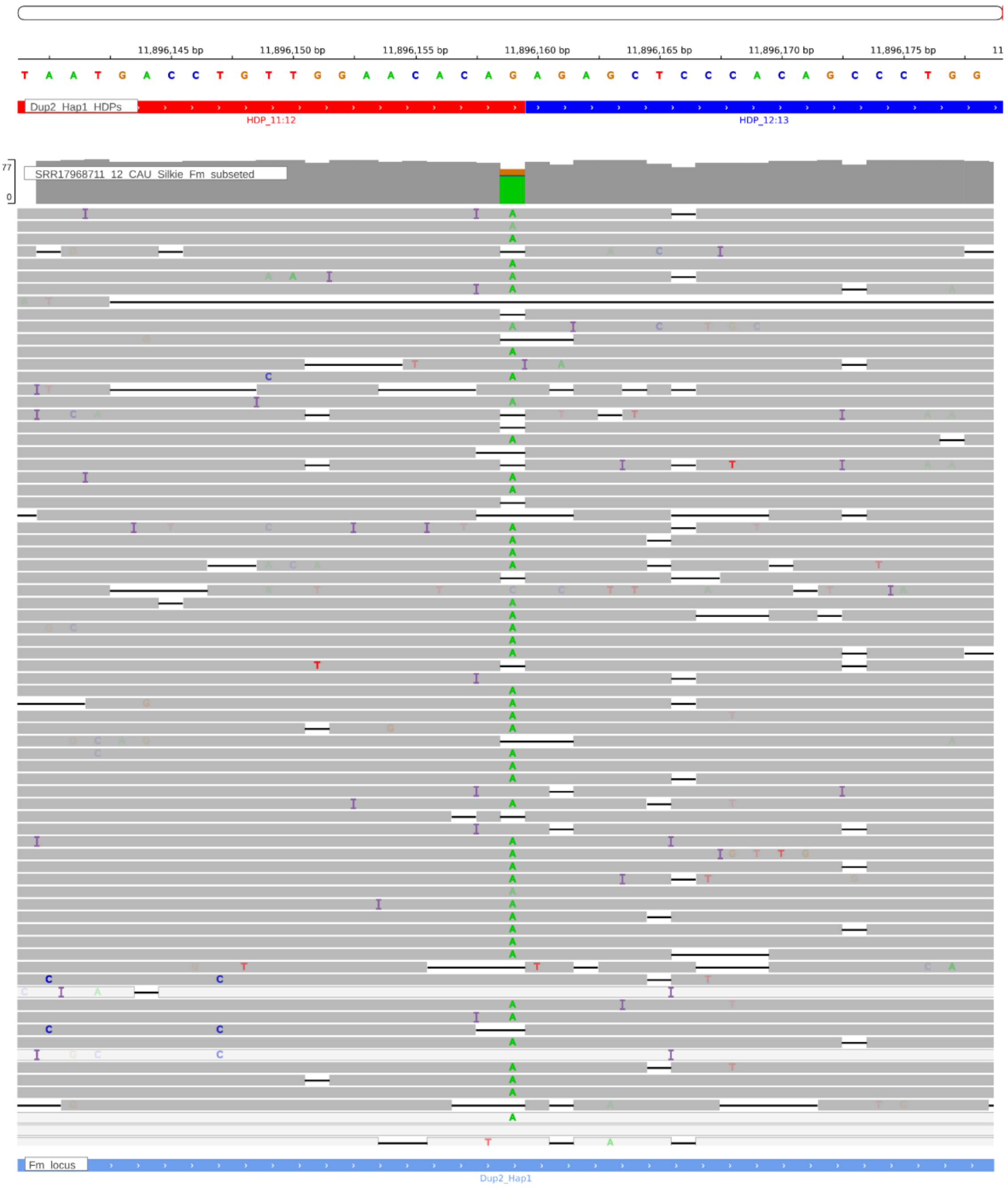


**Supplementary Figure. S148:** Screenshot from IGV showing the HDP 11896159 within the Dup2 region of the CAU Silkie genome. The CAU Silkie ONT long-reads were aligned to the CAU Silkie genome using minimap2, following the settings described by Zhu et al. The alignment reveals that support for the HDP within the genome is notably poor, indicating a mosaic genome assembly due to haplotype switching. The genomic base assembled at the given HDP position actually belongs to the Dup2r haplotype.

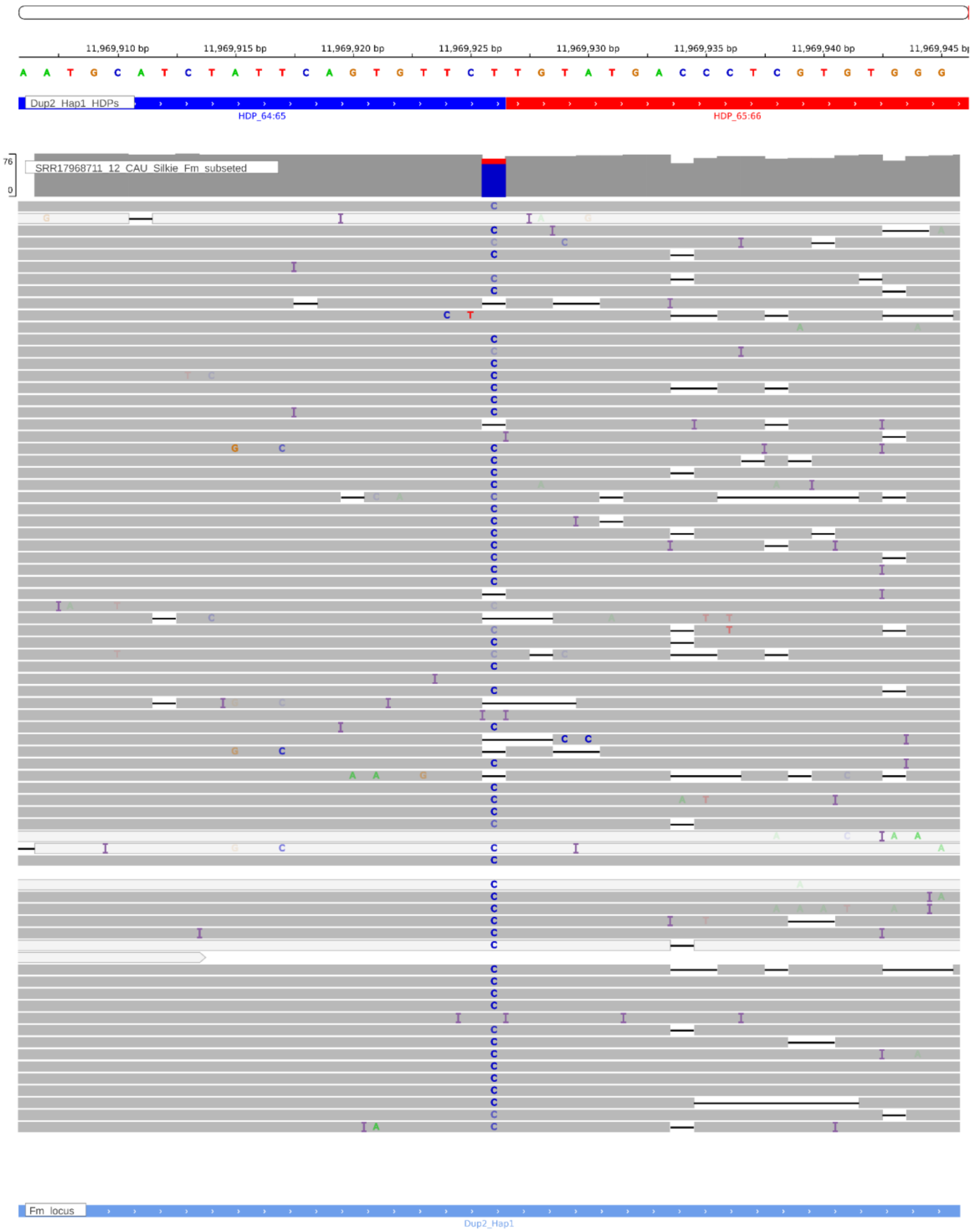


**Supplementary Figure. S149:** Screenshot from IGV showing the HDP 11969926 within the Dup2 region of the CAU Silkie genome. The CAU Silkie ONT long-reads were aligned to the CAU Silkie genome using minimap2, following the settings described by Zhu et al. The alignment reveals that support for the HDP within the genome is notably poor, indicating a mosaic genome assembly due to haplotype switching. The genomic base assembled at the given HDP position actually belongs to the Dup2r haplotype.

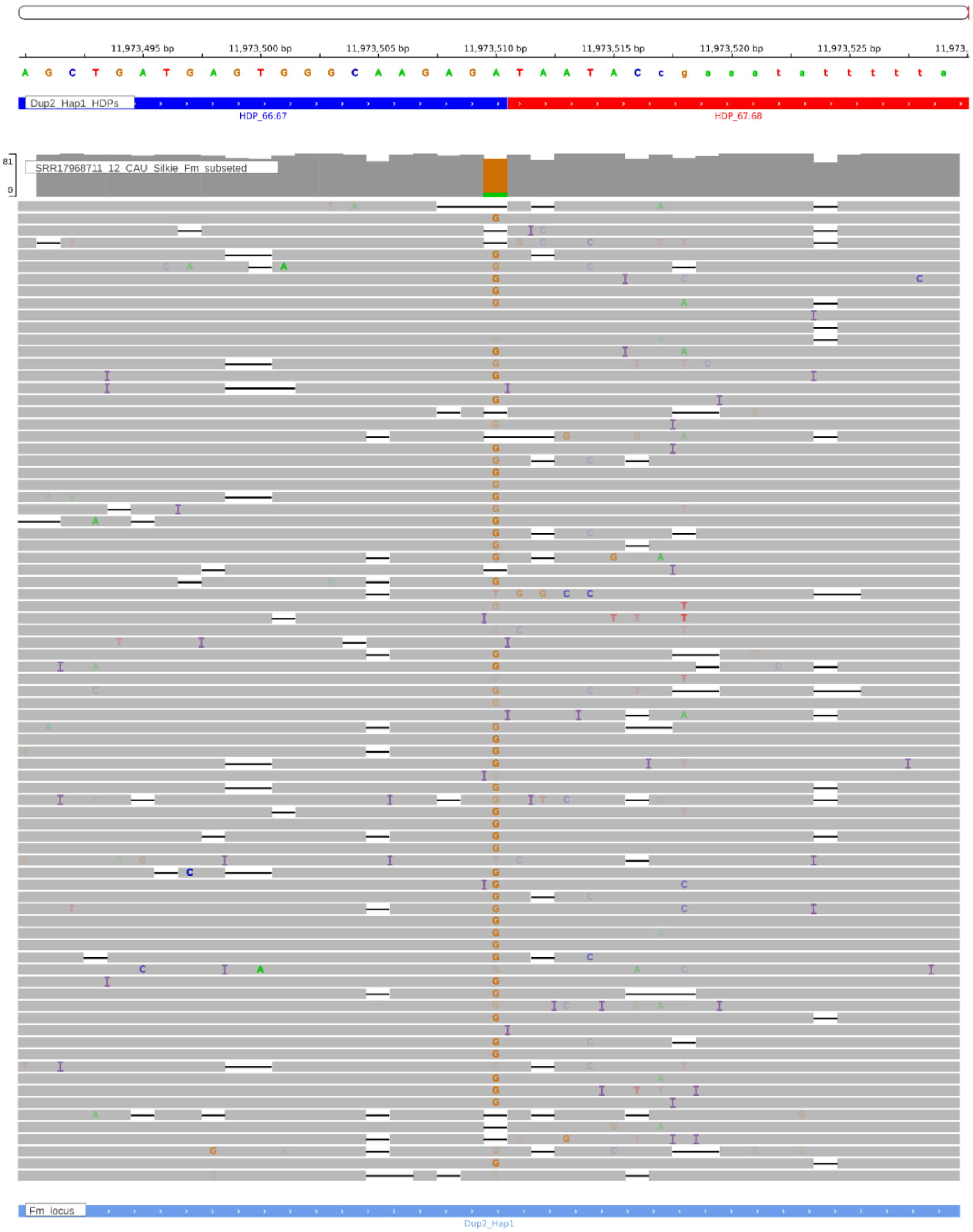


**Supplementary Figure. S150:** Screenshot from IGV showing the HDP 11973510 within the Dup2 region of the CAU Silkie genome. The CAU Silkie ONT long-reads were aligned to the CAU Silkie genome using minimap2, following the settings described by Zhu et al. The alignment reveals that support for the HDP within the genome is notably poor, indicating a mosaic genome assembly due to haplotype switching. The genomic base assembled at the given HDP position actually belongs to the Dup2r haplotype.

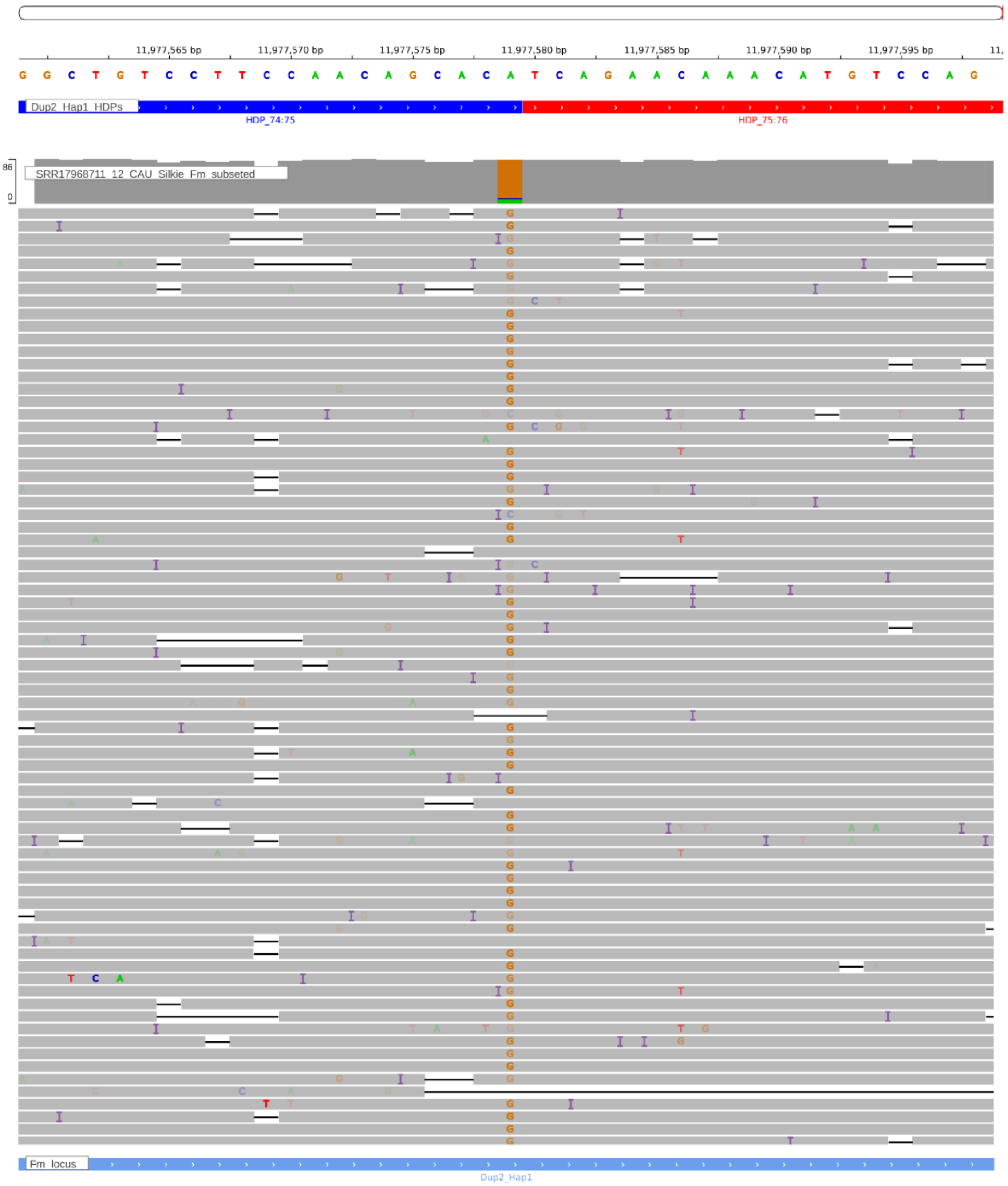


**Supplementary Figure. S151:** Screenshot from IGV showing the HDP 11977579 within the Dup2 region of the CAU Silkie genome. The CAU Silkie ONT long-reads were aligned to the CAU Silkie genome using minimap2, following the settings described by Zhu et al. The alignment reveals that support for the HDP within the genome is notably poor, indicating a mosaic genome assembly due to haplotype switching. The genomic base assembled at the given HDP position actually belongs to the Dup2r haplotype.

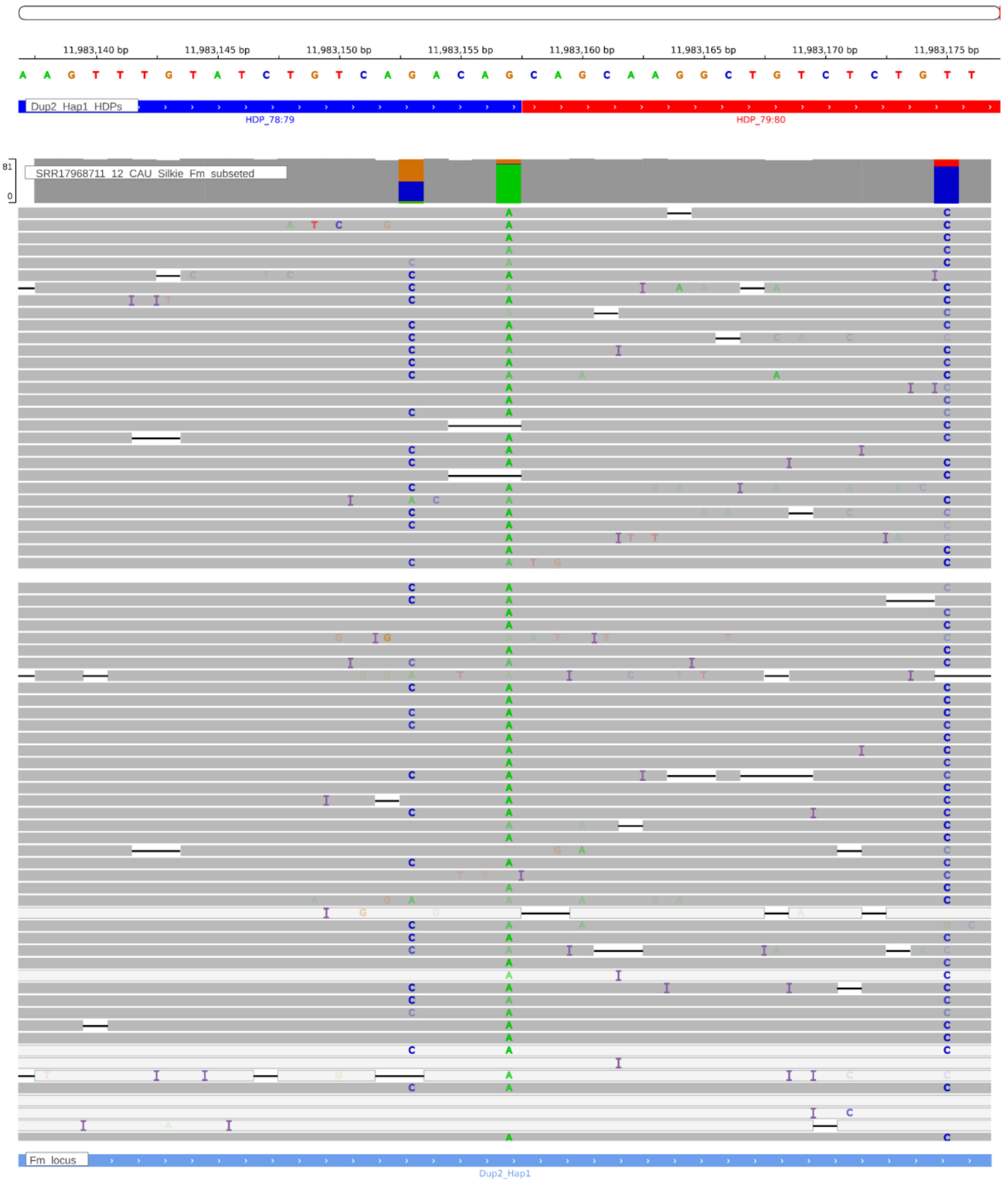


**Supplementary Figure. S152:** Screenshot from IGV showing the HDP 11983157 within the Dup2 region of the CAU Silkie genome. The CAU Silkie ONT long-reads were aligned to the CAU Silkie genome using minimap2, following the settings described by Zhu et al. The alignment reveals that support for the HDP within the genome is notably poor, indicating a mosaic genome assembly due to haplotype switching. The genomic base assembled at the given HDP position actually belongs to the Dup2r haplotype.

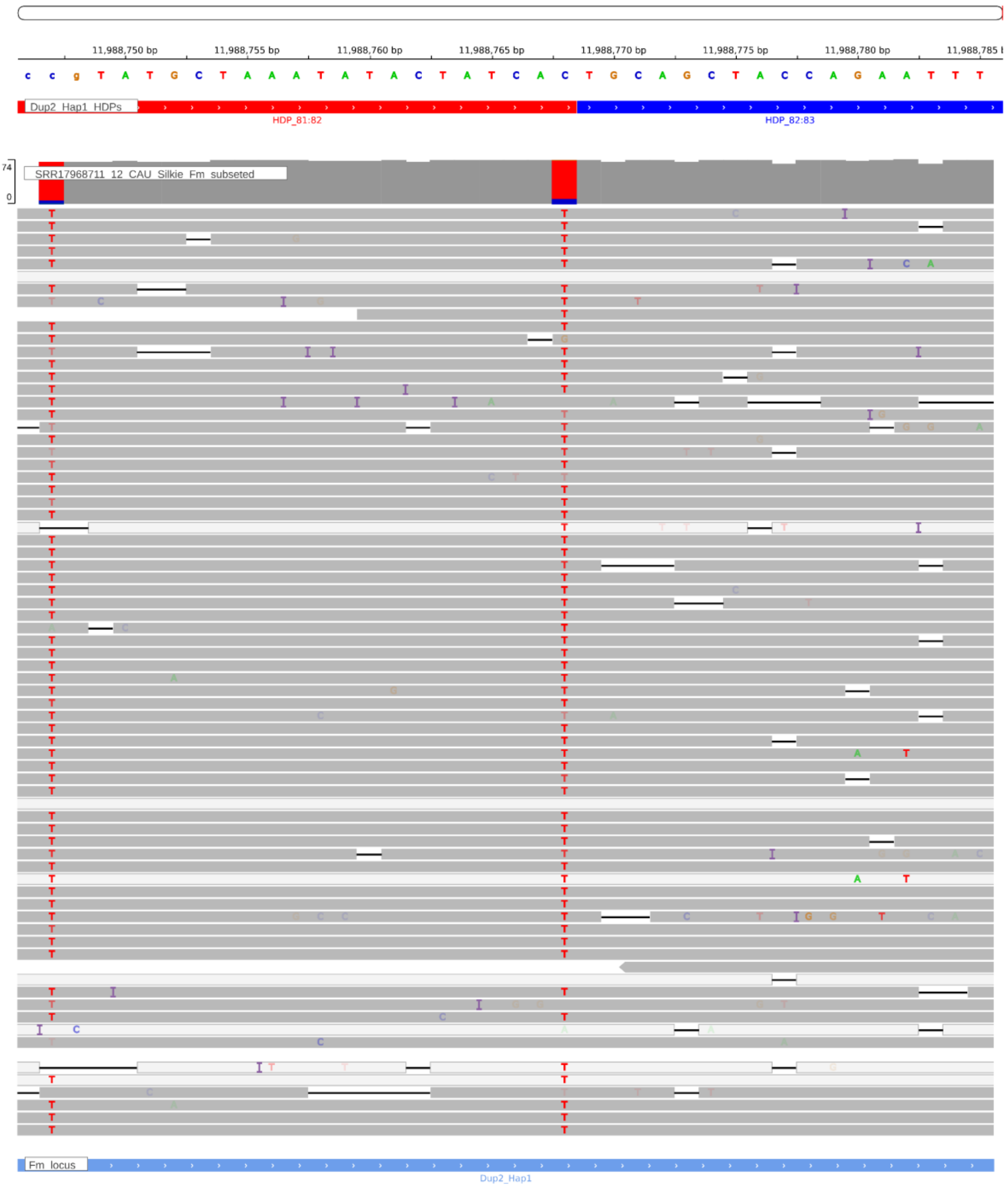


**Supplementary Figure. S153:** Screenshot from IGV showing the HDP 11988768 within the Dup2 region of the CAU Silkie genome. The CAU Silkie ONT long-reads were aligned to the CAU Silkie genome using minimap2, following the settings described by Zhu et al. The alignment reveals that support for the HDP within the genome is notably poor, indicating a mosaic genome assembly due to haplotype switching. The genomic base assembled at the given HDP position actually belongs to the Dup2r haplotype.

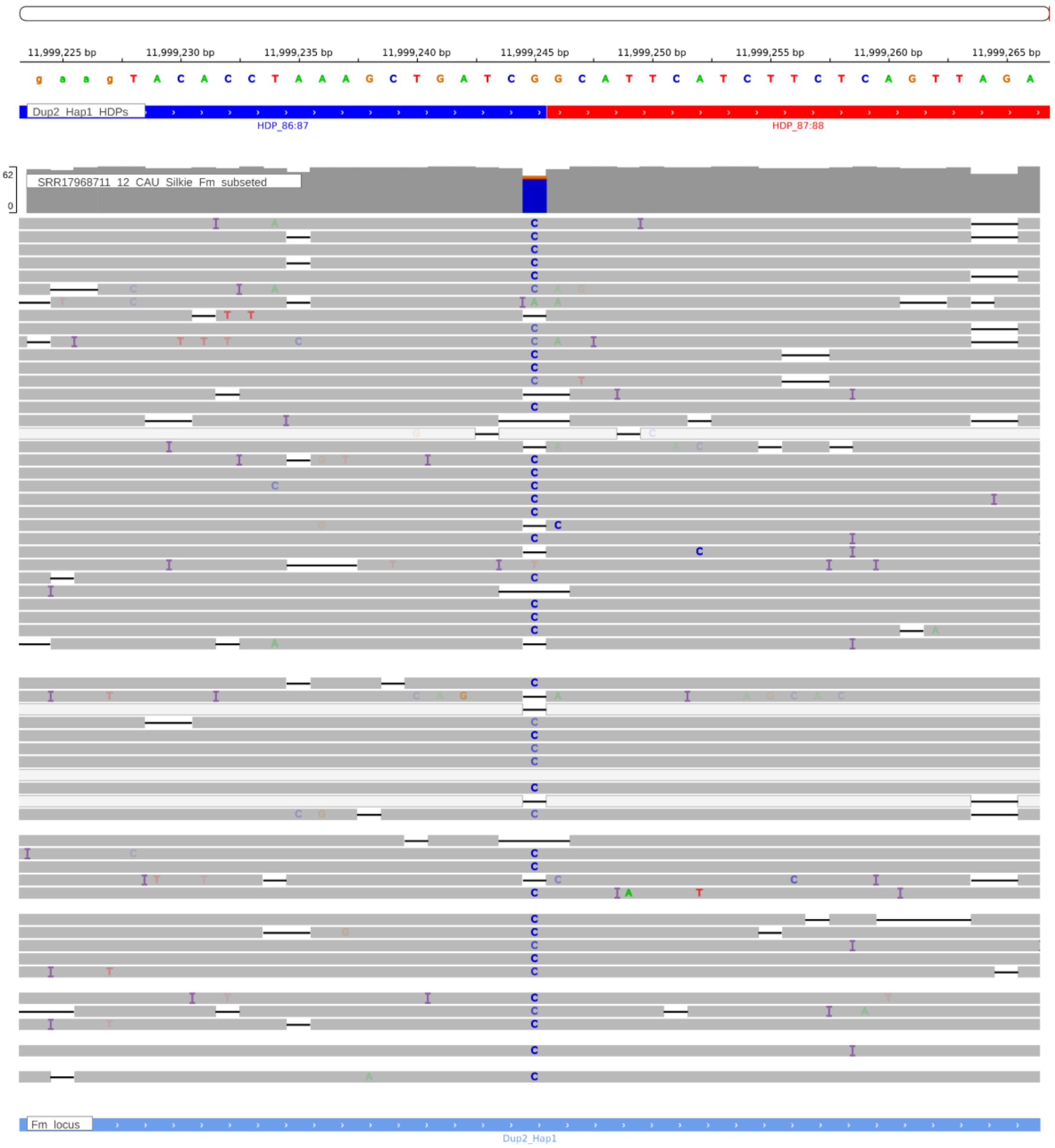


**Supplementary Figure. S154:** Screenshot from IGV showing the HDP 11999245 within the Dup2 region of the CAU Silkie genome. The CAU Silkie ONT long-reads were aligned to the CAU Silkie genome using minimap2, following the settings described by Zhu et al. The alignment reveals that support for the HDP within the genome is notably poor, indicating a mosaic genome assembly due to haplotype switching. The genomic base assembled at the given HDP position actually belongs to the Dup2r haplotype.

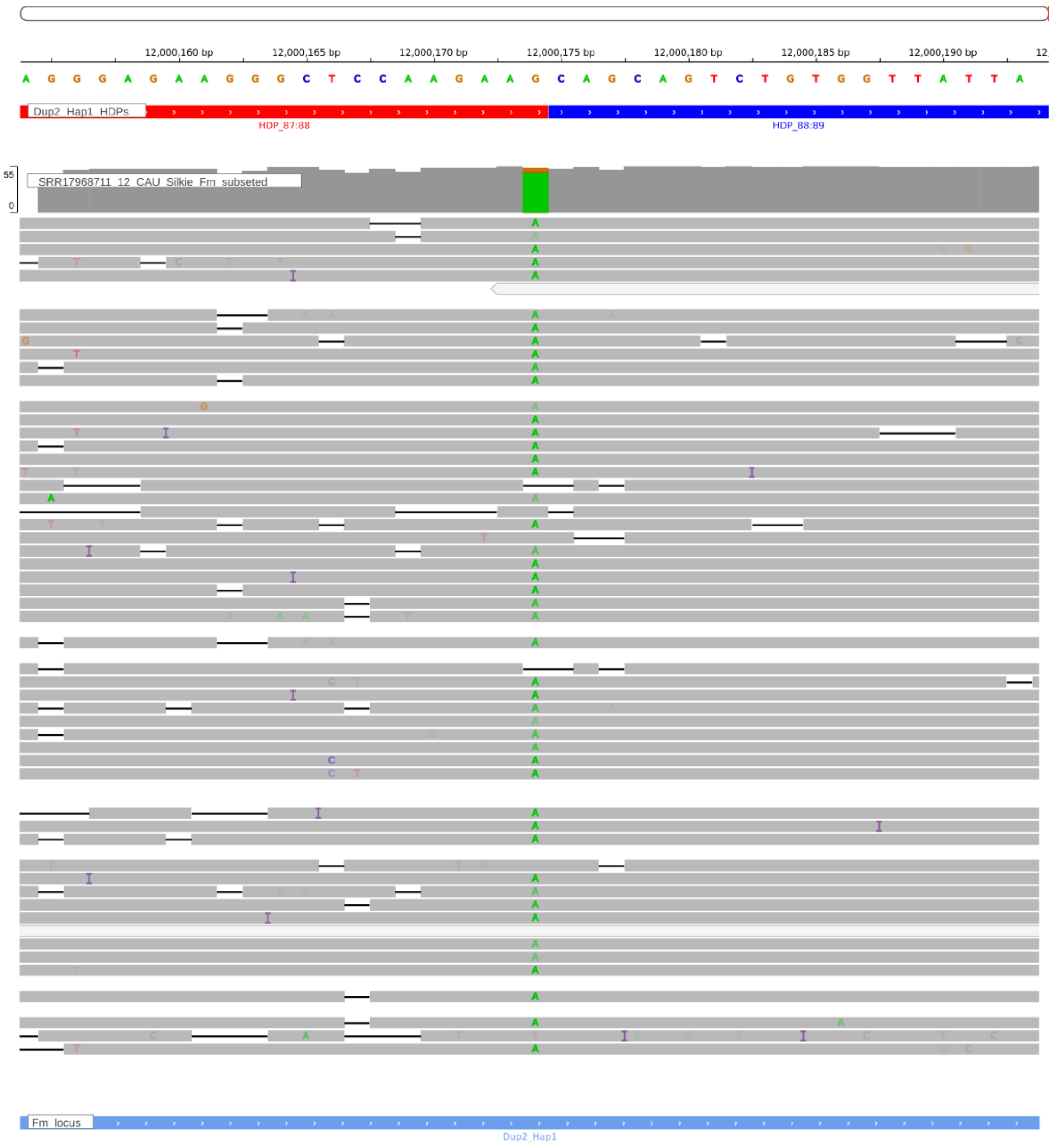


**Supplementary Figure. S155:** Screenshot from IGV showing the HDP 12000174 within the Dup2 region of the CAU Silkie genome. The CAU Silkie ONT long-reads were aligned to the CAU Silkie genome using minimap2, following the settings described by Zhu et al. The alignment reveals that support for the HDP within the genome is notably poor, indicating a mosaic genome assembly due to haplotype switching. The genomic base assembled at the given HDP position actually belongs to the Dup2r haplotype.

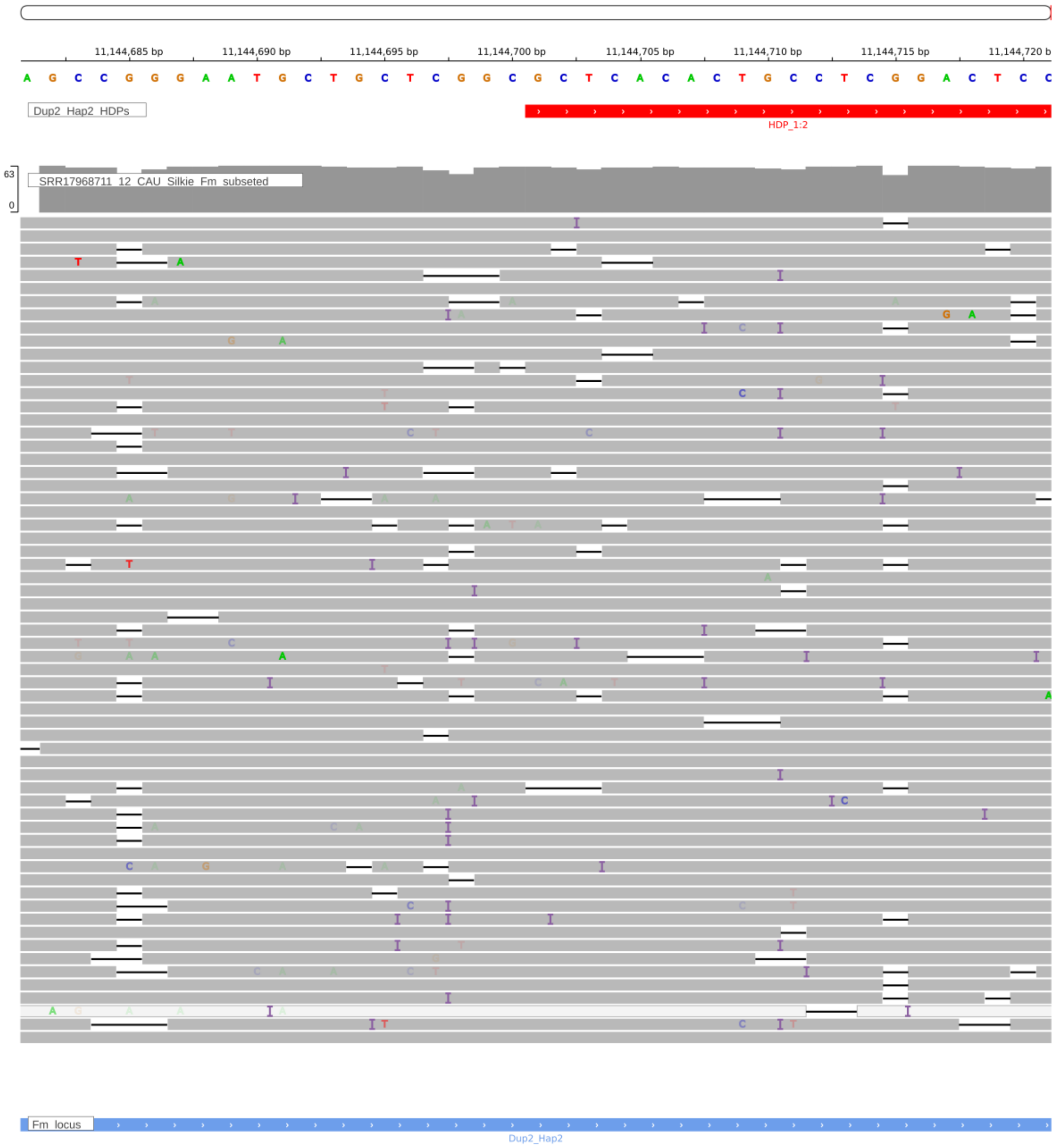


**Supplementary Figure. S156:** The IGV screenshot displays Dup2r HDP 11144700 correctly assembled in the CAU Silkie genome, supported by raw reads from a single haplotype mapped to the genome.

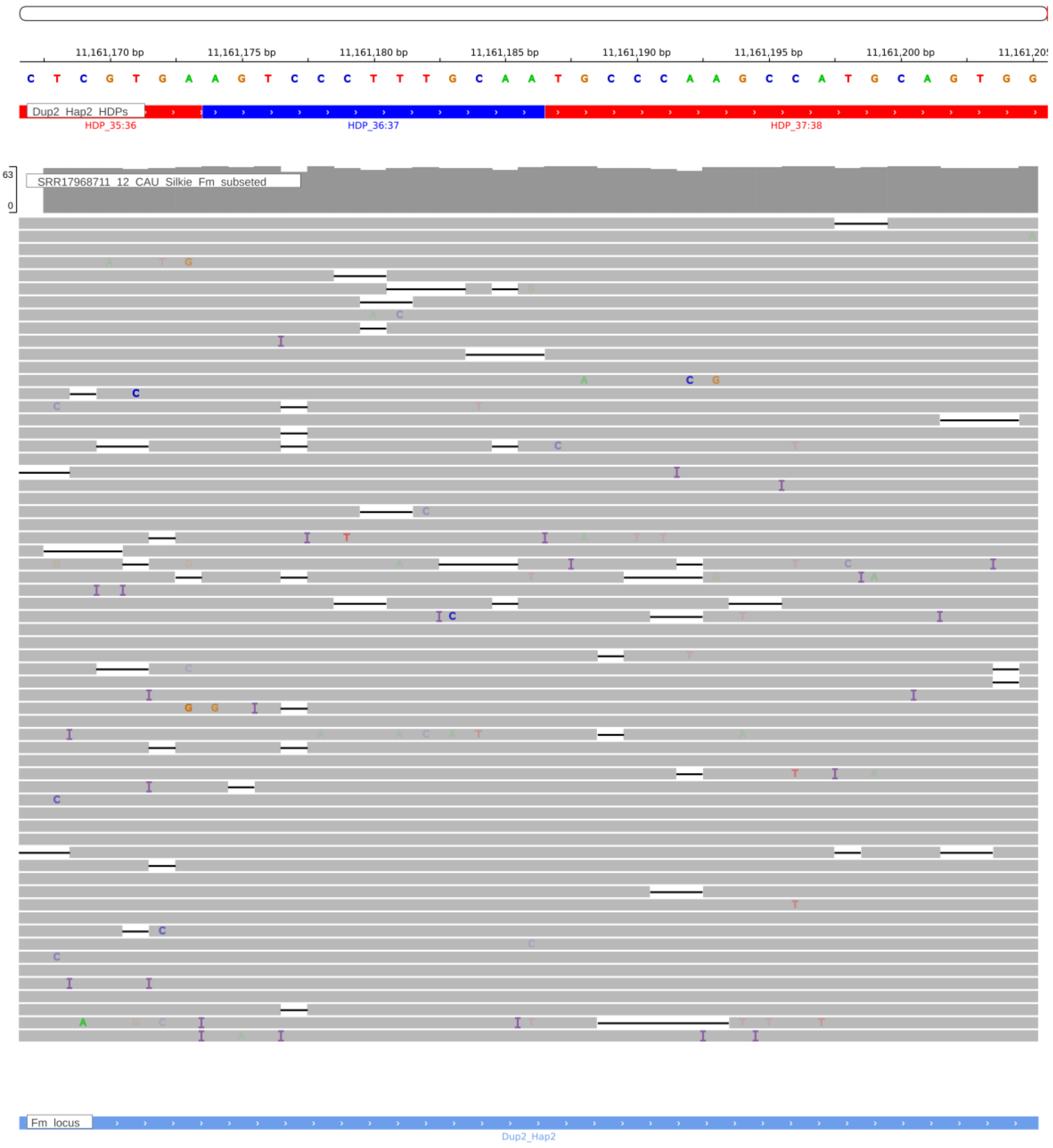


**Supplementary Figure. S157:** The IGV screenshot displays Dup2r HDP 11161186 correctly assembled in the CAU Silkie genome, supported by raw reads from a single haplotype mapped to the genome.

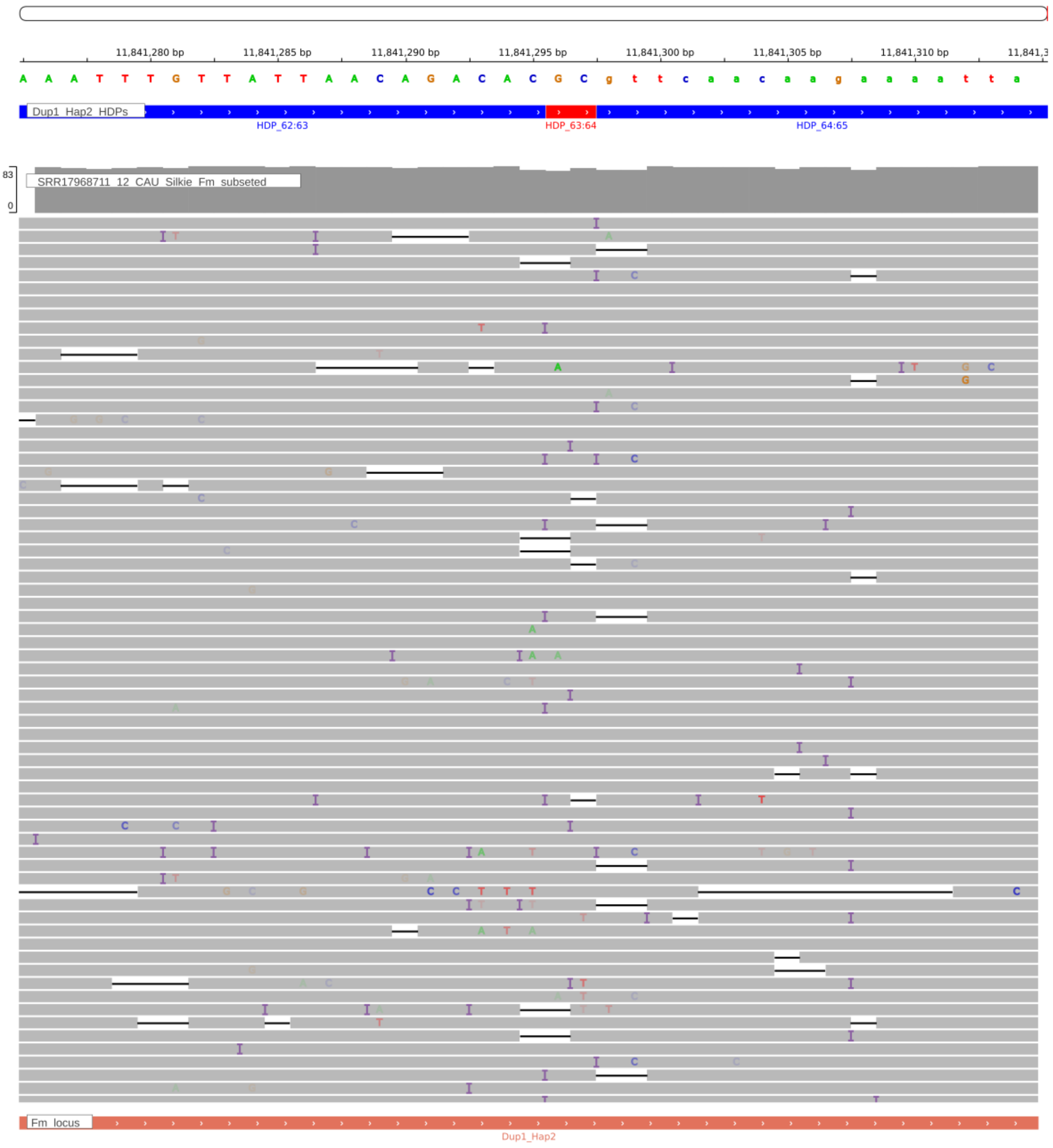


**Supplementary Figure. S158:** The IGV screenshot displays Dup1r HDP 11841295 correctly assembled in the CAU Silkie genome, supported by raw reads from a single haplotype mapped to the genome.

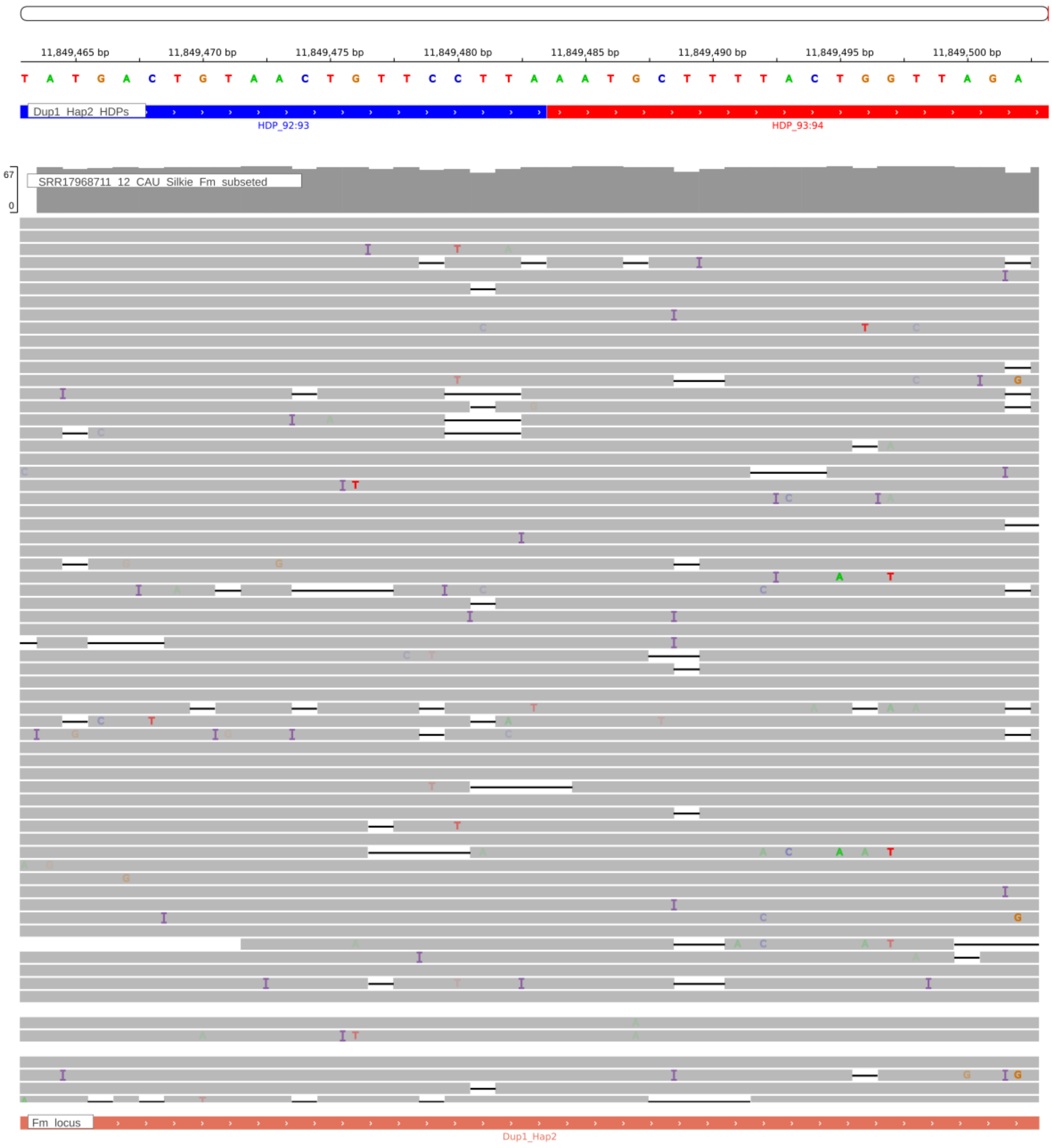


**Supplementary Figure. S159:** The IGV screenshot displays Dup1r HDP 11849483 correctly assembled in the CAU Silkie genome, supported by raw reads from a single haplotype mapped to the genome.

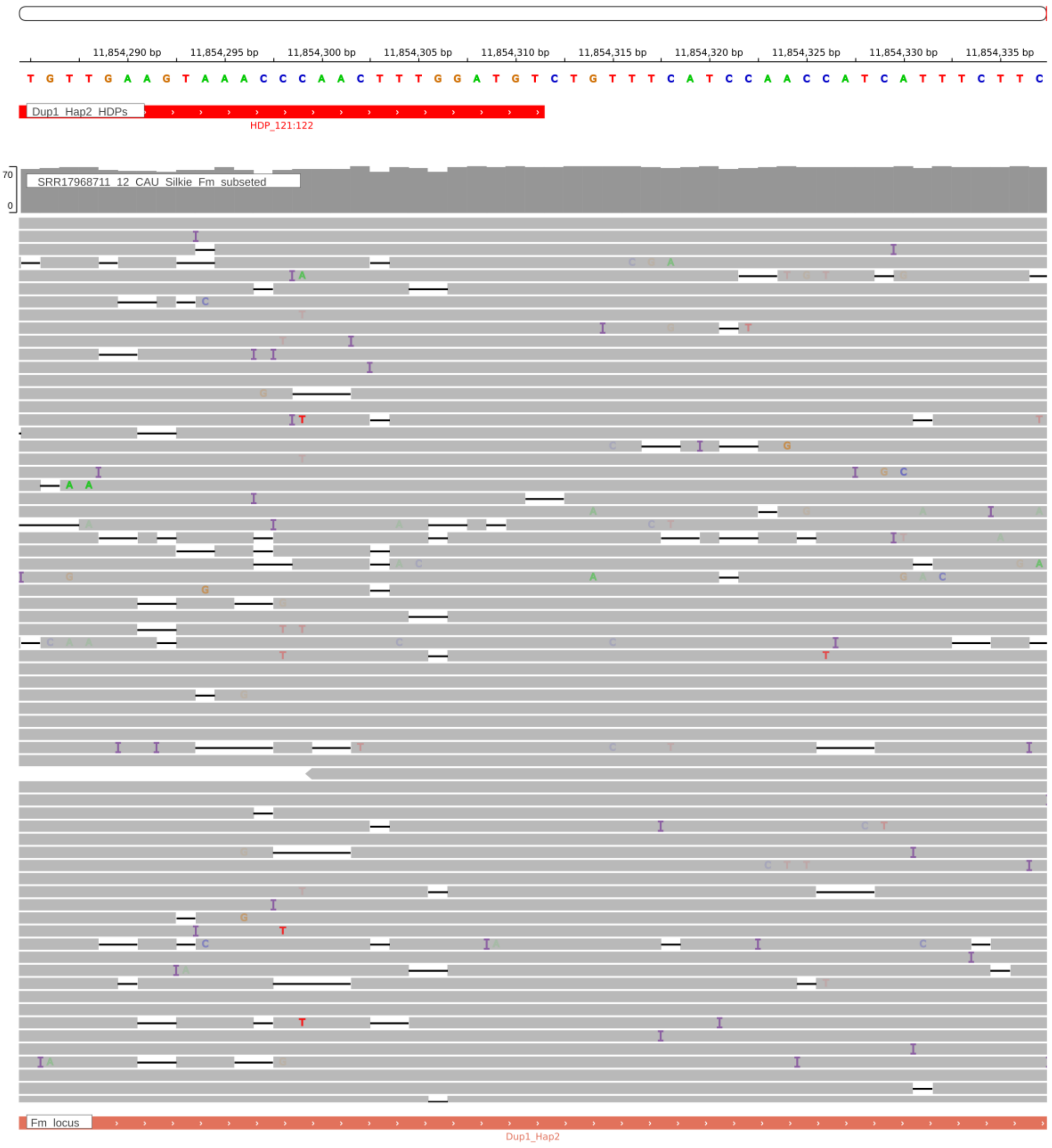


**Supplementary Figure. S160:** The IGV screenshot displays Dup1r HDP 11854311 correctly assembled in the CAU Silkie genome, supported by raw reads from a single haplotype mapped to the genome.

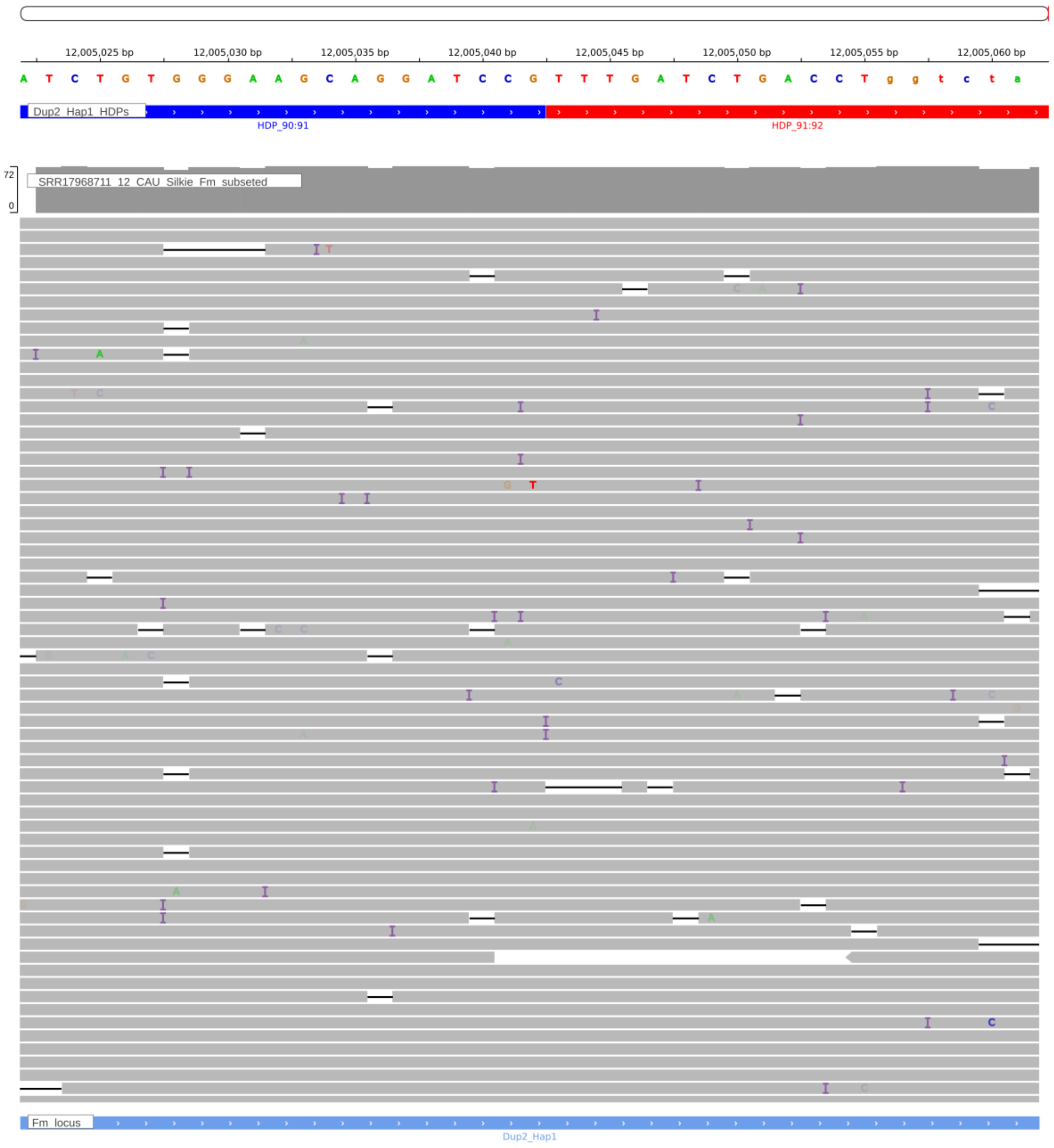


**Supplementary Figure. S161:** The IGV screenshot displays Dup2 HDP 12005042 correctly assembled in the CAU Silkie genome, supported by raw reads from a single haplotype mapped to the genome.

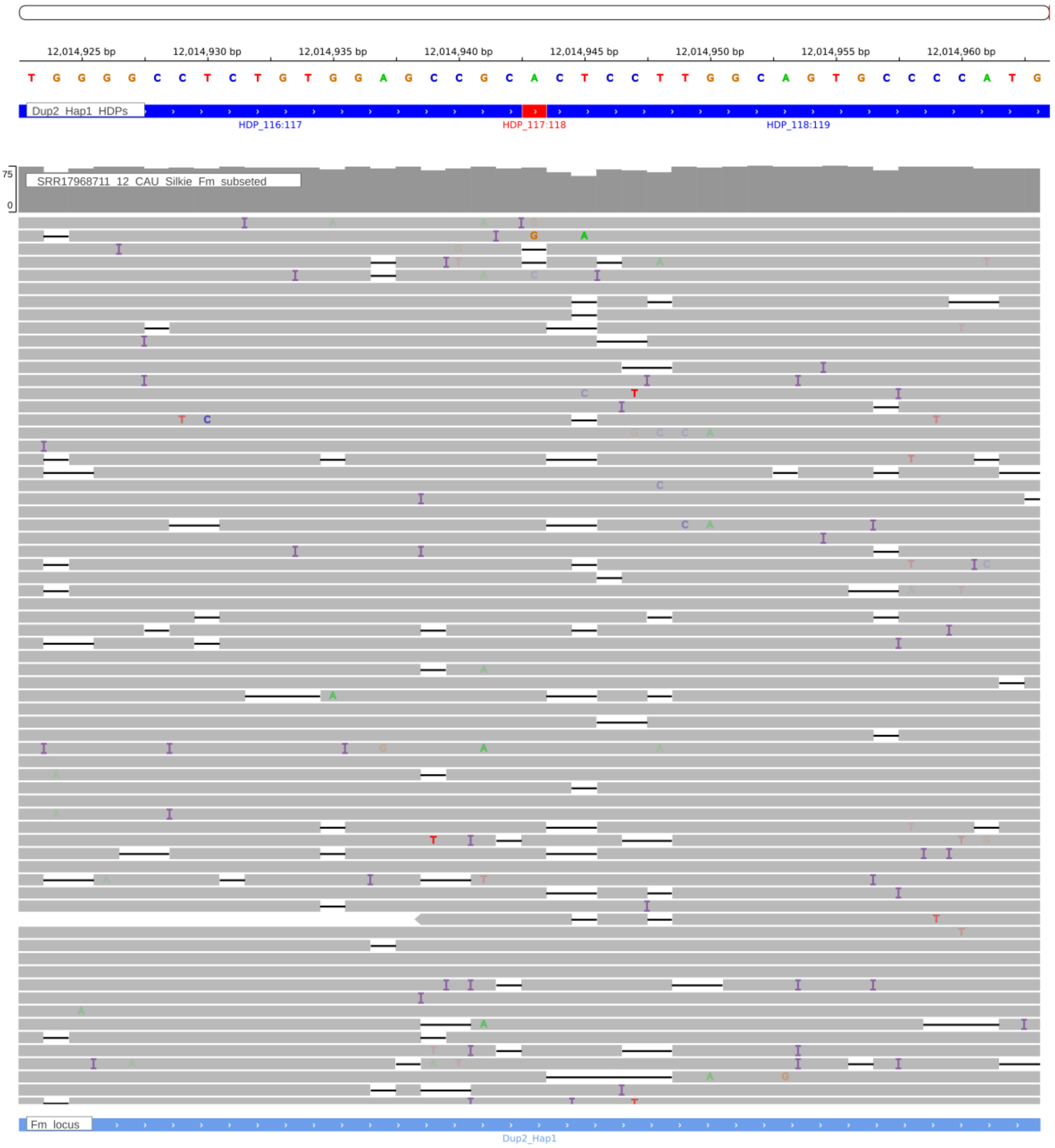


**Supplementary Figure. S162:** The IGV screenshot displays Dup2 HDP 12014943 correctly assembled in the CAU Silkie genome, supported by raw reads from a single haplotype mapped to the genome.

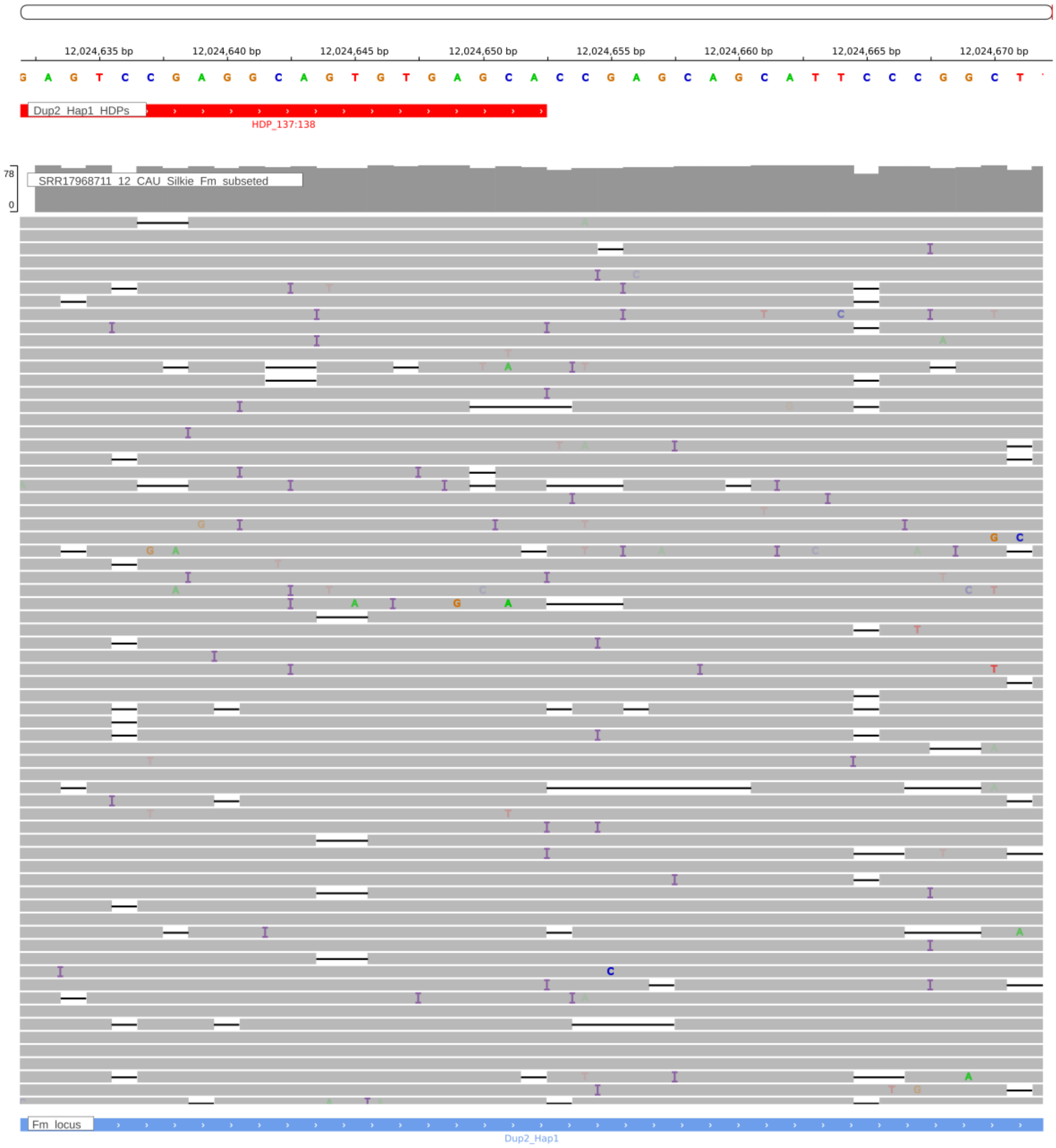


**Supplementary Figure. S163:** The IGV screenshot displays Dup2 HDP 12024652 correctly assembled in the CAU Silkie genome, supported by raw reads from a single haplotype mapped to the genome.
